# Sub-stoichiometric modifications of *Aplysia californica* tRNAs and tRNA fragments revealed by integrating intact and bottom-up mass spectrometry

**DOI:** 10.64898/2026.09.27.754811

**Authors:** Weichen Huang, William M. McGee, Erika Stark, Erica Wang, Kevin D. Clark

**Affiliations:** Department of Chemistry, Tufts University, Medford, MA, 02155, USA; Waters Corporation, Cambridge, MA, 02142, USA

**Keywords:** tRNA, mass spectrometry, RNA modification, tRNA halves, neuro-epitranscriptome

## Abstract

Transfer RNA (tRNA) modifications regulate protein synthesis, yet their sequence positions and stoichiometries are known only for a subset of tRNAs in model organisms. *Aplysia californica* is a key neurobiological model in which neuronal modified ribonucleosides have been linked to changes in animal behavior. However, *Aplysia* tRNA modifications have not been mapped, obscuring their roles in modulating neuronal functions. To address this knowledge gap, we leveraged hybridization pulldowns and an integrated mass spectrometry (MS) strategy combining nucleoside profiling, intact tRNA measurements, and bottom-up tRNA modification mapping to determine the full sequences of *Aplysia* tRNA^Glu^CUC and tRNA^Lys^UUU. Sub-stoichiometric modifications including dihydrouridine (D), 5-methylcytidine (m^5^C), 5,2’-O-dimethyluridine (m^5^Um) and 1-methyladenosine (m^1^A) on tRNA^Glu^CUC varied across tissues, suggesting tRNA modifications are regulated to accommodate translational needs. In addition to conserved thio-modifications in the anticodon loop of tRNA^Lys^UUU, we report 2-thiocytidine (s^2^C) at position 32, representing the first detection of s^2^C in eukaryotes. We also developed a generalizable method to directly detect tRNA halves by intact LC-MS, revealing endogenous tRNA cleavage patterns. Our findings illustrate the advantages of combining intact and bottom-up MS, reveal s^2^C as an unexpected tRNA modification in a classical neurobiological model and offer new insights into post-transcriptional regulation of animal physiology.

Graphical abstract

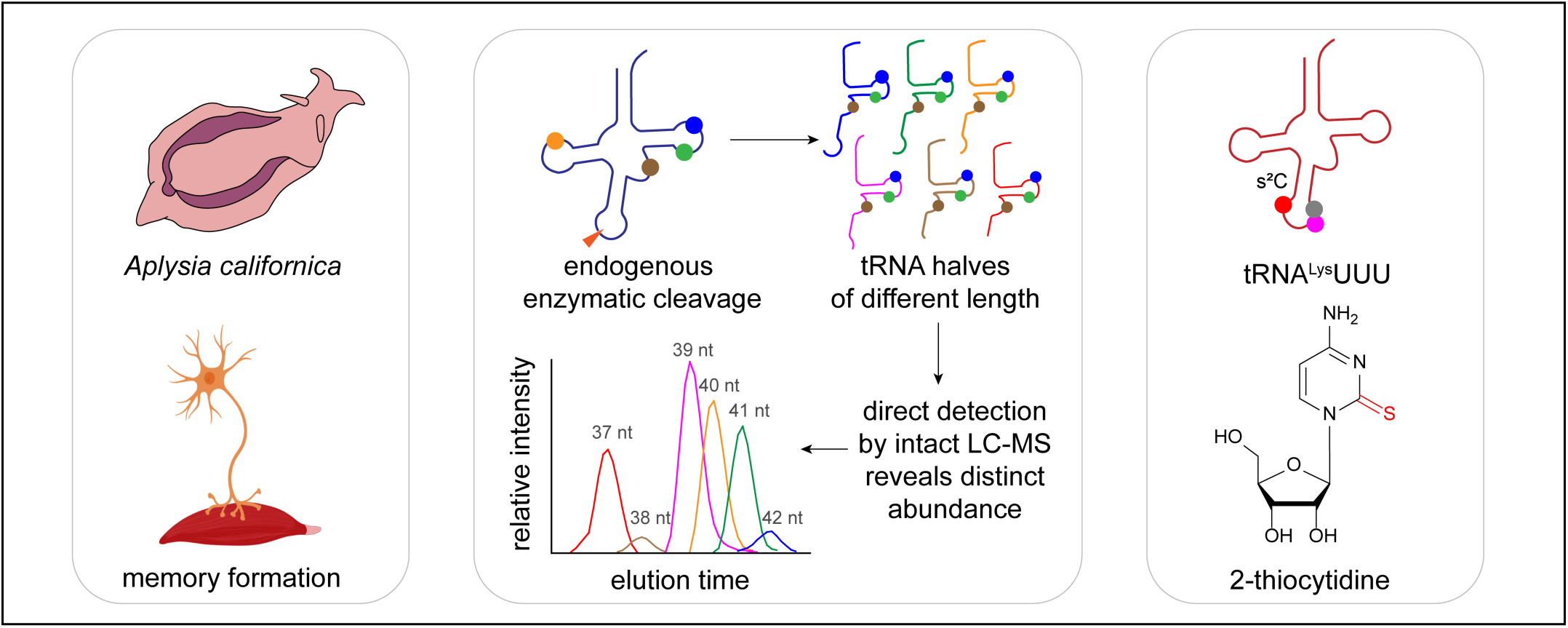

## Introduction

Transfer RNAs (tRNAs) function as adapter molecules that couple mRNA codons to their corresponding amino acids during protein synthesis, shaping the proteome to support fundamental cell functions. In response to endogenous signals and external stimuli, the tRNA population is dynamically regulated at the transcriptional level so that the relative abundance of different tRNAs can accommodate codon-biased translation, prioritizing production of certain proteins. Apart from expression levels, post-transcriptional modifications of tRNAs are increasingly recognized for their critical roles in regulating translation. More than 170 unique RNA modifications including methylation, thiolation, and transglycosylation have been reported to date (1), with more than 80% of known modifications found exclusively in tRNA molecules. Modifications fine-tune tRNA structure, stability, cellular localization, and interaction with proteins. In the brain, tRNAs and their modifications are linked to learning and memory, neurodevelopment, and neurodegenerative disorders (2–4). Beyond directly modulating transcription through decoding mRNA codons, tRNAs are cleaved into tRNA-derived fragments, which are involved in central nervous system (CNS) function by mediating the formation of stress granules and G-quadruplex structures (5–8). Nonetheless, due to the complexity of CNS in vertebrate animal models, connecting tRNA modification dynamics to biological functions or specific phenotypes remains a major challenge.

*Aplysia californica* is a marine mollusk and classical model for neurophysiology whose numerically simple CNS is composed of nine ganglia with ∼10,000 neurons (9), a relatively small number compared to the billions of cells in the human brain. Many *Aplysia* neurons are easily identified by their size (up to 1 mm in diameter) and characteristic positions in the CNS, allowing researchers to link specific neurons to biological functions and simple behaviors. *Aplysia* exhibit both non-associative and associative forms of learning, and have thus been a key model leveraged to elucidate numerous, highly conserved mechanisms of learning and memory including synaptic plasticity, neuronal transmission and gene expression (10–13).

Recent technical advancements have enabled discoveries linking the neuronal epitranscriptome to specific functions and behaviors in *Aplysia*. Optimized LC-MS/MS approaches have been developed to profile spatiotemporal patterns for dozens of neuronal RNA modifications in *Aplysia* (14–17), revealing characteristic tRNA modification dynamics during behavioral sensitization and habituation. Notably, levels of 1-methyladenosine (m^1^A) and 5-methoxycarbonylmethyl-2-thiouridine (mcm^5^s^2^U) were elevated in sensitized animals compared to naïve animals, which facilitated polyglutamine protein synthesis, increased neuron excitability, and ultimately led to behavioral change. Although m¹A and mcm⁵s²U have been mapped to tRNAs in other organisms and have important roles in modification hierarchy (18, 19) and codon-biased modulation of translation efficiency (20–22), no sequence information for any tRNA modification has been reported in *Aplysia*. Overcoming this knowledge gap would enable the identification of specific tRNAs involved in regulating animal behavior, reveal the functional importance of tRNA modification stoichiometries, and lay the foundation for mechanistic studies of how tRNA modifications influence neurophysiology.

Here, we developed an integrated mass spectrometry strategy to obtain the complete sequence of select *Aplysia* tRNAs predicted to bear learning-related modifications. By combining hybridization pulldowns, nucleosides analysis, intact tRNA analysis by liquid chromatography-mass spectrometry (LC-MS), and bottom-up modification mapping by LC-tandem mass spectrometry (LC-MS/MS) with parallel nuclease digestions, we determined the complete sequences of *Aplysia* tRNA^Glu^CUC and tRNA^Lys^UUU inclusive of modifications. Using this comprehensive approach, we detected 2-thiocytidine (s^2^C) at position 32 of *Aplysia* tRNA^Lys^UUU, which to our knowledge represents the first detection of s^2^C in a eukaryotic tRNA. mcm^5^s^2^U and ms^2^t^6^A were also mapped to the anticodon loop of tRNA^Lys^UUU, suggesting coordinated functions in codon recognition. tRNA^Glu^CUC displayed sub-stoichiometric modifications including dihydrouridine (D), 5-methylcytidine (m^5^C), 5,2’-O-dimethyluridine (m^5^Um) and m^1^A, some of which showed tissue-specific abundances. Intact LC-MS analysis of hybridization pulldowns revealed co-purified tRNA^Glu^CUC halves of different lengths that displayed the same modifications and stoichiometry profiles as those on full length tRNA^Glu^CUC. Differential abundances of these fragments suggested sequence preferences for endogenous tRNA cleavage. Altogether, this study reports a generalizable MS workflow for sequencing and quantifying tRNA modifications, expands the eukaryotic tRNA modification repertoire to include s^2^C, and reveals sub-stoichiometric modification patterns of *Aplysia* tRNAs and distinct abundances of endogenous tRNA halves. These findings lay the groundwork to fully leverage the *Aplysia* model for investigating how modifications of tRNAs and tRNA halves contribute to fundamental neuronal functions and animal behaviors.

## Materials and methods

### Safety statement

No unexpected or unusually high safety hazards were encountered.

### Codon frequency analysis of *Aplysia californica* RNA-seq data

Codon frequency analysis was conducted using the web-based platform Galaxy (https://usegalaxy.org/). Bulk-tissue RNA-seq datasets from *Aplysia* neurons and other tissues (Supplementary Material 1: Table S1) were *de novo* assembled with rnaSPAdes 4.3.0 (23), and codon frequency statistics were obtained from assembled mRNA transcripts by TransDecoder 5.5.0 (24) and EMBOSS cusp 5.0.0 (25) with nuclear genome genetic code, thus excluding mitochondria genome encoded mRNAs. Codon frequency statistics were analyzed and visualized by Microsoft Excel.

### *Aplysia californica* tissue isolation and total RNA extraction

Animals weighing 100–250 g were obtained from the National Resource for *Aplysia* (Miami, FL, USA) and housed in an aquarium with circulating, aerated artificial seawater (ASW; Instant Ocean, Blacksburg, VA, USA) at 14 °C for a minimum of 48 h prior to dissection. Animals were anesthetized by injections with 0.33 M MgCl_2_ (approximately 40% body volume) directly into the body cavity. Organs/tissues were surgically isolated, fast frozen by liquid nitrogen and homogenized by a mortar and pestle to a fine powder. Total RNA was extracted by liquid phase extraction with RNAzol (Sigma-Aldrich) following manufacturer’s instructions, and small RNA was fractionated by solid phase extraction using Monarch^®^ Spin RNA Cleanup Kit (New England Biolabs).

### DNA probe-pulldown of tRNA^Glu^CUC and tRNA^Lys^UUU

*Aplysia* tRNA^Glu^CUC and tRNA^Lys^UUU were isolated from small RNA fractions using biotinylated complementary DNA probes (IDT) and streptavidin-functionalized Sepharose beads (Cytiva). First, 100 μL small RNA (∼400-500 ng/μL), 4 μL DNA probe (10 μM), 2.6 μL 20× SSC buffer (3 M sodium chloride, 300 mM trisodium citrate, pH adjusted to 7.0 with HCl), and 1 μL RNase inhibitor (New England Biolabs) were combined in a 200 μL PCR tube and incubated at 50 °C for 30 min. A 100 μL aliquot of the streptavidin-functionalized beads suspension was washed twice with 1 mL 0.5× SSC buffer by gentle trituration, centrifuged at 8000*g* for 1 min, and the supernatant discarded. Next, the RNA: DNA probe mixture was combined with the beads and incubated at room temperature for 15 min. The mixture was then transferred to a 1.5 mL Ultrafree-MC tube (0.2 μm, Sigma-Aldrich), washed five times with 300 μL 0.5× SSC buffer by centrifugation at 5000*g* for 1 min, and flowthrough was discarded. Then, 100 μL of RNase-free water was added to the filter cartridge and incubated in a heat block at 75 °C for 15 min, then centrifuged at 5000*g* for 1 min. The flowthrough containing the target tRNA was collected. The final yield of tRNA^Glu^CUC or tRNA^Lys^UUU varied among different tissues, ranging from 500 to 1,200 ng tRNA (20∼50 picomole) per 500 mg of fresh tissue.

### Denaturing polyacrylamide gel electrophoresis

For polyacrylamide gel electrophoresis (PAGE), 10% denaturing polyacrylamide gels were prepared with 4.8 g urea dissolved in 2.5 mL 40% polyacrylamide/bis solution (Bio-Rad) combined with 1.0 mL 10× Tris/Borate/EDTA (TBE) buffer and 6.5 mL water. Gels were polymerized by 100 μL 10% ammonium persulfate and 15 μL tetramethylethylenediamine. For each gel, 0.5 μL low range ssRNA ladder was mixed with 0.5 μL of RNA loading dye (New England Biolabs) and 1.0 μL formamide (Thermo Fisher). tRNA samples were mixed with 1× volume of formamide and 1.0 μL RNA loading dye. RNA ladders and tRNA samples were denatured at 85 °C for 5 min, chilled on ice for 1 min, and loaded on gel. The gel was electrophoresed at 120 V for 40 min in 1× TBE buffer, stained with SYBR Green II (Thermo Fisher) for 10 min at room temperature, and visualized on a transilluminator.

### Enzymatic digestion of tRNA to nucleosides and analysis by LC-MS/MS

tRNA^Glu^CUC and tRNA^Lys^UUU samples were subjected to a two-step digestion to ribonucleosides using a combination of nuclease P1, snake venom phosphodiesterase and bacterial alkaline phosphatase (Fisher Scientific), following the method developed by Crain et al. (26). tRNA samples were incubated with nuclease P1 in 10 mM ammonium acetate at 45 °C for 2 h, followed by addition of snake venom phosphodiesterase and bacterial alkaline phosphatase and a second incubation in 100 mM ammonium bicarbonate at 37 °C for 2 h. For each sample, ∼250 ng of tRNA were digested in a final volume of 40 μL.

tRNA nucleoside digests were analyzed using an Agilent 1260 Infinity II HPLC system coupled to an Agilent QTOF 6530B with Dual AJS ESI operated in positive mode. The LC autosampler was kept at 4 °C to minimize sample degradation. LC separations were conducted using a ZORBAX Eclipse Plus C18 Narrow Bore RR 2.1 x 100 mm, 3.5 µm column (Agilent) and gradient elution with mobile phase A consisting of 5 mM ammonium acetate (pH 5.6) and mobile phase B as 60/40 mobile phase A/acetonitrile. Gradient elution was performed using a flow rate of 0.2 mL/min at 36 °C and the following conditions: 0–5 min 1% B, 7% B at 11 min, 10% B at 13 min, 15% B at 32 min, 70% B at 40 min, 100% B at 44 min, 100% B at 50 min, 1% B at 50.1 min, and a 10-min hold at 1% B prior to the start of the next injection. LC flow was directed to the mass spectrometer after diverting to waste for 1.5 min. The capillary voltage was 3 kV and the N_2_ drying gas was set to 5 L/min and 300 °C. Fragmentor voltage was 100 V. For MS1, a scan range of *m/z* 200–600 was used. Automatic MS/MS mode was used with a scan range of *m/z* 80-600, and collision energies were applied using a linear relationship between the following two points: 10 V for *m/z* 100, z=1 and 50 V for *m/z* 1,000, z=1. Additional LC-MS/MS method details are provided in Supplementary Material 1: Table S6 and Table S7. The identities of modified nucleosides were determined by expected monoisotopic mass values corresponding to protonated nucleosides ([M+H]^+^), by comparing the major MS/MS fragment ions to those reported in the MODOMICS database (1), and by LC retention times and selectivity which are characteristic for reversed phase chromatography on C18 columns (1, 27, 28).

### Fitting analysis for mass spectrum of s^2^C

For isotopic envelope fitting analysis of s^2^C, the theoretical electrospray ionization (ESI) mass spectrum for the [M+H] ion was simulated with Prot pi (https://www.protpi.ch/Calculator/MassSpecSimulator) to produce a distribution of theoretical mass-to-charge ratios (*m/z*) and corresponding probabilities. The approach used here was adapted from previous reports (28, 29). Briefly, *m/z* values matching [M+H], [M+H]+1 and +2 Da isotope peaks were picked to represent peaks in centroid data, and their corresponding probability values were collected. The observed *m/z* values and intensities were collected from *Aplisya* tRNA^Lys^UUU nucleoside samples from muscle. The theoretical and observed values were paired as 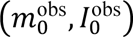, 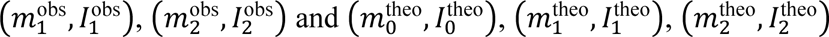, where m is for *m/z* values and *I* is for intensities. Intensity vectors for observed and theoretical envelopes were defined as 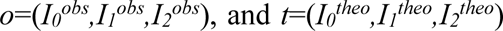, respectively. The intensity vectors are normalized by 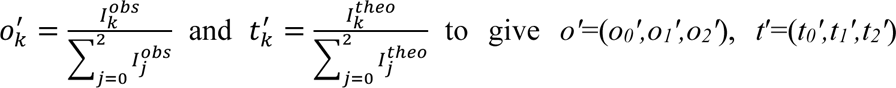. Finally, the cosine similarity is calculated by 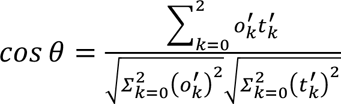, and a *cos θ* value closer to 1 indicates high similarity of the observed to theoretical isotopic envelope.

### RNase T1 and RNase 4 digestion of tRNA for bottom-up sequencing by LC-MS/MS

For RNase T1 digestion, 500-600 ng tRNA was digested with 150 U of RNase T1 (Thermo Scientific) in nuclease-free water in a total volume of 21.5 μL at 37 °C for 40 min. Next, 0.0125 U bacterial alkaline phosphatase was added and incubated in 0.1 M ammonium bicarbonate (pH 8.0) with a total volume of 24.2 μL at 37 °C for 40 min. For RNase 4 digestion, 500-600 ng tRNA was denatured in 1 M urea at 75 °C for 5 min, then digested with 1 μL RNase 4 and 3’ end repair mix (New England Biolabs) in 1×NEBuffer r1.1 (New England Biolabs) in a total volume of 15 μL at 37 °C for 60 min. Digested samples were loaded into the LC autosampler kept at 4 °C and promptly injected for LC-MS/MS analysis or stored at −80 °C until analysis. tRNA digests were then analyzed using an Agilent 1260 infinity II HPLC system coupled to an Agilent QTOF 6530B with Dual AJS ESI operated in negative mode. LC separations were conducted using a Waters XBridge BEH C18 column with 130 Å pores, 3.5 µm particles, 4.6 mm i.d. × 100 mm. Gradient elution was performed with mobile phase A consisting of 10 mM ammonium acetate (pH 7.0) and mobile phase B as 100% acetonitrile. Gradient elution with a flow rate of 0.300 mL/min at 50 °C was used with the following conditions: 0–10 min 0% B, 10% B at 30 min, 50% B at 35 min, 50% B at 40 min, 0% B at 45 min and a 10-min hold at 0% B prior to the start of the next injection. LC flow was directed to the mass spectrometer after diverting to waste for the initial 1.5 min to avoid introducing nonvolatile salts into the mass spectrometer. For mass spectrometry, a MS1 scan range of *m/z* 300–3200 was used. The capillary voltage was 4.5 kV and the N_2_ drying gas was set to 12 L/min and 350 °C, with fragmentor voltage of 180 V. Automatic MS/MS mode was used with a scan range of *m/z* 100-3200. For collision energies, the following formula was used: E=(slope) × (*m/z*)/100 + offset. For more details about the LC-MS/MS method see Supplementary Material 1: Table S4 and Table S5.

### tRNA modification mapping from MS/MS data

Among organisms for which tRNA^Lys^ modifications have been previously mapped, *Heterololigo bleekeri* (formerly *Loligo bleekeri*, a squid species in the phylum Mollusca) is phylogenetically closest to *Aplysia* and has a tRNA with high sequence similarity to *Aplysia* tRNA^Lys^UUU. Cytoplasmic tRNA^Lys^ sequences of *H. bleekeri* were retrieved from the Modomics database (Supplementary Material 1: Table S9) and used as a guide for mapping modifications of *Aplysia* tRNA^Lys^UUU. No modification information was available for *H. bleekeri* tRNA^Glu^, so tRNA^Glu^CUC sequences from mouse and human were used to predict conserved modifications on *Aplysia* tRNA^Glu^CUC. Based on the expected RNase T1 and RNase 4 digestion fragments (with and without modifications), *m/z* of ions in the electrospray series were simulated by Mongo Oligo Mass Calculator v2.07 (31). Extracted ion chromatograms (EICs) for these ions were plotted using MassHunter Qualitative Analysis 10.0 (Agilent). MS2 spectra for tRNA digestion products were manually annotated and further validated by Pytheas software package (32) with automated MS/MS data matching to RNA sequences. Pytheas outputs, which summarized matched product ions from each RNA precursor (Supplementary Materials 2 and 3), were used to visualize sequence coverage, following the ion type nomenclature defined by McLuckey et al. (33). Product ions from multiple MS/MS scans of the same precursor ions were combined when calculating and visualizing sequence coverage.

### Intact mass analysis of tRNA and tRNA fragments by ion-pairing reversed-phase LC-MS

tRNA pulldown samples suspended in nuclease-free water were analyzed on an I-Class UPLC that had been fully replumbed with MaxPeak High Performance Surfaces to minimize oligonucleotide loss from phosphate adsorption. This was coupled with a Xevo QToF G3 (Waters Corp., Cambridge, MA USA) with an electrospray ionization source operated in negative mode. LC separations were conducted with an Acquity Premier BEH C18 column (1.7 μm, 130 Å, 2.1 × 50 mm; Waters) and gradient elution with mobile phase A consisting of 5 mM dibutylamine (DBA), 25 mM 1,1,1,3,3,3-hexafluoroisopropanol (HFIP) and mobile phase B as 100% acetonitrile. Gradient elution was performed using a flow rate of 200 μL/min at 40 °C and the following conditions: 0–5 min 5% B, 15% B at 6 min, 21% B at 13 min, 23% B at 22 min, 70% B at 24 min, 70% B at 30 min, 5% B at 31 min, 5% B at 35 min. LC flow was directed to waste before 5 min and after 22 min. For MS, a scan range of *m/z* 1,000–50,000 was used. The cone voltage ramps range from 80 to 100 V, with desolvation pressure of 1,200 L/h, source voltage of 2.5 kV, and scan time of 1s. MS1 signal was collected from 5 to 22 min, and LC-MS data was analyzed by MassLynx v4.2 (Waters).

## Results and discussion

tRNA expression and modification stoichiometries are responsive to external stimuli and impact translation of stimulus-specific proteins in a codon-biased fashion (34–41). However, due to the complexity of conventional animal models, very little is known about how tRNA modifications connect to stimulus response at the organismal level (*i.e.*, behavioral change). Studying the stereotyped behaviors and numerically simple CNS of *Aplysia* represents a promising approach to overcome this challenge and elucidate the epitranscriptomic mechanisms that underlie learning, memory, and behavioral change. For example, higher levels of m^1^A and mcm^5^s^2^U in total tRNA from *Aplysia* neurons were linked to altered protein synthesis and neuron excitability during behavioral sensitization (42). To gain deeper insights toward epitranscriptomic regulation of translation in the CNS, these modifications must be mapped to specific tRNA sequences. We thus aimed to sequence *Aplysia* tRNAs, initially focusing on tRNAs that (i) harbor m^1^A and mcm^5^s^2^U modifications due to their previously identified roles in learning and memory and (ii) are highly expressed in *Aplysia* CNS and other tissues to maximize the amount of tRNA recovered from the limited amounts of tissue available from each animal.

### Prediction of *Aplysia* tRNA^Glu^CUC and tRNA^Lys^UUU as highly abundant cytoplasmic tRNAs by codon usage analysis

Eukaryotic genomes often contain hundreds of tRNA genes, of which several dozen are expressed (43). In the nuclear and mitochondrial genomes of *Aplysia*, there are a total of 171 nuclear plus 22 mitochondrial genes that potentially encode tRNAs (44, 45), but the exact number of transcribed tRNA species and their relative abundance across tissues is unknown. We estimated tRNA relative abundance based on predicted codon usage derived from existing RNA-seq data for *Aplysia* mRNAs, because the frequency of mRNA codons and the abundance of their cognate tRNAs are highly coordinated for optimized protein production, as observed in *Escherichia coli*, *Trypanosoma cruzi*, and humans (38, 46–49). To estimate the relative abundance of tRNAs in *Aplysia*, bulk-tissue RNA-seq datasets from *Aplysia* neurons and other tissues (Supplementary Material 1: Table S1) were *de novo* assembled with rnaSPAdes (23), and codon frequency statistics were obtained from assembled mRNA transcripts. GAG, ACA, AAG, AAA, and TCT were the five most frequent codons across the ten neuronal RNA-seq datasets analyzed (Figure 1), and among other tissues investigated, GAG was consistently ranked as the most abundant codon, while the ranking of other codons varied only slightly (Supplementary Material 1: Table S2). We further filtered the high frequency codons to identify cognate tRNAs with learning-related post-transcriptional modifications m^1^A and mcm^5^s^2^U (42). The m^1^A58 modification is highly conserved in the T-loop of multiple tRNAs including tRNA^Glu^CUC (1, 50, 51). We thus selected tRNA^Glu^CUC (URS0000928747_6500), which had no isodecoders, for subsequent modification mapping due to its predicted high expression levels across all tissues. Among tRNAs that harbor mcm^5^s^2^U at the anticodon position 34 (52–54), only tRNA^Lys^UUU was among the tRNAs predicted to be most abundant in CNS tissues. For tRNA^Lys^UUU, there are two isodecoders in the *Aplysia* nuclear genome, and we selected the tRNA encoded by seven copies (URS0000947ED9_6500), anticipating it would be more highly expressed relative to its single-copy isodecoder (55, 56). In sum, the predicted high abundances, expected m^1^A and mcm^5^s^2^U modification statuses, and previously reported links to translation and neuronal activities motivated us to isolate tRNA^Glu^CUC and tRNA^Lys^UUU to sequence the full complement of their modifications.

**Figure 1.**
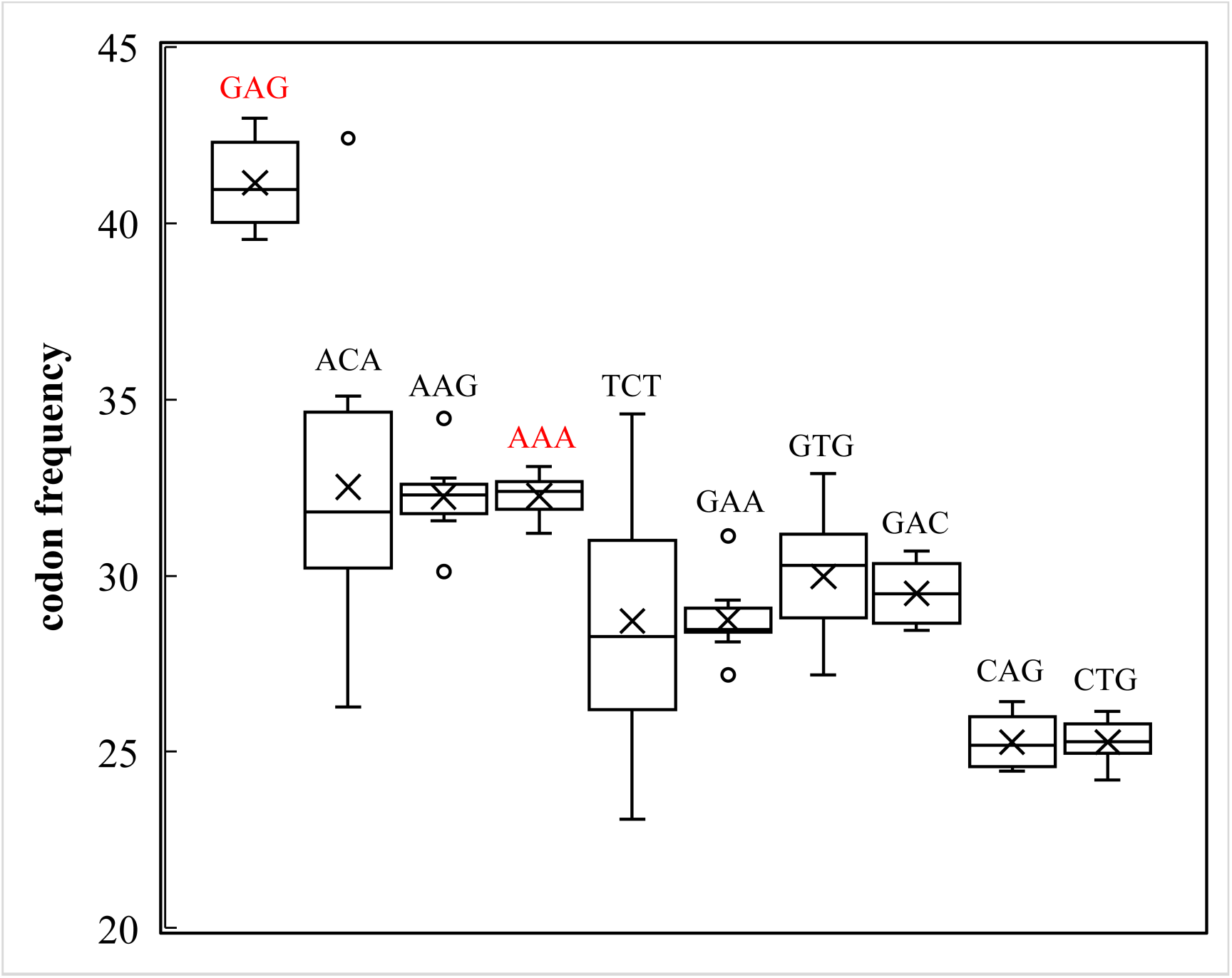
Codon frequency boxplot from ten pleural-pedal ganglia RNA-seq datasets of *Aplysia californica*. Frequency is defined as the expected number of codons, given the input sequence(s), per 1000 bases (EMBOSS: cusp). Frequency box plots of top ten codons are shown, with median values included for quartile calculation, and cross markers represent mean values and circles are outliers, which are values < Q1 − 1.5×IQR or > Q3 + 1.5×IQR (IQR= interquartile range).

### Isolation of *Aplysia* tRNA^Glu^CUC and tRNA^Lys^UUU and profiling of modified ribonucleosides by LC-MS/MS

Given the high sequence similarity among tRNA isodecoders, tRNA isolation by complementary DNA-probes can lead to contamination due to non-specific hybridization. To mitigate this, we designed DNA probes based on comparative multi-sequence alignment analysis of *Aplysia* tRNAs predicted from the nuclear genome (44) and mitochondrial genome (45). tRNA^Lys^UUU and tRNA^Glu^CUC each differ from their most similar tRNA sequences by more than three nucleotides at the 3’ end region (Supplementary Materials 4 & 5), which is sufficient for DNA probe-based isolation of high-purity tRNAs (57). Biotinylated DNA probes (30 nt) were thus designed complementary to the 3’ region of tRNA^Lys^UUU or tRNA^Glu^CUC (Supplementary Material 1: Table S3) and used in conjunction with streptavidin agarose beads to isolate tRNAs from the small RNA fraction of multiple different tissues. Polyacrylamide gel analysis (PAGE) of tRNA^Glu^CUC showed intact and lower molecular weight bands (i.e., <50 nt) (Figure 2A), indicating co-purification of tRNA fragments. Since the DNA probe used for isolation was complementary to the 3’ end of tRNA^Glu^CUC, the fragment bands likely correspond to 3’ tRNA^Glu^CUC fragments (investigated in subsequent section). tRNA^Lys^UUU showed obvious bands near 80 nt and negligible signal elsewhere, indicating high purity tRNA preparations (Figure 2B). Overall, these results showed low levels of contamination with ribosomal RNAs or other small RNAs.

**Figure 2.**
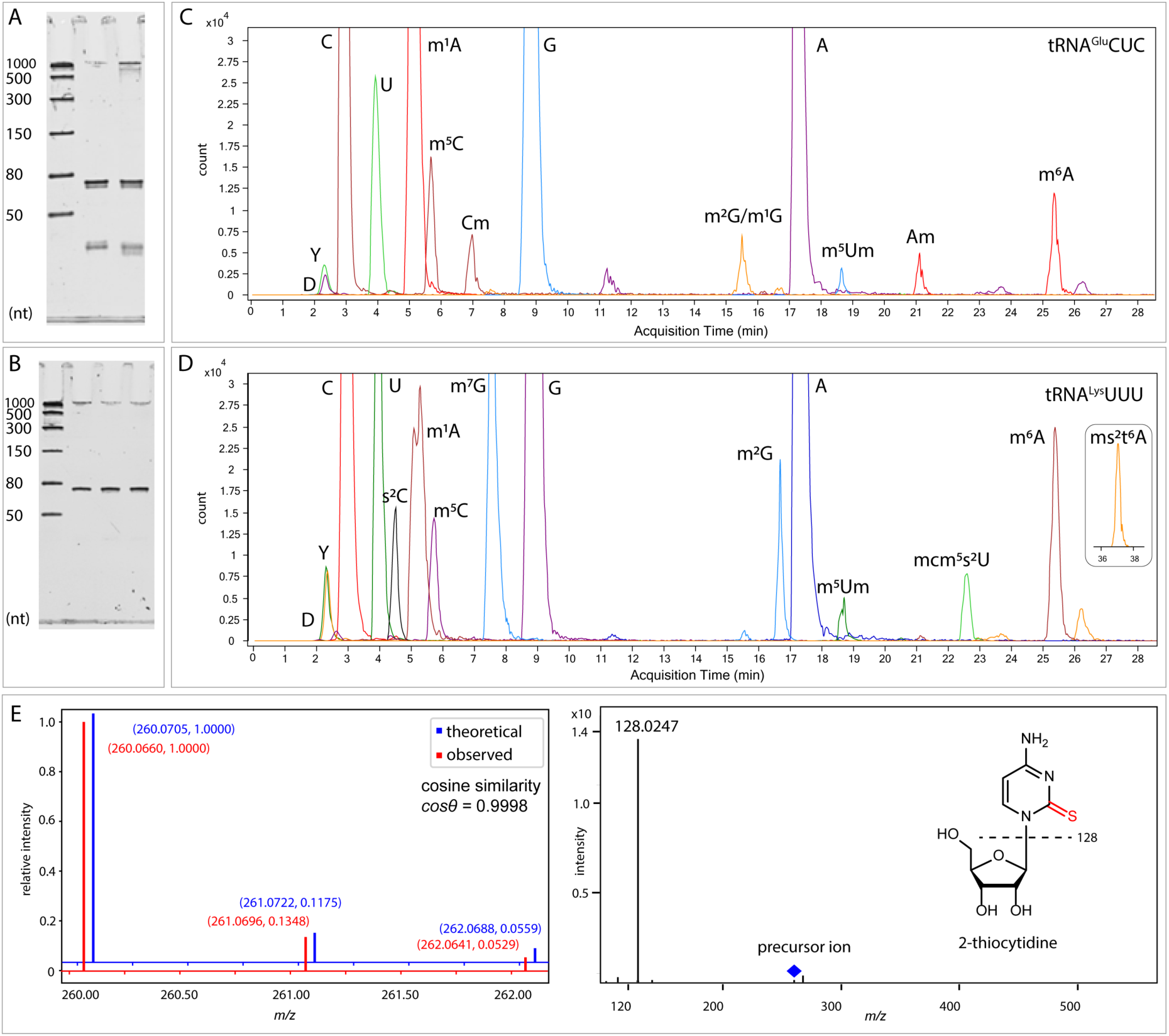
Nucleoside profiling analysis. Denaturing polyacrylamide gel electrophoresis (10%) for *Aplysia* tRNA^Glu^CUC (A) and tRNA^Lys^UUU (B) samples. ssRNA ladders were added on the left most lane on each gel, and the length of ladders (top to bottom) are: 1000, 500, 300, 150, 80 and 50 nt, respectively. For each gel, preparations from different tissues were loaded. For tRNA^Glu^CUC: muscle (2^nd^ lane), accessary genital mass (3^rd^ lane). For tRNA^Lys^UUU: all three lanes were from heart. Ribonucleoside profiles are shown by overlaid EICs of *Aplysia* tRNA^Glu^CUC (C) and tRNA^Lys^UUU (D). Unlabeled peaks are contaminating substances other than nucleosides, as verified by MS/MS spectra. Peak shape was smoothed to enhance visual clarity and ms^2^t^6^A eluted between 36 and 38 min (inset). (E) Isotopic envelope fitting analysis for s^2^C, and MS/MS spectrum showing characteristic product ion of 128 Da. Theoretical and observed *m/z* values along with relative intensities are shown in parenthesis.

Ribonucleoside compositions of each tRNA sample were then elucidated by digestion to nucleosides and LC-MS/MS analysis. A total of thirteen and fifteen ribonucleosides were detected from tRNA^Glu^CUC and tRNA^Lys^UUU, respectively (Figure 2C & D). The identities of ribonucleosides were verified by manual inspection of extracted ion chromatograms (EICs) and comparison of accurate masses, MS/MS product ions, and relative LC elution orders to known values (1). In tRNA^Glu^CUC, three methylated adenosine nucleosides, namely m^1^A, 2’-O-methyladenosine (Am), and N6-methyladenosine (m^6^A) were detected. The origin of m^6^A requires further verification, as Dimroth rearrangement of m^1^A to m^6^A is known to occur during the two-step digestion method. Two methylated cytidine nucleosides were detected and assigned as 5-methylcytidine (m^5^C) and 2’-O-methylcytidine (Cm), respectively. One species of methylated guanosine was detected, which eluted between guanosine and adenosine; MS/MS product ion indicates methylation on the base, suggesting either N1-methylguanosine (m^1^G) or N2-methylguanosine (m^2^G). 5,2’-O-dimethyluridine (m^5^Um), a modification commonly found in the T loop (1, 58), was also detected in tRNA^Glu^CUC digest. The *Aplysia* tRNA^Glu^CUC modification pattern resembles tRNA^Glu^CUC of human, mouse and yellow lupin (50, 59, 60) particularly the hypomodified anticodon loop, which may make tRNA^Glu^CUC a target for ribonuclease cleavage to generate tRNA fragments (61), as discussed in later sections.

In nucleoside digest samples from *Aplysia* tRNA^Lys^UUU, a total of fifteen ribonucleosides were detected (Figure 2D). Among these, two species of methylated adenosine (m^1^A and m^6^A), one species of methylated cytidine (m^5^C), and two species of methylated guanosine (m^2^G and m^7^G) were confirmed by the same approach applied for tRNA^Glu^CUC samples. The nucleosides data unambiguously identified mcm^5^s^2^U and 2-methylthio-N6-threonylcarbamoyladenosine (ms^2^t^6^A), which are commonly reported in eukaryotic tRNA^Lys^ in the anticodon loop (52, 54, 62). Notably, we also detected 2-thiocytidine (s^2^C), which would signify the first report of s^2^C in eukaryotes. The observed isotope envelope matched the theoretical simulation for s^2^C with a cosine similarity value of 0.9998, and the expected product ion of 128 Da was observed (Figure 2E), collectively supporting detection of s^2^C. All detected nucleosides match the LC retention order and MS1 data with high accuracy, and the confirmation of m^1^A and mcm^5^s^2^U in *Aplysia* tRNA^Glu^CUC and tRNA^Lys^UUU motivated us to map modifications in these two tRNAs.

### LC-MS/MS analysis revealed sub-stoichiometric modifications on *Aplysia* tRNA^Glu^CUC and tRNA^Lys^UUU

For LC-MS/MS sequencing of tRNA modifications, digestion by ribonuclease is desirable to generate smaller fragments amenable to LC separation and MS detection. However, digestion with a single ribonuclease limits sequence coverage and sometimes misses regions due to structural hinderance or cleavage-blocking modifications. We thus employed a parallel nuclease digestion strategy with RNase T1 and RNase 4 to increase sequence coverage and potentially identify sub-stoichiometric modifications of *Aplysia* tRNA^Glu^CUC and tRNA^Lys^UUU. RNase T1 cleaves the phosphodiester bond to the 3’ of guanosine residues (modifications such as 7-methylation blocks cleavage), while RNase 4 cleaves with selectivity for U followed by a purine and is reportedly insensitive to modifications. Combining ribonucleoside profiles, manual inspection of EICs, tandem MS spectra verification, and comparative analysis of tRNA modification positions across multiple organisms (Supplementary Material 1: Table S9), modifications were mapped to tRNA^Glu^CUC and tRNA^Lys^UUU as summarized in Figure 3. All expected RNase T1 digestion fragments ≥ 2 nt were observed, confirmed by EICs for multiple charge states and by MS/MS data (Supplementary Materials 2 & 3). Combined with the detected RNase 4 digestion fragments, sequence coverages of 88% (66/75 nt) for tRNA^Glu^CUC and 89% (68/76 nt) for tRNA^Lys^UUU were achieved. Cleavage after G_46_ in tRNA^Lys^UUU was not observed, suggesting that 7-methylguanosine (m^7^G) resides at this position, as this modification prevents RNase T1 cleavage (63) and was detected by nucleosides analysis. In contrast, N2-methylguanosine (m^2^G) does not interfere with RNase T1 and was mapped to position 10 on tRNA^Lys^UUU. For tRNA^Glu^CUC, our current data could not distinguish between m^1^G or m^2^G. Although m^2^G on position 10 is a common modification on different tRNAs among various species (64, 65), the occurrence of m^1^G on *Aplysia* tRNA^Glu^CUC could not be ruled out.

**Figure 3.**
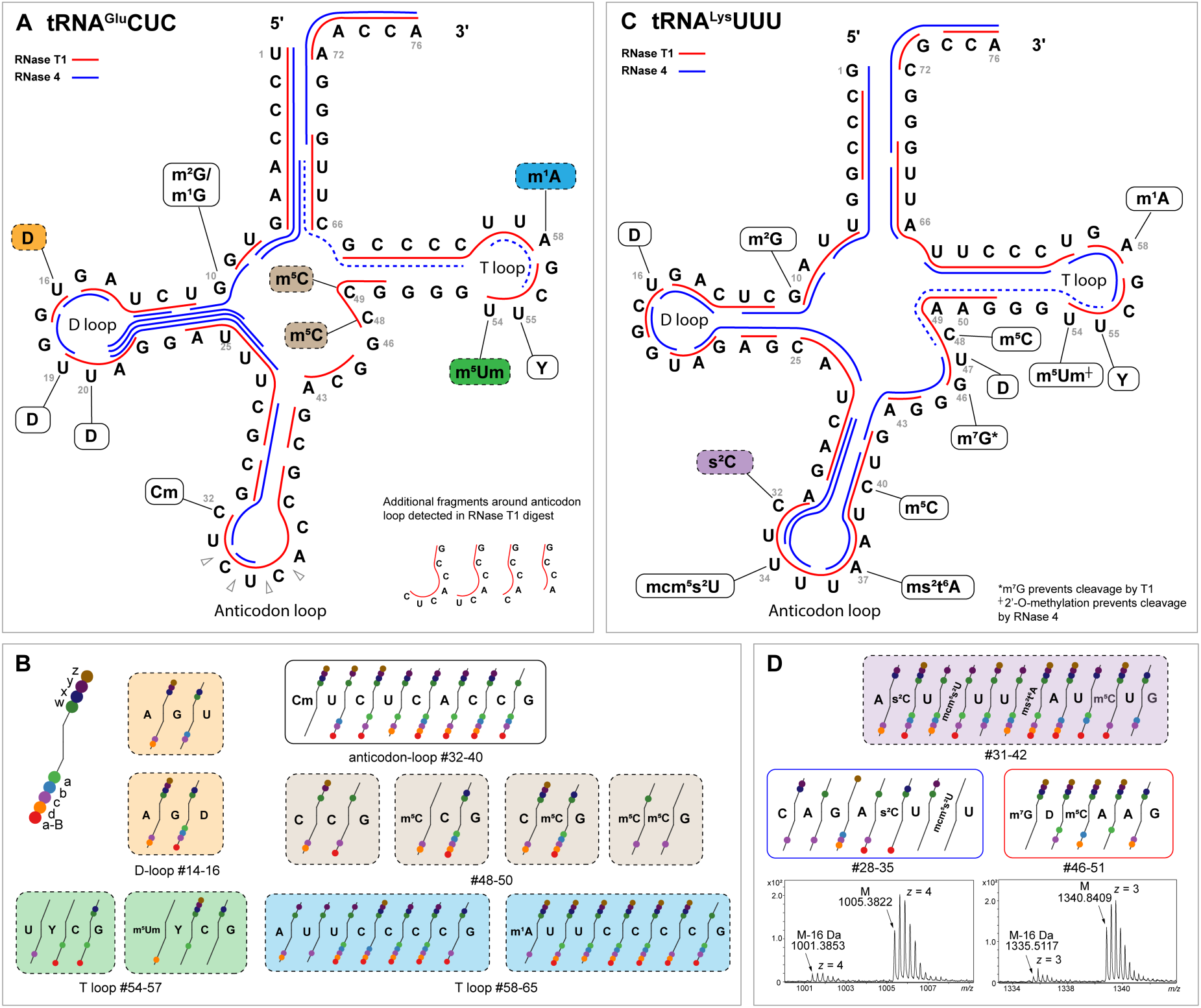
(A): Mapping modifications of tRNA^Glu^CUC by parallel nuclease digestion and LC-MS/MS. Observed fragments for RNase T1 (red lines) and RNase 4 (blue) are indicated. Dashed blue lines show predicted fragments that were undetected, and sub-stoichiometric modifications are labeled by colored squares. (B): MS/MS product ions coverage for tRNA^Glu^CUC digest. Fragments with sub-stoichiometric modification are enclosed by dashed lines, and color scheme match that in part A. (C): Sequence map for tRNA^Lys^UUU. (D): MS/MS product ions coverage for tRNA^Lys^UUU digest fragments, and mass spectra for the RNase T1-digest fragment spanning A31 to G42, with the isotopic envelopes of M and M-16 Da (z=3 and 4) variants presented.

Some nucleotide positions on tRNAs are modified only in a fraction of molecules rather than in every copy, a phenomenon known as sub-stoichiometric modification. Hypomodified tRNAs are structurally and functionally different from mature tRNAs, and tRNAs lacking specific modifications affect translation efficiencies in a codon biased fashion, as reported in model organisms. In mouse, the absence of queuosine (Q) at the wobble position leads to stalling of ribosomes on Q-decoded codons and a global imbalance in translation elongation speed, leading to learning and memory deficits (3). In *Drosophila*, loss of tRNA wobble uridine methylation resulted in decreased selenoprotein levels and has been linked to associative memory impairment (4). However, modifications in *Aplysia* tRNAs have not been mapped, and thus their stoichiometries have not been characterized.

Here, we found a series of sub-stoichiometric modifications on *Aplysia* tRNAs. For tRNA^Glu^CUC, EICs of nucleolytic fragments and MS/MS mapping results support sub-stoichiometric m^5^C_48_, m^5^C_49_ in the variable loop, as well as m^5^Um_54_ and m^1^A_58_ in the T-loop (Figure 3B, Supplementary Material 2). Deconvoluted spectra from intact mass analysis of tRNA^Glu^CUC from muscle and spermatheca showed a dominant monoisotopic mass value of 24188.490 Da (n=4), closely matching a theoretical mass of tRNA^Glu^CUC with all mapped modifications except a methyl group (24188.225 Da, mass accuracy = 11.0 ppm, Supplementary Material 1: Table S11), hence referred to as tRNA^Glu^CUC(–Me). A weaker signal for tRNA^Glu^CUC with deconvoluted mass of 24202.328 Da was also observed, matching the theoretical mass of 24202.241 Da (mass accuracy = 3.6 ppm) with complete modification status, referred to as tRNA^Glu^CUC(+Me). The tRNA^Glu^CUC(–Me) variant is most likely caused by a combination of methylation statuses at m^5^C_48_, m^5^C_49_, and m^1^A_58_, since m^1^A and m^5^C are reversible modifications (66–70). This suggests that on a given tRNA^Glu^CUC, if any one position is demethylated, the methyl-eraser enzyme is less likely to further act on this tRNA, making it less likely to be doubly- or triply-demethylated; an alternative explanation for this observation would be the affinity of methyl-writer enzyme to the tRNA^Glu^CUC substrate is diminished upon initial methylation, reducing likelihood of methylation at all three sites. In addition, RNase 4 digest revealed sub-stoichiometric incorporation of dihydrouridine at position 16 in the D-loop of tRNA^Glu^CUC (Figure 3B, Supplementary Material 2). Dihydrouridine is one of the most common modified nucleosides found in the D loop and increases the flexibility of the tRNA backbone, contributing to proper tRNA folding, recognition by aminoacyl-tRNA synthetases, and overall structural stability of the tRNA (71, 72). Rider et al. reported that other tRNA modifications are prerequisites for uracil reduction by DUS, indicating that D is incorporated at a later step in tRNA maturation (73). Although sequencing methods were developed for detection of D, and substrate specificity of dihydrouridine synthase has been revealed (74–76), tissue-specific dihydrouridine modification patterns have not been investigated. Our results revealed differential D_16_ stoichiometry on tRNA^Glu^CUC across *Aplysia* tissues (Supplementary Material 2). For example, D_16_ levels in tRNA^Glu^CUC from the accessory genital mass were lower than that of heart or muscle. It was reported that in *E. coli* D levels affect aminoacylation of specific tRNAs and modulate translation at corresponding codons (77). For a multi-cell organism like *Aplysia*, the distinct levels of D_16_ on tRNA^Glu^CUC may reflect differential codon usage in different tissues.

For tRNA^Lys^UUU, mcm^5^s^2^U, ms^2^t^6^A and m^5^C were mapped to anticodon loop positions 34, 37 and 40, respectively (Figure 3C; Supplementary Material 3), and matched conserved modification positions reported for *Heterololigo bleekeri* tRNA^Lys^ (74, 75). Notably, 2-thiocytidine (s^2^C) was mapped to position 32, despite s^2^C having not been reported in tRNAs of *Heterololigo bleekeri* nor any other eukaryotes to date. Deconvoluted spectra from intact mass analysis of tRNA^Lys^UUU showed a dominant monoisotopic mass of 24,898.387 Da (averaged, n=3; Supplementary Material 1: Table S11), which closely matches the theoretical monoisotopic mass (24,898.280 Da, 4.3 ppm) predicted by combining data from bottom-up modification mapping analysis. For the RNase T1-digest fragment spanning A_31_ to G_42_, two additional variants of M-16 Da and M-32 Da were detected, indicating potential sub-stoichiometric modifications (Figure 3D; Supplementary Material 3). These variants were also detected by intact mass analysis: for M-16 the mass accuracy is 1.6 ppm; for M-32, mass accuracy could not be calculated because it was not obvious to us what sequence/modification would result in M-32. Since the precursors of mcm^5^s^2^U_34_ or ms^2^t^6^A_37_ (i.e., mcm^5^U, cm^5^s^2^U, t^6^A) were not detected in the nucleosides analysis of tRNA^Lys^UUU, the M-16 Da variants detected at oligonucleotide and intact tRNA levels were likely due to hypomodification on C_32_, but the identity of the M-32 Da variant deserves further investigation. One possibility is that other substoichiometric S-containing modifications were present and not detected during nucleosides analysis. As noted above, LC-MS/MS data from RNase 4 digestion improved sequence coverage from 89% (68/76) to 97% (74/76) for *Aplysia* tRNA^Lys^UUU, by providing RNA fragments that covered consecutive guanosine residues. Our experiments showed RNase 4 cleavage products for all nucleotides to the 3’ of uridine (U/A, U/G and U/U), and there were several unassigned products that indicate unexpected cleavage selectivity. Very few expected RNase 4 fragments around the anticodon loop and T loop of *Aplysia* tRNA^Glu^CUC were observed. Thus, parallel RNase 4 and T1 digests only improved the coverage of tRNA^Glu^CUC from 88% (66/75) to 91% (68/75). Since tRNA^Glu^CUC has no uridine residues from position 36 to 58, it is possible that longer fragments produced by RNase 4 digestion were not well separated/ionized by our LC-MS/MS method and were not detected. Nonetheless, our strategy addressed the limitations of RNase T1 mapping which cannot sequence consecutive guanosine residues. Moreover, RNase 4 digestion provided additional fragments to validate sub-stoichiometric modifications such as D_16_ in tRNA^Glu^CUC, and provided additional support for s^2^C_32_, mcm^5^s^2^U_34_, ms^2^t^6^A_37_ and m^5^C_40_ on anticodon loop and anticodon arm of tRNA^Lys^UUU.

### Characterization of tRNA halves from *Aplysia* tRNA^Glu^CUC and tRNA^Lys^UUU

Transfer RNA-derived small RNAs (tsRNAs) are small non-coding RNAs produced by enzymatic cleavage of tRNAs. Based on length and tRNA cleavage position, they are classified as tRNA halves (or stress-induced RNAs, tiRNAs, ∼31-40 nt, cleavage at anticodon loop) and tRNA-derived fragments (tRFs, ∼14-30 nt, cleavage at D and T loops in addition to anticodon loop) such as tRF-1, tRF-2, tRF-3, tRF-5 and i-tRF (80, 81). tsRNAs are involved in various cellular processes including stress granule formation, ribosome assembly, and regulation of histone levels, and are attracting particular interest in neurons as disease-specific tsRNA profiles have been identified in a number of CNS disorders (8). However, modifications of tsRNAs such as m^1^A and m^3^C impede reverse transcription (82, 83), making NGS-based detection of tsRNAs challenging. Moreover, the end chemistry of tsRNAs necessitates specialized library preparation protocols (8, 84, 85). Although ligation-independent cloning approaches are emerging for detection of tsRNAs (86, 87), the high copy number and similarities between tRNA genes makes sequence mapping challenging (43), hindering modification detection and quantification. We developed an LC-MS based method to directly detect tsRNAs regardless of modification status and end chemistry. Gel electrophoresis results (Figure 2A, B) and the detection of additional, unexpected digest fragments mapping to the anticodon loop and anticodon stem of tRNA^Glu^CUC that would not result from RNase T1 cleavage (see Figure 3A inset) prompted us to profile tsRNAs by intact mass analysis. Six different 3’ tsRNAs were consistently detected by IP-RPLC-MS in tRNA^Glu^CUC samples from spermatheca and muscle (Figure 4). Deconvoluted mass values showed that the length of these six tsRNAs were 37, 38, 39, 40, 41, and 42 nt, respectively. The 39-nt tsRNA was the most abundant fragment based on estimated peak height from EICs (Figure 4B). Notably, for each of the six tsRNAs, signal for the –Me variants were more abundant than the +Me variants, as exemplified by 39-nt tsRNA^Glu^CUC (Figure 4C). While not observed by gel electrophoresis, intact mass analysis revealed tsRNA^Lys^UUU of 40 and 41 nt in pooled samples consisting of heart and reproductive tissue (Figure 4E, F). The deconvoluted mass values correspond to fully modified variants, while hypomodified variants were not detected (Supplementary Material 1: Figure S6).

**Figure 4.**
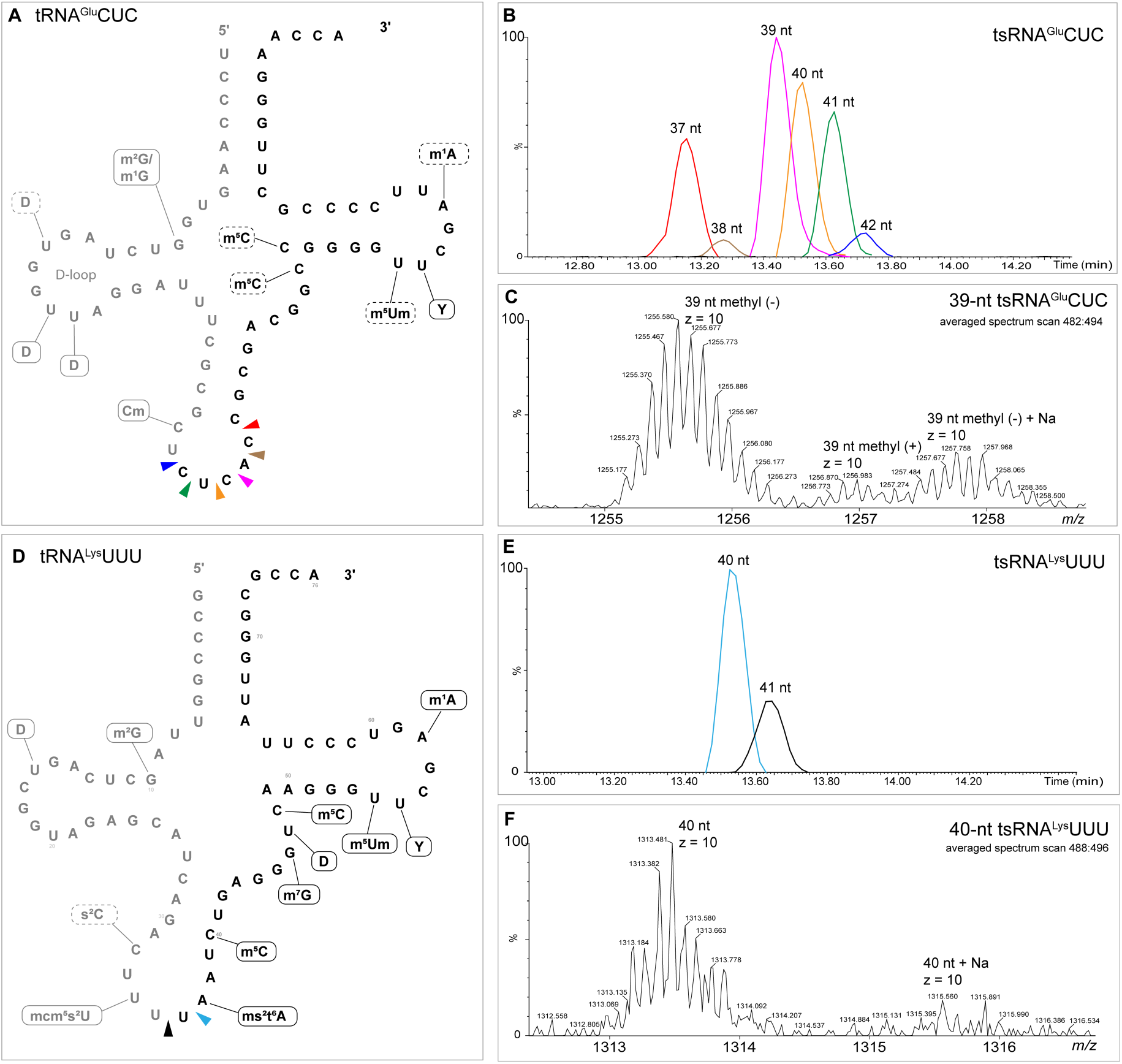
tRNA halves of *Aplysia* tRNA^Glu^CUC and tRNA^Lys^UUU. (A): Cleavage sites on tRNA^Glu^CUC corresponding to detected tsRNAs, with colored triangles matching EICs in B. (B): EICs of detected tsRNAs (methyl (-), *z*=10) in spermatheca sample, smoothed by Savitzky-Golay method. (C): Averaged mass spectrum of scan 482-494 from spermatheca sample, with three isotopic envelopes of 39 nt tsRNA labeled. (D): Cleavage sites on tRNA^Lys^UUU corresponding to detected tsRNAs with colored triangles matching EICs in E. (E): EICs of detected tsRNAs (*z*=10) in heart + reproductive tissue pooled sample, smoothed by Savitzky-Golay method. (F): Averaged mass spectrum of scan 488-496 from heart + reproductive tissue pooled sample, with two isotopic envelopes of 40 nt tsRNA labeled.

Based on our intact mass analysis, all detected *Aplysia* 3’ tsRNAs have hydroxyl groups on both ends (Supplementary Material 1: Table S12, S13), consistent with angiogenin-cleavage end chemistry. Angiogenin (ANG) is an endoribonuclease of RNase A superfamily involved in the cleavage at tRNA anticodon-loop, induced by diverse cellular stimuli, including stress (88, 89), sex-hormone signaling (90), and mycobacterial infection (91). Members of the RNase A superfamily typically cleave 3’ of pyrimidines, with a marked preference for cytidine compared to uridine (92), and ANG shows highest specificity for cleavage between 5’-C and A-3’, followed by 5’-C–G-3’ (93). Our data show that the 39-nt 3’ tsRNA^Glu^CUC (cleavage between C_36_ and A_37_) is the most abundant, followed by 40-nt, 41-nt, and 37-nt tsRNAs (Figure 4B), which resembles the cleavage specificity of ANG. However, the RNase A superfamily is vertebrate-specific (94), so it is unclear which ribonuclease cleaves these tRNAs in *Aplysia*. Both +Me and –Me variants were detected for all six tsRNAs derived from tRNA^Glu^CUC, indicating both variants are substrates for tRNA cleavage.

tRNA modifications may either protect or promote enzymatic tRNA cleavage depending on modification identity and position (61). For example, loss of methylation at tRNA C_38_ in eukaryotes leads to accumulation of 5’-tsRNAs (95–97), while queuosine (Q) modification can either protect cognate tRNAs against ribonuclease cleavage (98) or mark Q-containing tRNAs as targets of ribonuclease (99). *Aplysia* tRNA^Glu^CUC is hypomodified at the anticodon loop, so the relative abundance of tsRNA^Glu^CUC likely represents a cleavage pattern largely based on sequence preference of the cleaving enzyme(s) rather than modification. On the other hand, *Aplysia* tRNA^Lys^UUU harbors mcm^5^s^2^U at position 34, a modification that was reported to mark tRNA as cleavage targets in yeast (100). However, the abundance and number of tsRNA^Lys^UUU detected was far lower than that of tsRNA^Glu^CUC, suggesting s^2^C32 and ms^2^t^6^A37 on the anticodon loop may have protective effects. Overall, tsRNA generation appears to be dependent on tRNA sequence and modification status, and future characterization of tsRNAs and their modifications would allow a refined set of rules for tRNA cleavage to be established.

tsRNAs have been reported to be involved in neurological disorders such as Alzheimer’s, Parkinson’s, and Huntington’s disorder, with underlying mechanisms including modulation of translation and gene expression (7, 8, 101). tsRNAs inhibit translation initiation by a mechanism dependent on G-quadruplex secondary structure (G4) formation, where tsRNAs displace eIF4F complexes from the m^7^GTP cap of mRNAs and lead to stress granule formation (102–106). These studies focused on 5’ tsRNAs and used synthetic oligonucleotides without naturally occurring post-transcriptional modifications in their experiments, so whether endogenous 3’ tsRNAs assemble into G4 and how modifications affect the structure and function deserves further investigation. In *Aplysia* tsRNA^Glu^CUC, the T arm contains four consecutive guanosine residues (position G_50_-G_53_), raising the possibility of intermolecular G4 formation, and these tsRNAs may serve as a reservoir for fast response to external stimuli, such as during learning paradigms. Beyond intracellular regulation, tsRNAs may also serve cell-to-cell signaling functions, as synaptic vesicles isolated from the CNS of *Torpedo californica* (Pacific electric ray) and mouse contain 5’ tsRNAs which may regulate local protein synthesis (6). Worth noting is that an NGS-based method was employed to detect tsRNAs, and library preparation may render 3’ tsRNAs in synaptic vesicles underrepresented.

### Two key enzymes in s^2^C biosynthesis pathway identified in *Aplysia californica* genome

The presence of s^2^C was confirmed by LC-MS-based nucleoside analysis in *Escherichia coli* (107, 108), *Salmonella enterica* (109, 110) and *Methanococcales* (archaea) (111, 112) but this modification has yet to be reported in eukaryotic organisms. Three key enzymes involved in s^2^C biosynthesis were previously identified in *E. coli* and *S. enterica*. IscS, a cysteine desulfurase, is required for the biosynthesis of all thiolated nucleosides (110, 113) as it initiates the pathway by extracting sulfur from L-cysteine and generating a persulfide intermediate that serves as the universal sulfur donor for downstream reactions. IscU, an iron-sulfur cluster assembly scaffold protein, then uses sulfur from IscS, together with iron, to assemble a [4Fe–4S] cluster (114) that is subsequently transferred to TtcA, the dedicated 2-thiocytidine synthetase (109). Cluster-bound TtcA accepts persulfide sulfur from IscU and catalyzes the redox-dependent insertion of this sulfur atom at the C2 position of C_32_ within the anticodon loop of *E. coli* tRNA^Arg^ACG, tRNA^Arg^CCG, tRNA^Arg^UCU, and tRNA^Ser^GCU (108, 115). While IscU is also involved in biosynthesis of ms^2^i^6^A_37_ (116), TtcA is specific to s^2^C, and genes homologous to TtcA have been identified in bacteria and archaea, and some eukaryotes including *Saccharomyces cerevisiae* and *Giardia lamblia* (protozoan) (109, 115). Nonetheless, evidence of eukaryotic s^2^C from LC-MS/MS has been lacking until the present study.

After we detected s^2^C_32_ in nucleosides digests and mapped s^2^C in *Aplysia* tRNA^Lys^UUU via RNase T1 and RNase 4 digestions, we searched for homologs of the three key enzymes IscS, IscU and TtcA in *Aplysia californica* genome. With the genome annotation on NCBI, the iron-sulfur cluster assembly scaffold protein IscU was located to LOC101853926. Sequence comparison revealed that the *Aplysia* IscU homolog shares 75% amino acid sequence identity with *E. coli* IscU at the aligned region. A predicted IscS homologous gene (LOC101849150), targeted to mitochondria, shares 61% amino acid sequence identity with *E. coli* IscS at the aligned region. Blastp search using *E. coli* TtcA as the query retrieved an uncharacterized protein encoded by LOC101861551, with 34% identities and 50% positives within the aligned regions. Domain analysis by NCBI’s CD-Search revealed that this 1177-aa peptide contains a CsdA domain (Selenocysteine lyase/Cysteine desulfurase, IscS, #COG0520) and a TtcA-like domain (#cd24138) (Figure 5; Supplementary Material 6). NCBI RNA-seq coverage for this gene showed clear intron-exon junctions, and comparable expression level across exons, suggesting it is expressed as a consecutive mRNA transcript.

**Figure 5.**
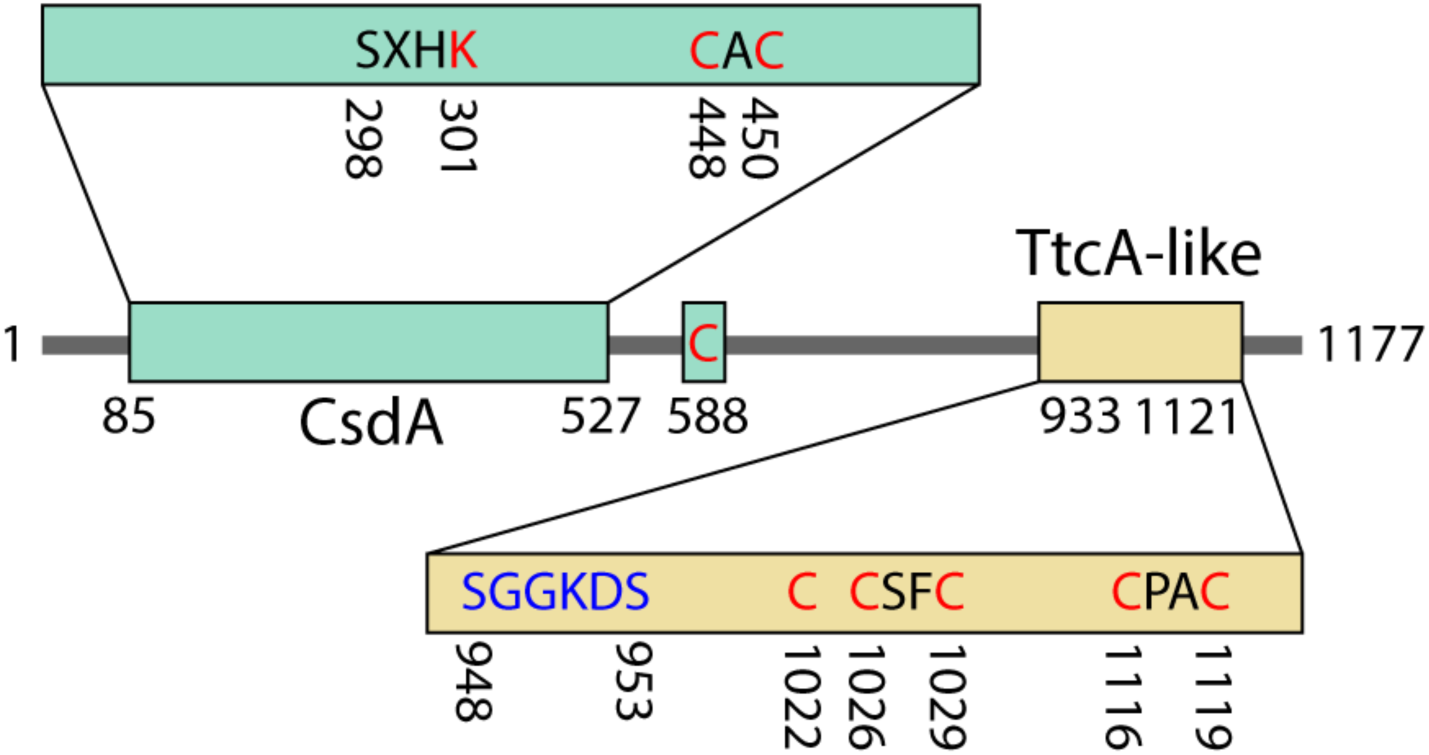
Domain structures of *Aplysia* IscS-TtcA The two conserved domains are illustrated as colored boxes and conserved amino acid residues are displayed. Red fonts indicate functionally critical residues revealed by previous studies, and blue font marks the PP-loop.

The *E. coli* genes encoding IscS (gene ID: 947004) and TtcA (gene ID: 948967) are more than 1.2 million base pairs apart on the circular chromosome. In contrast, LOC101861551 in the *Aplysia* genome encodes a peptide containing both domains, suggesting that the ancestral genes were fused. To investigate if this gene fusion is specific to *Aplysia* or to a broader group of organisms, the peptide sequence of *Aplysia* IscS-TtcA was used to conduct blastp searches against available genomes of the phylum Mollusca on NCBI. The top 100 hits were retrieved, aligned by MuscleWS, and peptide sequences shorter than 1000 aa were removed from the alignment. The aligned sequences are from 28 mollusk species, spanning different classes including Bivalvia, Cephalopoda, Gastropoda and Polyplacophora (Supplementary Material 7). Blastp search retrieved homologous sequences from even more distant species such as birds and insects (data not shown), indicating the fusion of these two genes in a wide range of taxonomic groups. Whether this fused gene was a result of parallel evolution or was inherited from common ancestor deserves further investigation.

Among the aligned peptides, the lysine residue (Lys206 in *E. coli* IscS, Lys301 in *Aplysia* IscS-TtcA) for pyridoxal 5′-phosphate (PLP) binding (117, 118) is conserved (Figure 5). Although the catalytic cysteine residue (Cys328 in *E. coli* IscS and Cys381 in human cysteine desulfurase NFS1 (117, 119)) was not identified among the fused peptide sequences, Cys448, Cys450 and Cys588 (*Aplysia* IscS-TtcA) are highly conserved in the alignment and could be candidate active residues. The conserved PP-loop (ATP-binding) motif (SGGKDS) and putative catalytic cysteine residues critical for TtcA activity (109, 115, 120, 121) are highly conserved among the aligned sequences, although instead of a CXXC-C-CXXC structure found in bacteria, the aligned mollusk sequences showed a C-CSFC-CPAC arrangement. Together this suggests that the fused peptide may utilize alternative cysteines as active residues of the CsdA domain, while the TtcA domain functionally resembles the non-fused counterpart. The region between these two conserved domains (405 aa in *Aplysia*) shows low levels of conservation among the aligned peptide sequences, indicating it is structurally flexible and is not under strong evolutionary constraints. This region may either act as a linker or is cleaved post-translationally to produce separate peptides.

Enzymes in the same metabolic pathway often evolve balanced expression levels, as supported by large-scale comparative studies showing that pathways tend to preserve stoichiometric ratios of enzyme synthesis across species (122). The fused IscS-TtcA gene may enable coordinated expression and balanced stoichiometry of these two enzymes, perhaps allowing efficient installation of s^2^C on tRNA. However, if IscS is required for biosynthesis of all thiolated ribonucleosides in these eukaryotes, as shown in bacteria (110, 113), the coupled expression of TtcA which is specific to s^2^C would seemingly be a waste of resources, unless benefits from the coupled expression outweigh the cost, or there are additional IscS genes in these eukaryotic genomes that encode enzymes for biosynthesis of other thiolated ribonucleosides. To test this, we used the IscS-TtcA fused peptide sequence as a query and performed a blastp search against the *Aplysia* genome and identified a stand-alone IscS/CsdA-like protein (encoded by LOC101846496). This stand-alone IscS/CsdA-like protein only shares 44% identities with the CsdA domain of the fused peptide, indicating their diverged functions. We thus hypothesize that the stand-alone IscS/CsdA-like protein is involved in thiolation of other nucleosides, and the IscS-TtcA fused enzyme has evolved to be dedicated to s^2^C biosynthesis, indicating s^2^C stoichiometry is under a more precise regulation.

### Coordination of s^2^C, mcm^5^s^2^U and ms^2^t^6^A at the anticodon loop of tRNA^Lys^UUU enhances cognate codon pairing

Across all characterized tRNA structures, C_32_ and A_38_ engage in a conserved bifurcated hydrogen bond (O2 of C_32_ to N6 of A_38_), resulting in a noncanonical mismatch base pair positioned at the anticodon stem–loop junction (123). s^2^C_32_ restricts tRNA wobble decoding by altering this C_32_-A_38_ cross-loop interaction (124). Found at the wobble position of tRNA^Lys^, tRNA^Glu^ and tRNA^Gln^, mcm^5^s^2^U_34_ prevents wobble pairing with near-cognate codons by stabilizing Watson-Crick based pairing with A-ending codons (125) and enhancing binding of cognate tRNAs to the ribosome A-site (22). ms^2^t^6^A_37_ rigidifies the anticodon loop to maintain the correct geometry for base pairing, reducing near-cognate codon pairing (126). Together, we hypothesize that the coordination of s^2^C_32_, mcm^5^s^2^U_34_ and ms^2^t^6^A_37_ on *Aplysia* tRNA^Lys^UUU restricts wobble flexibility and ensures tRNA^Lys^UUU exclusively decodes codon AAA but not AAG. On the other hand, neither C_32_ nor C_34_ in reported eukaryotic tRNA^Lys^CUU (except the squid *Heterololigo bleekeri* (75)) is modified, because cytidine on tRNA and adenosine on mRNA won’t form stable base pairing, reducing the probability of tRNA^Lys^CUU wobble-pairing with AAA codons. Similarly, the wobble position C_34_ in *Aplysia* tRNA^Glu^CUC is not modified, but for tRNA^Glu^UUC, U_34_ is more likely modified to avoid wobble-paring with GAG. Indeed, U_34_ at tRNA^Glu^UUC is modified in both prokaryote and eukaryotes (127–131).

Codons AAA and AAG both encode lysine and exhibit comparable abundances in *Aplysia* transcriptome (frequency of AAA: AAG=0.91), but the occurrence of repetitive lysine codons (AAA)_4_ among coding sequences is only seven, far lower than that of (AAG)_4,_ which is 287. This phenomenon is also pronounced in various other organisms (Figure 6; Supplementary Material 1: Table S15), but few studies have investigated the mechanism behind this imbalance between two synonymous codons. Koutmou et al. reported that consecutive AAA codons diminishes protein expression more than synonymous AAG codons, and they attributed this to ribosome sliding on homopolymeric adenosine sequences, which would result in nonsense-mediated-decay (132). Therefore, a plausible theory is that AAA codon is subjected to stronger selection over AAG codon, and AAA is being purged during evolution. However, this study did not investigate how modifications on tRNAs modulate the translation of (AAA)_4_ or (AAG)_4_ containing mRNAs. In *Aplysia* genome, for the seven (AAA)_4_ containing coding sequences, the coordination of s^2^C, mcm^5^s^2^U and ms^2^t^6^A at the anticodon loop of tRNA^Lys^UUU is likely involved in tight recognition of AAA codons, preventing ribosome sliding and ensuring accurate production of peptides containing consecutive lysine residues. The expression of these seven genes is thus likely modulated by s^2^C stoichiometry, and the exact mechanism deserves further experimental investigation.

**Figure 6.**
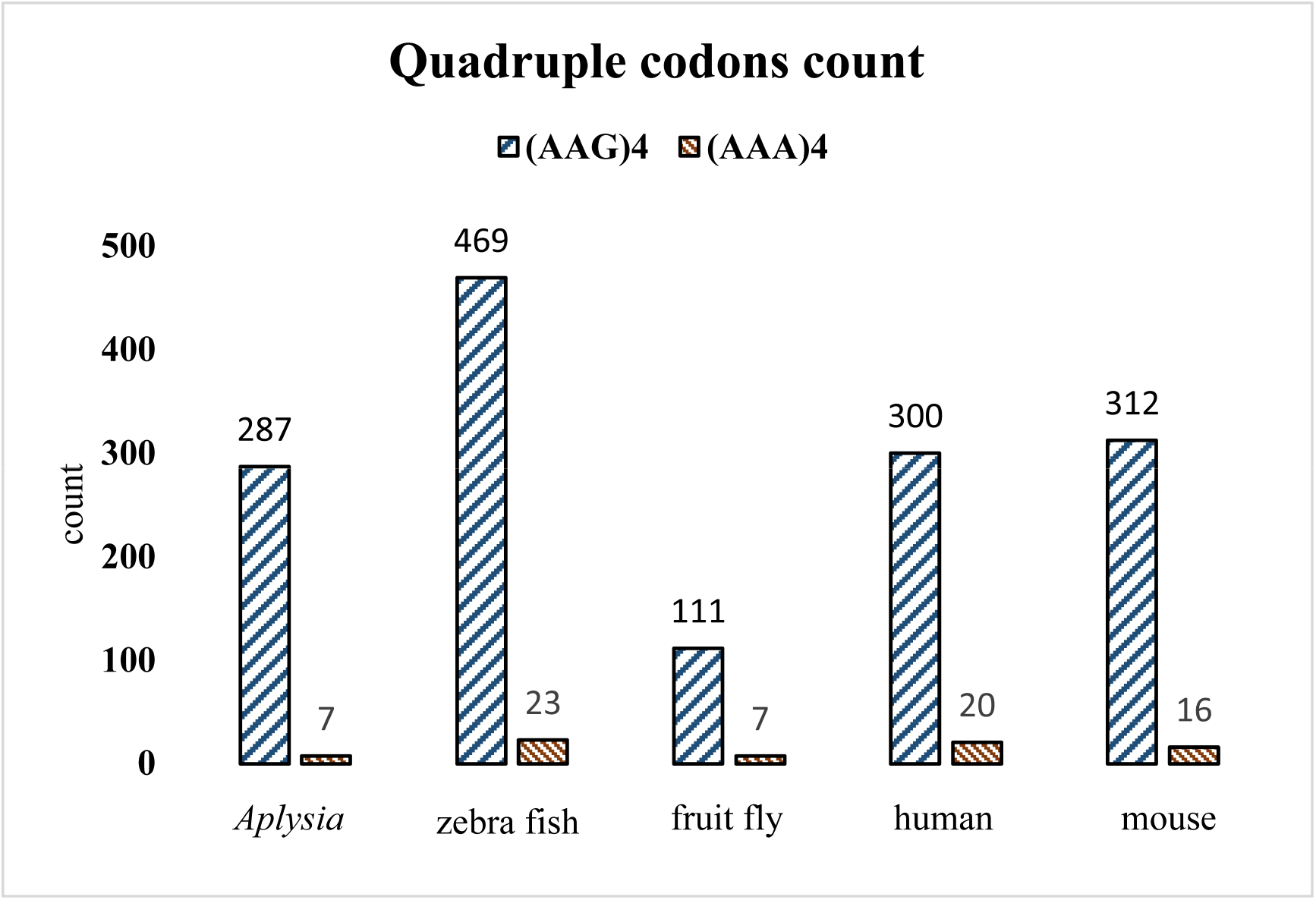
Quadruple codons count in eukaryote genomes, calculated from non-redundant coding sequences, excluding mitochondrial genes.

### Conclusion

*Aplysia californica* is a classical neurobiological model with decades-long impact in unraveling the molecular underpinnings of learning, memory, and behavior. Recent evidence revealed a link between tRNA modifications, protein synthesis, neuron excitability, and ultimately behavioral change. However, until now, no complete sequence modification information for any tRNA molecules in this organism has been available, severely limiting progress in understanding how tRNA modifications affect fundamental cellular functions and neurophysiology. Here, tRNA^Glu^CUC and tRNA^Lys^UUU were isolated from *Aplysia* tissues and an integrated strategy employed to map modifications. LC-MS/MS analysis mapped s^2^C, mcm^5^s^2^U and ms^2^t^6^A to the anticodon loop of tRNA^Lys^UUU and are essential for decoding fidelity and prevention of ribosome sliding. Sub-stoichiometric modifications including m^1^A were detected on tRNA^Glu^CUC and 3’ tsRNAs of different length. The distinct abundances of tsRNA^Glu^CUC of different length indicate a cleavage mechanism by an enzyme homologous to ANG, but the exact enzyme involved deserves further investigation. tsRNA^Glu^CUC contains consecutive guanosine residues, a structure prerequisite for intermolecular G4 formation which could regulate translation. Together, these results revealed potential origins of elevated m^1^A and mcm^5^s^2^U levels during behavioral sensitization in *Aplysia* previously reported. In addition, this study presents the first report of s^2^C on eukaryotic tRNA, and 3’ tsRNA detection by LC-MS, providing a new strategy for future studies targeting tsRNAs that are challenging to detect by RNA-seq due to their end chemistry and modification profiles.

## Supplementary data

Supplementary data is available at NAR online.

Supplementary material 1: Public datasets used in this study, LC-MS/MS analysis methods, tRNA intact analysis methods and results.

Supplementary material 2: tRNA^Glu^CUC information. Part1: RNase 4 data. Part2: RNase T1 data. Part 3. RNase T1 digested fragments from tsRNA^Glu^CUC

Supplementary material 3: tRNA^Lys^UUU information. Part1: RNase 4 data. Part2: RNase T1 data.

Supplementary material 4: Neighbor-joining tree of *Aplysia californica* tRNA sequences.

Supplementary material 5: Multi-sequence alignment of *Aplysia californica* tRNA sequences.

Supplementary material 6: *Aplysia californica* IscS-TtcA gene information

Supplementary material 7: Species taxonomy information

Supplementary material 8: s^2^C fitting analysis

## Acknowledgments

The authors are grateful to Yu-Li Shih and Gabriella Floro for their generous help with LC-MS/MS analysis of nucleosides.

## References

1. Sordyl, D., Boileau, E., Bernat, A., et al. (2026) MODOMICS: a database of RNA modifications and related information. 2025 update and 20th anniversary. Nucleic Acids Res., 54, D219–D225.

2. Blaze, J. and Akbarian, S. (2022) The tRNA regulome in neurodevelopmental and neuropsychiatric disease. Mol. Psychiatry, 27, 3204–3213.

3. Cirzi, C., Dyckow, J., Legrand, C., et al. (2023) Queuosine-tRNA promotes sex-dependent learning and memory formation by maintaining codon-biased translation elongation speed. EMBO J., 42, e112507.

4. Madhwani, K.R., Sayied, S., Ogata, C.H., et al. (2024) tRNA modification enzyme-dependent redox homeostasis regulates synapse formation and memory. Proc. Natl. Acad. Sci., 121, e2317864121.

5. Fagan, S.G., Helm, M. and Prehn, J.H.M. (2021) tRNA-derived fragments: A new class of non-coding RNA with key roles in nervous system function and dysfunction. Prog. Neurobiol., 205, 102118.

6. Li, H., Wu, C., Aramayo, R., et al. (2015) Synaptic vesicles contain small ribonucleic acids (sRNAs) including transfer RNA fragments (trfRNA) and microRNAs (miRNA). Sci. Rep., 5, 14918.

7. Mathew, B.A., Katta, M., Ludhiadch, A., et al. (2023) Role of tRNA-derived fragments in neurological disorders: a review. Mol. Neurobiol., 60, 655–671.

8. Winek, K. and Soreq, H. (2025) Emerging roles of transfer RNA fragments in the CNS. Brain, 148, 2631–2645.

9. Moroz, L.L. (2011) Aplysia. Curr. Biol., 21, R60–R61.

10. Bailey, C.H. and Kandel, E.R. (1993) Structural changes accompanying memory storage. Annu. Rev. Physiol., 55, 397–426.

11. Bartsch, D., Ghirardi, M., Skehel, P.A., et al. (1995) Aplysia CREB2 represses long-term facilitation: Relief of repression converts transient facilitation into long-term functional and structural change. Cell, 83, 979–992.

12. Castellucci, V., Pinsker, H., Kupfermann, I., et al. (1970) Neuronal mechanisms of habituation and dishabituation of the gill-withdrawal reflex in Aplysia. Science, 10.1126/science.167.3926.1745.

13. Kandel, E.R. (2001) The molecular biology of memory storage: a dialogue between genes and synapses. Science, 10.1126/science.1067020.

14. Clark, K.D., Philip, M.C., Tan, Y., et al. (2020) Biphasic liquid microjunction extraction for profiling neuronal RNA modifications by liquid chromatography–tandem mass spectrometry. Anal. Chem., 92, 12647–12655.

15. Clark, K.D., Rubakhin, S.S. and Sweedler, J.V. (2022) Characterizing RNA modifications in single neurons using mass spectrometry. J. Vis. Exp. JoVE, 10.3791/63940.

16. Clark, K.D., Rubakhin, S.S. and Sweedler, J.V. (2021) Single-neuron RNA modification analysis by mass spectrometry: characterizing RNA modification patterns and dynamics with single-cell resolution. Anal. Chem., 93, 14537–14544.

17. Huang, H.X., Floro, G.M., Asare, E., et al. (2026) Neuronal tRNA Modifications in Aplysia californica are Repatterned during Behavioral Habituation. Sci. Signal. Accepted, pre-print: 10.1101/2025.11.13.688303.

18. Kadaba, S., Krueger, A., Trice, T., et al. (2004) Nuclear surveillance and degradation of hypomodified initiator tRNAMet in S. cerevisiae. Genes Dev., 18, 1227–1240.

19. Yared, M.-J., Yoluç, Y., Catala, M., et al. (2023) Different modification pathways for m1A58 incorporation in yeast elongator and initiator tRNAs. Nucleic Acids Res., 51, 10653– 10667.

20. Björk, G.R., Huang, B., Persson, O.P., et al. (2007) A conserved modified wobble nucleoside (mcm5 s2 U) in lysyl-tRNA is required for viability in yeast. RNA, 13, 1245–1255.

21. Linder, B., Sharma, P., Wu, J., et al. (2025) tRNA modifications tune m6A-dependent mRNA decay. Cell, 188.

22. Rezgui, V.A.N., Tyagi, K., Ranjan, N., et al. (2013) tRNA tKUUU, tQUUG, and tEUUC wobble position modifications fine-tune protein translation by promoting ribosome A-site binding. Proc. Natl. Acad. Sci., 110, 12289–12294.

23. Bushmanova, E., Antipov, D., Lapidus, A., et al. (2019) rnaSPAdes: a de novo transcriptome assembler and its application to RNA-Seq data. GigaScience, 8, giz100.

24. Haas, B. (2025) TransDecoder.

25. Rice, P., Longden, I. and Bleasby, A. (2000) EMBOSS: The European Molecular Biology Open Software Suite. Trends Genet., 16, 276–277.

26. Crain, P.F. (1990) [42] Preparation and enzymatic hydrolysis of DNA and RNA for mass spectrometry. In Methods in Enzymology, Mass Spectrometry. Academic Press, Vol. 193, pp. 782–790.

27. Gehrke, C.W., Kuo, K.C. and Zumwalt, R.W. (1980) Chromatography of nucleosides. J. Chromatogr. A, 188, 129–147.

28. Su, D., Chan, C.T.Y., Gu, C., et al. (2014) Quantitative analysis of ribonucleoside modifications in tRNA by HPLC-coupled mass spectrometry. Nat. Protoc., 9, 828–841.

29. Harwood, T.V., Treen, D.G.C., Wang, M., et al. (2023) BLINK enables ultrafast tandem mass spectrometry cosine similarity scoring. Sci. Rep., 13, 13462.

30. Zhang, X., Wu, R. and Qu, Z. (2023) A cosine-similarity-based deconvolution method for analyzing data-independent acquisition mass spectrometry data. Appl. Sci., 13, 5969.

31. Rozenski, J. (1999) Mongo Oligo Mass Calculator.

32. D’Ascenzo, L., Popova, A.M., Abernathy, S., et al. (2022) Pytheas: a software package for the automated analysis of RNA sequences and modifications via tandem mass spectrometry. Nat. Commun., 13, 2424.

33. Mcluckey, S.A., Van Berkel, G.J. and Glish, G.L. (1992) Tandem mass spectrometry of small, multiply charged oligonucleotides. J. Am. Soc. Mass Spectrom., 3, 60–70.

34. Ando, D., Rashad, S., Begley, T.J., et al. (2025) Decoding codon bias: the role of tRNA modifications in tissue-specific translation. Int. J. Mol. Sci., 26, 706.

35. Brandmayr, C., Wagner, M., Brückl, T., et al. (2012) Isotope-based analysis of modified tRNA nucleosides correlates modification density with translational efficiency. Angew. Chem. Int. Ed Engl., 51, 11162–11165.

36. Chan, C., Pham, P., Dedon, P.C., et al. (2018) Lifestyle modifications: coordinating the tRNA epitranscriptome with codon bias to adapt translation during stress responses. Genome Biol., 19, 228.

37. Dedon, P.C. and Begley, T.J. (2022) Dysfunctional tRNA reprogramming and codon-biased translation in cancer. Trends Mol. Med., 28, 964–978.

38. Dittmar, K.A., Goodenbour, J.M. and Pan, T. (2006) Tissue-specific differences in human transfer RNA expression. PLOS Genet., 2, e221.

39. Giguère, S., Wang, X., Huber, S., et al. (2024) Antibody production relies on the tRNA inosine wobble modification to meet biased codon demand. Science, 383, 205–211.

40. Mitchener, M.M., Begley, T.J. and Dedon, P.C. (2023) Molecular coping mechanisms: reprogramming tRNAs to regulate codon-biased translation of stress response proteins. Acc. Chem. Res., 56, 3504–3514.

41. Pang, Y.L.J., Abo, R., Levine, S.S., et al. (2014) Diverse cell stresses induce unique patterns of tRNA up- and down-regulation: tRNA-seq for quantifying changes in tRNA copy number. Nucleic Acids Res., 42, e170.

42. Clark, K.D., Lee, C., Gillette, R., et al. (2021) Characterization of neuronal RNA modifications during non-associative Learning in Aplysia reveals key roles for tRNAs in behavioral sensitization. ACS Cent. Sci., 7, 1183–1190.

43. Goodenbour, J.M. and Pan, T. (2006) Diversity of tRNA genes in eukaryotes. Nucleic Acids Res., 34, 6137–6146.

44. Aplysia californica genome assembly AplCal3.0 NCBI.

45. Knudsen, B., Kohn, A.B., Nahir, B., et al. (2006) Complete DNA sequence of the mitochondrial genome of the sea-slug, Aplysia californica: conservation of the gene order in Euthyneura. Mol. Phylogenet. Evol., 38, 459–469.

46. Berg, O.G. and Kurland, C.G. (1997) Growth rate-optimised tRNA abundance and codon usage1. J. Mol. Biol., 270, 544–550.

47. Hill, A.M., To, K. and Wilke, C.O. (2025) Availability of charged tRNAs drives maximal protein synthesis at intermediate levels of codon usage bias. 10.1101/2025.06.16.659965.

48. Ikemura, T. (1981) Correlation between the abundance of Escherichia coli transfer RNAs and the occurrence of the respective codons in its protein genes. J. Mol. Biol., 146, 1–21.

49. Silva, H.G.S., Kimura, S., Lima, P.L.C., et al. (2025) Integrating tRNA gene epigenomics and expression with codon usage unravels an intricate connection with translatome dynamics in Trypanosoma cruzi. mBio, 16, e01622–25.

50. Barciszewska, M. and Barciszewski, J. (1988) Yellow lupin cytoplasmic tRNAGlu is not a cofactor in chlorophyll biosynthesis. Mol. Biol. Rep., 13, 11–14.

51. Peterson, D., Schön, A. and Söll, D. (1988) The nucleotide sequences of barley cytoplasmic glutamate transfer RNAs and structural features essential for formation of δ-aminolevulinic acid. Plant Mol. Biol., 11, 293–299.

52. Hedgcoth, C., Hayenga, K., Harrison, M., et al. (1984) Lysine tRNAs from rat liver: lysine tRNA sequences are highly conserved. Nucleic Acids Res., 12, 2535–2541.

53. Madison, J.T. and Boguslawski, S.J. (1974) Partial digestion of a yeast lysine transfer ribonucleic acid and reconstruction of the nucleotide sequence. Biochemistry, 13, 524– 527.

54. Raba, M., Limburg, K., Burghagen, M., et al. (1979) Nucleotide sequence of three isoaccepting lysine tRNAs from rabbit liver and SV40-transformed mouse fibroblasts. Eur. J. Biochem., 97, 305–318.

55. Iben, J.R. and Maraia, R.J. (2014) tRNA gene copy number variation in humans. Gene, 536, 376–384.

56. McDonald, M.J., Chou, C.-H., Swamy, K.B., et al. (2015) The evolutionary dynamics of tRNA-gene copy number and codon-use in E. coli. BMC Evol. Biol., 15, 163.

57. Bugga, P., Asthana, V. and Drezek, R. (2024) Simulation-guided tunable DNA probe design for mismatch tolerant hybridization. PLOS ONE, 19, e0305002.

58. Lei, H.-T., Wang, Z.-H., Li, B., et al. (2023) tModBase: deciphering the landscape of tRNA modifications and their dynamic changes from epitranscriptome data. Nucleic Acids Res., 51, D315–D327.

59. Smardo, F.L. and Calvet, J.P. (1987) Sequence analysis of the glutamate tRNA family: evidence for pseudogenes. Gene, 57, 213–220.

60. Smardo, F.L. and Calvet, J.P. (1987) Human glutamate tRNA forms stable hybrids in vitro with 28S ribosomal RNA. Nucleic Acids Res., 15, 661–681.

61. Lyons, S.M., Fay, M.M. and Ivanov, P. (2018) The role of RNA modifications in the regulation of tRNA cleavage. FEBS Lett., 592, 2828–2844.

62. Kang, B., Miyauchi, K., Matuszewski, M., et al. (2017) Identification of 2-methylthio cyclic N6-threonylcarbamoyladenosine (ms2ct6A) as a novel RNA modification at position 37 of tRNAs. Nucleic Acids Res., 45, 2124–2136.

63. Tomikawa, C. (2018) 7-methylguanosine modifications in transfer RNA (tRNA). Int. J. Mol. Sci., 19, 4080.

64. Armengaud, J., Urbonavicius, J., Fernandez, B., et al. (2004) N2-methylation of guanosine at position 10 in tRNA is catalyzed by a THUMP domain-containing, S-adenosylmethionine-dependent methyltransferase, conserved in Archaea and Eukaryota. J. Biol. Chem., 279, 37142–37152.

65. Purushothaman, S.K., Bujnicki, J.M., Grosjean, H., et al. (2005) Trm11p and Trm112p Are both Required for the Formation of 2-Methylguanosine at Position 10 in Yeast tRNA. Mol. Cell. Biol., 25, 4359–4370.

66. Chan, C.T.Y., Pang, Y.L.J., Deng, W., et al. (2012) Reprogramming of tRNA modifications controls the oxidative stress response by codon-biased translation of proteins. Nat. Commun., 3, 937.

67. Chen, X., Yuan, Y., Zhou, F., et al. (2025) RNA m5C modification: from physiology to pathology and its biological significance. Front. Immunol., 16.

68. Liu, F., Clark, W., Luo, G., et al. (2016) ALKBH1-mediated tRNA demethylation regulates translation. Cell, 167, 816–828.e16.

69. Lu, Y., Yang, L., Feng, Q., et al. (2024) RNA 5-methylcytosine modification: regulatory molecules, biological functions, and human diseases. Genomics Proteomics Bioinformatics, 22, qzae063.

70. Zhang, C. and Jia, G. (2018) Reversible RNA modification N1-methyladenosine (m1A) in mRNA and tRNA. Genomics Proteomics Bioinformatics, 16, 155–161.

71. Dyubankova, N., Sochacka, E., Kraszewska, K., et al. (2015) Contribution of dihydrouridine in folding of the D-arm in tRNA. Org. Biomol. Chem., 13, 4960–4966.

72. Finet, O., Yague-Sanz, C., Marchand, F., et al. (2022) The Dihydrouridine landscape from tRNA to mRNA: a perspective on synthesis, structural impact and function. RNA Biol., 19, 735–750.

73. Rider, L.W., Ottosen, M.B., Gattis, S.G., et al. (2009) Mechanism of dihydrouridine synthase 2 from yeast and the importance of modifications for efficient tRNA reduction. J. Biol. Chem., 284, 10324–10333.

74. Draycott, A.S., Schaening-Burgos, C., Rojas-Duran, M.F., et al. (2022) Transcriptome-wide mapping reveals a diverse dihydrouridine landscape including mRNA. PLOS Biol., 20, e3001622.

75. Ji, J., Yu, N.J. and Kleiner, R.E. (2024) Sequence- and structure-specific tRNA dihydrouridylation by hDUS2. ACS Cent. Sci., 10, 803–812.

76. Ju, C.-W., Li, H., Jiang, B., et al. (2025) Quantitative CRACI reveals transcriptome-wide distribution of RNA dihydrouridine at base resolution. Nat. Commun., 16, 8863.

77. Schultz, S.K., Hossain, N., Barnes, L., et al. (2025) The tRNA dihydrouridine synthase DusA has a distinct mechanism in optimizing tRNAs for translation. 10.1101/2025.08.28.672980.

78. Matsuo, M., Abe, Y., Saruta, Y., et al. (1995) Mollusk genes encoding lysine tRNA(UUU) contain introns. Gene, 165, 249–253.

79. Matsuo, M., Yokogawa, T., Nishikawa, K., et al. (1995) Highly specific and efficient cleavage of squid tRNALys catalyzed by magnesium ions. J. Biol. Chem., 270, 10097–10104.

80. Liu, B., Cao, J., Wang, X., et al. (2021) Deciphering the tRNA-derived small RNAs: origin, development, and future. Cell Death Dis., 13, 24.

81. Yu, X., Xie, Y., Zhang, S., et al. (2021) tRNA-derived fragments: mechanisms underlying their regulation of gene expression and potential applications as therapeutic targets in cancers and virus infections. Theranostics, 11, 461–469.

82. Potapov, V., Fu, X., Dai, N., et al. (2018) Base modifications affecting RNA polymerase and reverse transcriptase fidelity. Nucleic Acids Res., 46, 5753–5763.

83. Tepe, M.L., Chen, Y., Carso, A., et al. (2025) MapID-based quantitative mapping of chemical modifications and expression of human transfer RNA. Cell Chem. Biol., 32, 752–766.e7.

84. Shigematsu, M., Matsubara, R., Gumas, J., et al. (2025) Angiogenin-catalyzed cleavage within tRNA anticodon-loops identified by cP-RNA-seq. Biosci. Biotechnol. Biochem., 89, 398– 405.

85. Thompson, J.E., Venegas, F.D. and Raines, R.T. (1994) Energetics of catalysis by ribonucleases: fate of the 2’, 3’-cyclic phosphodiester intermediate. Biochemistry, 33, 7408–7414.

86. Gustafsson, H.T., Ferguson, L., Galan, C., et al. (2025) Deep sequencing of yeast and mouse tRNAs and tRNA fragments using OTTR. eLife, 14, e77616.

87. Scacchetti, A., Shields, E.J., Trigg, N.A., et al. (2024) A ligation-independent sequencing method reveals tRNA-derived RNAs with blocked 3′ termini. Mol. Cell, 84, 3843–3859.e8.

88. Fu, H., Feng, J., Liu, Q., et al. (2009) Stress induces tRNA cleavage by angiogenin in mammalian cells. FEBS Lett., 583, 437–442.

89. Yamasaki, S., Ivanov, P., Hu, G., et al. (2009) Angiogenin cleaves tRNA and promotes stress-induced translational repression. J. Cell Biol., 185, 35–42.

90. Honda, S., Loher, P., Shigematsu, M., et al. (2015) Sex hormone-dependent tRNA halves enhance cell proliferation in breast and prostate cancers. Proc. Natl. Acad. Sci., 112, E3816–E3825.

91. Pawar, K., Shigematsu, M., Sharbati, S., et al. (2020) Infection-induced 5′-half molecules of tRNAHisGUG activate Toll-like receptor 7. PLOS Biol., 18, e3000982.

92. Raines, R.T. (1998) Ribonuclease A. Chem. Rev., 98, 1045–1066.

93. Russo, N., Acharya, K.R., Vallee, B.L., et al. (1996) A combined kinetic and modeling study of the catalytic center subsites of human angiogenin. Proc. Natl. Acad. Sci., 93, 804–808.

94. Goo, S.M. and Cho, S. (2013) The expansion and functional diversification of the mammalian ribonuclease A superfamily epitomizes the efficiency of multigene families at generating biological novelty. Genome Biol. Evol., 5, 2124–2140.

95. Blanco, S., Dietmann, S., Flores, J.V., et al. (2014) Aberrant methylation of tRNAs links cellular stress to neuro-developmental disorders. EMBO J., 33, 2020–2039.

96. Schaefer, M., Pollex, T., Hanna, K., et al. (2010) RNA methylation by Dnmt2 protects transfer RNAs against stress-induced cleavage. Genes Dev., 24, 1590–1595.

97. Tuorto, F., Liebers, R., Musch, T., et al. (2012) RNA cytosine methylation by Dnmt2 and NSun2 promotes tRNA stability and protein synthesis. Nat. Struct. Mol. Biol., 19, 900– 905.

98. Wang, X., Matuszek, Z., Huang, Y., et al. (2018) Queuosine modification protects cognate tRNAs against ribonuclease cleavage. RNA, 24, 1305–1313.

99. Ogawa, T., Tomita, K., Ueda, T., et al. (1999) A cytotoxic ribonuclease targeting specific transfer RNA anticodons. Science, 283, 2097–2100.

100. Lu, J., Esberg, A., Huang, B., et al. (2008) Kluyveromyces lactis γ-toxin, a ribonuclease that recognizes the anticodon stem loop of tRNA. Nucleic Acids Res., 36, 1072–1080.

101. Anderson, P. and Ivanov, P. (2014) tRNA fragments in human health and disease. FEBS Lett., 588, 4297–4304.

102. Ivanov, P., Emara, M.M., Villen, J., et al. (2011) Angiogenin-induced tRNA fragments inhibit translation initiation. Mol. Cell, 43, 613–623.

103. Ivanov, P., O’Day, E., Emara, M.M., et al. (2014) G-quadruplex structures contribute to the neuroprotective effects of angiogenin-induced tRNA fragments. Proc. Natl. Acad. Sci. U. S. A., 111, 18201–18206.

104. Jackowiak, P., Hojka-Osinska, A., Gasiorek, K., et al. (2017) Effects of G-quadruplex topology on translational inhibition by tRNA fragments in mammalian and plant systems in vitro. Int. J. Biochem. Cell Biol., 92, 148–154.

105. Lyons, S.M., Gudanis, D., Coyne, S.M., et al. (2017) Identification of functional tetramolecular RNA G-quadruplexes derived from transfer RNAs. Nat. Commun., 8, 1127.

106. Lyons, S.M., Kharel, P., Akiyama, Y., et al. (2020) eIF4G has intrinsic G-quadruplex binding activity that is required for tiRNA function. Nucleic Acids Res., 48, 6223–6233.

107. Carbon, J., David, H. and Studier, M.H. (1968) Thiobases in Escherichia coli transfer RNA: 2-thiocytosine and 5-methylaminomethyl-2-thiouracil. Science, 161, 1146–1147.

108. Yamada, Y., Saneyoshi, M. and Nishimura, S. (1970) Isolation and characterization of 2-thiocytidine from a serine transfer ribonucleic acid of Escherichia coli. FEBS Lett., 7, 207–210.

109. Jäger, G., Leipuviene, R., Pollard, M.G., et al. (2004) The conserved cys-X1-X2-cys motif present in the TtcA protein Is required for the thiolation of cytidine in position 32 of tRNA from Salmonella enterica serovar Typhimurium. J. Bacteriol., 186, 750–757.

110. Nilsson, K., Lundgren, H.K., Hagervall, T.G., et al. (2002) The cysteine desulfurase IscS is required for synthesis of all five thiolated nucleosides present in tRNA from Salmonella enterica serovar Typhimurium. J. Bacteriol., 184, 6830–6835.

111. Best, A.N. (1978) Composition and Characterization of tRNA from Methanococcus vannielii. J. Bacteriol., 133, 240–250.

112. McCloskey, J.A., Graham, D.E., Zhou, S., et al. (2001) Post-transcriptional modification in archaeal tRNAs: identities and phylogenetic relations of nucleotides from mesophilic and hyperthermophilic Methanococcales. Nucleic Acids Res., 29, 4699–4706.

113. Lauhon, C.T. (2002) Requirement for IscS in biosynthesis of all thionucleosides in Escherichia coli. J. Bacteriol., 184, 6820–6829.

114. Chandramouli, K., Unciuleac, M.-C., Naik, S., et al. (2007) Formation and Properties of [4Fe-4S] Clusters on the IscU Scaffold Protein. Biochemistry, 46, 6804–6811.

115. Bouvier, D., Labessan, N., Clémancey, M., et al. (2014) TtcA a new tRNA-thioltransferase with an Fe-S cluster. Nucleic Acids Res., 42, 7960–7970.

116. Leipuviene, R., Qian, Q. and Björk, G.R. (2004) Formation of thiolated nucleosides present in tRNA from Salmonella enterica serovar Typhimurium occurs in two principally distinct pathways. J. Bacteriol., 186, 758–766.

117. Pang, Y., Wang, J., Gao, X., et al. (2023) Roles of conserved active site residues in the IscS cysteine desulfurase reaction. Front. Microbiol., 14.

118. Shi, R., Proteau, A., Villarroya, M., et al. (2010) Structural basis for Fe–S cluster assembly and tRNA thiolation mediated by IscS protein–protein interactions. PLOS Biol., 8, e1000354.

119. Lauhon, C.T., Skovran, E., Urbina, H.D., et al. (2004) Substitutions in an active site loop of Escherichia coli IscS result in specific defects in Fe-S cluster and thionucleoside biosynthesis in vivo. J. Biol. Chem., 279, 19551–19558.

120. Čavužić, M. and Liu, Y. (2017) Biosynthesis of sulfur-containing tRNA modifications: a comparison of bacterial, archaeal, and eukaryotic pathways. Biomolecules, 7, 27.

121. Romsang, A., Duang-nkern, J., Khemsom, K., et al. (2018) Pseudomonas aeruginosa ttcA encoding tRNA-thiolating protein requires an iron-sulfur cluster to participate in hydrogen peroxide-mediated stress protection and pathogenicity. Sci. Rep., 8, 11882.

122. Lalanne, J.-B., Taggart, J.C., Guo, M.S., et al. (2018) Evolutionary convergence of pathway-specific enzyme expression stoichiometry. Cell, 173, 749–761.e38.

123. Auffinger, P. and Westhof, E. (1999) Singly and bifurcated hydrogen-bonded base-pairs in tRNA anticodon hairpins and ribozymes. J. Mol. Biol., 292, 467–483.

124. Vangaveti, S., Cantara, W.A., Spears, J.L., et al. (2020) A structural basis for restricted codon recognition mediated by 2-thiocytidine in tRNA containing a wobble position inosine. J. Mol. Biol., 432, 913–929.

125. Schaffrath, R. and Leidel, S.A. (2017) Wobble uridine modifications–a reason to live, a reason to die⁈. RNA Biol., 14, 1209–1222.

126. Su, C., Jin, M. and Zhang, W. (2022) Conservation and diversification of tRNA t6A-modifying enzymes across the three domains of Life. Int. J. Mol. Sci., 23, 13600.

127. Altwegg, M. and Kubli, E. (1980) The nucleotide sequence of glutamate tRNA4 of Drosophila melanogaster. Nucleic Acids Res., 8, 215–223.

128. Andachi, Y., Yamao, F., Muto, A., et al. (1989) Codon recognition patterns as deduced from sequences of the complete set of transfer RNA species in Mycoplasma capricolum. J. Mol. Biol., 209, 37–54.

129. Chan, J.C., Yang, J.A., Dunn, M.J., et al. (1982) The nucleotide sequence of a glutamine tRNA from rat liver. Nucleic Acids Res., 10, 3755–3758.

130. Sakurai, M., Ohtsuki, T., Suzuki, T., et al. (2005) Unusual usage of wobble modifications in mitochondrial tRNAs of the nematode Ascaris suum. FEBS Lett., 579, 2767–2772.

131. Wong, T.W., McCutchan, T., Kohli, J., et al. (1979) The nucleotide sequence of the major glutamate transfer RNA from Schizosaccharomyces pombe. Nucleic Acids Res., 6, 2057– 2068.

132. Koutmou, K.S., Schuller, A.P., Brunelle, J.L., et al. (2015) Ribosomes slide on lysine-encoding homopolymeric A stretches. eLife, 4, e05534.

