## Supplementary material for "Sub-stoichiometric modifications of *Aplysia californica* tRNAs and tRNA fragments revealed by integrating intact and bottom-up mass spectrometry": Supplementary_material_1_LCMS_method_intact_analysis.docx

| **Table S1. sample information for RNA-seq datasets used for codon frequency analysis** | | |
| --- | --- | --- |
| **SRA ID** | **BioSample ID** | **tissue** |
| SRR26488150 | SAMN37934075 | pleural-pedal ganglia |
| SRR26488151 | SAMN37934074 | pleural-pedal ganglia |
| SRR26488153 | SAMN37934072 | pleural-pedal ganglia |
| SRR26488155 | SAMN37934115 | pleural-pedal ganglia |
| SRR26488156 | SAMN37934114 | pleural-pedal ganglia |
| SRR26488157 | SAMN37934113 | pleural-pedal ganglia |
| SRR26488158 | SAMN37934112 | pleural-pedal ganglia |
| SRR26488159 | SAMN37934111 | pleural-pedal ganglia |
| SRR26488160 | SAMN37934110 | pleural-pedal ganglia |
| SRR26488161 | SAMN37934109 | pleural-pedal ganglia |
| SRR17649334 | SAMN24439644 | gills |
| SRR17649337 | SAMN24439641 | hepatopancreas |
| SRR17649339 | SAMN24439639 | salivary gland |
| SRR17649338 | SAMN24439640 | digestive system |
| SRR17649336 | SAMN24439642 | heart |

| **Table S2. Top ten most frequent codons in *Aplysia californica* transcriptome** | | | | |
| --- | --- | --- | --- | --- |
| **gills** | **hepatopancreas** | **salivary gland** | **digestive system** | **heart** |
| GAG | GAG | GAG | GAG | GAG |
| GAC | GAC | AAG | GAC | GAC |
| AAG | AAG | GAC | AAG | AAG |
| CTG | CTG | CTG | CTG | CTG |
| CAG | CAG | CAG | CAG | CAG |
| GTG | GTG | AAA | GTG | AAA |
| AAA | AAA | GTG | AAA | GTG |
| AAC | AAC | GAA | AAC | GAA |
| GAA | GAA | AAC | GAA | AAC |
| ATG | ATG | ATG | ATG | ATG |

| **Table S3. Biotinylated DNA probes for tRNA pull-down** | |
| --- | --- |
| **sequence name** | **sequence** |
| *A.c* tRNA_Glu_CUC_3'-probe | /5Biosg/TTCCCAAGCGGGGAATCGAACCCCGGCCGT |
| *A.c* tRNA_Lys_UUU_3'-probe | /5Biosg/CGCCCAATAAGGGACTCGAACCCTTGACCC |

| **Table S4. Liquid chromatography parameters for analysis of *Aplysia* *californica* individual tRNAs and their enzymatic digestion products (RNA oligos)** | |
| --- | --- |
| HPLC system | Agilent 1260 infinity II HPLC |
| Column | XBridge BEH C18 Column, 130Å, 3.5 µm, 4.6 mm × 100 mm (Waters) |
| UV wavelength | 260 nm (diode array detector) |
| UV peak width | >0.05 min (1 s response time; 5 Hz) |
| Column compartment temperature | 50 °C |
| Flow rate | 0.3 mL/min |
| Mobile phase A | 10 mM ammonium acetate in LC-MS grade water, pH = 7.0 |
| Mobile phase B | 100% Acetonitrile |
| Gradient | 0.00-10.00 min: 0.0% B  10.00-30.00 min: 0.0-10.0% B  30.00-35.00 min: 10.0-50.0% B  35.00-40.00 min: 50.0-50.0% B  40.00-45.00 min: 50.0-0.0% B |
| Maximum pressure | 400 bar |
| Post time | 10 min |

| **Table S5. Mass spectrometry parameters for analysis of *Aplysia* *californica* individual tRNAs and their enzymatic digestion products (RNA oligos)** | |
| --- | --- |
| Mass spectrometer | Agilent 6530B Accurate-Mass QTOF-MS system |
| Source | Agilent Dual AJS ESI |
| Ion mode | negative |
| Drying gas temperature | 350 °C |
| Drying gas flow | 12 L/min |
| Nebulizer | 30 psi |
| Sheath gas temperature | 400 |
| Sheath gas flow | 12 L/min |
| Fragmentor voltage | 180 V |
| Capillary voltage | 4500 V |
| Nozzle voltage | 2000 V |
| Skimmer | 65 V |
| Oct 1 RF Vpp | 750 V |
| MS1 m/z range | 300-3200 |
| MS1 acquisition rate | 3 spectra/s |
| MS2 m/z range | 100-3200 |
| MS2 acquisition rate | 3 spectra/s |
| collision energy formula | E=(slope) × (m/z)/100 + offset |
| charge state, slope, offset | 1, 3.3, 2  2, 3.3, 2.24  3, 3.3, -3.8  >3, 3.1, -3.46 |

| **Table S6. Liquid chromatography parameters for analysis of nucleosides from *Aplysia* *californica* individual tRNAs** | |
| --- | --- |
| HPLC system | Agilent 1260 infinity II HPLC |
| Column | ZORBAX Eclipse Plus C18 Narrow Bore RR 2.1 x 100 mm, 3.5 µm column (Agilent) |
| UV wavelength | 260 nm (diode array detector) |
| UV peak width | >0.1 min (2 s response time; 2.5 Hz) |
| Column compartment temperature | 36 °C |
| Flow rate | 0.2 mL/min |
| Mobile phase A | 5 mM ammonium acetate in LC-MS grade water, pH = 5.5 |
| Mobile phase B | 5 mM ammonium acetate in 40/60 acetonitrile/H2O |
| Gradient | 0.00 min: 1.0% B  5.00 min: 1.0% B  11.00 min: 7.0% B  13.00 min: 10.0% B  32.00 min: 15.0% B  40.00 min: 70.0% B  44.00 min: 100.0% B  50.00 min: 100.0% B  50.10 min: 1.0% B |
| Maximum pressure | 400 bar |
| Post time | 10 min |

| **Table S7. Mass spectrometry parameters for analysis of nucleosides from *Aplysia* *californica* individual tRNAs** | |
| --- | --- |
| Mass spectrometer | Agilent 6530B Accurate-Mass QTOF-MS system |
| Source | Agilent Dual AJS ESI |
| Ion mode | positive |
| Drying gas temperature | 300 °C |
| Drying gas flow | 5 L/min |
| Nebulizer | 35 psi |
| Sheath gas temperature | 325 |
| Sheath gas flow | 8 L/min |
| Fragmentor voltage | 100 V |
| Capillary voltage | 3000 V |
| Nozzle voltage | 1000 V |
| Skimmer | 65 V |
| Oct 1 RF Vpp | 400 V |
| MS1 m/z range | 200-600 |
| MS1 acquisition rate | 2 spectra/s |
| MS2 m/z range | 100-600 |
| MS2 acquisition rate | 2 spectra/s |

| **Table S8. MGF data file export parameters** | |
| --- | --- |
| Agilent MassHunter Qualitative Analysis 10.0 | |
| Export content | Entire data file |
| Qualitative method | One export file per data file |
| Maximum spike width | 2 |
| Required valley | 70 |
| Height filters options | Uncheck |
| Maximum number of peaks | Uncheck |
| Isotope model | Peptide |
| Charge state options | Uncheck |

| **Table S9. tRNA sequence information from Modomics database** | | | |
| --- | --- | --- | --- |
| **species** | **RNA type** | **cellular localization** | **sequence** |
| *Heterololigo bleekeri* | tRNA^Lys^ | cytoplasmic | GCCUCCAUA[L](https://genesilico.pl/modomics/modifications/191)CUCAG[D](https://genesilico.pl/modomics/modifications/190)CGGUAGAGCA[P](https://genesilico.pl/modomics/modifications/185)CAGACU[N](https://genesilico.pl/modomics/modifications/76)UU[H](https://genesilico.pl/modomics/modifications/72)A[Ѯ](https://genesilico.pl/modomics/modifications/98)CUGAGG[7](https://genesilico.pl/modomics/modifications/203)[D](https://genesilico.pl/modomics/modifications/190)[?](https://genesilico.pl/modomics/modifications/186)UGGGG[\](https://genesilico.pl/modomics/modifications/296)[P](https://genesilico.pl/modomics/modifications/185)CG[Ѣ](https://genesilico.pl/modomics/modifications/204)GUCCCCAU[L](https://genesilico.pl/modomics/modifications/191)UGGG[?](https://genesilico.pl/modomics/modifications/186)UCCA |
| *Heterololigo bleekeri* | tRNA^Lys^ | cytoplasmic | UCCCG[L](https://genesilico.pl/modomics/modifications/191)CUA[L](https://genesilico.pl/modomics/modifications/191)CUCAG[D](https://genesilico.pl/modomics/modifications/190)CGGUAGAGCACGAGA[Ѵ](https://genesilico.pl/modomics/modifications/70)UCUU[6](https://genesilico.pl/modomics/modifications/231)A[P](https://genesilico.pl/modomics/modifications/185)CUCGGG[7](https://genesilico.pl/modomics/modifications/203)[D](https://genesilico.pl/modomics/modifications/190)[?](https://genesilico.pl/modomics/modifications/186)GUGGG[\](https://genesilico.pl/modomics/modifications/296)[P](https://genesilico.pl/modomics/modifications/185)CG[Ѣ](https://genesilico.pl/modomics/modifications/204)GCCCCACGUUGGGAGCCA |
| *Heterololigo bleekeri* | tRNA^Lys^ | cytoplasmic | GCCUUCAUA[L](https://genesilico.pl/modomics/modifications/191)CUCAG[D](https://genesilico.pl/modomics/modifications/190)CGGUAGAGCA[P](https://genesilico.pl/modomics/modifications/185)CAGACU[N](https://genesilico.pl/modomics/modifications/76)UU[H](https://genesilico.pl/modomics/modifications/72)A[Ѯ](https://genesilico.pl/modomics/modifications/98)CUGAGG[7](https://genesilico.pl/modomics/modifications/203)[D](https://genesilico.pl/modomics/modifications/190)[?](https://genesilico.pl/modomics/modifications/186)UGGGG[\](https://genesilico.pl/modomics/modifications/296)[P](https://genesilico.pl/modomics/modifications/185)CG[Ѣ](https://genesilico.pl/modomics/modifications/204)GUCCCCAU[L](https://genesilico.pl/modomics/modifications/191)CGGG[?](https://genesilico.pl/modomics/modifications/186)UCCA |
| *Mus musculus* | tRNA^Glu^ | cytoplasmic | UCCCUGGUG[L](https://genesilico.pl/modomics/modifications/191)UC[P](https://genesilico.pl/modomics/modifications/185)AGUGG[D](https://genesilico.pl/modomics/modifications/190)[P](https://genesilico.pl/modomics/modifications/185)AGGAUUCGGCGCUCUCACCGCCGCGGC[??](https://genesilico.pl/modomics/modifications/186)GGG[\](https://genesilico.pl/modomics/modifications/296)[P](https://genesilico.pl/modomics/modifications/185)CGAUUCCCGGUCAGGGAACCA |
| *Homo sapiens* | tRNA^Glu^ | cytoplasmic | UCCCUGGUG[L](https://genesilico.pl/modomics/modifications/191)UC[P](https://genesilico.pl/modomics/modifications/185)AGUGG[D](https://genesilico.pl/modomics/modifications/190)[P](https://genesilico.pl/modomics/modifications/185)AGGAUUCGGCGCUCUCACCGCCGCGGC[??](https://genesilico.pl/modomics/modifications/186)GGG[\](https://genesilico.pl/modomics/modifications/296)[P](https://genesilico.pl/modomics/modifications/185)CGAUUCCCGGUCAGGGAACCA |

References:

Matsuo,M., Yokogawa,T., Nishikawa,K., Watanabe,K. and Okada,N. (1995) Highly Specific and Efficient Cleavage of Squid tRNALys Catalyzed by Magnesium Ions (∗). *Journal of Biological Chemistry*, **270**, 10097–10104.

Matsuo,M., Abe,Y., Saruta,Y. and Okada,N. (1995) Mollusk genes encoding lysine tRNA(UUU) contain introns. *Gene*, **165**, 249–253.

Smardo,F.L. and Calvet,J.P. (1987) Sequence analysis of the glutamate tRNA family: evidence for pseudogenes. *Gene*, **57**, 213–220.

Smardo,F.L. and Calvet,J.P. (1987) Human glutamate tRNA forms stable hybrids in vitro with 28S ribosomal RNA. *Nucleic Acids Res*, **15**, 661–681.

| **Table S10. tRNA information for intact mass analysis** | | | | |
| --- | --- | --- | --- | --- |
| **tRNA type** | **length** | **sequence** | **formula and monoisotopic mass*** | **highest isotopic peak* at specific charge** |
| tRNA^Glu^CUC, -CCA tail, methyl (-), | 75 nt | 5'-P-UCCCA AGUGL(K) UCUAG DGGDD AGGAU UUCGC G?UCU CACCG CGACG GC?GG GG\PC G**"**UUC CCCGC UUGGG AACCA-OH-3' | C715H902N275O533P75,  24188.225 | *m/z* = 1208.904 or 1208.954  *z* = -20 |
| tRNA^Glu^CUC,  -CCA tail, methyl (+) | 75 nt | 5'-P-UCCCA AGUGL(K) UCUAG DGGDD AGGAU UUCGC G?UCU CACCG CGACG G??GG GG\PC G**"**UUC CCCGC UUGGG AACCA-OH-3' | C716H904N275O533P75,  24202.241 | *m/z* = 1209.605 or 1209.655  *z* = -20 |
| tRNA^Glu^CUC,  -CC tail, methyl (-) | 74 nt | 5'-P-UCCCA AGUGL(K) UCUAG DGGDD AGGAU UUCGC G?UCU CACCG CGACG GC?GG GG\PC G**"**UUC CCCGC UUGGG AACC-OH-3' | C705H890N270O527P74,  23859.173 | *m/z =* 1192.452  *z = -*20 |
| tRNA^Glu^CUC,  no tail, methyl (-) | 72 nt | 5'-P-UCCCA AGUGL(K) UCUAG DGGDD AGGAU UUCGC G?UCU CACCG CGACG GC?GG GG\PC G**"**UUC CCCGC UUGGG AA-OH-3' | C687H866N264O513P72,  23249.090 | *m/z* = 1161.947  *z* = -20 |
| tRNA^Lys^UUU | 76 nt | 5'-P-GCCCG GUUAL(K) CUCAG DCGGU AGAGC AUCAG A%U3U U[AU? UGAGG 7D?AA GGG\P CG"GU CCCUU AUUGG GCGCC A-OH-3' | C738H925N285O542S3P76, 24898.280 | *m/z* = 1244.457  *z* = -20 |

*Monoisotopic mass was calculated by <https://www.sisweb.com/referenc/tools/exactmass.htm>? Isotopic envelope pattern was simulated by <https://www.envipat.eawag.ch/index.php> The highest isotopic peak refers to the highest peak in an isotopic envelope at a specific charge state.

**Figure S1. Extracted ion chromatogram for tRNA^Glu^CUC intact mass analysis**

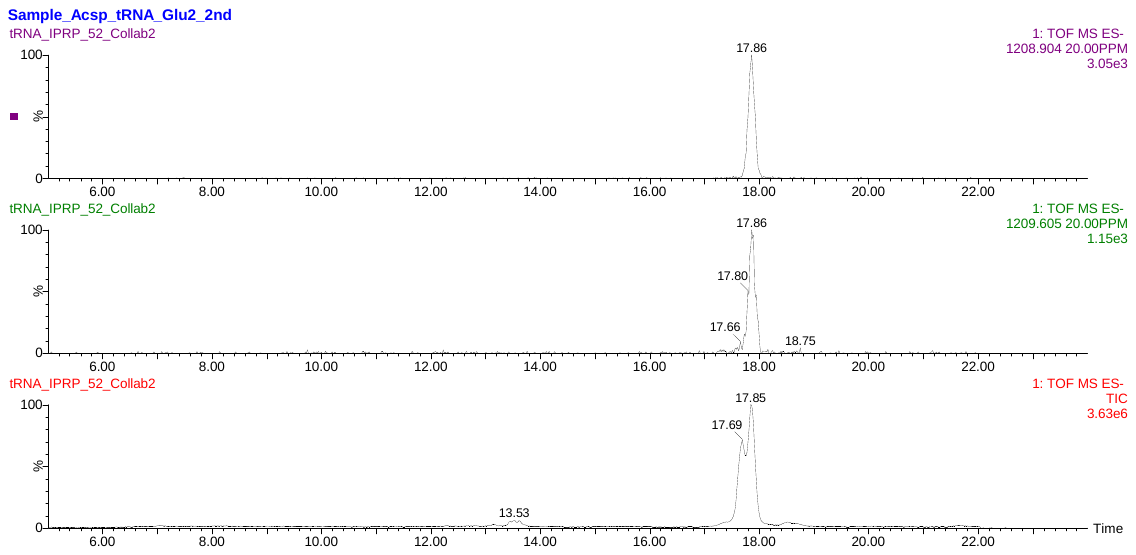

TIC

tRNA^Glu^CUC, -CCA tail, methyl (+)

tRNA^Glu^CUC, -CCA tail, methyl (-)

A

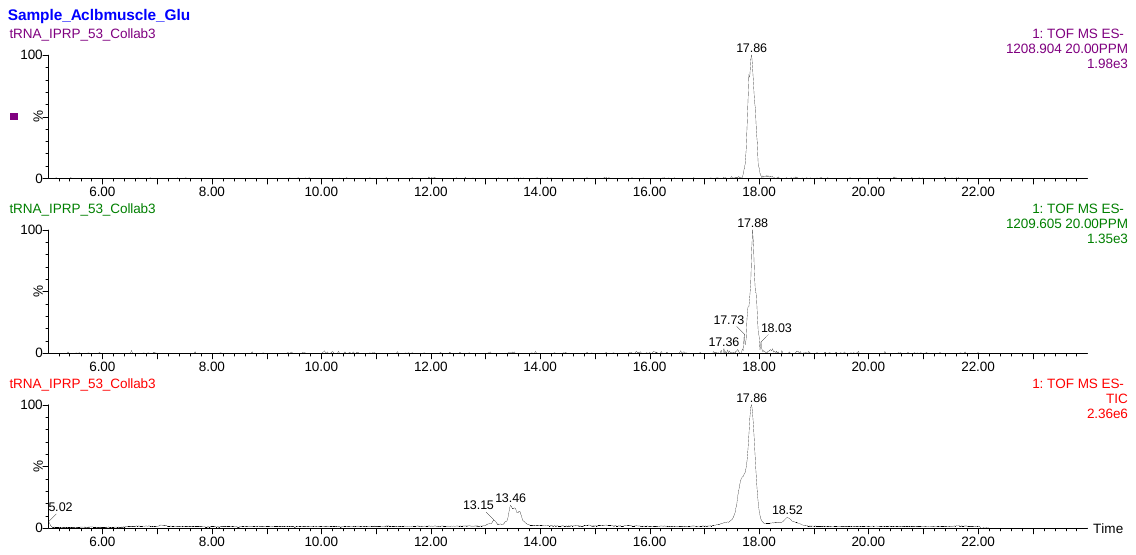

TIC

tRNA^Glu^CUC, -CCA tail, methyl (-)

tRNA^Glu^CUC, -CCA tail, methyl (+)

B

**Figure S1. Extracted ion chromatogram for tRNA^Glu^CUC intact mass analysis**

(A): Extracted ion chromatograms for the spermatheca tRNA^Glu^CUC sample (Collab2). Upper panel: full length tRNA^Glu^CUC, methyl (-), *m/z* = 1208.904, *z* = -20. Middle panel: methyl (+), *m/z* = 1209.605, *z* = -20. Lower panel: total ion chromatogram. (B): Extracted ion chromatograms for the muscle tRNA^Glu^CUC sample (Collab3). Upper panel: full length tRNA^Glu^CUC, methyl (-), *m/z* = 1208.904, *z* = -20. Middle panel: methyl (+), *m/z* = 1209.605, *z* = -20. Lower panel: total ion chromatogram.

**Figure S2.** **Mass spectra for full length tRNA^Glu^CUC intact mass analysis**

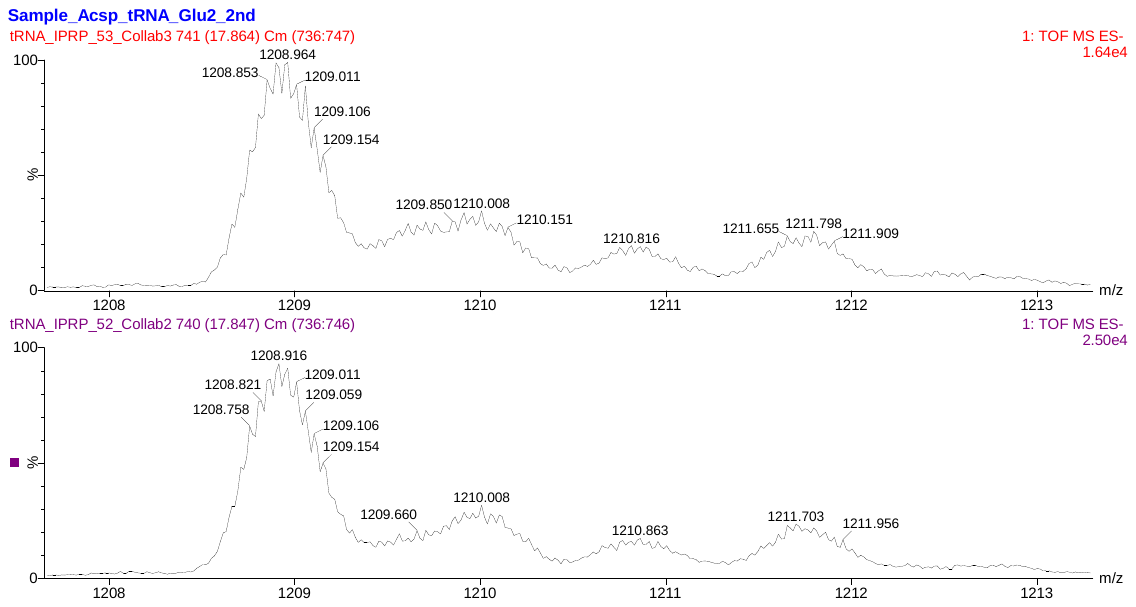

M+Na

M+Na

methyl (+)

methyl (-)

methyl (+)

methyl (-)

**Figure S2.** **Mass spectra for full length tRNA^Glu^CUC intact mass analysis**

Averaged mass spectra from scan 736 to 747 (upper panel, muscle sample) and 736 to 746 (lower panel, spermatheca sample). The isotopic envelopes are labelled: methyl (-) corresponds to *z* = -20 for monoisotopic mass = 24188.225; methyl (+) corresponds to *z* = -20 for monoisotopic mass = 24202.241; M+Na corresponds to the sodium adduct. Note that the isotopic envelopes of methyl (-) and M+Na overlapped.

**Figure S3. Extracted ion chromatograms for tRNA^Lys^UUU intact mass analysis**

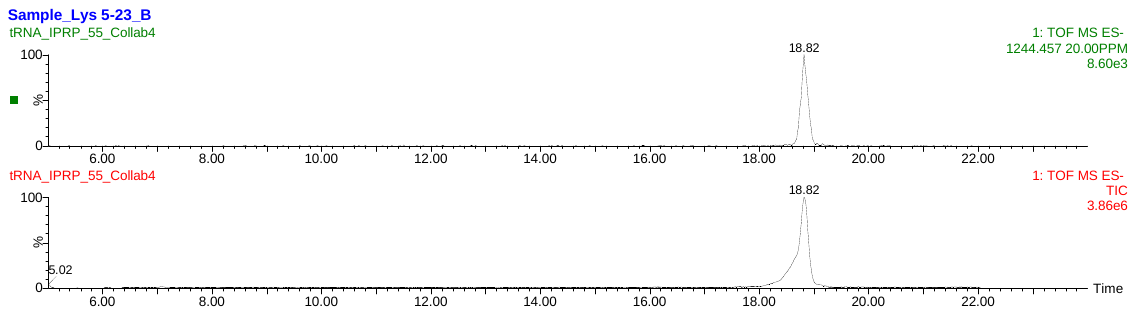

A

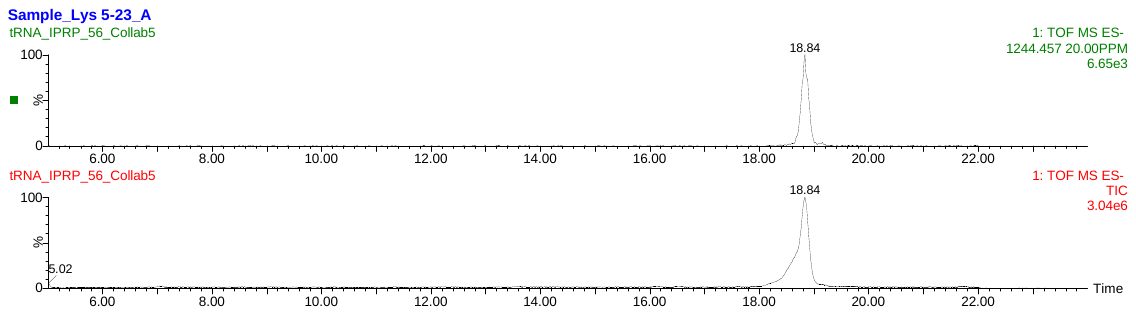

B

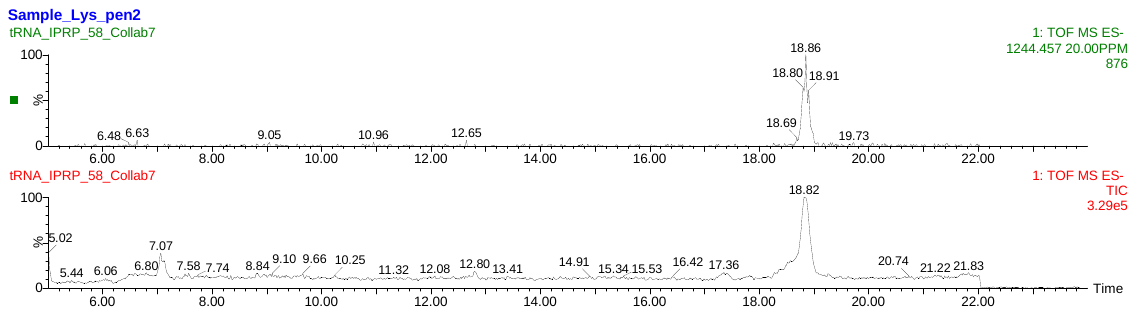

C

**Figure S3. Extracted ion chromatograms for tRNA^Lys^UUU intact mass analysis**

Extracted ion and total ion chromatograms for the tRNA^Lys^UUU samples, for *m/z* = 1244.457, *z* = -20. (A) and (B): pooled samples of heart + penis. (C): reproductive organ sample.

**Figure S4.** **Mass spectra for full length tRNA^Lys^UUU intact mass analysis**

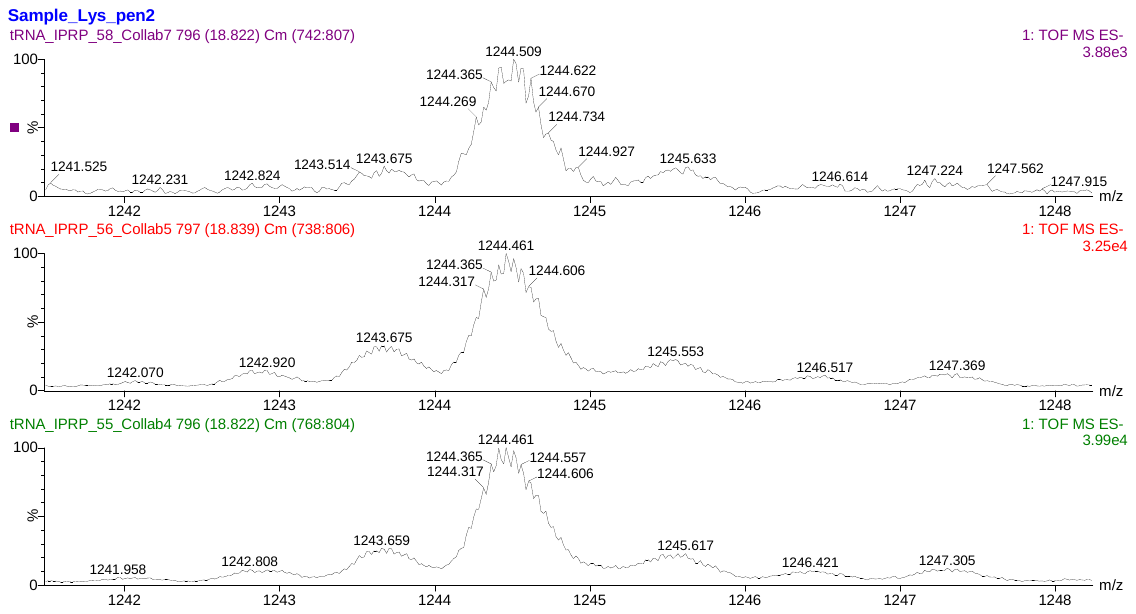

M

M+Na

M-16 Da

M-32 Da

M

M+Na

M-16 Da

M-32 Da

M-32 Da

M

M-16 Da

M+Na

M+Na

M+Na

tRNA^Lys^UUU

tRNA^Lys^UUU

**Figure S4.** **Mass spectra for full length tRNA^Lys^UUU intact mass analysis**

Averaged mass spectra for tRNA^Lys^UUU samples. Collab4 and Collab5: pooled heart + reproductive; Collab7: reproductive. For *m/z* = 1244.457, *z* = -20. The isotopic envelopes for M, M-16 Da, M-32 Da and M+Na are labeled.

| **Table S11. Deconvolution results for intact mass analysis** | |
| --- | --- |
| **Sample name** | **deconvoluted intact mass (most abundant)** |
| Collab 2-rep1 | 24188.13 Da |
| Collab 2-rep2 | 24188.33 Da |
| Collab 3-rep1 | 24188.71 Da |
| Collab 3-rep2 | 24188.79 Da |
| Collab 4 | 24898.37 Da |
| Collab 5 | 24898.37 Da |
| Collab 7 | 24898.42 Da |

| **Table S12. tsRNA^Glu^CUC information for intact mass analysis** | | | |
| --- | --- | --- | --- |
| **length and type** | **sequence (5' to 3')** | **formula and monoisotopic mass*** | **highest isotopic peak* and its charge state** |
| 37 nt methyl (-) | CG CGACG GC?GG GG\PC G"UUC CCCGC UUGGG AACCA | C355H447N141O258P36,  11926.675 | *m/z* = 1192.160  *z* = -10 |
| 37 nt methyl (+) | CG CGACG G??GG GG\PC G"UUC CCCGC UUGGG AACCA | C356H449N141O258P36,  11940.690 | *m/z =* 1193.562  *z =* -10 |
| 38 nt methyl (-) | CCG CGACG GC?GG GG\PC G"UUC CCCGC UUGGG AACCA | C364H459N144O265P37,  12231.716 | *m/z* = 1222.665  *z* = -10 |
| 38 nt methyl (+) | CCG CGACG G??GG GG\PC G"UUC CCCGC UUGGG AACCA | C365H461N144O265P37,  12245.732 | *m/z =*1224.066  *z =* -10 |
| 39 nt methyl (-) | ACCG CGACG GC?GG GG\PC G"UUC CCCGC UUGGG AACCA | C374H471N149O271P38,  12560.769 | *m/z* = 1255.570  *z* = -10 |
| 39 nt methyl (+) | ACCG CGACG G??GG GG\PC G"UUC CCCGC UUGGG AACCA | C375H473N149O271P38,  12574.784 | *m/z =* 1256.971  *z* = -10 |
| 40 nt methyl (-) | CACCG CGACG GC?GG GG\PC G"UUC CCCGC UUGGG AACCA | C383H483N152O278P39,  12865.810 | *m/z* = 1286.074  *z* = -10 |
| 40 nt methyl (+) | CACCG CGACG G??GG GG\PC G"UUC CCCGC UUGGG AACCA | C384H485N152O278P39,  12879.826 | *m/z* = 1287.475  *z* = -10 |
| 41 nt methyl (-) | UCACCGCGAC GGC?GGGG\P CG"UUCCCCG CUUGGGAACC A | C392H494N154O286P40,  13171.835156 | *m/z* = 1316.676  *z* = -10 |
| 41 nt methyl (+) | UCACCGCGAC GG??GGGG\P CG"UUCCCCG CUUGGGAACC A | C393H496N154O286P40,  13185.851 | *m/z =* 1318.078  *z* = -10 |
| 42 nt methyl (-) | CUCACCGCGA CGGC?GGGG\ PCG"UUCCCC GCUUGGGAAC CA | C401H506N157O293P41,  13476.876446 | *m/z* = 1347.281  *z* = -10 |
| 42 nt methyl (+) | CUCACCGCGA CGG??GGGG\ PCG"UUCCCC GCUUGGGAAC CA | C402H508N157O293P41,  13490.892 | *m/z =* 1348.682  *z* = -10 |

*Monoisotopic mass was calculated by <https://www.sisweb.com/referenc/tools/exactmass.htm>?; Isotopic envelope pattern was simulated by <https://www.envipat.eawag.ch/index.php> The highest isotopic peak refers to the highest peak in an isotopic envelope at a specific charge state.

**Figure S5. Extracted ion chromatograms for tsRNA^Glu^CUC**

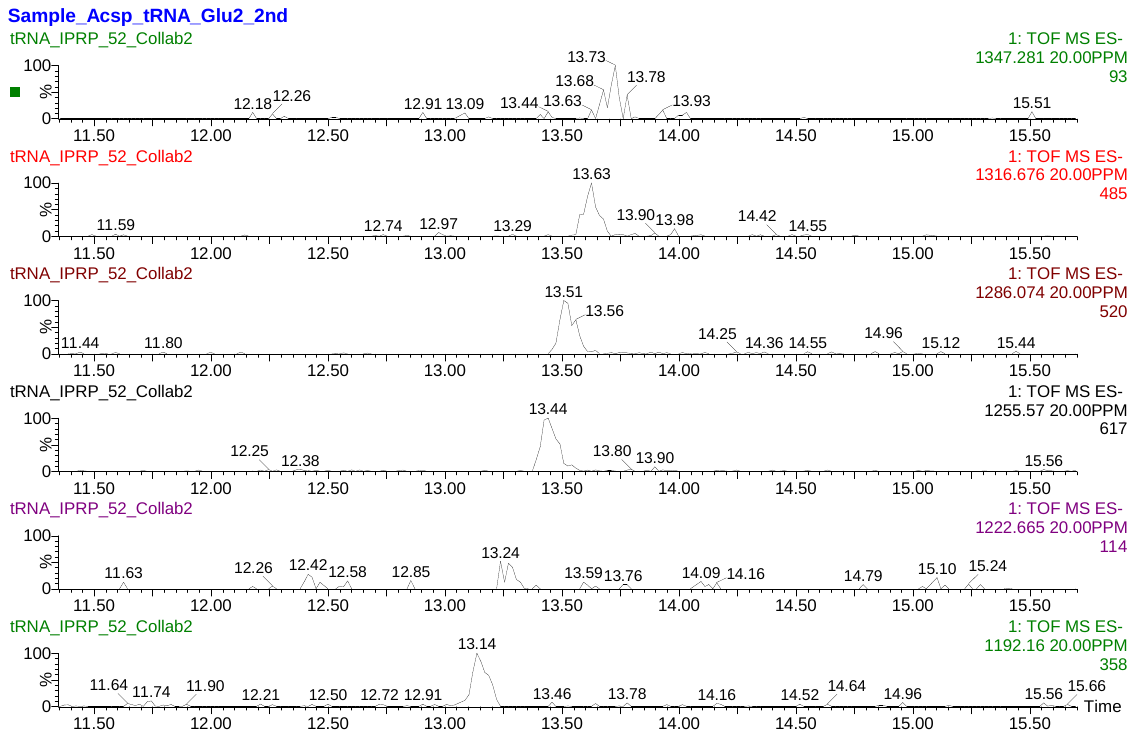

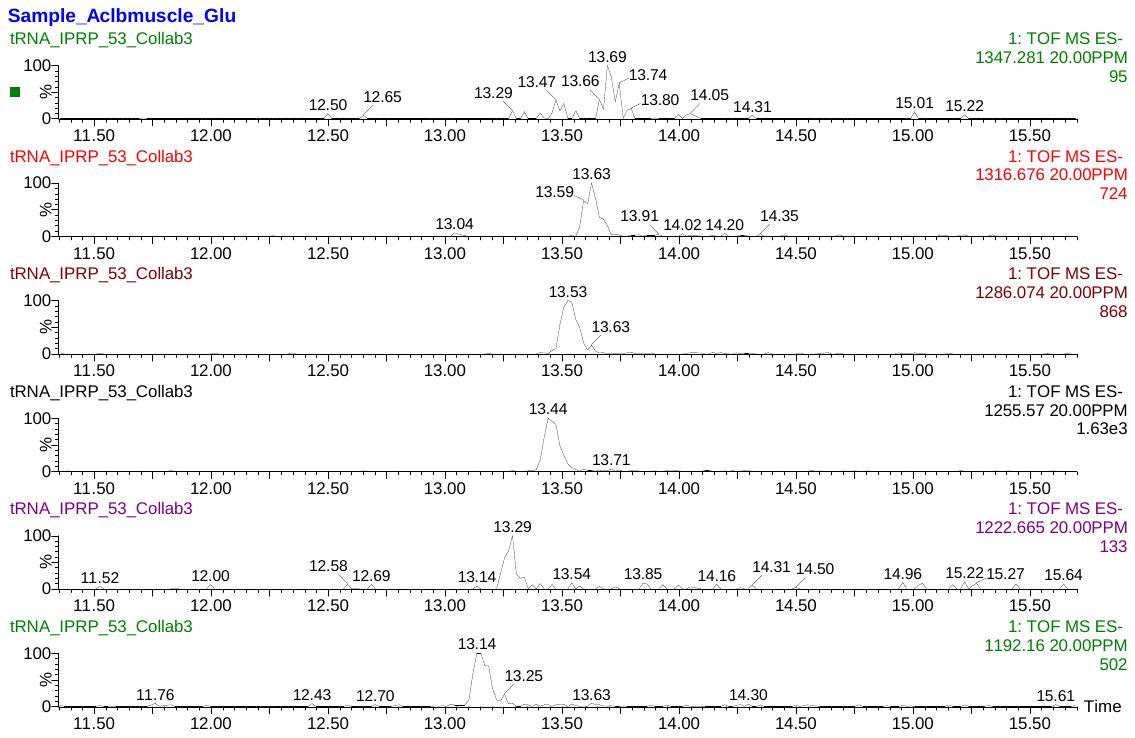

**Figure S5. Extracted ion chromatograms for tsRNA^Glu^CUC**

EICs for methyl (-) tsRNAs. Upper panel: spermatheca sample (Collab2). Lower panel: muscle sample (Collab3). See Table S11 for correspondence with tsRNA length. Horizontal axis (time) was zoomed in to enhance visualization of these EIC peaks.

| **Table S13. tsRNA^Lys^UUU information for intact mass analysis** | | | |
| --- | --- | --- | --- |
| **length** | **sequence (5' to 3')** | **formula and monoisotopic mass*** | **highest isotopic peak* and its charge state** |
| 39 nt | AU? UGA GG7 D?A AGG G\P CG" GUC CCU UAU UGG GCG CCA | C377H473N147O274P38, 12618.763 | *m/z* = 1261.369,  *z* = -10 |
| 40 nt | [AU? UGA GG7 D?A AGG G\P CG" GUC CCU UAU UGG GCG CCA | C393H494N153O284SP39, 13138.841 | *m/z* = 1313.377 or 1313.477,  *z* = -10 |
| 41 nt | U[AU?UGAGG 7D?AAGGG\P CG"GUCCCUU AUUGGGCGCC A | C402H505N155O292SP40,  13444.866 | *m/z* = 1344.080,  *z* = -10 |
| 42 nt | UU[AU?UGAGG 7D?AAGGG\P CG"GUCCCUU AUUGGGCGCC A | C411H516N157O300SP41,  13750.891 | *m/z =* 1374.682,  *z =* -10 |
| 43 nt | 3UU[AU?UGAGG 7D?AAGGG\P CG"GUCCCUU AUUGGGCGCC A | C423H531N159O309S2P42,  14144.915 | *m/z =* 1414.085,  *z =* -10 |

*Monoisotopic mass was calculated by <https://www.sisweb.com/referenc/tools/exactmass.htm>?; Isotopic envelope pattern was simulated by <https://www.envipat.eawag.ch/index.php> The highest isotopic peak refers to the highest peak in an isotopic envelope at a specific charge state.

**Figure S6. Extracted ion chromatograms for tsRNA^Lys^UUU**

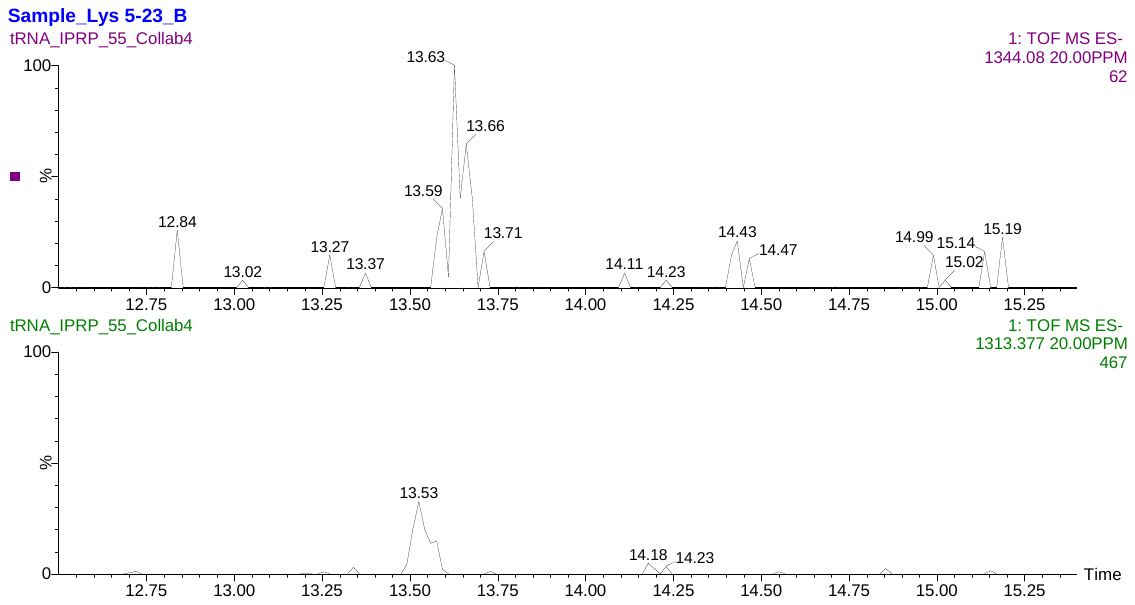

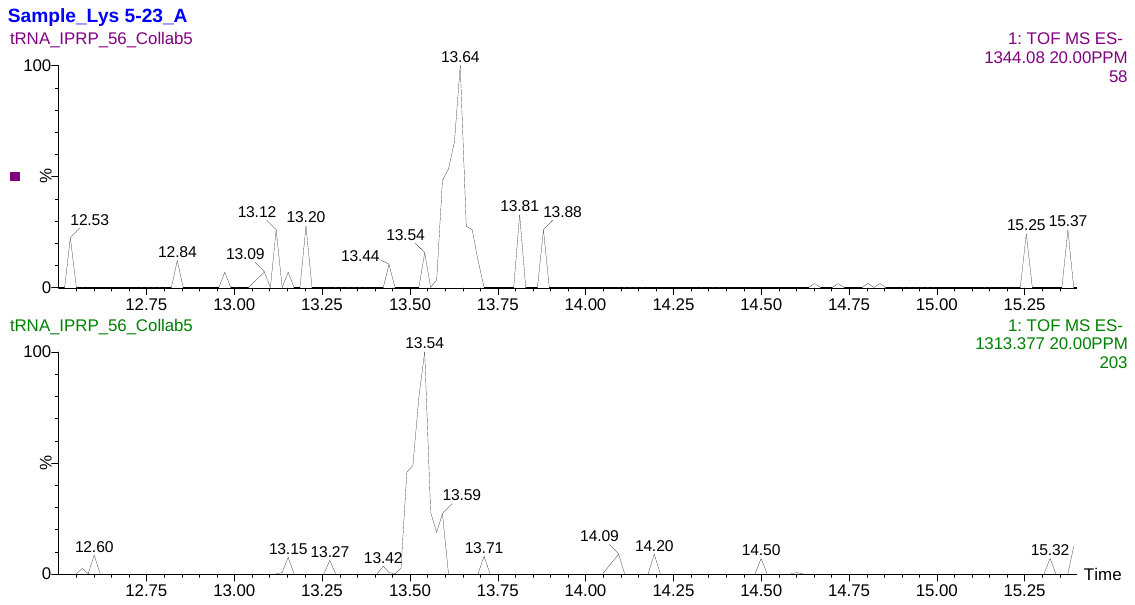

**Figure S6. Extracted ion chromatograms for tsRNA^Lys^UUU**

EICs for tsRNA^Lys^UUU of 40 and 41 nt. Upper part: reproductive + heart pooled sample; lower part: reproductive sample. For 40 nt tsRNA: *m/z* = 1313.377, *z* = -10. For 41 nt tsRNA: *m/z* = 1344.080, *z* = -10. Horizontal axis (time) was zoomed in to enhance visualization of these EIC peaks.

| **Table S14. Genomes used for AAA and AAG codon statistics** | |
| --- | --- |
| **species** | **NCBI genome assembly** |
| *Aplysia californica* | AplCal3.0 |
| *Danio rerio* | GRCz12tu |
| *Drosophila melanogaster* | Release 6 plus ISO1 MT |
| *Homo sapiens* | GRCh38.p14 |
| *Mus musculus* | GRCm39 |

| **Table S15. AAA and AAG codon statistics** | | |
| --- | --- | --- |
| **species** | **(AAG)4** | **(AAA)4** |
| *Aplysia californica* | 287 | 7 |
| *Danio rerio* | 469 | 23 |
| *Drosophila melanogaster* | 111 | 7 |
| *Homo sapiens* | 300 | 20 |
| *Mus musculus* | 312 | 16 |
