## Supplementary material for "Sub-stoichiometric modifications of *Aplysia californica* tRNAs and tRNA fragments revealed by integrating intact and bottom-up mass spectrometry": Supplementary_material_2_tRNA_GluCUC.docx

**Supplementary material 2: tRNA^Glu^CUC information**

**Sequence information:**

Organism: *Aplysia californica*

RNA type: tRNA^Glu^CUC

RNAcentral link: <https://rnacentral.org/rna/URS0000928747/6500>

RNA sequence (unmodified, with CCA tail, 5'-3'):

UCCCAAGUGGUCUAGUGGUUAGGAUUUCGCGCUCUCACCGCGACGGCCGGGGUUCGAUUCCCCGCUUGGGAA-CCA

Length: 75

| **List of ribonucleosides detected in *A. californica* tRNA^Glu^CUC digest** | | | | | | |
| --- | --- | --- | --- | --- | --- | --- |
| **RNAMods code** | **short name*** | **full name** | **theoretical [M+H]^+^** | **observed [M+H]^+^** | **mass error (ppm)*** | **product ion(s) detected** |
| P | Y | pseudouridine | 245.0773 | 245.0731 | 17.14 | 209, 179, 155 |
| D | D | dihydrouridine | 247.093 | 247.0886 | 17.81 | 115 |
| C | C | cytidine | 244.0933 | 244.0893 | 16.39 | 112 |
| U | U | uridine | 245.0773 | 245.0730 | 17.55 | 113 |
| " | m^1^A | 1-methyladenosine | 282.1202 | 282.1155 | 16.66 | 150 |
| ? | m^5^C | 5-methylcytidine | 258.1090 | 258.1048 | 16.27 | 126 |
| B | Cm | 2'-O-methylcytidine | 258.1090 | 258.1048 | 16.27 | 112 |
| G | G | guanosine | 284.0995 | 284.0950 | 15.84 | 152 |
| L/K | m^2^G/m^1^G | N2-methylguanosine/1-methylguanosine | 298.1151 | 298.1099 | 17.44 | 166 |
| A | A | adenosine | 268.1046 | 268.0999 | 17.53 | 136 |
| \ | m^5^Um | 5,2'-O-dimethyluridine | 273.1086 | 273.1031 | 20.14 | 127 |
| : | Am | 2'-O-methyladenosine | 282.1202 | 282.1152 | 17.72 | 136 |
| = | m^6^A | N6-methyladenosine | 282.1202 | 282.1150 | 18.43 | 150 |

*mass error was calculated using the observed [M+H]^+^ value at the peak maximum: mass error = (observed [M+H]^+^ - theoretical[M+H]^+^)/ theoretical[M+H]^+^ × 10^6^. The dataset used was: 20250418_Ac#1unk3tRNAGlu (spermatheca).

**Overlaid EICs for ribonucleosides detected in *A. californica* tRNA^Glu^CUC digest**

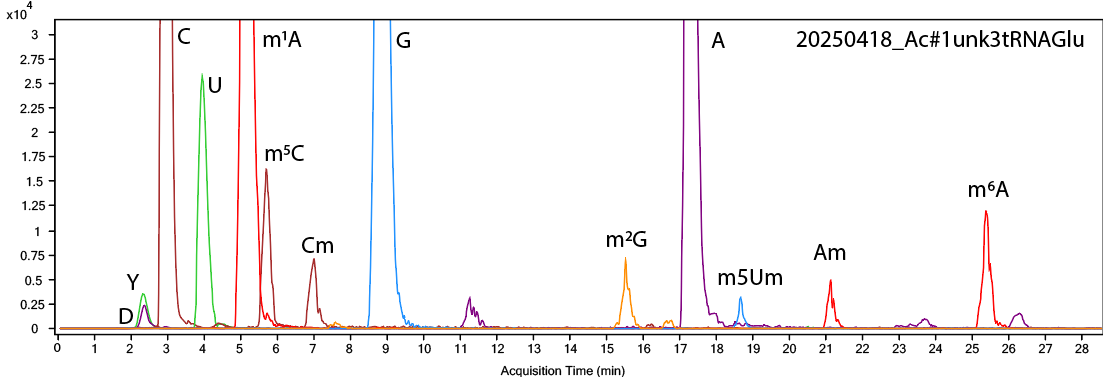

m^2^G/m^1^G

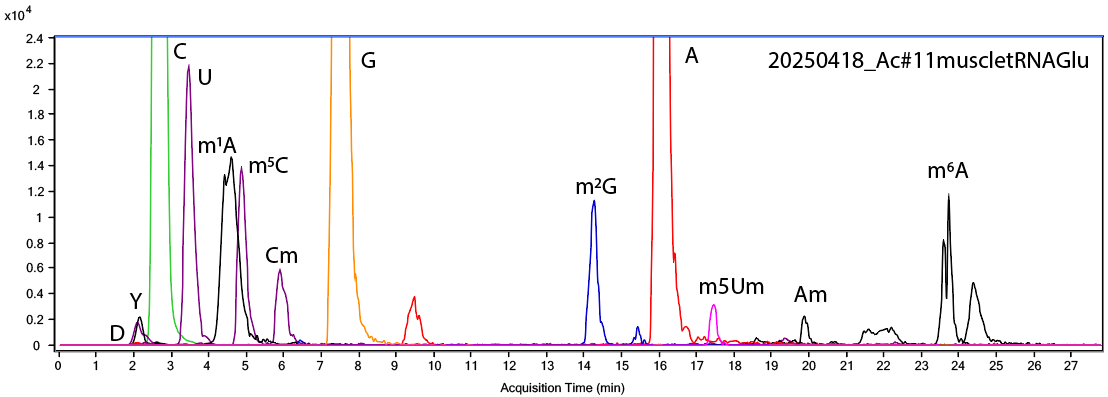

**Part 1. RNase 4 data**

| **list of LC-MS/MS data files used for analysis by RNase 4 digestion** | |
| --- | --- |
| **data file name** | **tissue** |
| 20250310_tRNA-Glu-1+5-R4 | heart (pooled from multiple animals) |
| 20250310_tRNA-Glu-R4-insitu | heart (pooled from multiple animals) |
| 20250317_tRNA-Glu-1+5 | heart (pooled from multiple animals) |
| 20250318_tRNA-Glu-2+3-R4 | heart (pooled from multiple animals) |
| 20250318_tRNA-Glu-2+3-R4-rep2* | heart (pooled from multiple animals) |
| *Pytheas results are based on this data file unless otherwise indicated | |

**Note:**

To simply MS/MS data matching, the position system used throughout this document is absolute position numbering, not that of Sprinzl et al. (1998) based on secondary structure.

| **digested fragment information** | |
| --- | --- |
| **enzyme** | RNase 4 |
| **unmodified sequence** | UCCCAAGU |
| **modified sequence** | N/A |
| **position on *A. c* tRNA^Glu^CUC** | [1-8] |
| **5’ end** | P |
| **3’ end** | OH |
| **charge state detected** | -3 |
| **theoretical m/z** | 848.4380 |
| **ppm used for EIC extraction** | 20 |
| **detection of MS1 signal in data files** | |
| 20250310_tRNA-Glu-1+5-R4 | Y |
| 20250310_tRNA-Glu-R4-insitu | Y |
| 20250317_tRNA-Glu-1+5 | Y |
| 20250318_tRNA-Glu-2+3-R4 | Y |
| 20250318_tRNA-Glu-2+3-R4-rep2 | Y |

*Y = Yes, MS1 signal detected; N = No, MS1 signal not detected

**Corresponding MS1 EIC**

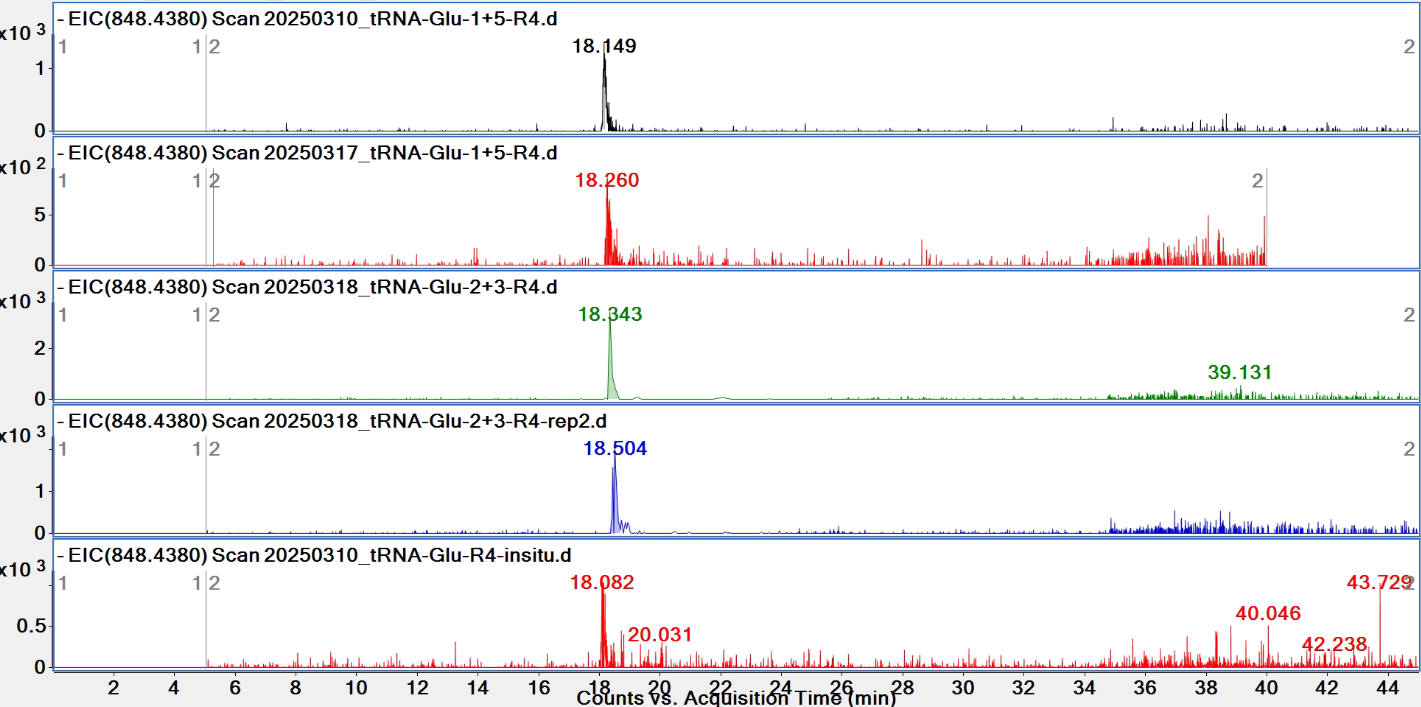

**Pytheas results**

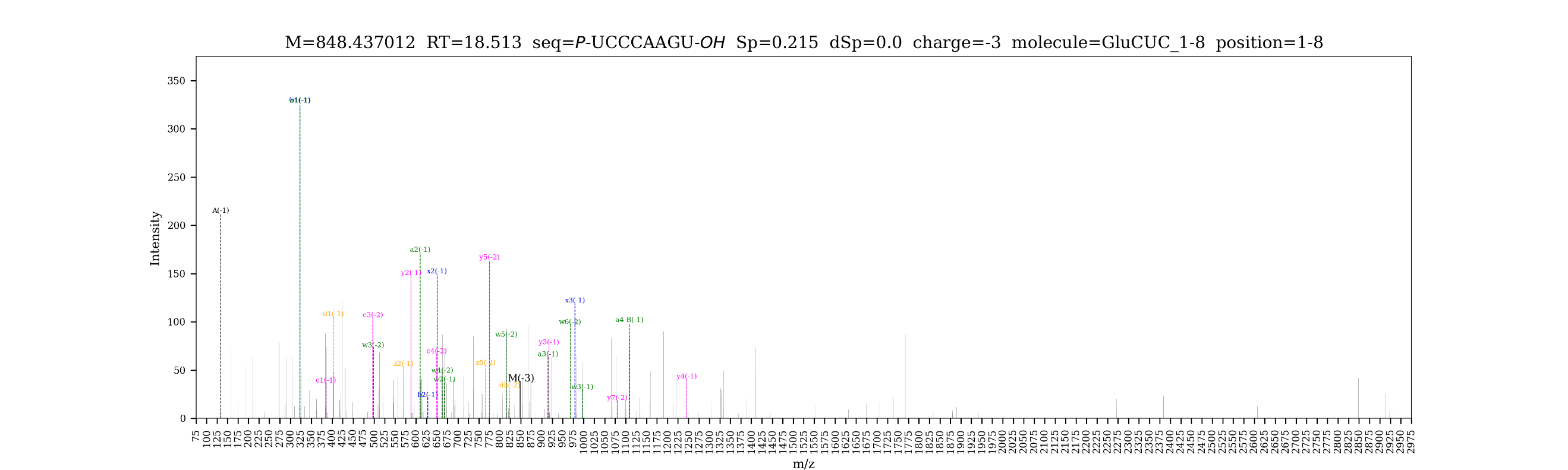

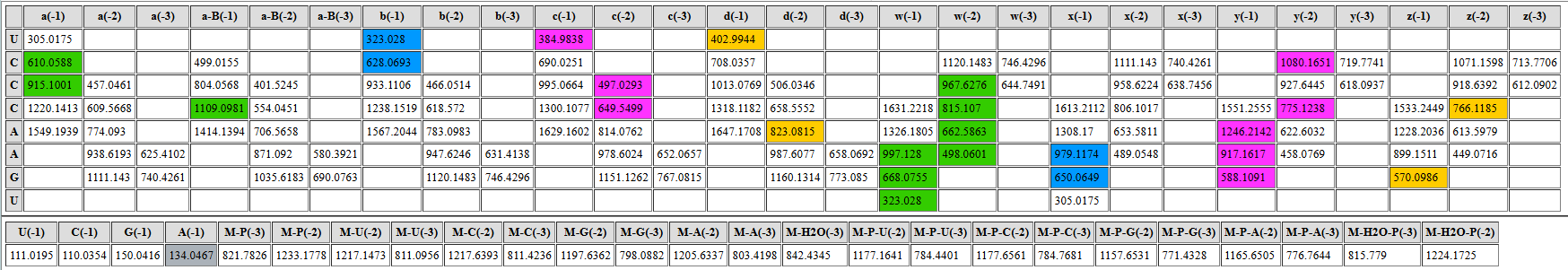

| **digested fragment information** | |
| --- | --- |
| **enzyme** | RNase 4 |
| **unmodified sequence** | AAGU |
| **modified sequence** | N/A |
| **position on *A. c* tRNA^Glu^CUC** | [5-8] |
| **5’ end** | OH |
| **3’ end** | OH |
| **charge state detected** | -1, -2 |
| **theoretical m/z** | 1246.2142, 622.6032 |
| **ppm used for EIC extraction** | 20 |
| **detection of MS1 signal in data files** | |
| 20250310_tRNA-Glu-1+5-R4 | Y, Y |
| 20250310_tRNA-Glu-R4-insitu | N, Y |
| 20250317_tRNA-Glu-1+5 | N, Y |
| 20250318_tRNA-Glu-2+3-R4 | Y, Y |
| 20250318_tRNA-Glu-2+3-R4-rep2 | Y, Y |

**EICs**

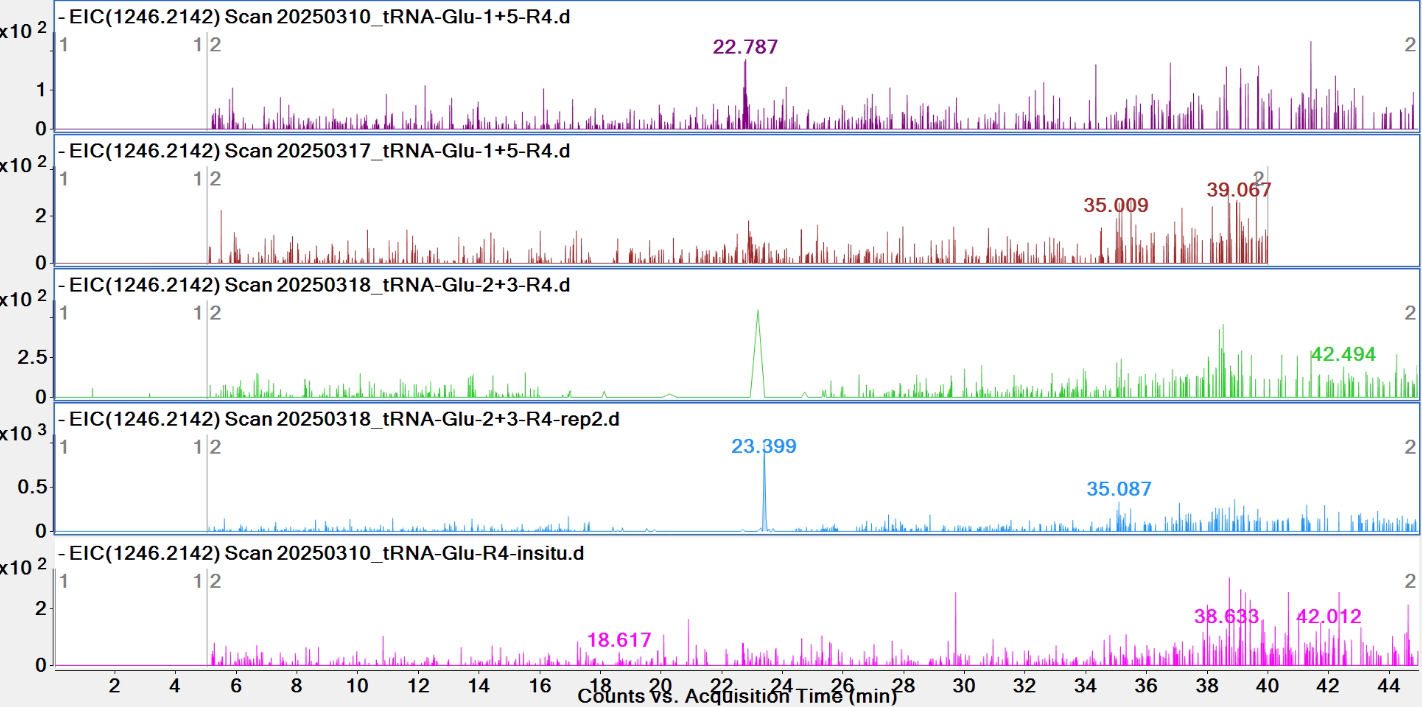

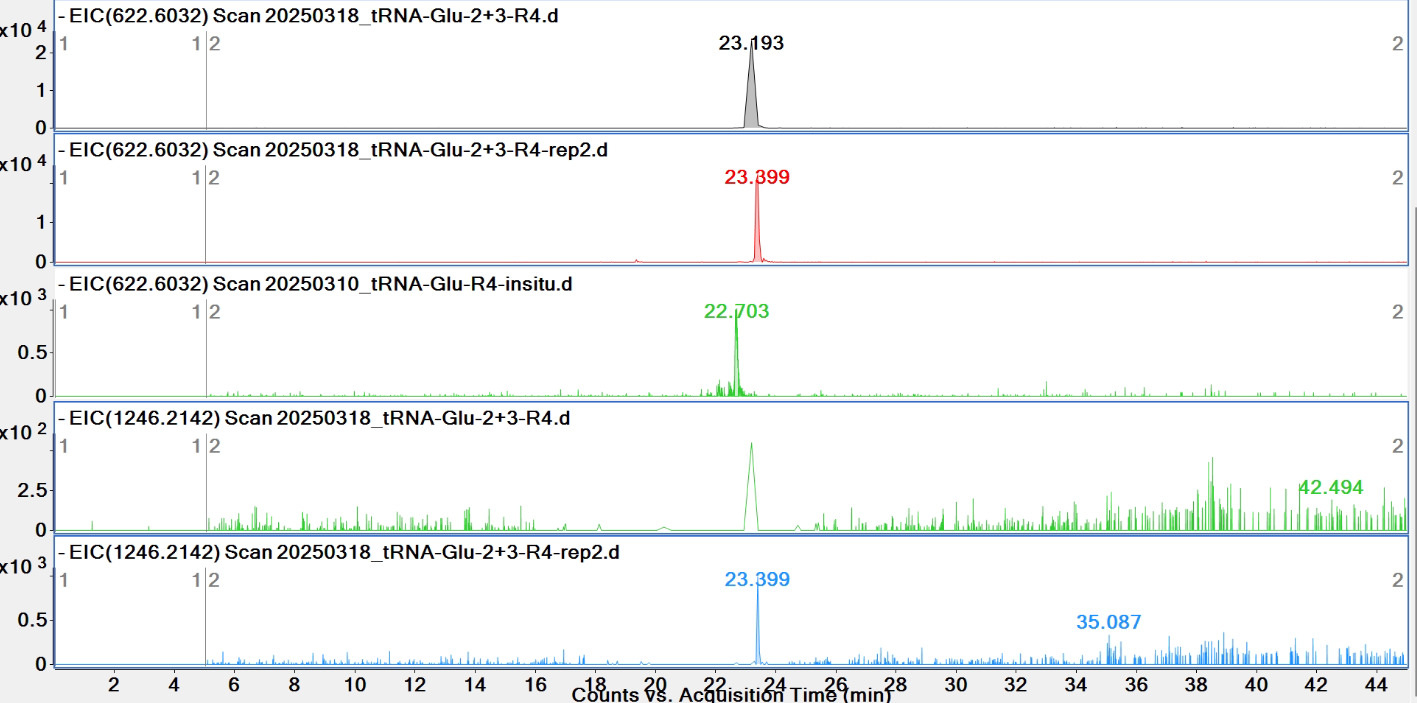

**Pytheas results**

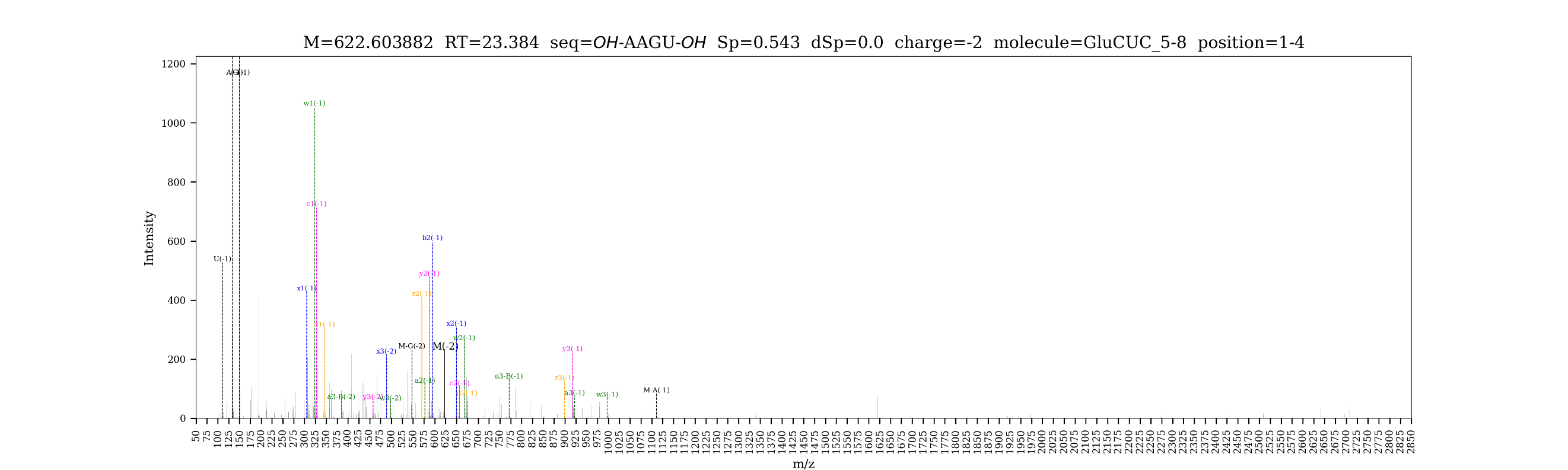

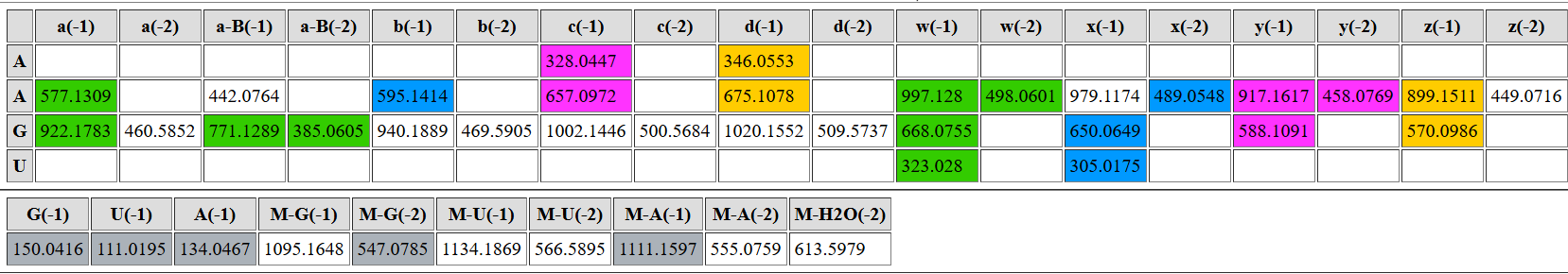

| **digested fragment information** | |
| --- | --- |
| **enzyme** | RNase 4 |
| **unmodified sequence** | GGU |
| **modified sequence** | G[m2G/m1G]U |
| **position on *A. c* tRNA^Glu^CUC** | [9-11] |
| **5’ end** | OH |
| **3’ end** | OH |
| **charge state detected** | -1, -2 |
| **theoretical m/z** | 947.171, 473.081 |
| **ppm used for EIC extraction** | 20 |
| **detection of MS1 signal in data files** | |
| 20250310_tRNA-Glu-1+5-R4 | Y, Y |
| 20250310_tRNA-Glu-R4-insitu | N, Y |
| 20250317_tRNA-Glu-1+5 | Y, Y |
| 20250318_tRNA-Glu-2+3-R4 | Y, Y |
| 20250318_tRNA-Glu-2+3-R4-rep2 | Y, Y |

**EICs**

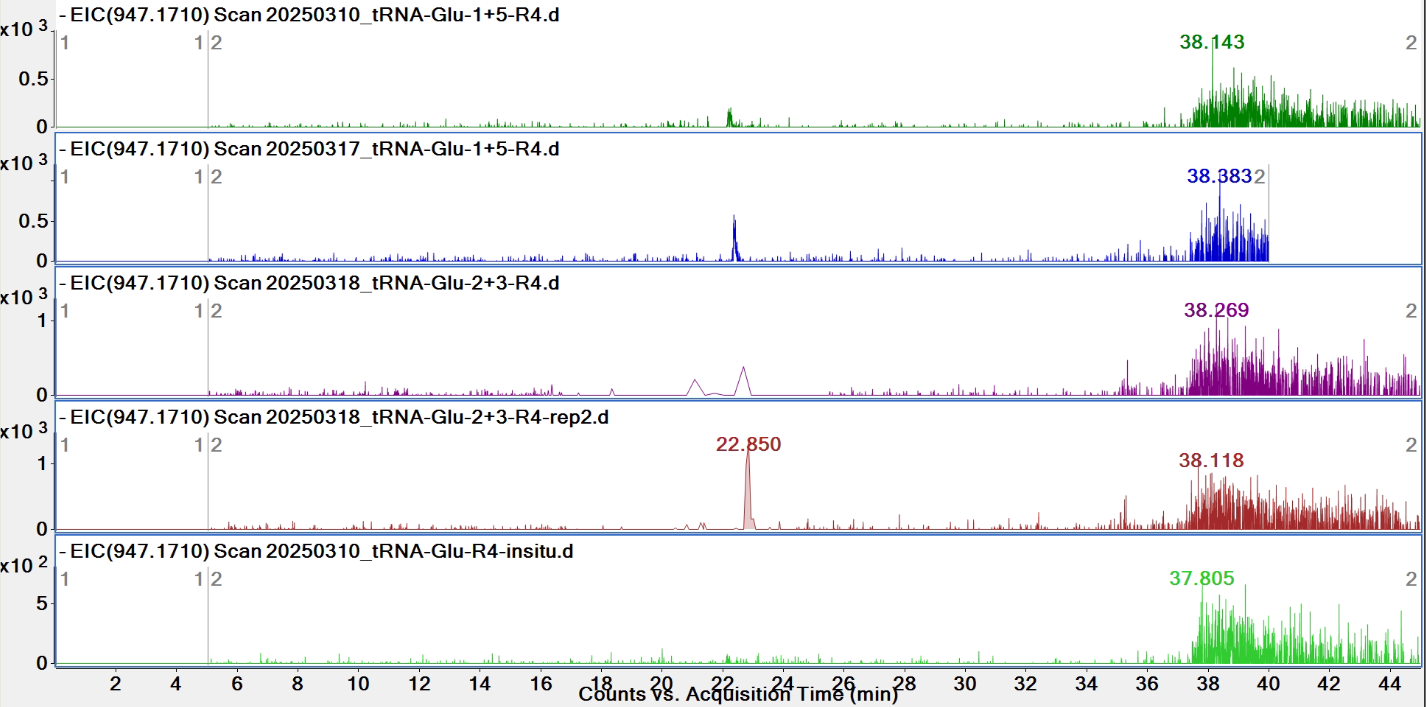

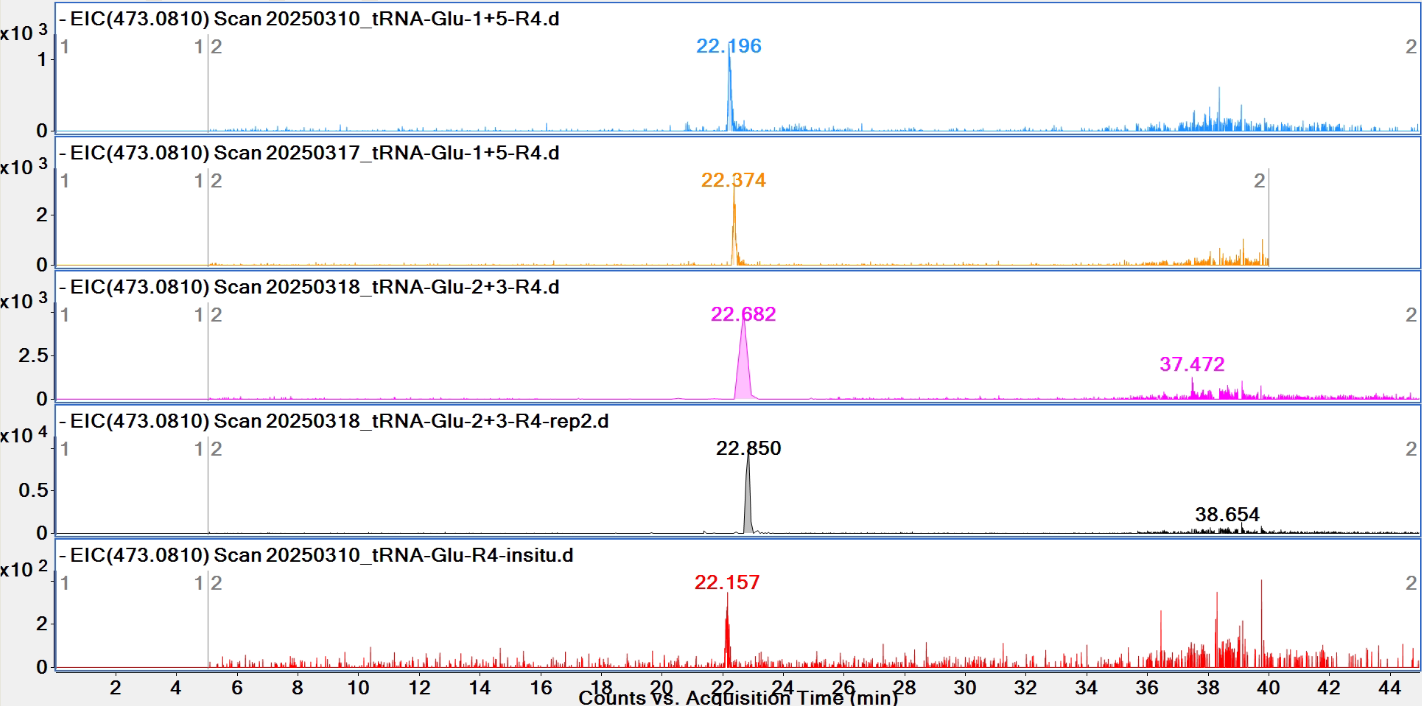

**Pytheas results**

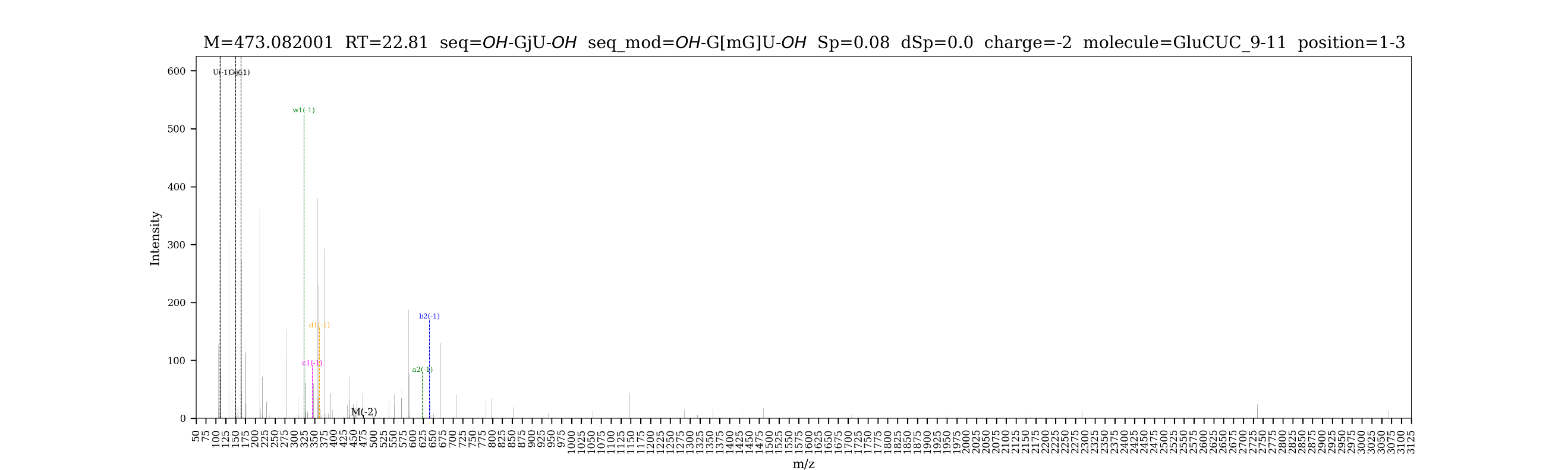

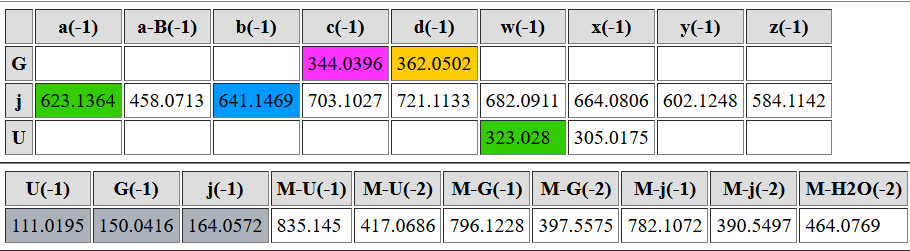

| **digested fragment information** | |
| --- | --- |
| **enzyme** | RNase 4 |
| **unmodified sequence** | CU |
| **modified sequence** | N/A |
| **position on *A. c* tRNA^Glu^CUC** | [12-13] |
| **5’ end** | OH |
| **3’ end** | OH |
| **charge state detected** | -1 |
| **theoretical m/z** | 548.103 |
| **ppm used for EIC extraction** | 20 |
| **detection of MS1 signal in data files** | |
| 20250310_tRNA-Glu-1+5-R4 | N |
| 20250310_tRNA-Glu-R4-insitu | N |
| 20250317_tRNA-Glu-1+5 | N |
| 20250318_tRNA-Glu-2+3-R4 | Y |
| 20250318_tRNA-Glu-2+3-R4-rep2 | Y |

**EICs**

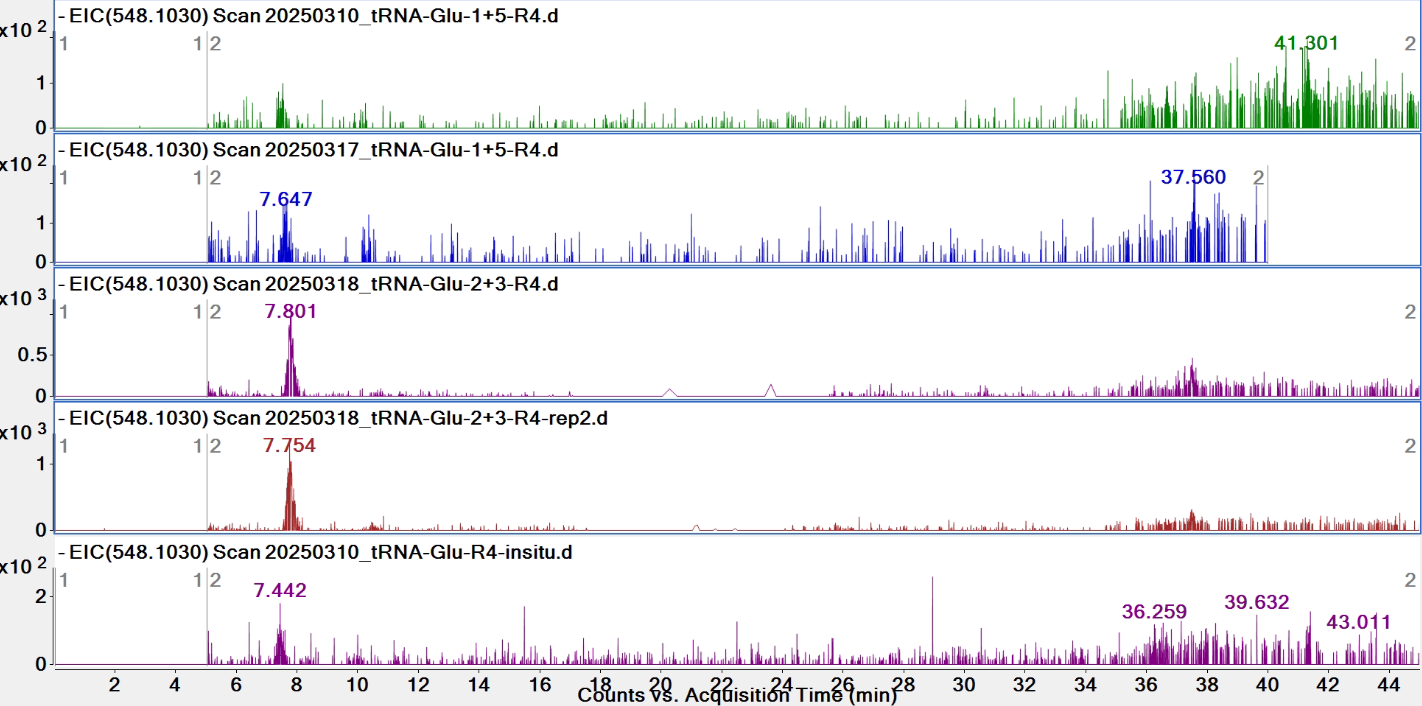

**Pytheas results:** Pytheas does not match 2-nt RNA

| **digested fragment information** | |
| --- | --- |
| **enzyme** | RNase 4 |
| **unmodified sequence** | AGU |
| **modified sequence** | AGD |
| **position on *A. c* tRNA^Glu^CUC** | [14-16] |
| **5’ end** | OH |
| **3’ end** | OH |
| **charge state detected** | -1, -2 |
| **theoretical m/z** | unmodified: 917.1617, 458.0769  modified: 919.1773, 459.0847 |
| **ppm used for EIC extraction** | 20 |
| **detection of MS1 signal in data files** | |
| 20250310_tRNA-Glu-1+5-R4 | unmodified: Y, Y; modified: Y, Y |
| 20250310_tRNA-Glu-R4-insitu | unmodified: Y, Y; modified: Y, Y |
| 20250317_tRNA-Glu-1+5 | unmodified: Y, Y; modified: Y, Y |
| 20250318_tRNA-Glu-2+3-R4 | unmodified: Y, Y; modified: Y, Y |
| 20250318_tRNA-Glu-2+3-R4-rep2 | unmodified: Y, Y; modified: Y, Y |

**EICs**

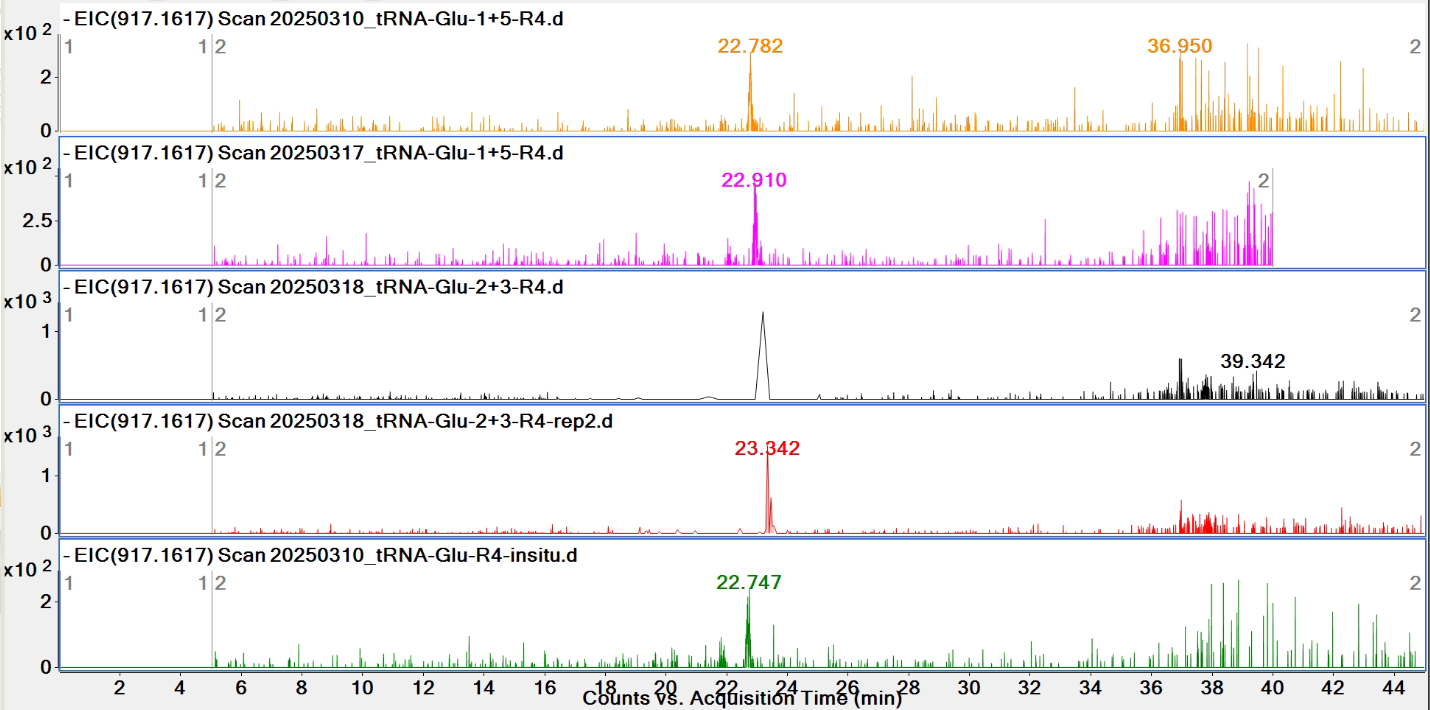

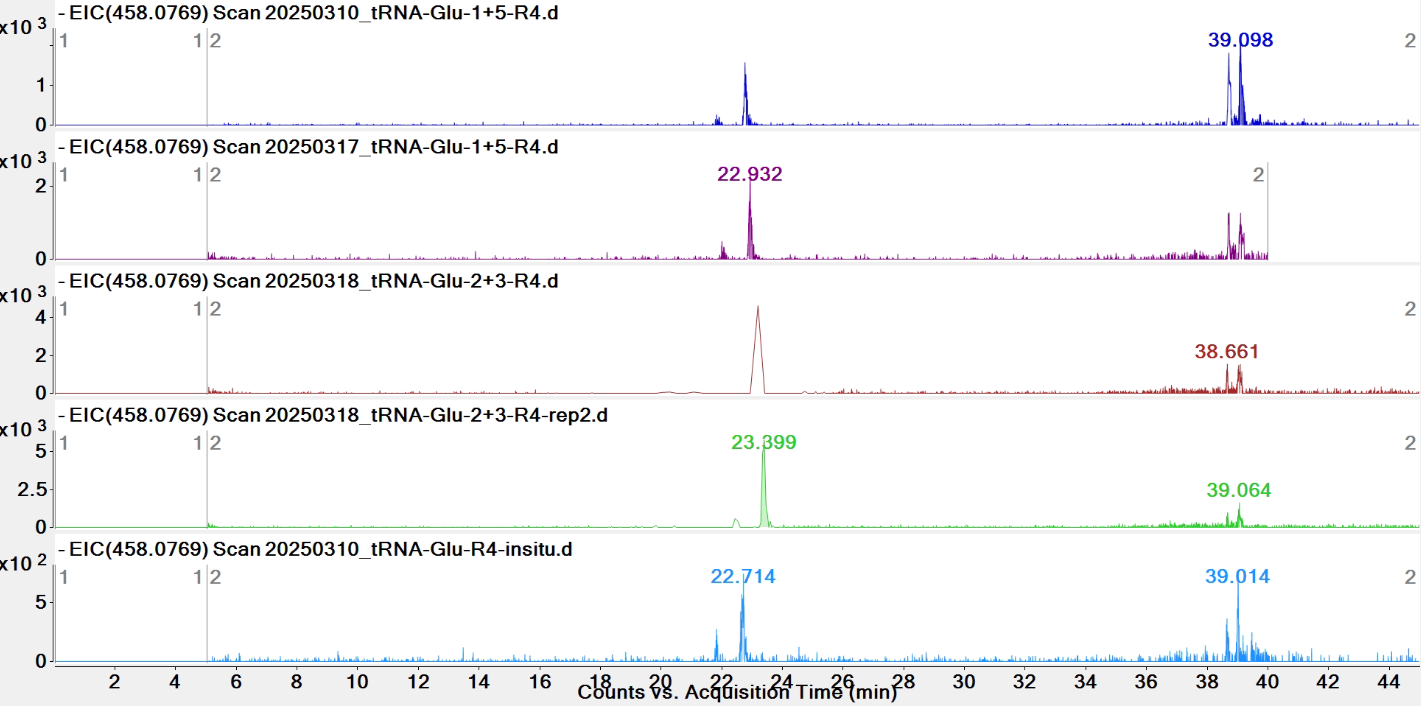

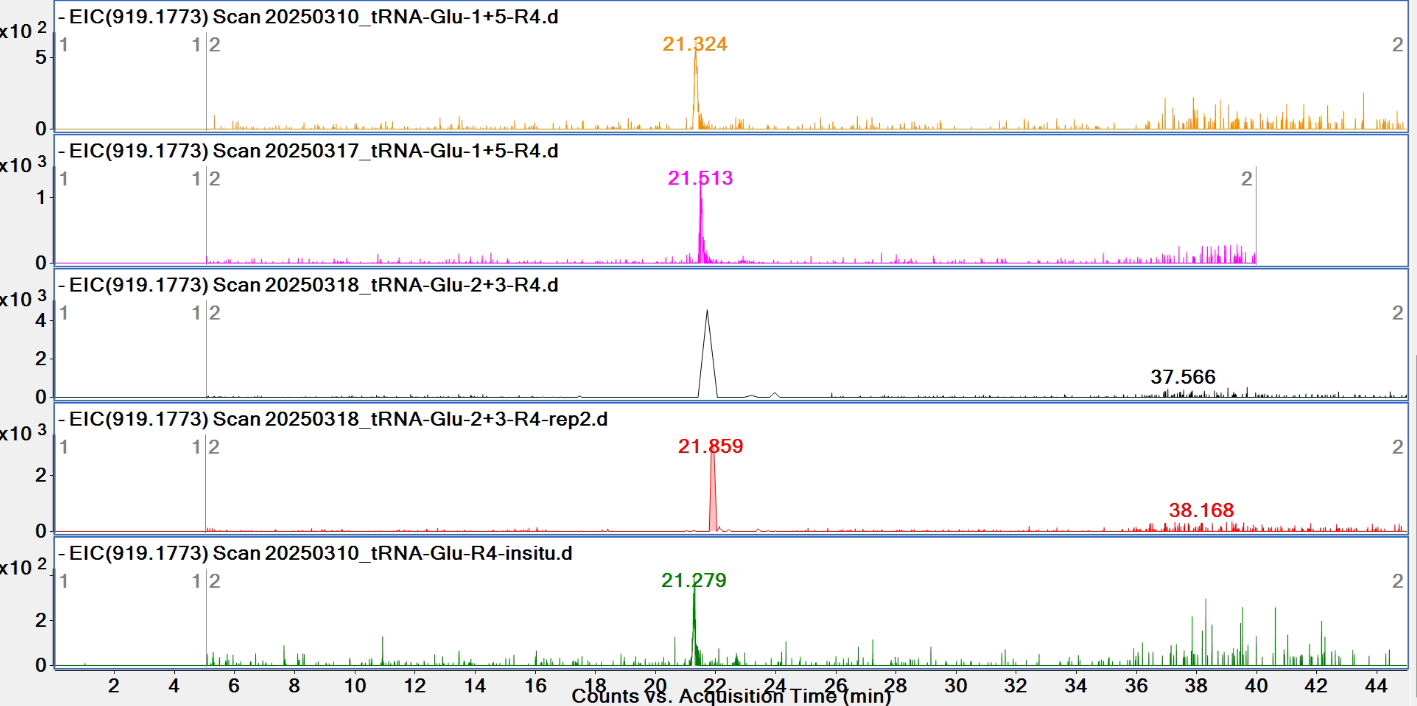

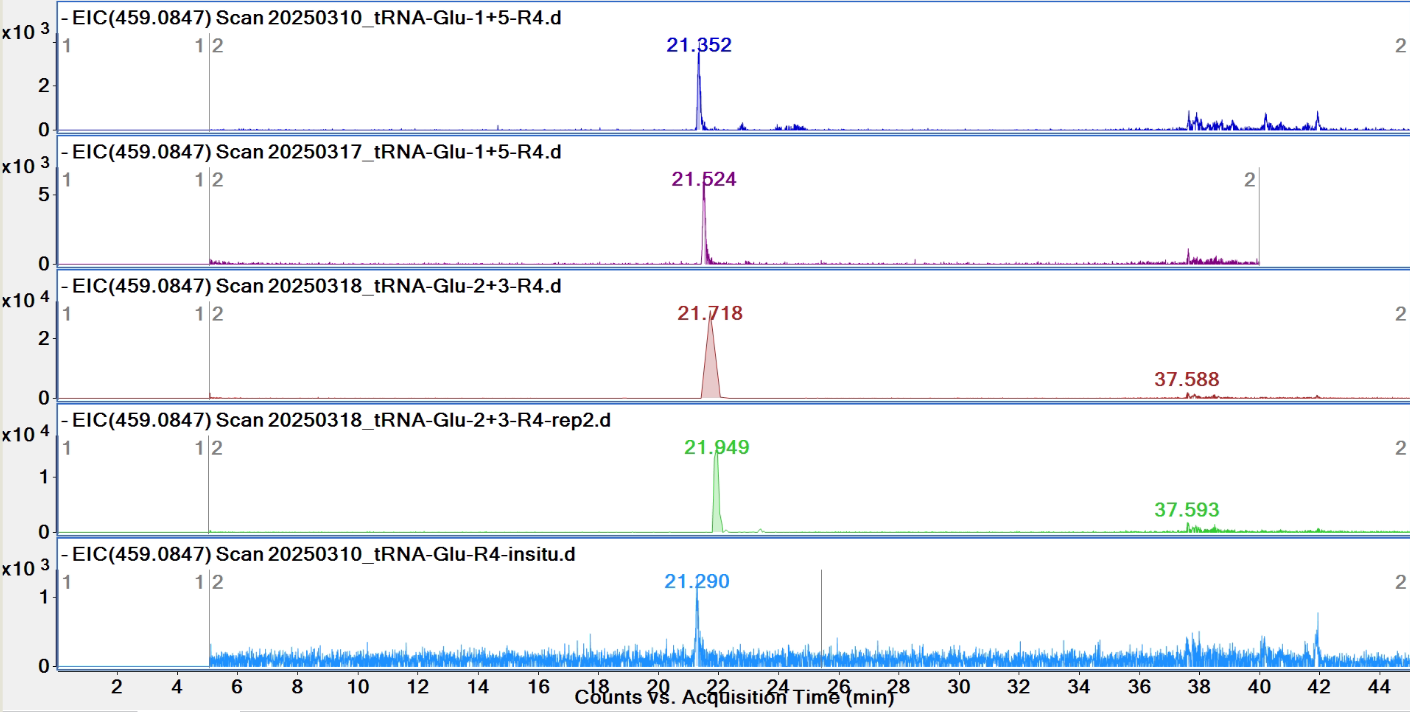

**Pytheas results**

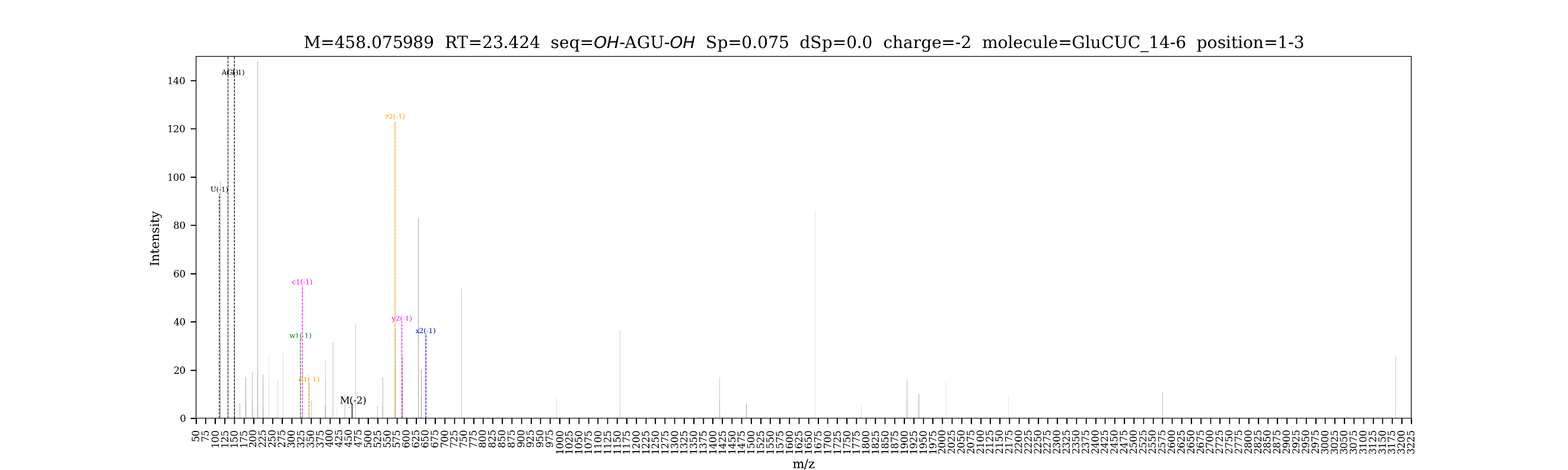

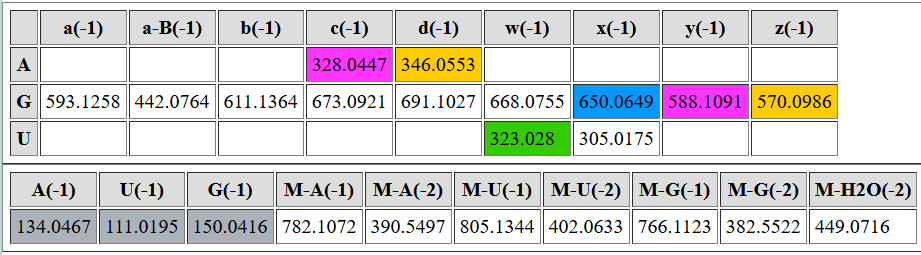

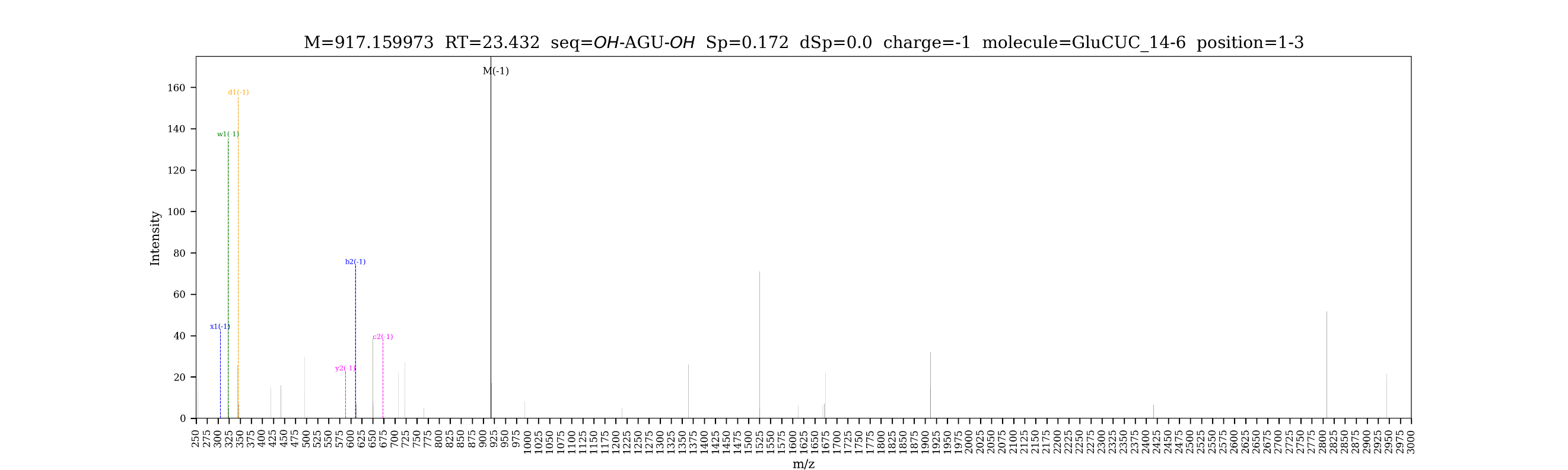

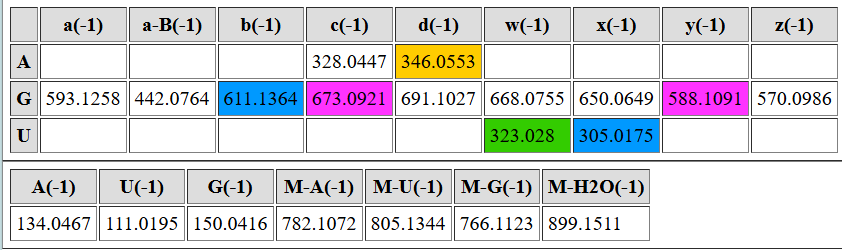

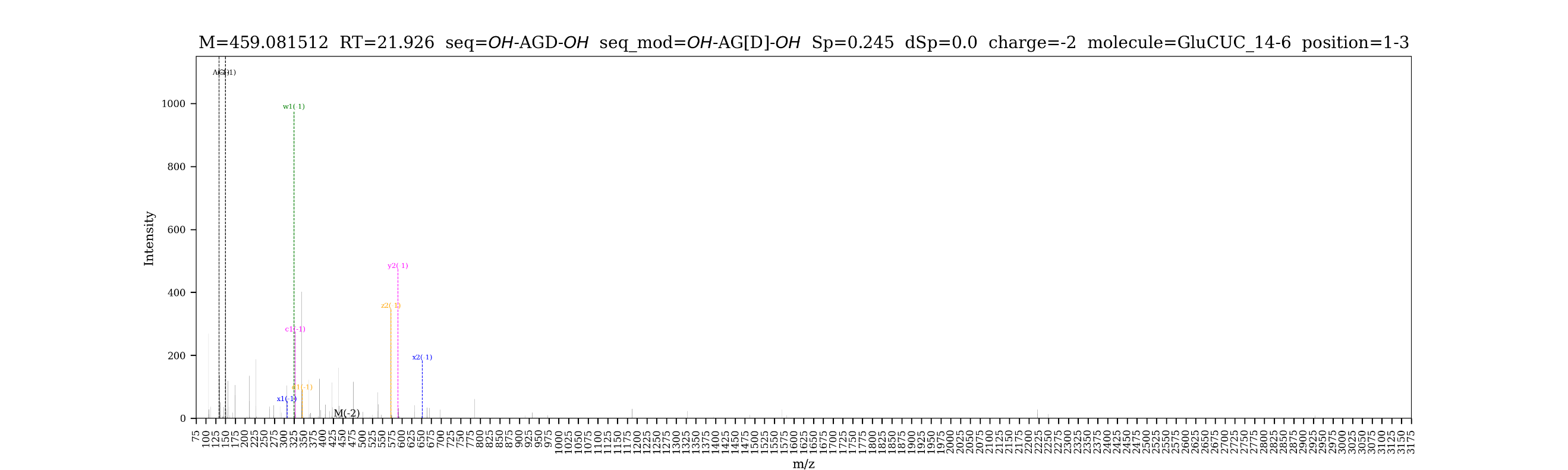

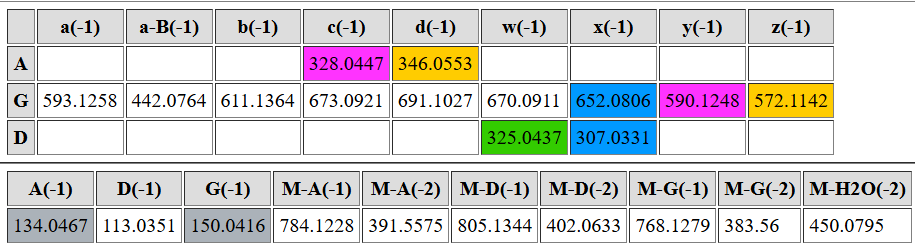

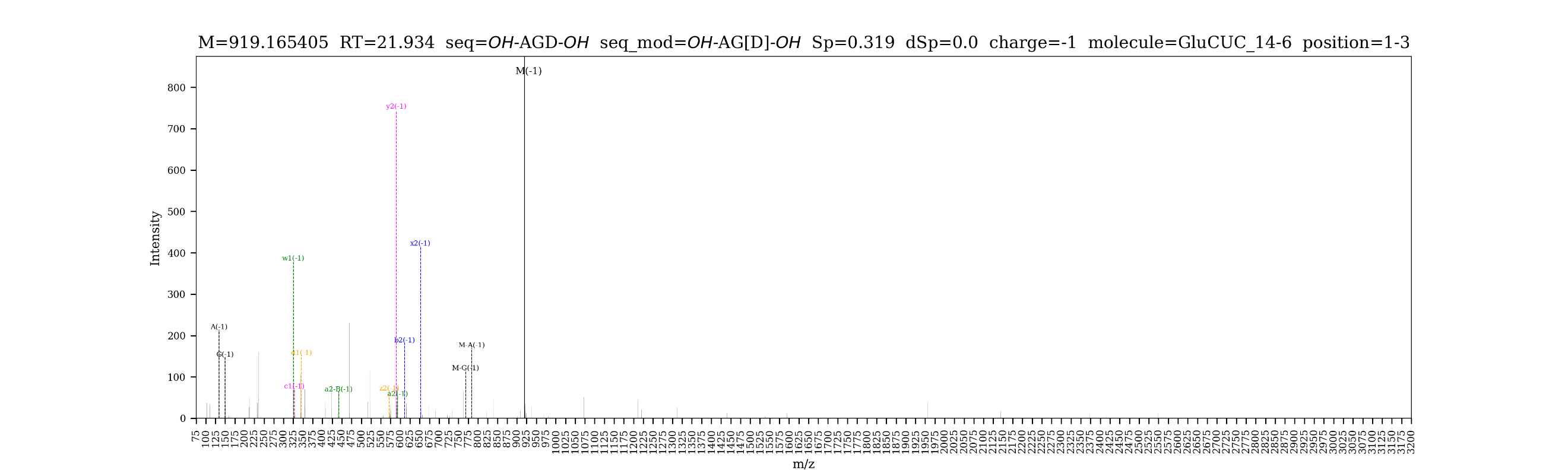

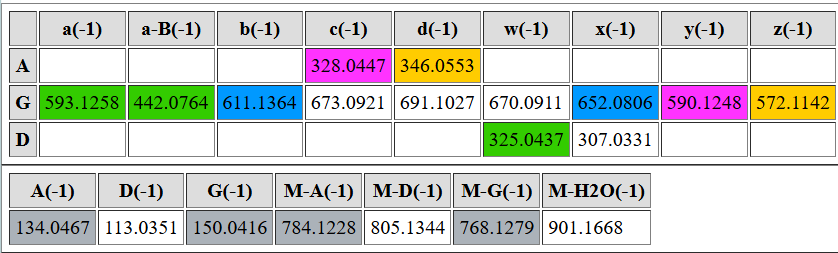

| **digested fragment information** | |
| --- | --- |
| **enzyme** | RNase 4 |
| **unmodified sequence** | GGU |
| **modified sequence** | GGD |
| **position on *A. c* tRNA^Glu^CUC** | [17-19] |
| **5’ end** | OH |
| **3’ end** | OH |
| **charge state detected** | -1, -2 |
| **theoretical m/z** | unmodified: 933.1566, 466.0744  modified: 935.1722, 467.0822 |
| **ppm used for EIC extraction** | 20 |
| **detection of MS1 signal in data files** | |
| 20250310_tRNA-Glu-1+5-R4 | unmodified: N, Y; modified: N, Y |
| 20250310_tRNA-Glu-R4-insitu | unmodified: N, Y; modified: N, Y |
| 20250317_tRNA-Glu-1+5 | unmodified: N, Y; modified: Y, Y |
| 20250318_tRNA-Glu-2+3-R4 | unmodified: N, N; modified: Y, Y |
| 20250318_tRNA-Glu-2+3-R4-rep2 | unmodified: N, Y; modified: Y, Y |

**EICs**

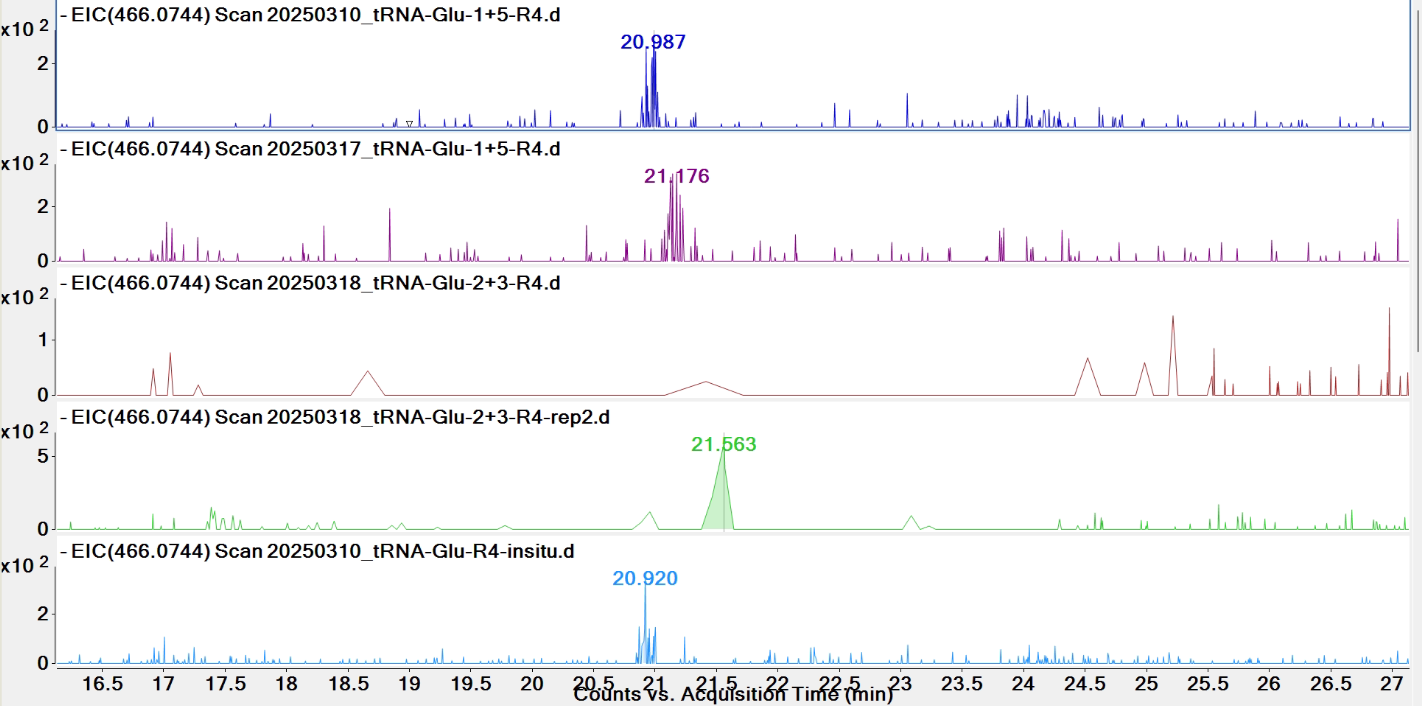

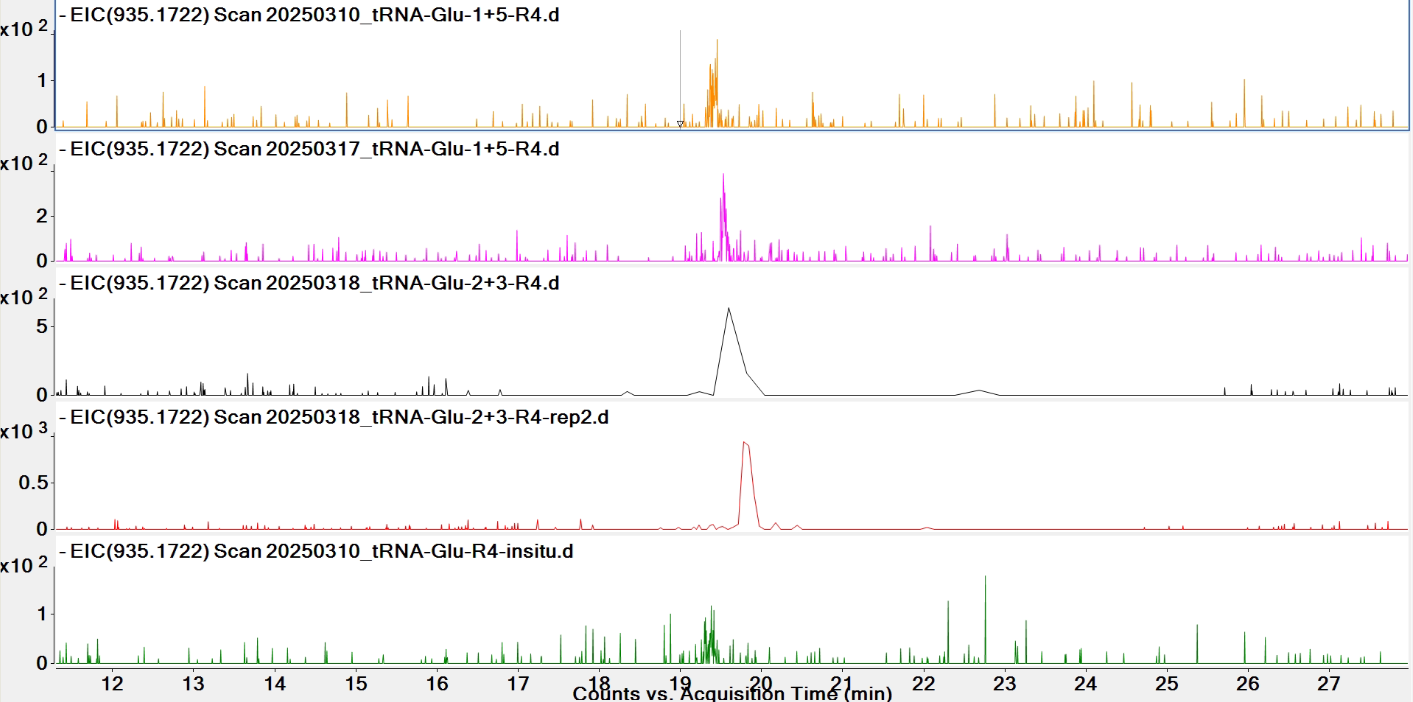

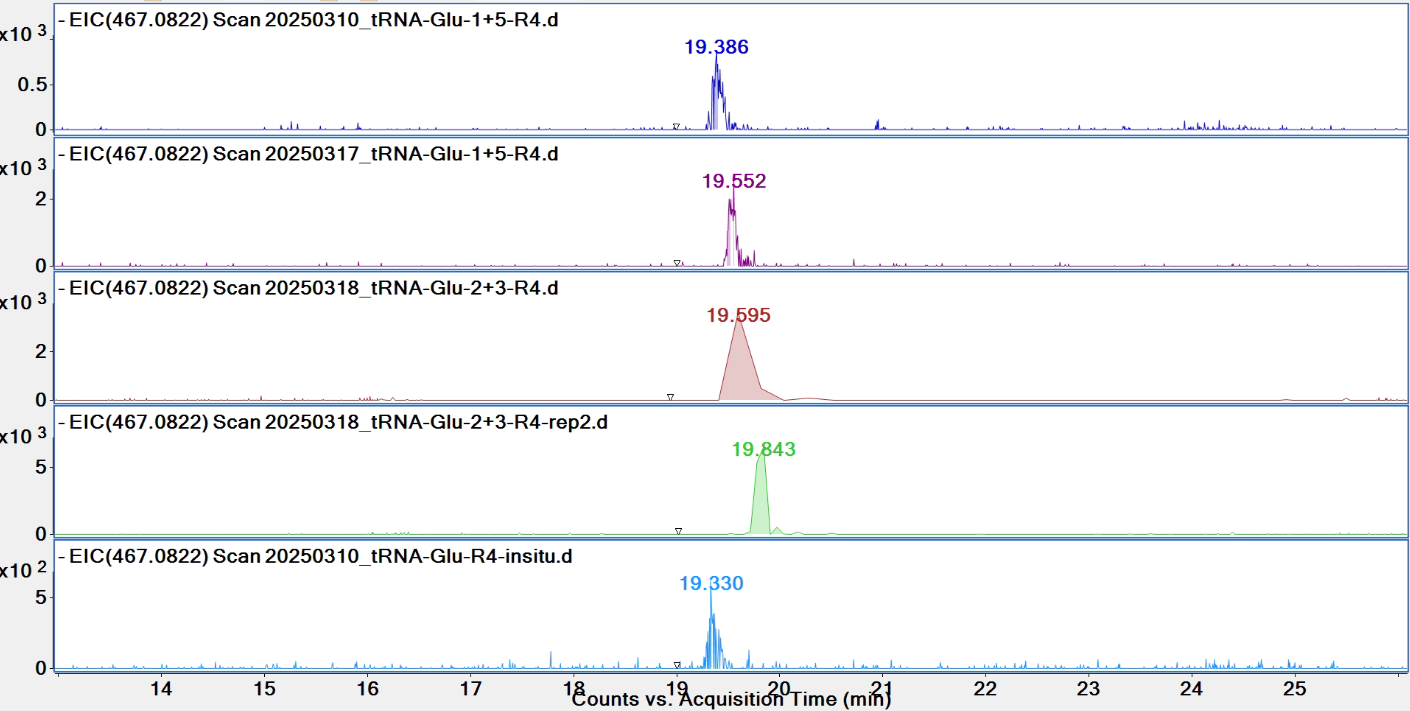

**Pytheas results**

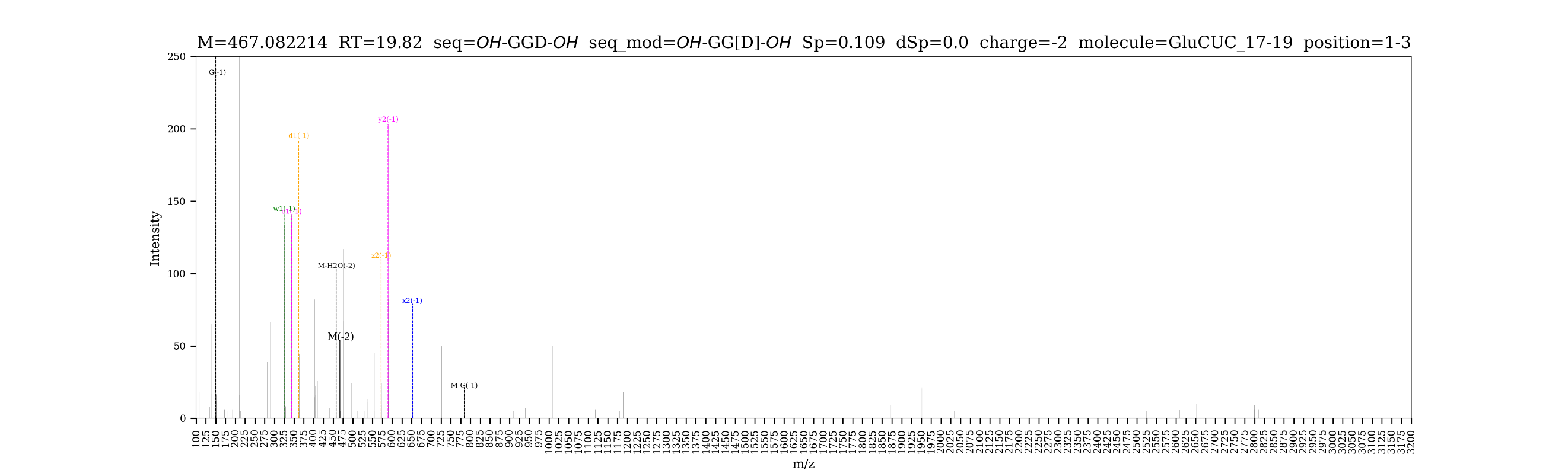

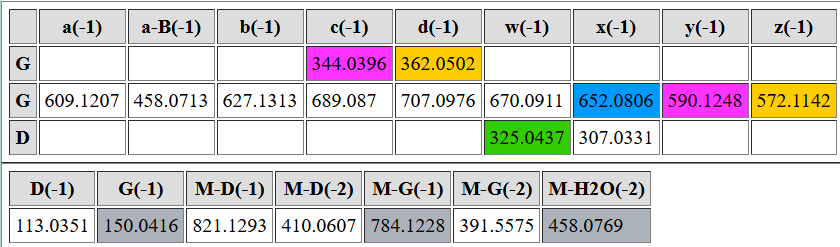

| **digested fragment information** | |
| --- | --- |
| **enzyme** | RNase 4 |
| **unmodified sequence** | AGGAUUU |
| **modified sequence** | N/A |
| **position on *A. c* tRNA^Glu^CUC** | [21-27] |
| **5’ end** | OH |
| **3’ end** | OH |
| **charge state detected** | -2, -3 |
| **theoretical m/z** | 1101.1522, 733.7655 |
| **ppm used for EIC extraction** | 20 |
| **detection of MS1 signal in data files** | |
| 20250310_tRNA-Glu-1+5-R4 | N, N |
| 20250310_tRNA-Glu-R4-insitu | N, N |
| 20250317_tRNA-Glu-1+5 | N, N |
| 20250318_tRNA-Glu-2+3-R4 | N, N |
| 20250318_tRNA-Glu-2+3-R4-rep2 | N, Y |

**EICs**

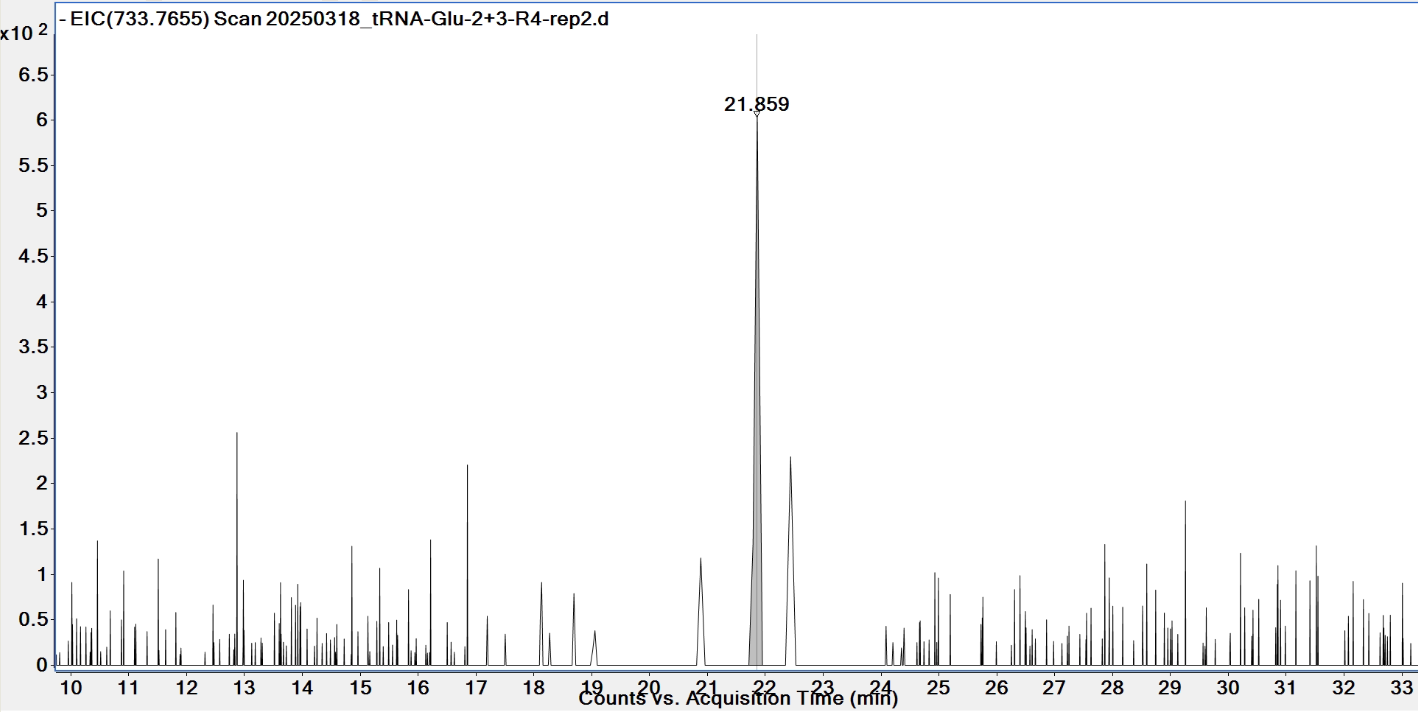

**Pytheas results:** no match

**See the next page for shorter fragments.**

| **digested fragment information** | |
| --- | --- |
| **enzyme** | RNase 4 |
| **unmodified sequence** | AGGAUU |
| **modified sequence** | N/A |
| **position on *A. c* tRNA^Glu^CUC** | [21-26] |
| **5’ end** | OH |
| **3’ end** | OH |
| **charge state detected** | -2, -3 |
| **theoretical m/z** | 948.1396, 631.7571 |
| **ppm used for EIC extraction** | 20 |
| **detection of MS1 signal in data files** | |
| 20250310_tRNA-Glu-1+5-R4 | Y, N |
| 20250310_tRNA-Glu-R4-insitu | N, N |
| 20250317_tRNA-Glu-1+5 | Y, N |
| 20250318_tRNA-Glu-2+3-R4 | Y, N |
| 20250318_tRNA-Glu-2+3-R4-rep2 | Y, Y |

**EICs**

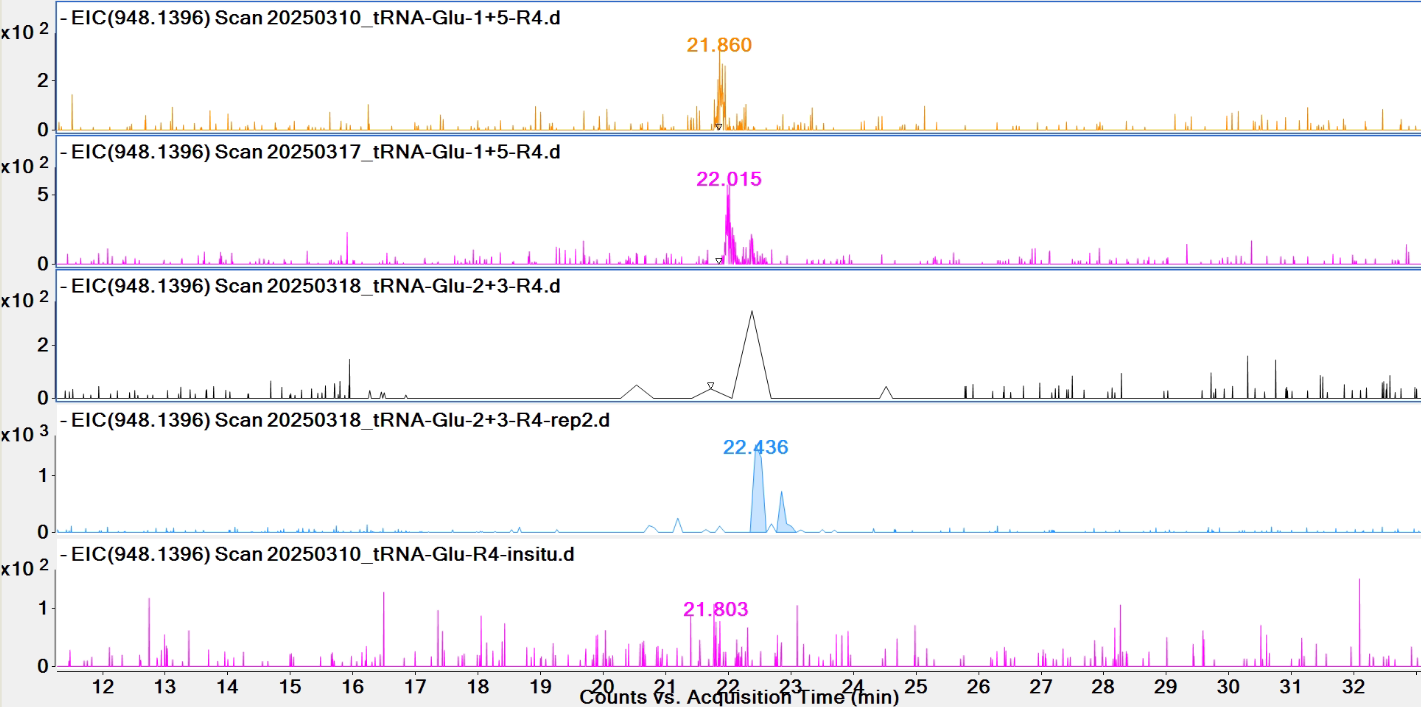

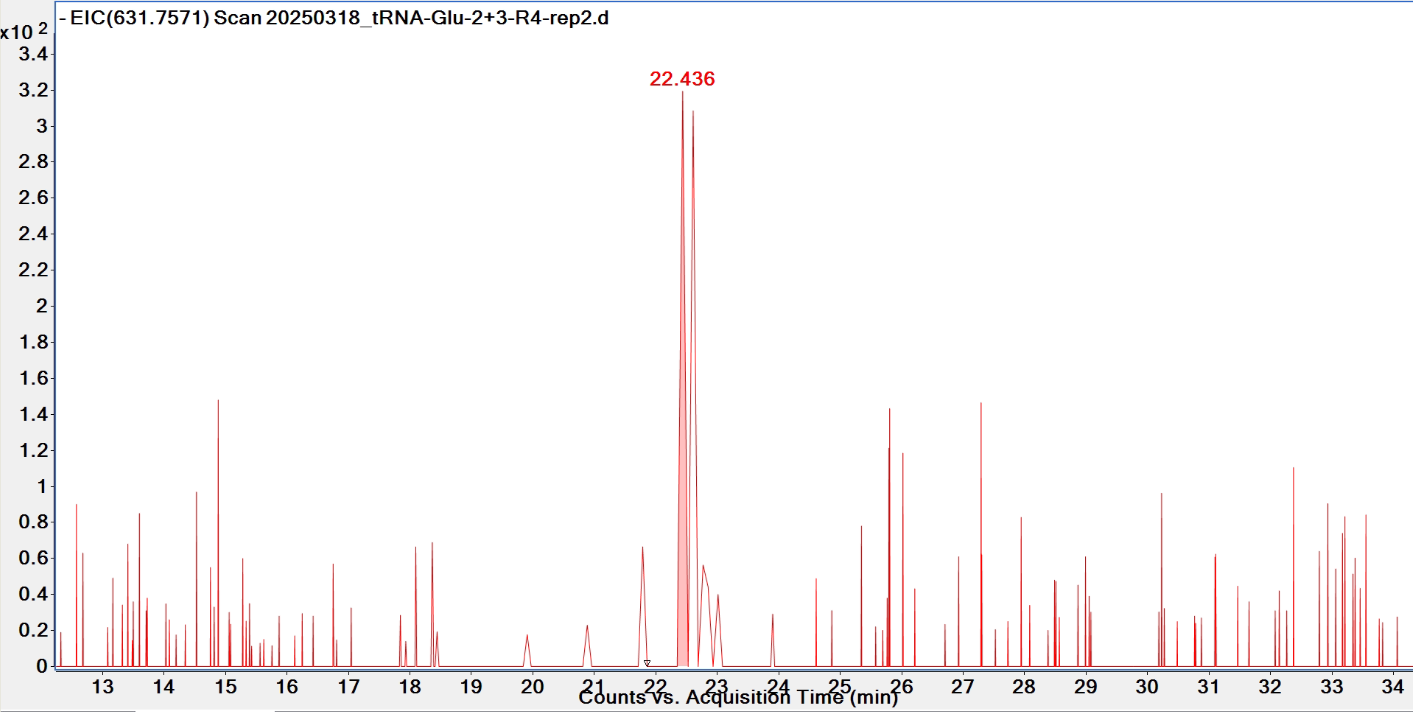

**Pytheas results**

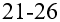

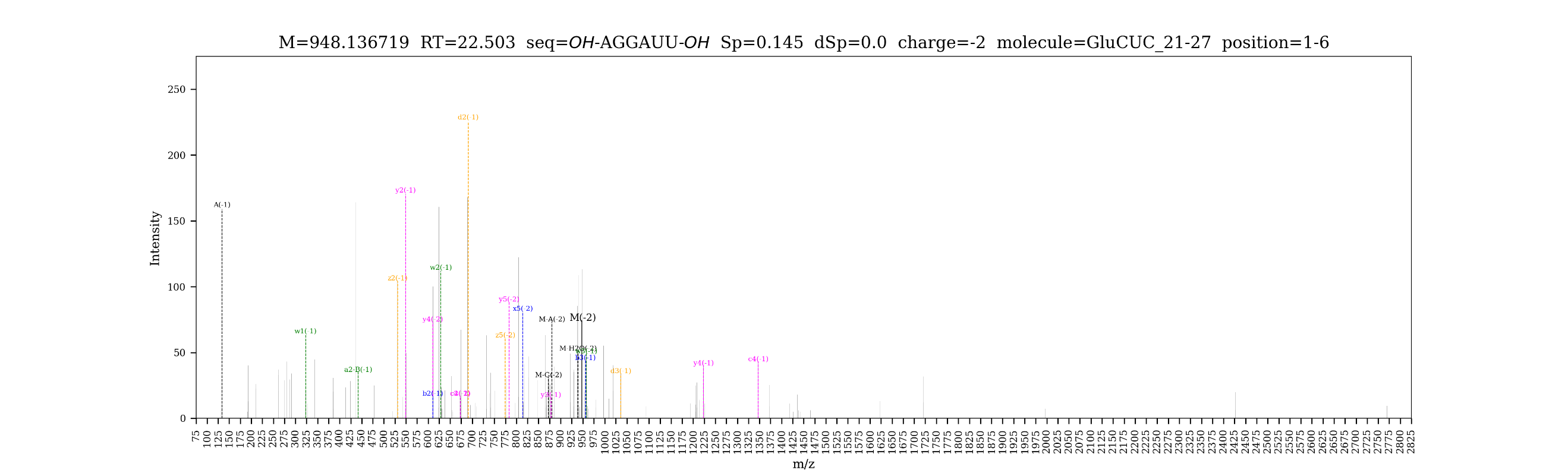

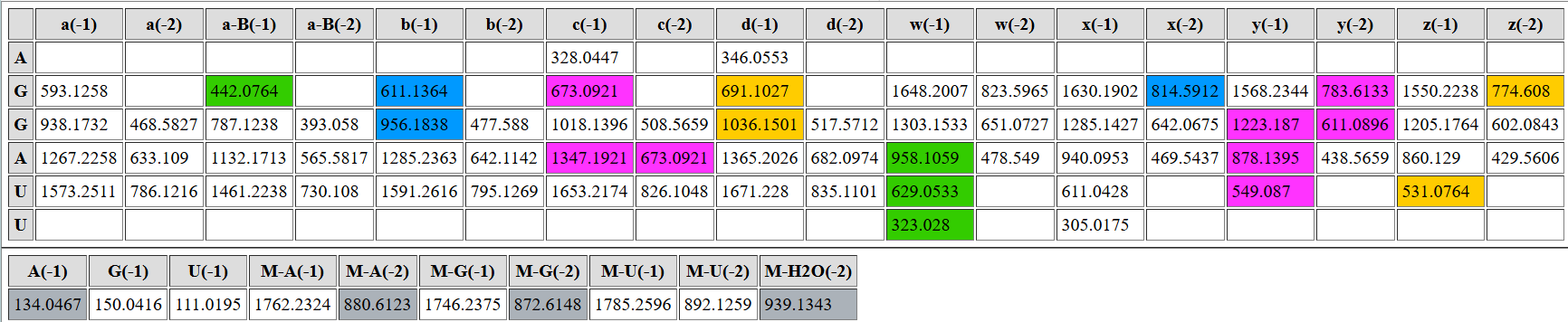

| **digested fragment information** | |
| --- | --- |
| **enzyme** | RNase 4 |
| **unmodified sequence** | AGGAU |
| **modified sequence** | N/A |
| **position on *A. c* tRNA^Glu^CUC** | [21-25] |
| **5’ end** | OH |
| **3’ end** | OH |
| **charge state detected** | -2 |
| **theoretical m/z** | 795.1269 |
| **ppm used for EIC extraction** | 20 |
| **detection of MS1 signal in data files** | |
| 20250310_tRNA-Glu-1+5-R4 | Y |
| 20250310_tRNA-Glu-R4-insitu | N |
| 20250317_tRNA-Glu-1+5 | Y |
| 20250318_tRNA-Glu-2+3-R4 | Y |
| 20250318_tRNA-Glu-2+3-R4-rep2 | Y |

**EICs**

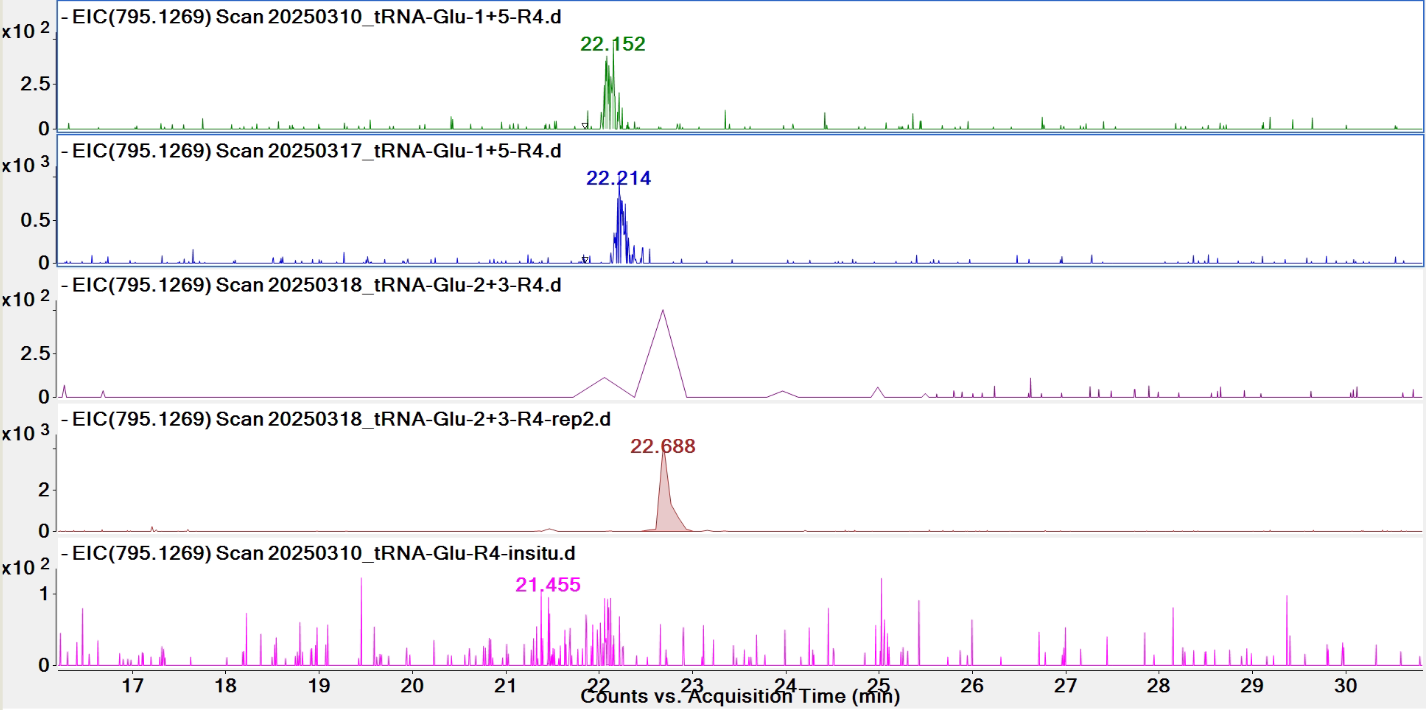

**Pytheas results**

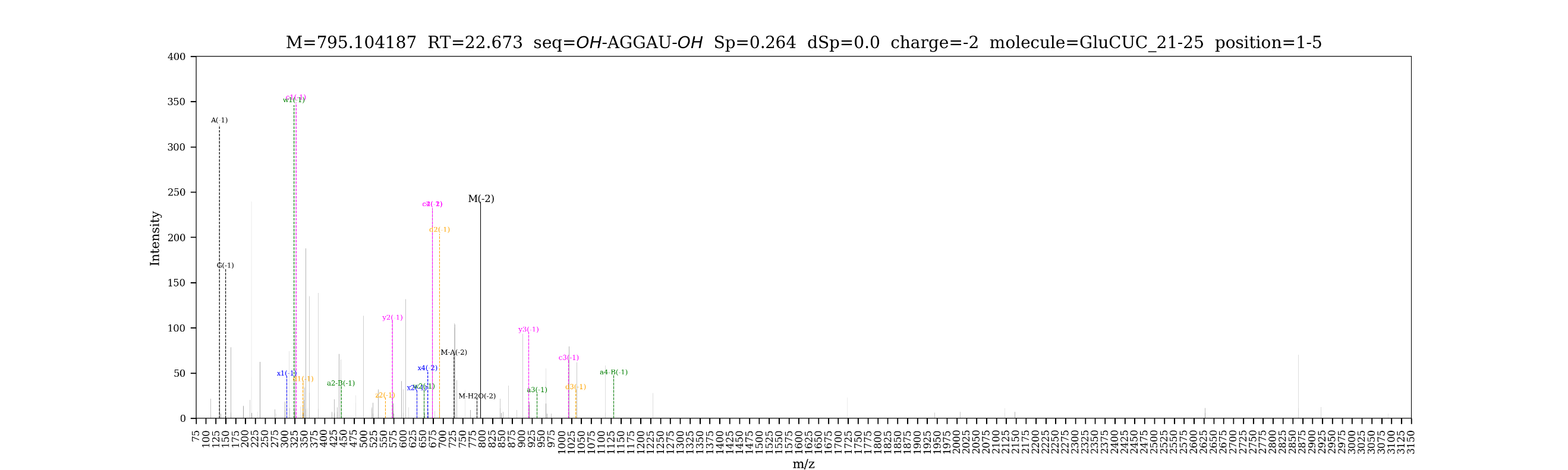

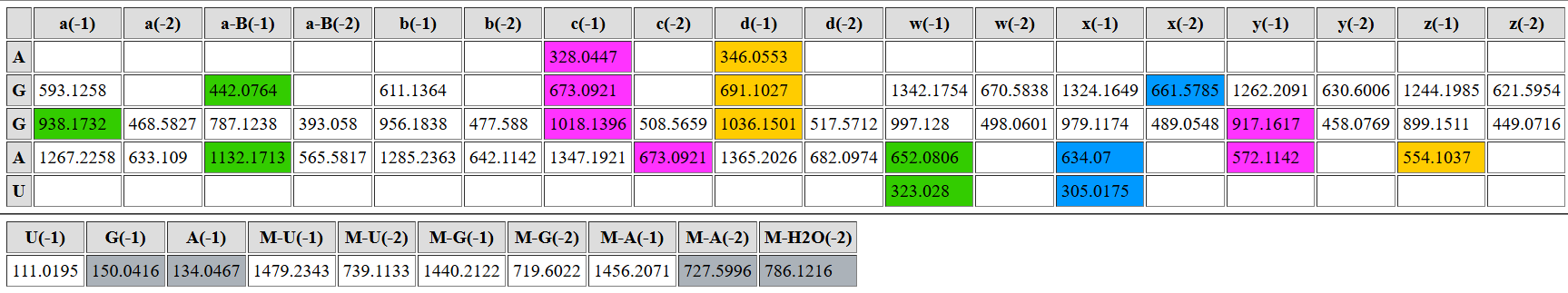

| **digested fragment information** | |
| --- | --- |
| **enzyme** | RNase 4 |
| **unmodified sequence** | CGCGCU |
| **modified sequence** | CGCG[mC]U |
| **position on *A. c* tRNA^Glu^CUC** | [28-33] |
| **5’ end** | OH |
| **3’ end** | OH |
| **charge state detected** | -2, -3 |
| **theoretical m/z** | modified: 930.6441, 620.0935 |
| **ppm used for EIC extraction** | 20 |
| **detection of MS1 signal in data files** | |
| 20250310_tRNA-Glu-1+5-R4 | N, Y |
| 20250310_tRNA-Glu-R4-insitu | N, N |
| 20250317_tRNA-Glu-1+5 | Y, Y |
| 20250318_tRNA-Glu-2+3-R4 | Y, Y |
| 20250318_tRNA-Glu-2+3-R4-rep2 | Y, Y |

**EICs**

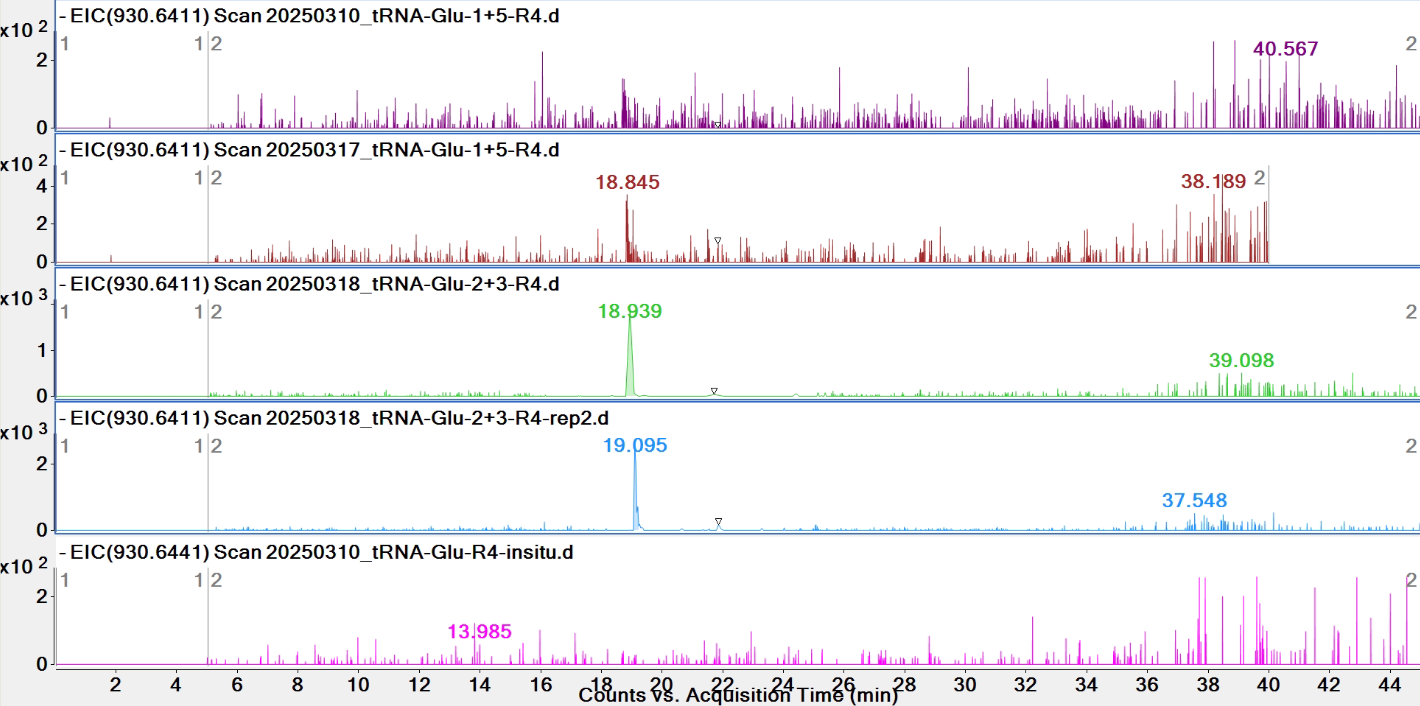

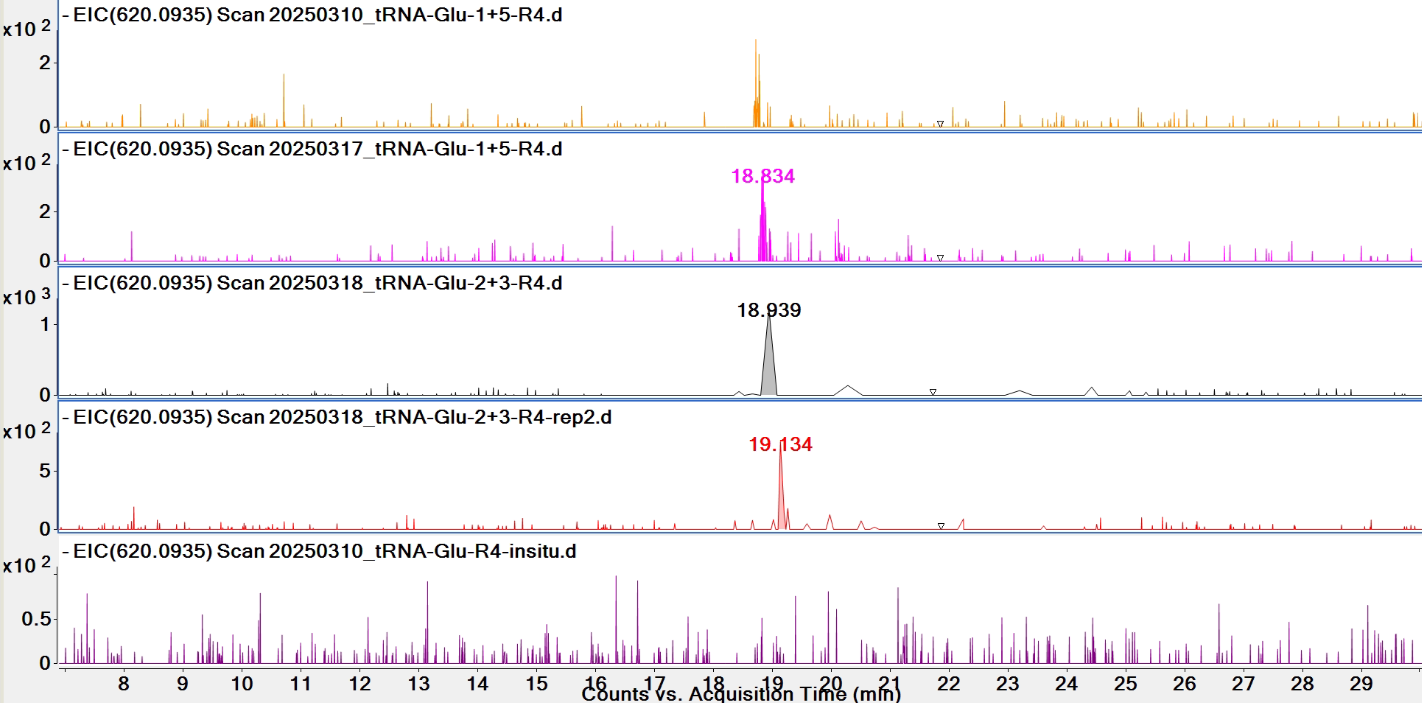

**Pytheas results: no MS2 match**

| **digested fragment information** | |
| --- | --- |
| **enzyme** | RNase 4 |
| **unmodified sequence** | CU |
| **modified sequence** | N/A |
| **position on *A. c* tRNA^Glu^CUC** | [34-35] |
| **5’ end** | OH |
| **3’ end** | OH |
| **charge state detected** | -1 |
| **theoretical m/z** | 548.1030 |
| **ppm used for EIC extraction** | 20 |
| **detection of MS1 signal in data files** | |
| 20250310_tRNA-Glu-1+5-R4 | N |
| 20250310_tRNA-Glu-R4-insitu | N |
| 20250317_tRNA-Glu-1+5 | N |
| 20250318_tRNA-Glu-2+3-R4 | Y |
| 20250318_tRNA-Glu-2+3-R4-rep2 | Y |

**EICs**

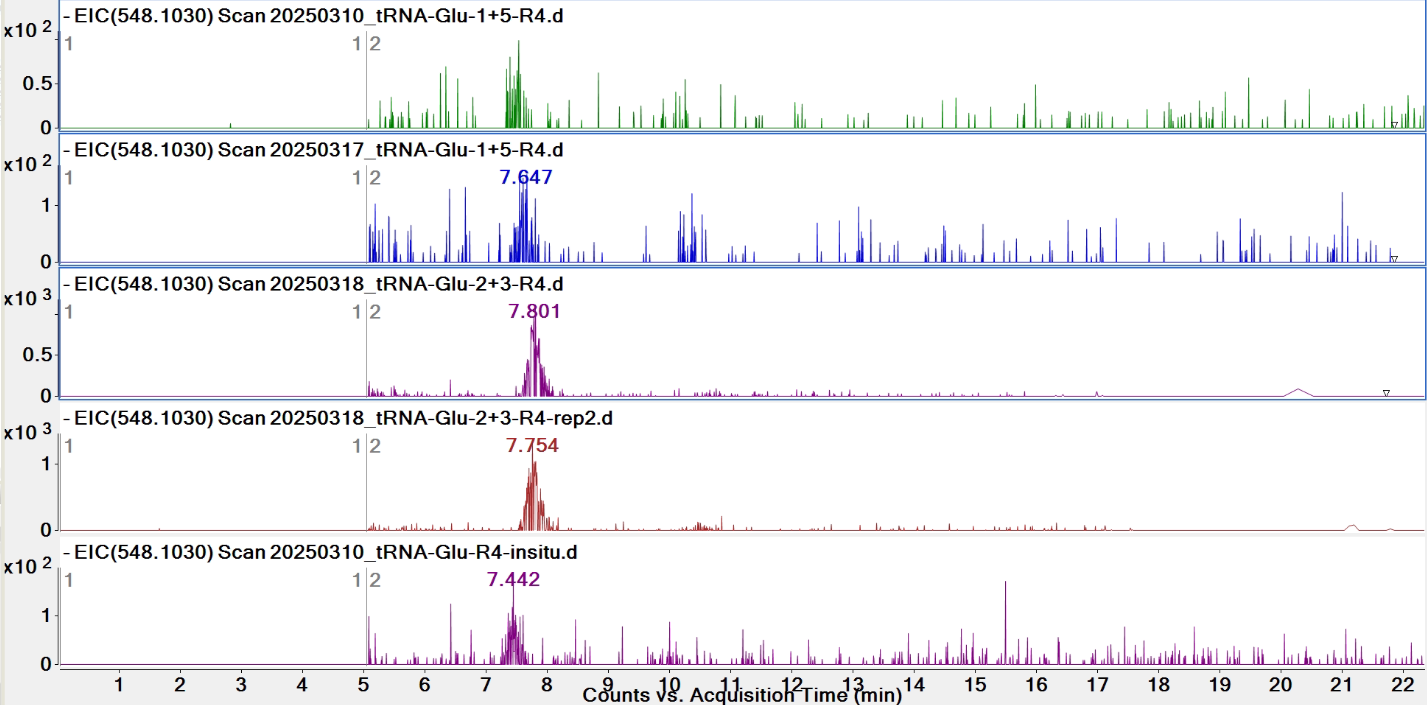

**Pytheas results:** no MS2 match (too short)

| **digested fragment information** | |
| --- | --- |
| **enzyme** | RNase 4 |
| **unmodified sequence** | CGAU |
| **modified sequence** | CG[mA]U |
| **position on *A. c* tRNA^Glu^CUC** | [55-58] |
| **5’ end** | OH |
| **3’ end** | OH |
| **charge state detected** | -2 |
| **theoretical m/z** | unmodified: 610.5976  modified: 617.6054 |
| **ppm used for EIC extraction** | 20 |
| **detection of MS1 signal in data files** | |
| 20250310_tRNA-Glu-1+5-R4 | N; N |
| 20250310_tRNA-Glu-R4-insitu | N; N |
| 20250317_tRNA-Glu-1+5 | N; N |
| 20250318_tRNA-Glu-2+3-R4 | N; N |
| 20250318_tRNA-Glu-2+3-R4-rep2 | N; N |

**EICs**

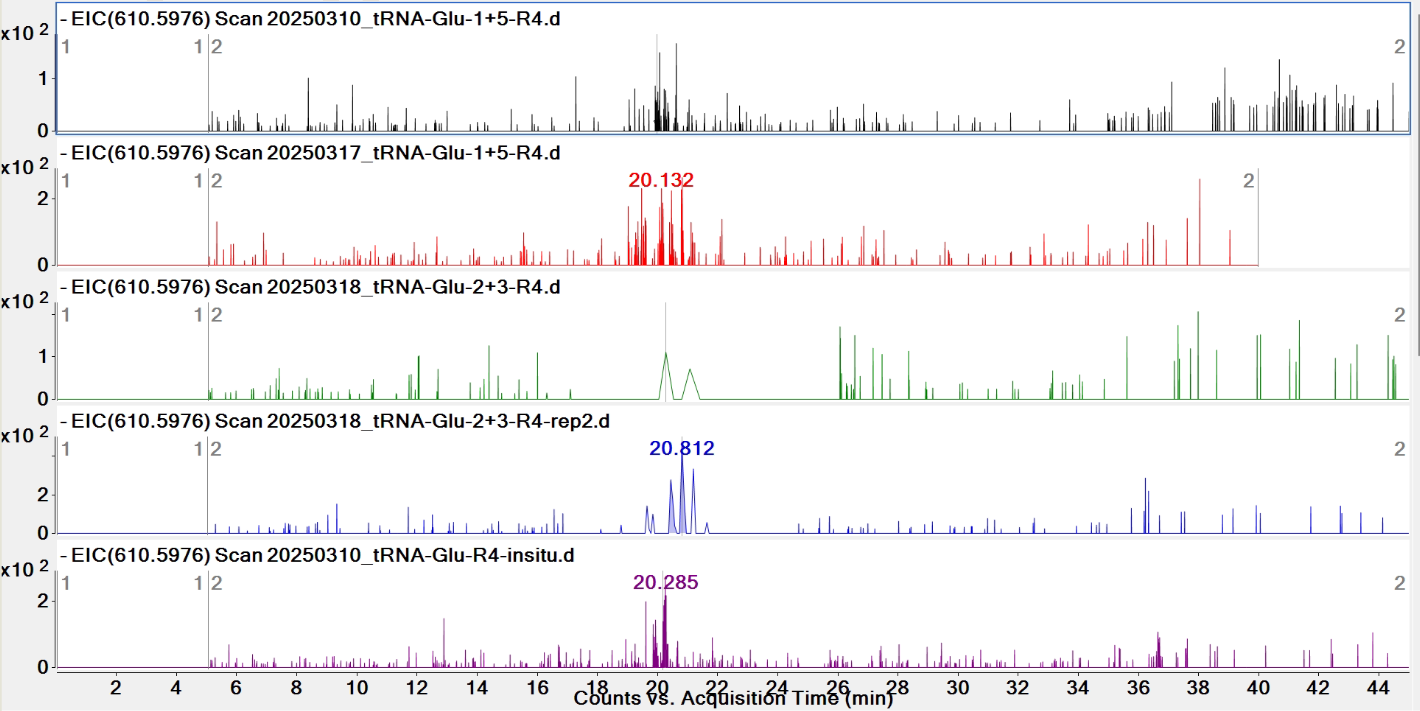

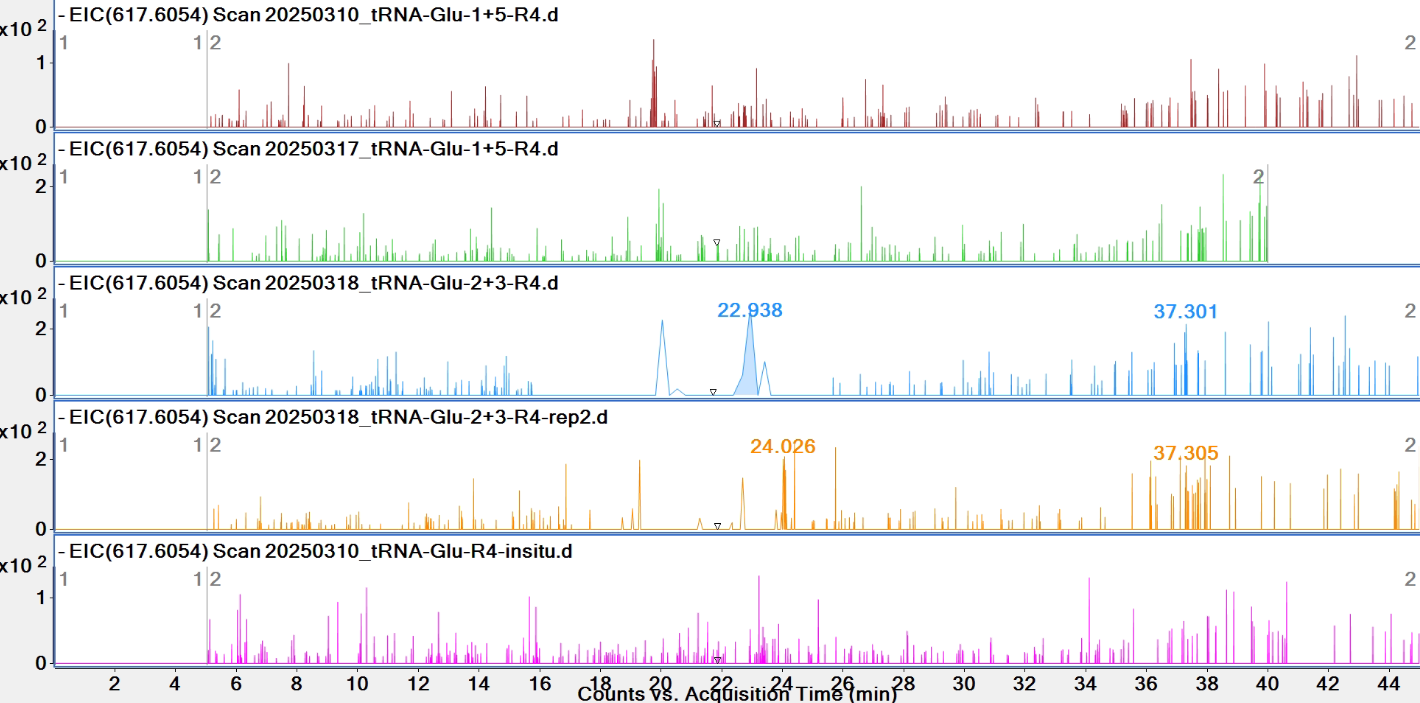

**Pytheas results:** no MS2 match

| **digested fragment information** | |
| --- | --- |
| **enzyme** | RNase 4 |
| **unmodified sequence** | CGAUU |
| **modified sequence** | CG[mA]UU |
| **position on *A. c* tRNA^Glu^CUC** | [55-59] |
| **5' end** | OH |
| **3' end** | OH |
| **charge state detected** | -2 |
| **theoretical *m/z*** | unmodified: 763.6102  modified: 770.6180 |
| **ppm used for EIC extraction** | 20 |
| **detection of MS1 signal in data files** | |
| 20250310_tRNA-Glu-1+5-R4 | N; N |
| 20250310_tRNA-Glu-R4-insitu | N; N |
| 20250317_tRNA-Glu-1+5 | N; N |
| 20250318_tRNA-Glu-2+3-R4 | N; N |
| 20250318_tRNA-Glu-2+3-R4-rep2 | N; N |

**EICs**

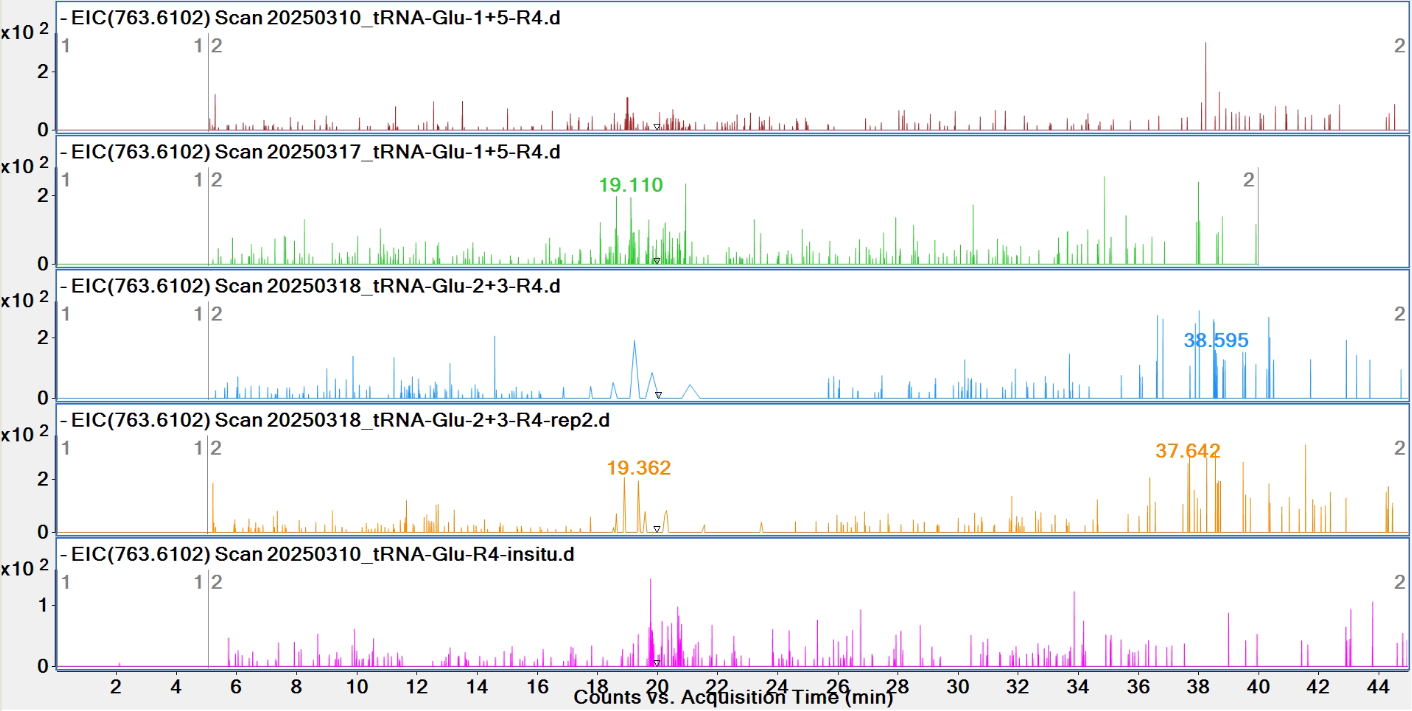

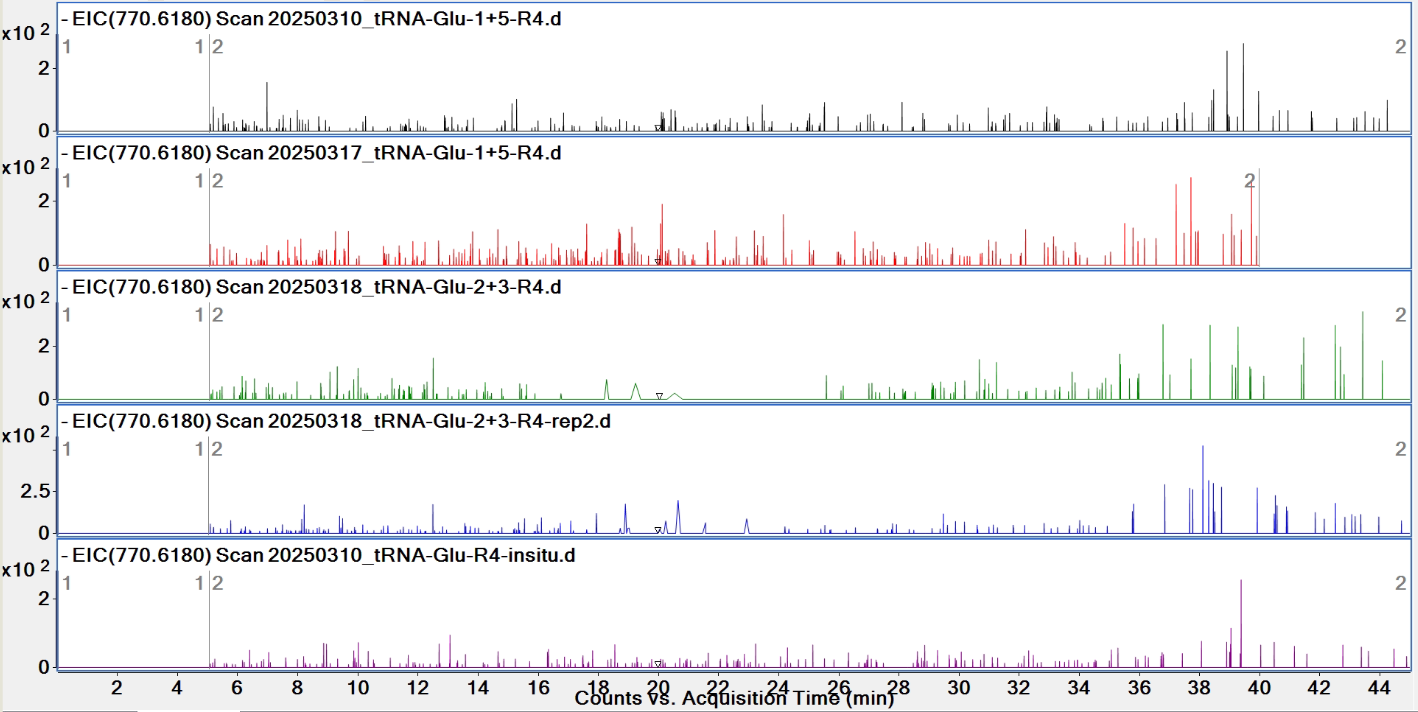

**Pytheas results: no match**

| **digested fragment information** | |
| --- | --- |
| **enzyme** | RNase 4 |
| **unmodified sequence** | CCCCGCU |
| **modified sequence** | N/A |
| **position on *A. c* tRNA^Glu^CUC** | [60-66] |
| **5’ end** | OH |
| **3’ end** | OH |
| **charge state detected** | -2, -3 |
| **theoretical m/z** | 1056.1539, 703.7667 |
| **ppm used for EIC extraction** | 20 |
| **detection of MS1 signal in data files** | |
| 20250310_tRNA-Glu-1+5-R4 | N; N |
| 20250310_tRNA-Glu-R4-insitu | N; N |
| 20250317_tRNA-Glu-1+5 | N; N |
| 20250318_tRNA-Glu-2+3-R4 | N; N |
| 20250318_tRNA-Glu-2+3-R4-rep2 | N; N |

**EICs**

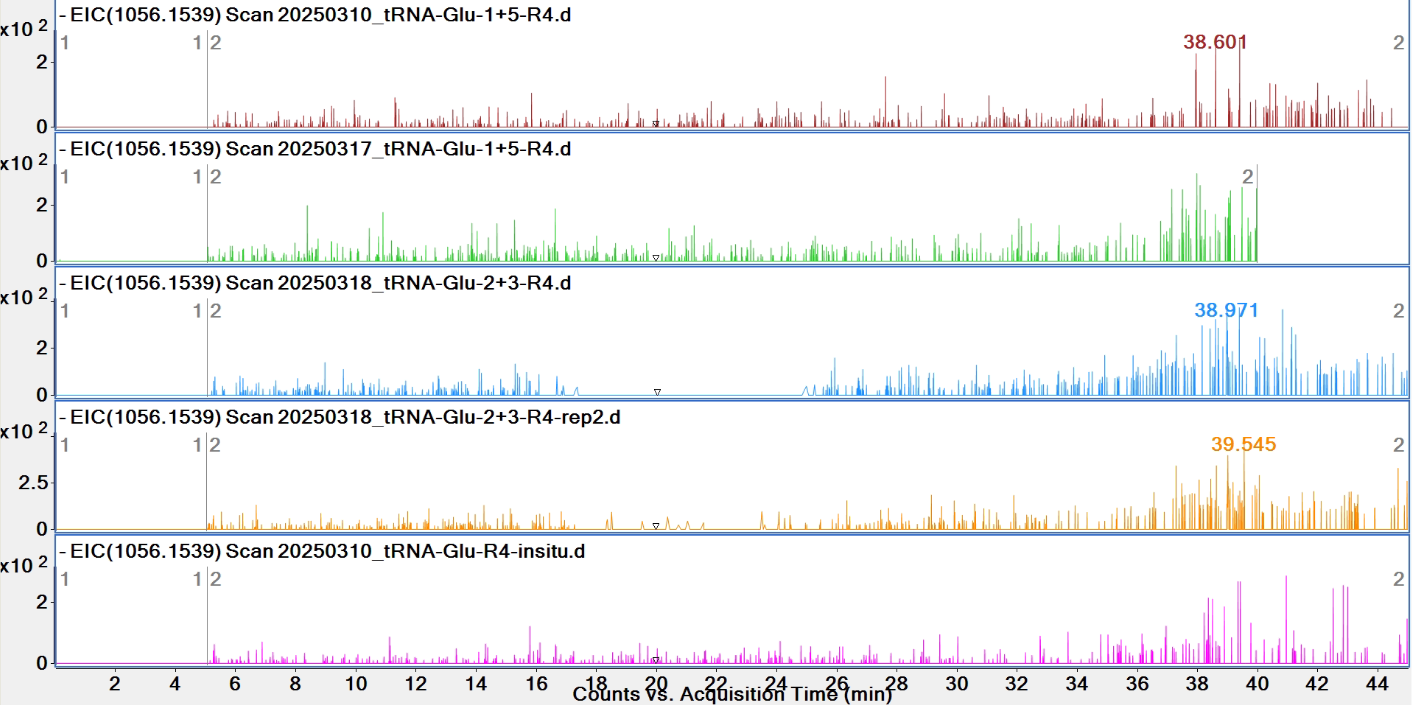

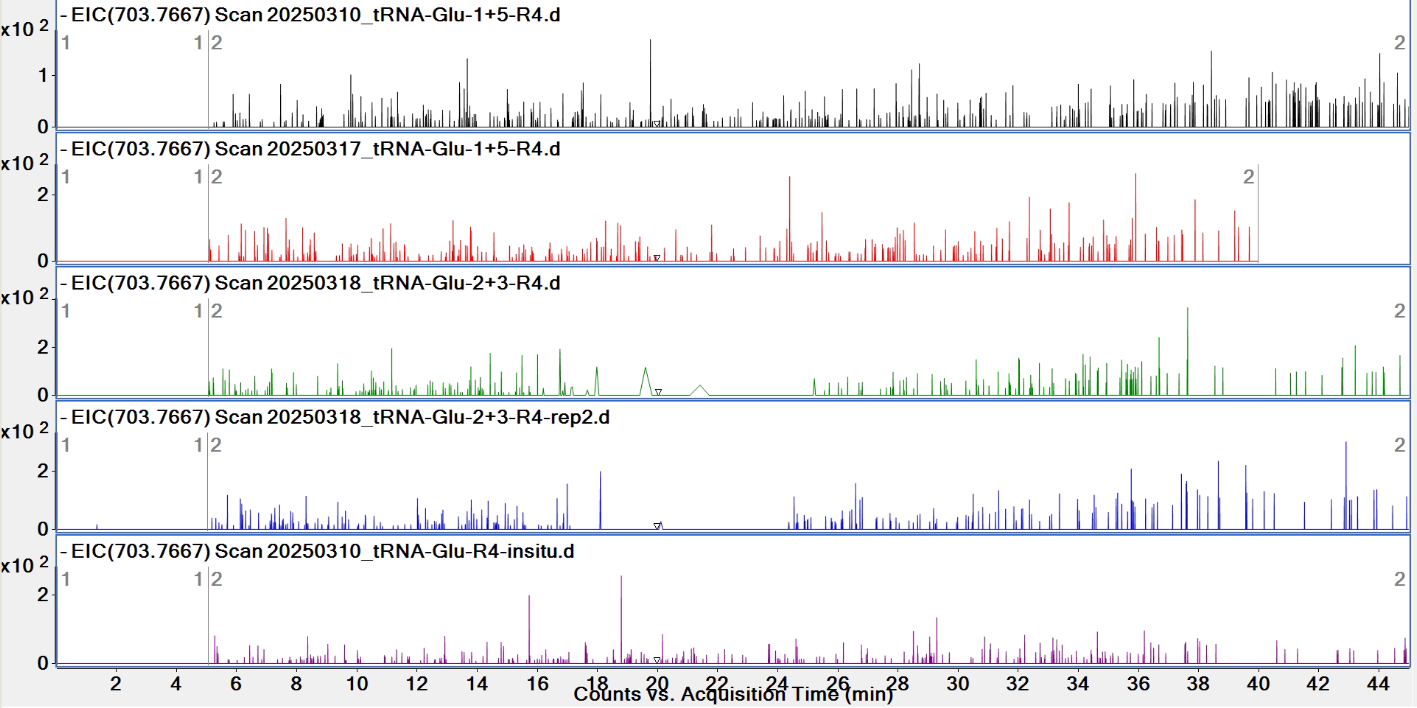

**Pytheas results: no match**

| **digested fragment information** | |
| --- | --- |
| **enzyme** | RNase 4 |
| **unmodified sequence** | CCCCGCUU |
| **modified sequence** | N/A |
| **position on *A. c* tRNA^Glu^CUC** | [60-67] |
| **5’ end** | OH |
| **3’ end** | OH |
| **charge state detected** | -2, -3 |
| **theoretical m/z** | 1209.1665, 805.7751 |
| **ppm used for EIC extraction** | 20 |
| **detection of MS1 signal in data files** | |
| 20250310_tRNA-Glu-1+5-R4 | N; N |
| 20250310_tRNA-Glu-R4-insitu | N; N |
| 20250317_tRNA-Glu-1+5 | N; N |
| 20250318_tRNA-Glu-2+3-R4 | N; N |
| 20250318_tRNA-Glu-2+3-R4-rep2 | N; N |

**EICs**

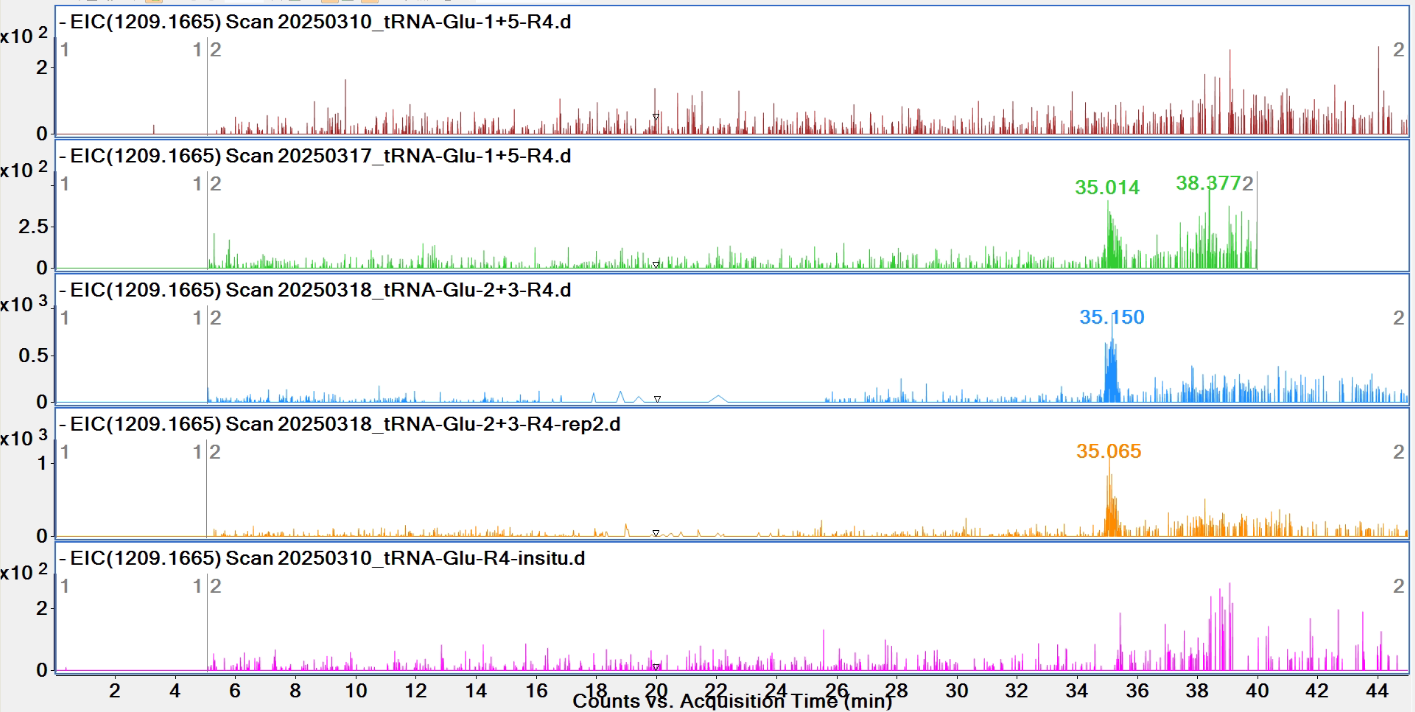

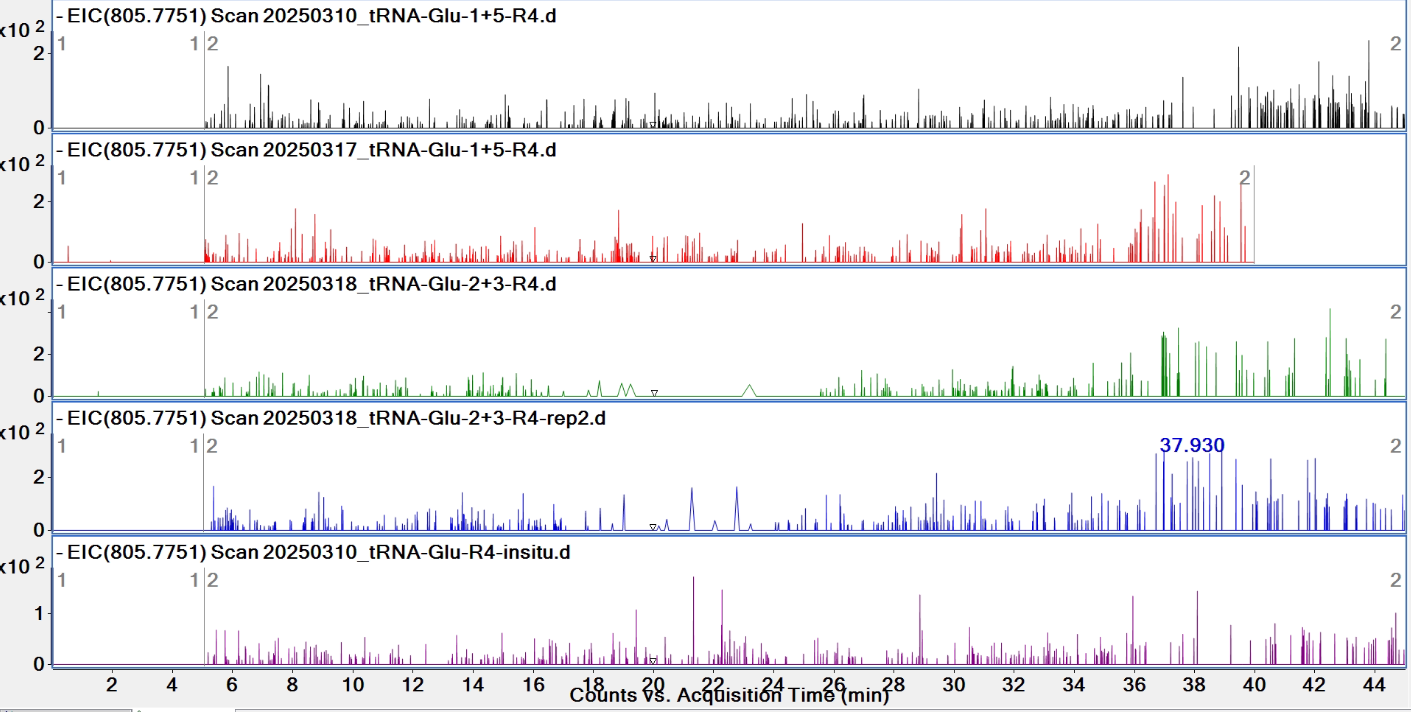

**Pytheas results: no match**

| **digested fragment information** | |
| --- | --- |
| **enzyme** | RNase 4 |
| **unmodified sequence** | GGGAACCA |
| **modified sequence** | N/A |
| **position on *A. c* tRNA^Glu^CUC** | [68-75] |
| **5’ end** | OH |
| **3’ end** | OH |
| **charge state detected** | -2, -3 |
| **theoretical m/z** | 1284.2055, 855.8011 |
| **ppm used for EIC extraction** | 20 |
| **detection of MS1 signal in data files** | |
| 20250310_tRNA-Glu-1+5-R4 | N, Y |
| 20250310_tRNA-Glu-R4-insitu | Y, Y |
| 20250317_tRNA-Glu-1+5 | Y, Y |
| 20250318_tRNA-Glu-2+3-R4 | Y, Y |
| 20250318_tRNA-Glu-2+3-R4-rep2 | Y, Y |

**EICs**

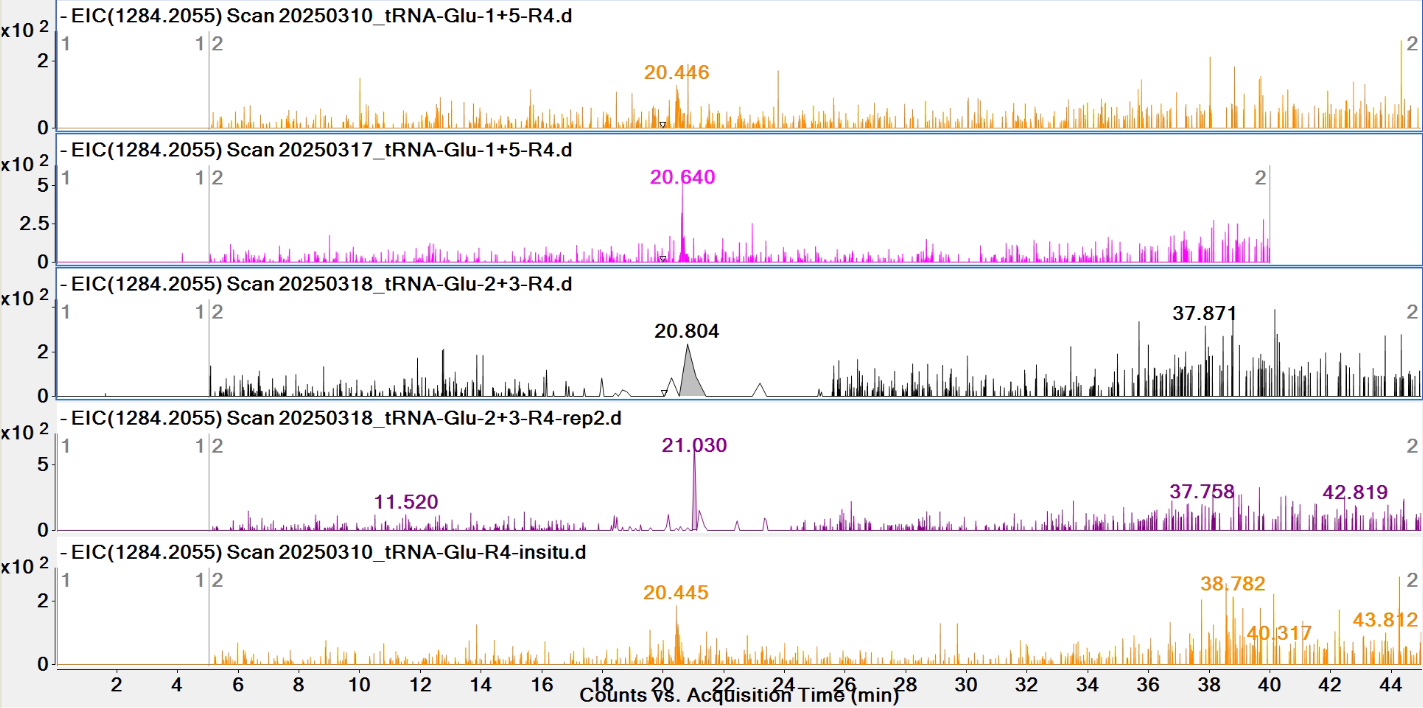

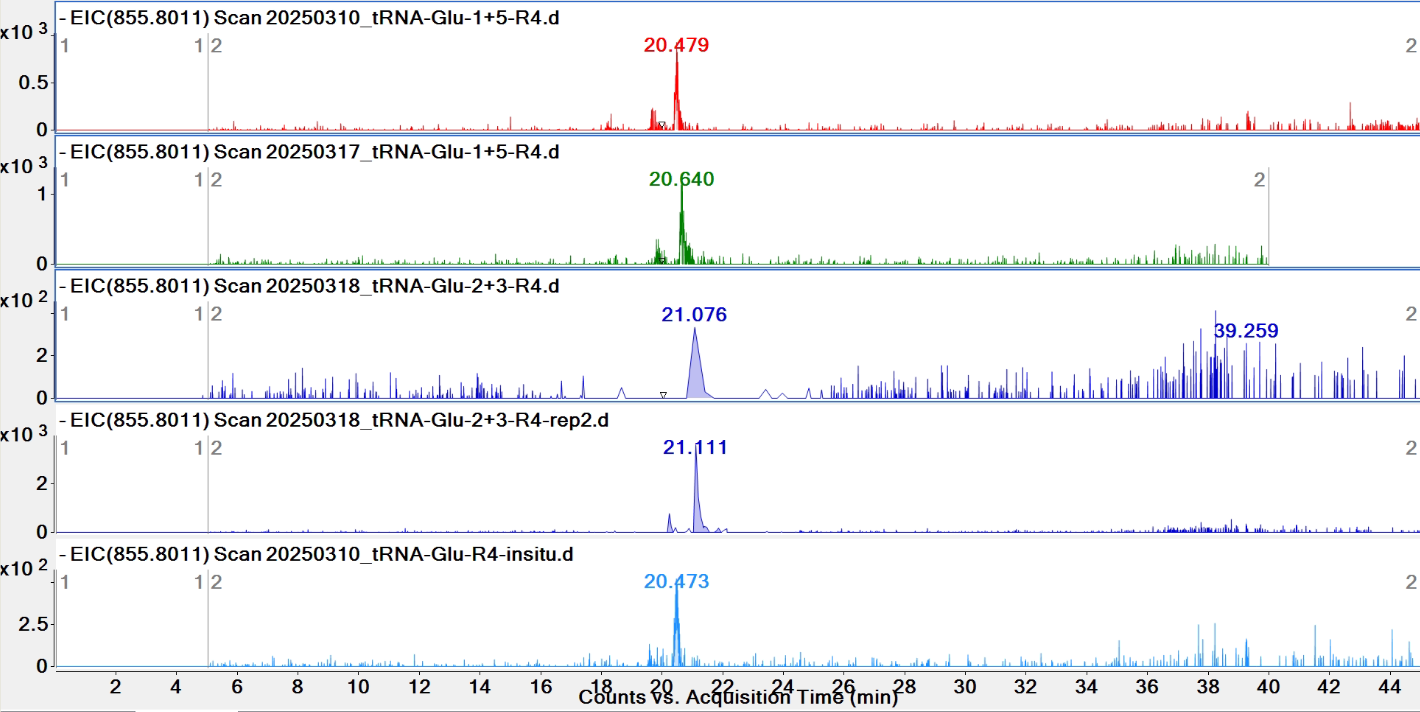

**Pytheas results**

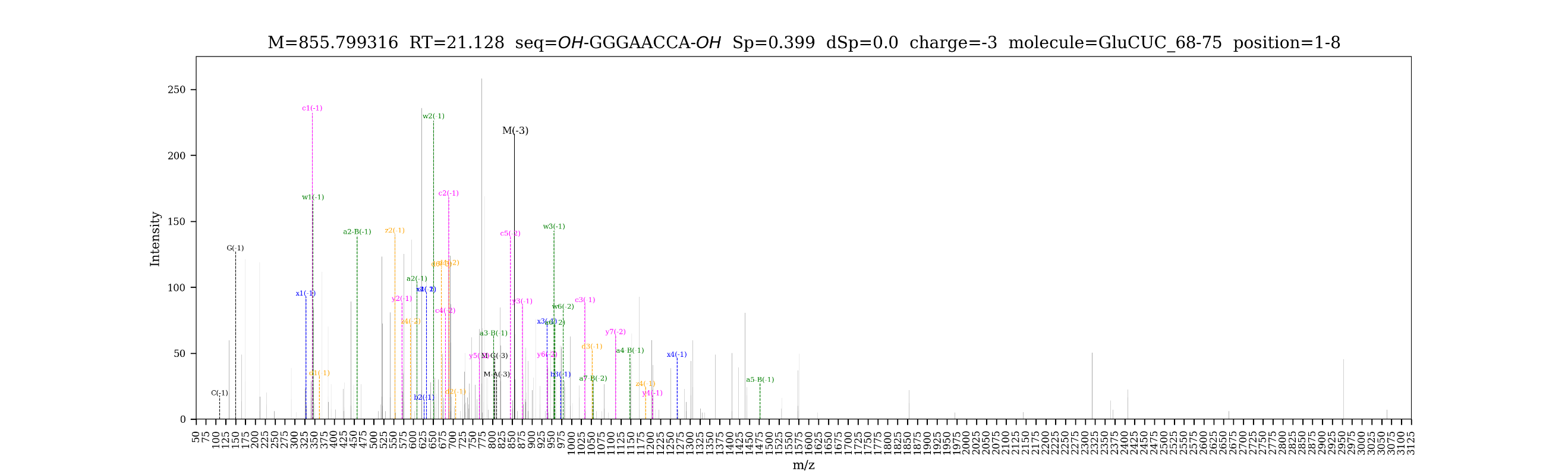

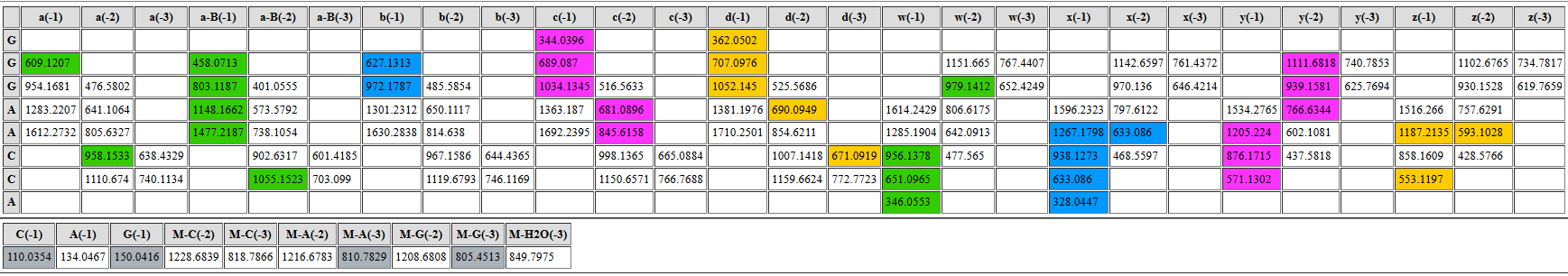

**Part 2. RNase T1 data**

| **list of LC-MS/MS data files used for analysis by RNase T1 digestion** | |
| --- | --- |
| **data file name** | **tissue** |
| 20250514_Glu_2-6_T1 | heart (pooled from multiple animals) |
| 20250514_Glu_7-11-A_T1 | heart (pooled from multiple animals) |
| 20250514_Glu_unk3_T1 | spermatheca (pooled from multiple animals) |
| 20250518_Glu_combined | heat + spermatheca |
| 20250524_tRNA_Glu_T1_heart* | heart (pooled from multiple animals) |
| 20250626_Ac#16_muscle_Glu-2_T1 | muscle (from one animal) |
| *Pytheas results are based on this data file unless otherwise indicated | |

| **digested fragment information** | |
| --- | --- |
| **enzyme** | RNase T1 |
| **unmodified sequence** | UCCCAAG |
| **modified sequence** | N/A |
| **position on *A. c* tRNA^Glu^CUC** | [1-7] |
| **5' end** | OH |
| **3' end** | OH |
| **charge state detected** | -2, -3 |
| **theoretical m/z** | 1080.1651, 719.7741 |
| **ppm used for EIC extraction** | 20 |
| **detection of MS1 signal in data files** | |
| 20250514_Glu_2-6_T1 | Y, Y |
| 20250514_Glu_7-11-A_T1 | Y, Y |
| 20250514_Glu_unk3_T1 | Y, Y |
| 20250518_Glu_combined | Y, Y |
| 20250524_tRNA_Glu_T1_heart | Y, Y |
| 20250626_Ac#16_muscle_Glu-2_T1 | Y, Y |

**EICs**

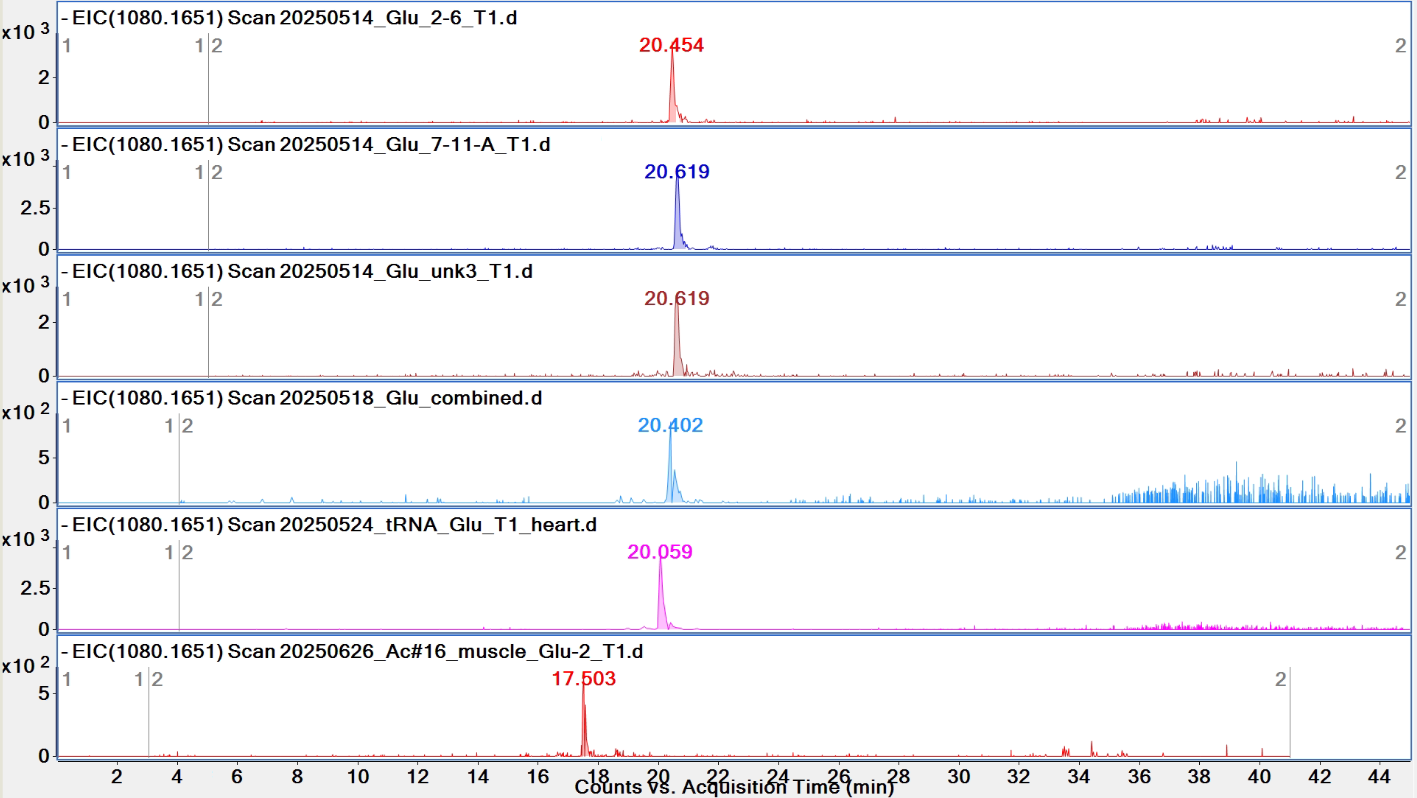

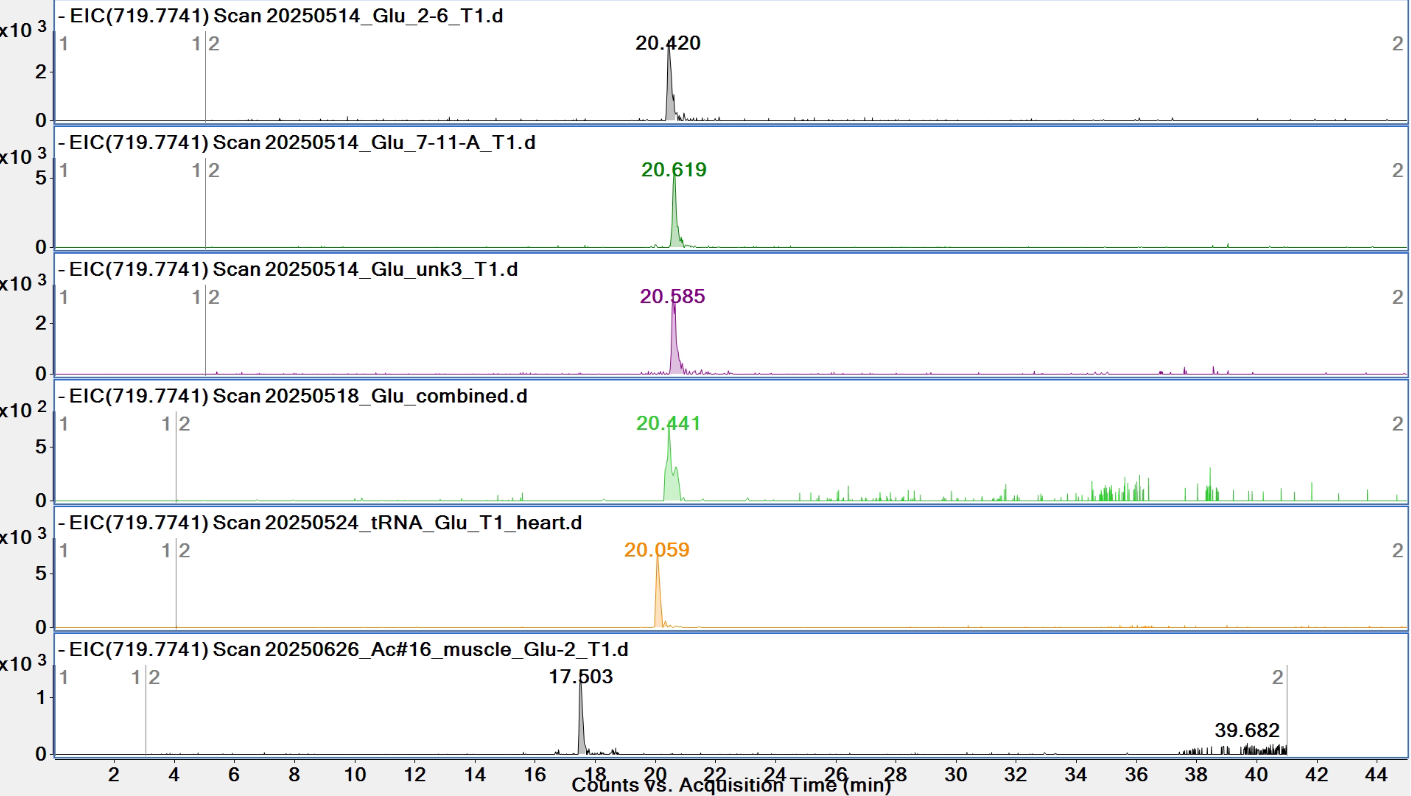

**Pytheas results**

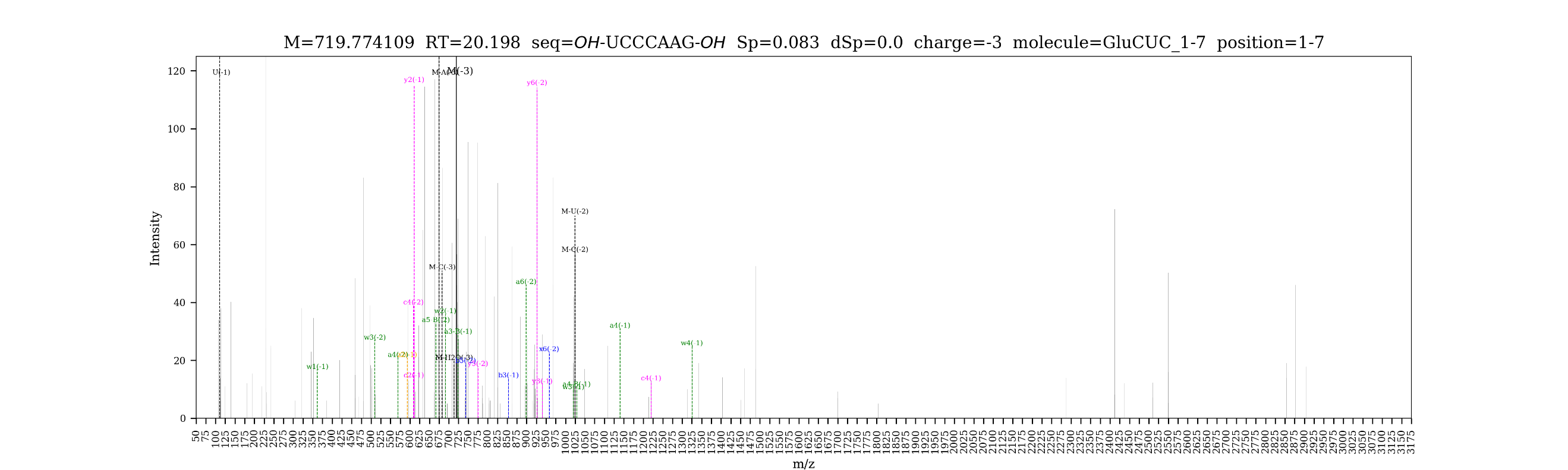

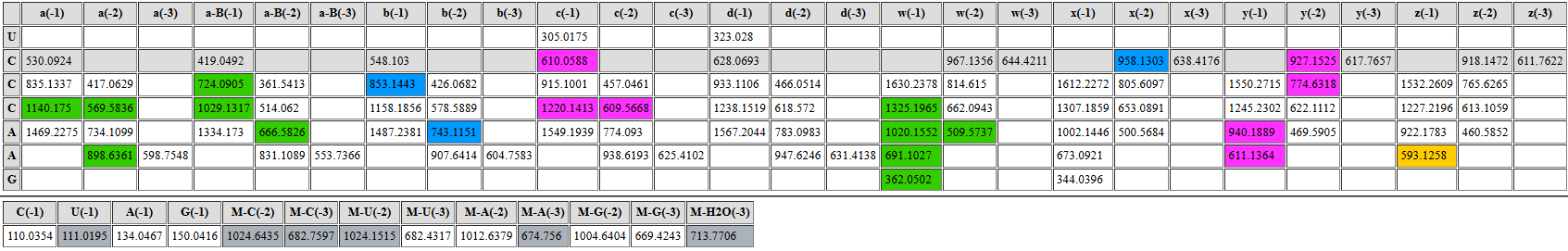

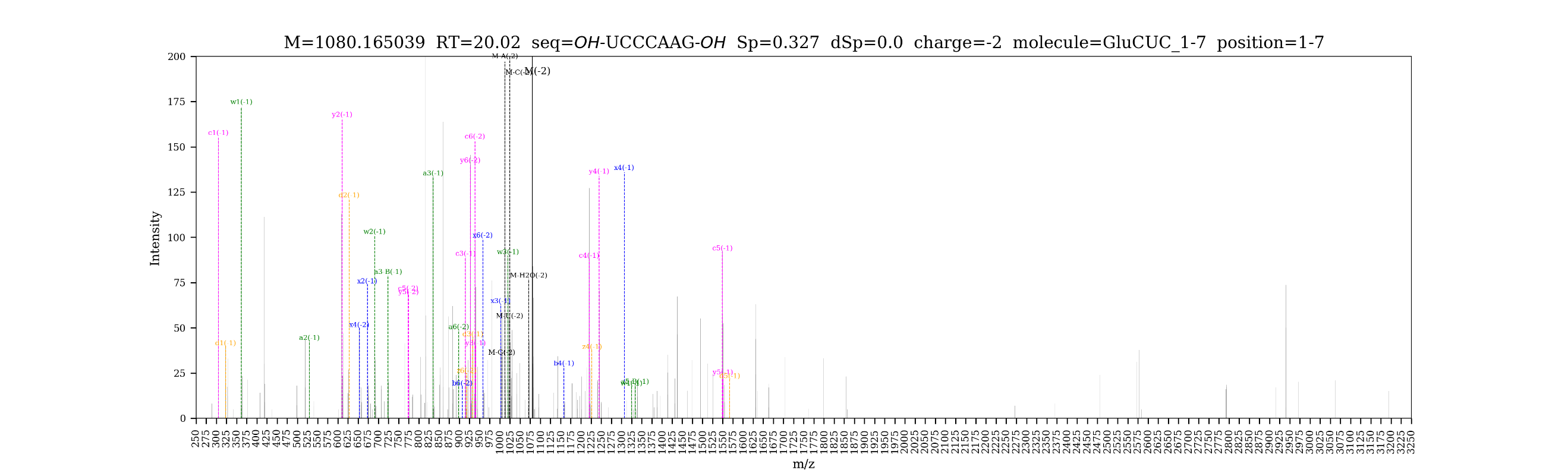

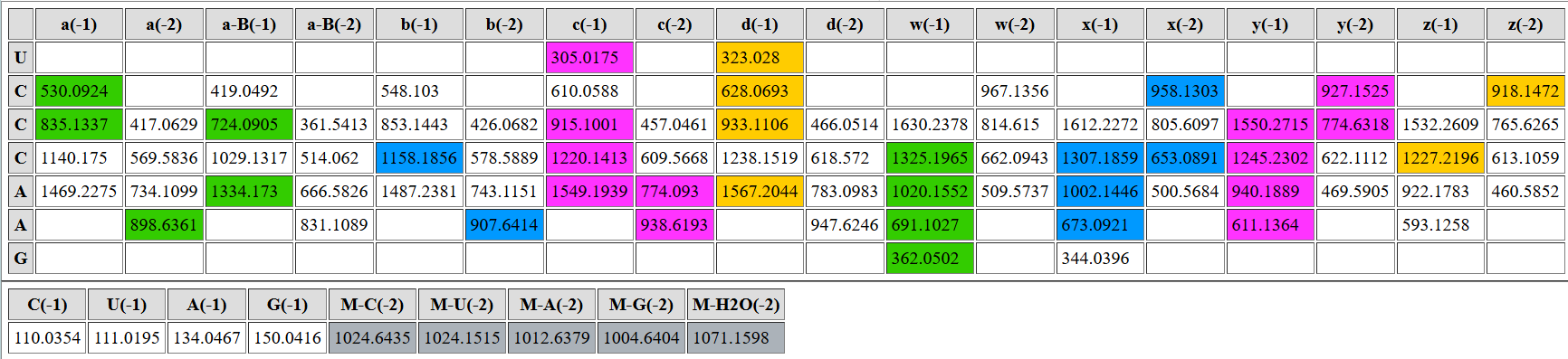

| **digested fragment information** | |
| --- | --- |
| **enzyme** | RNase T1 |
| **unmodified sequence** | UG |
| **modified sequence** | N/A |
| **position on *A. c* tRNA^Glu^CUC** | [8-9] |
| **5’ end** | OH |
| **3’ end** | OH |
| **charge state detected** | -1 |
| **theoretical m/z** | 588.1092 |
| **ppm used for EIC extraction** | 20 |
| **detection of MS1 signal in data files** | |
| 20250514_Glu_2-6_T1 | Y |
| 20250514_Glu_7-11-A_T1 | Y |
| 20250514_Glu_unk3_T1 | Y |
| 20250518_Glu_combined | Y |
| 20250524_tRNA_Glu_T1_heart | Y |
| 20250626_Ac#16_muscle_Glu-2_T1 | Y |

**EICs**

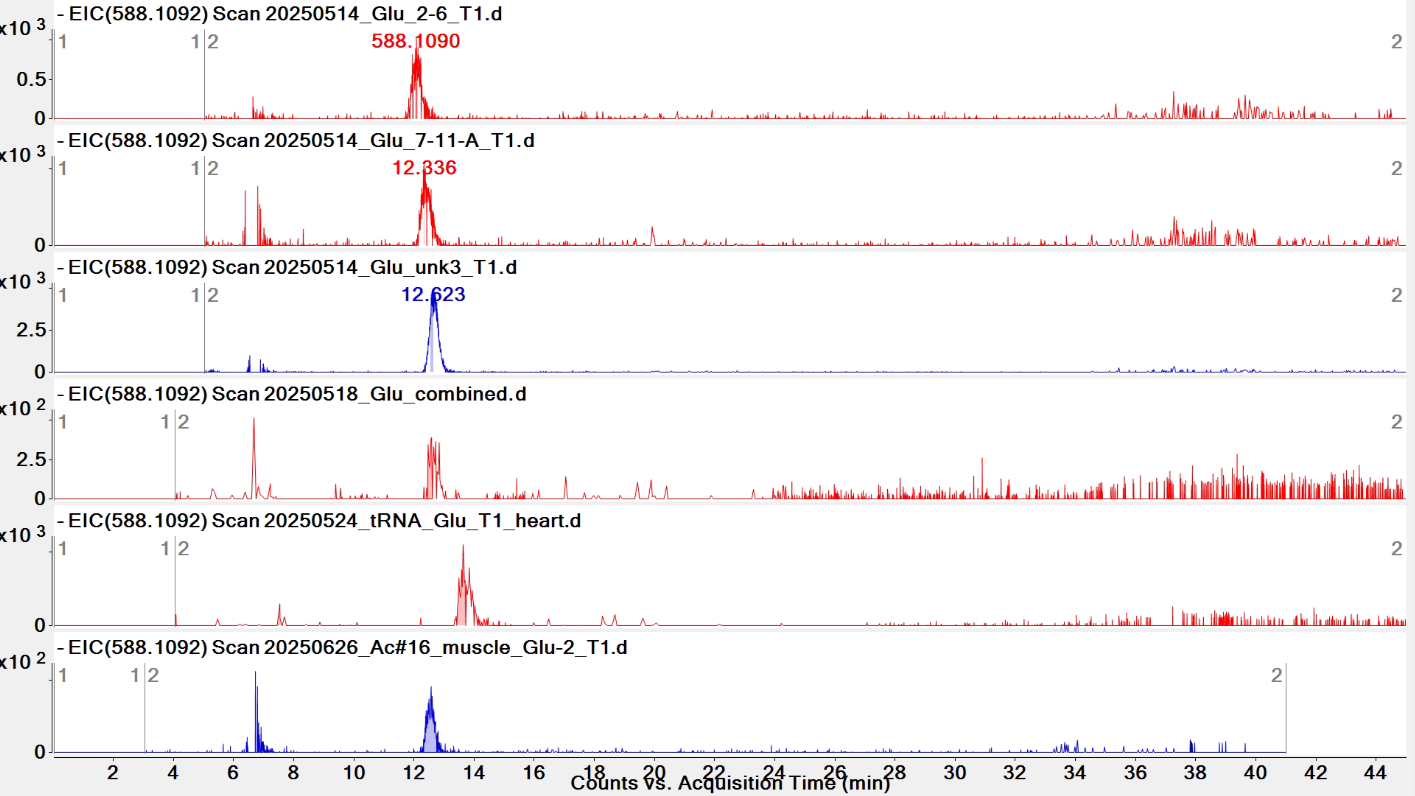

**Pytheas results: no match**

| **digested fragment information** | |
| --- | --- |
| **enzyme** | RNase T1 |
| **unmodified sequence** | UCUAG |
| **modified sequence** | N/A |
| **position on *A. c* tRNA^Glu^CUC** | [11-15] |
| **5’ end** | OH |
| **3’ end** | OH |
| **charge state detected** | -2 |
| **theoretical m/z** | 763.6102 |
| **ppm used for EIC extraction** | 20 |
| **detection of MS1 signal in data files** | |
| 20250514_Glu_2-6_T1 | Y |
| 20250514_Glu_7-11-A_T1 | Y |
| 20250514_Glu_unk3_T1 | Y |
| 20250518_Glu_combined | Y |
| 20250524_tRNA_Glu_T1_heart | Y |
| 20250626_Ac#16_muscle_Glu-2_T1 | Y |

**EICs**

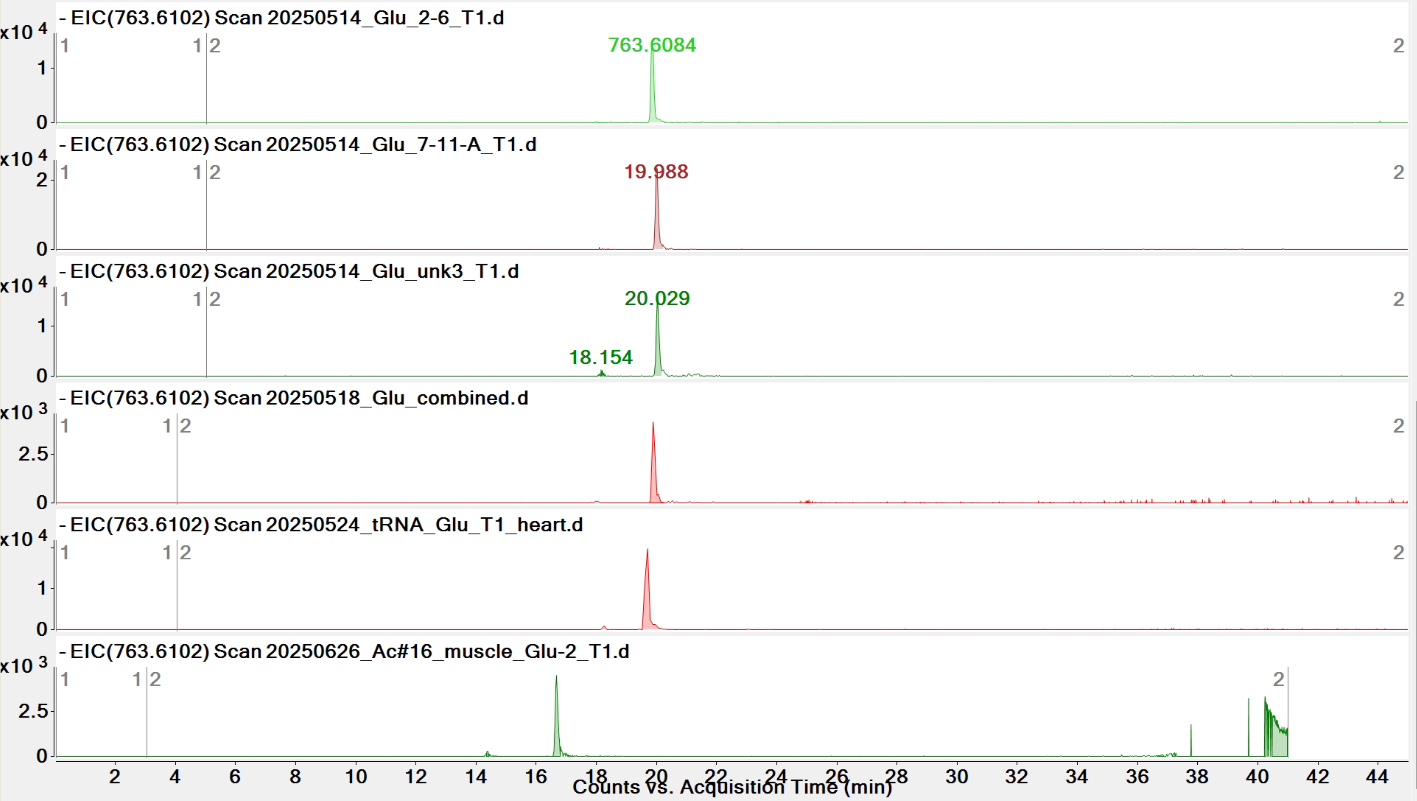

**Pytheas results**

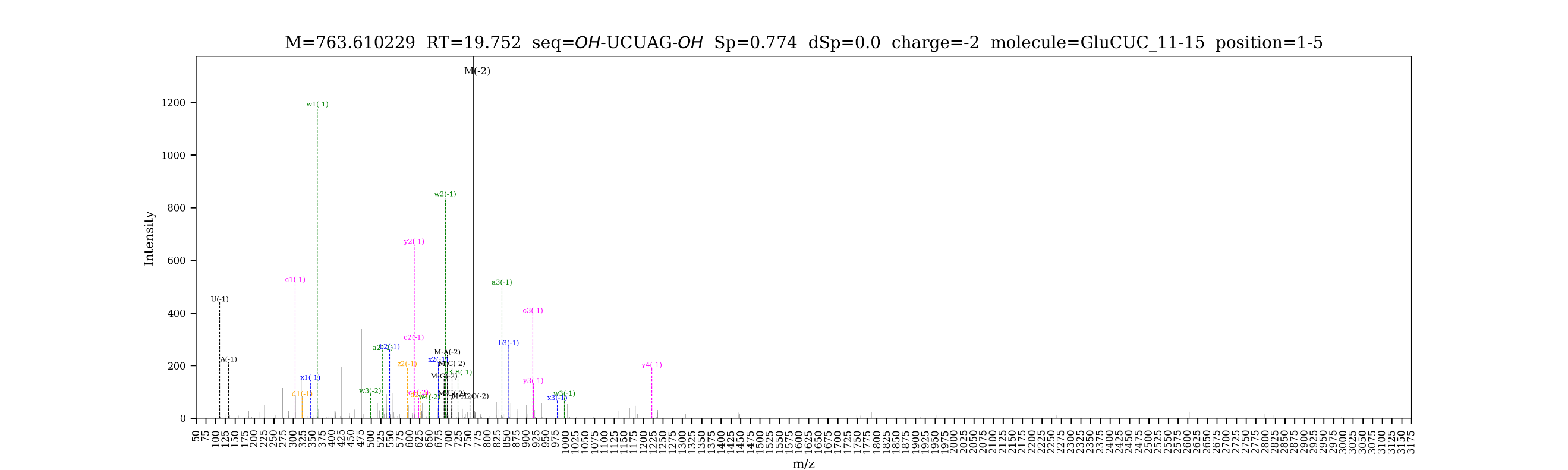

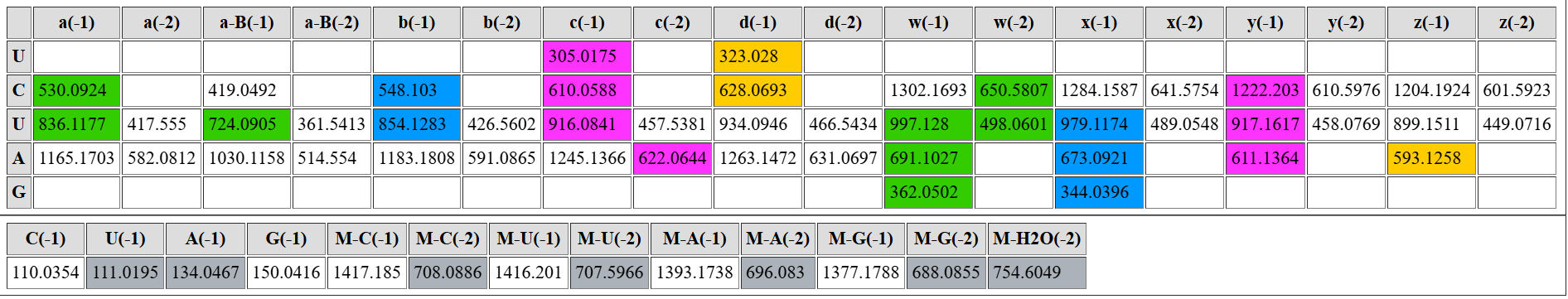

| **digested fragment information** | |
| --- | --- |
| **enzyme** | RNase T1 |
| **unmodified sequence** | UG |
| **modified sequence** | DG |
| **position on *A. c* tRNA^Glu^CUC** | [16-17] |
| **5' end** | OH |
| **3' end** | OH |
| **charge state detected** | -1 |
| **theoretical m/z** | unmodified: 588.1092  modified: 590.1248 |
| **ppm used for EIC extraction** | 20 |
| **detection of MS1 signal in data files** | |
| 20250514_Glu_2-6_T1 | Y; Y |
| 20250514_Glu_7-11-A_T1 | Y; Y |
| 20250514_Glu_unk3_T1 | Y; Y |
| 20250518_Glu_combined | Y; Y |
| 20250524_tRNA_Glu_T1_heart | Y; Y |
| 20250626_Ac#16_muscle_Glu-2_T1 | Y; Y |

**EICs**

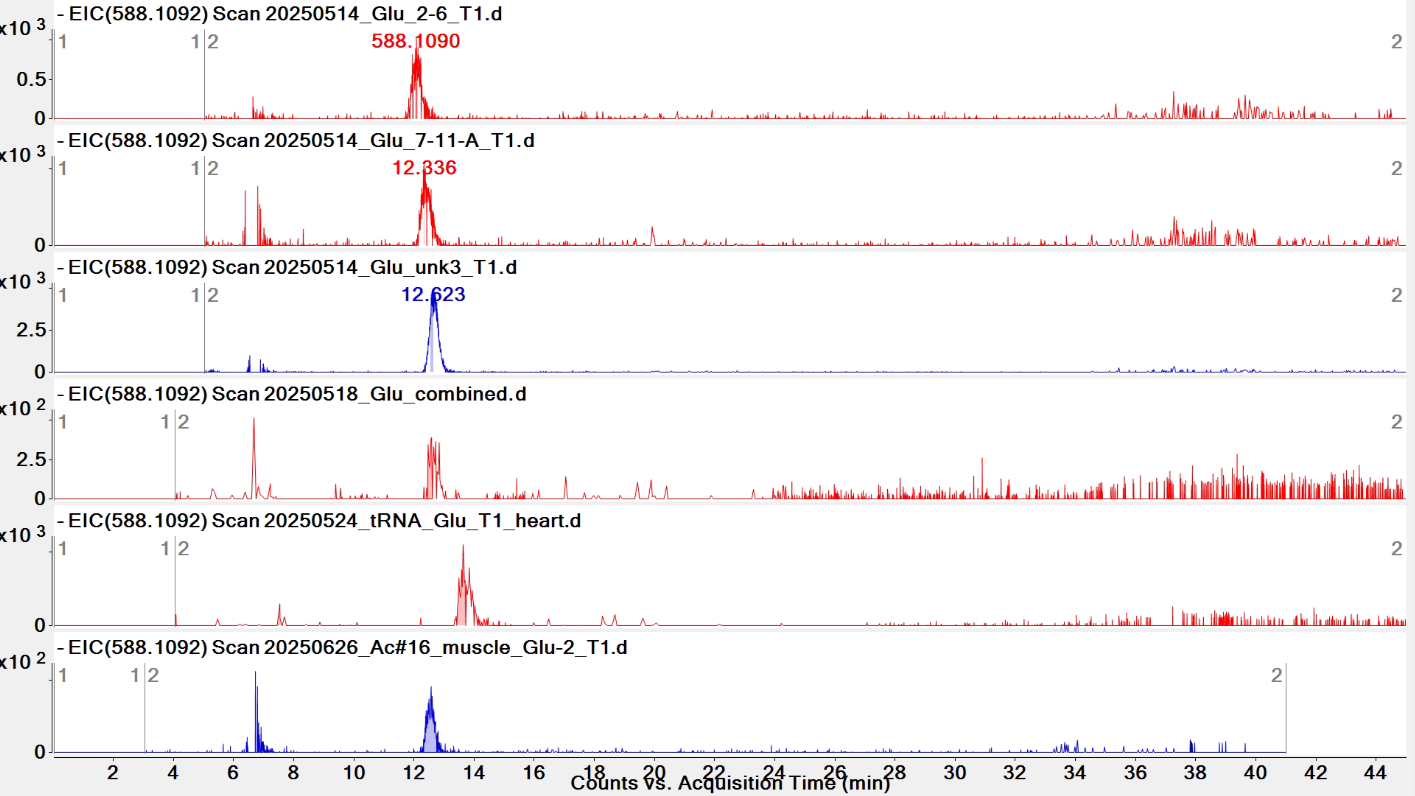

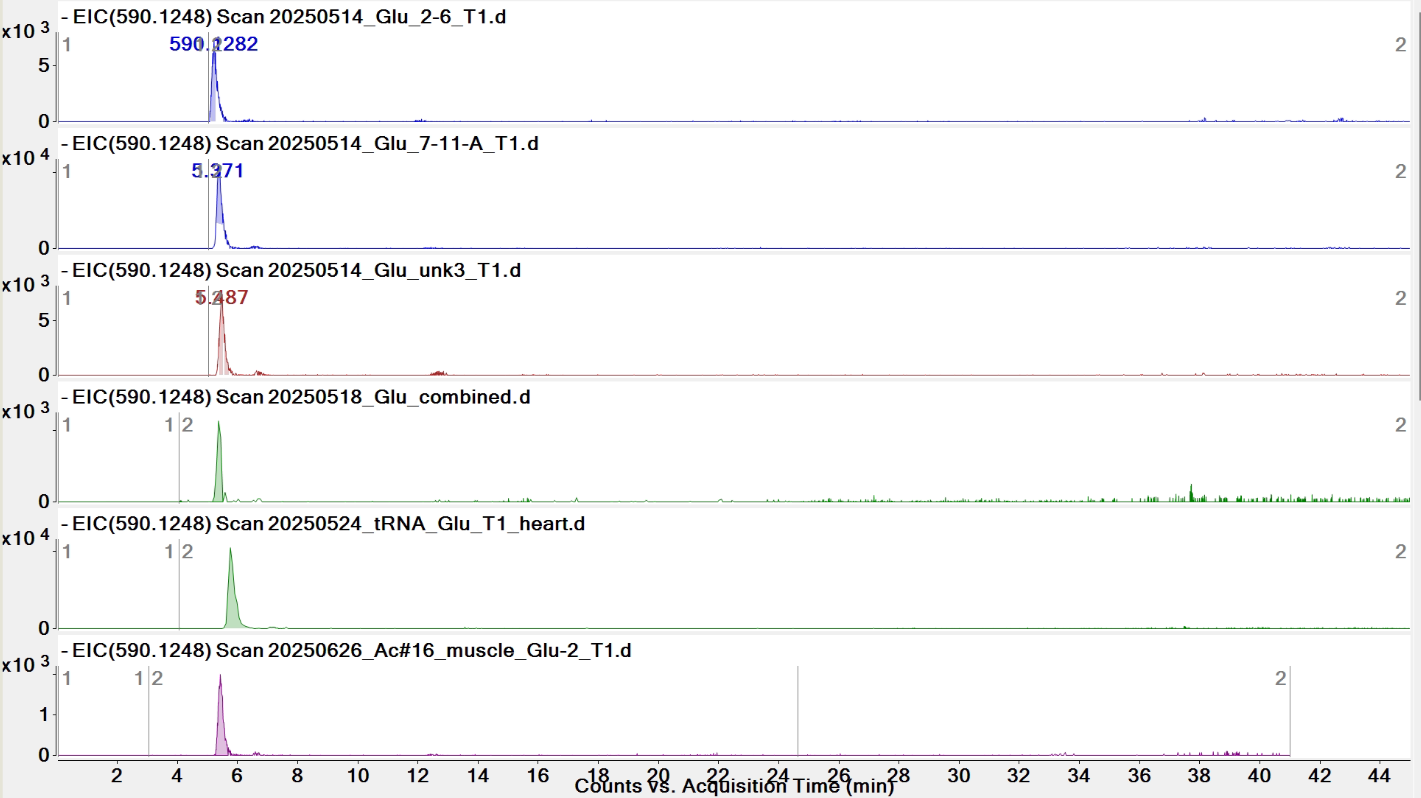

**Pytheas results: no match**

**Comparison of UG/DG EICs**

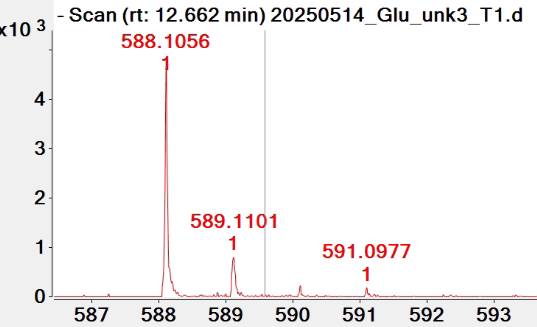

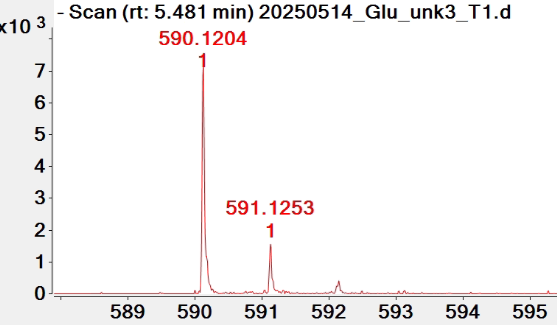

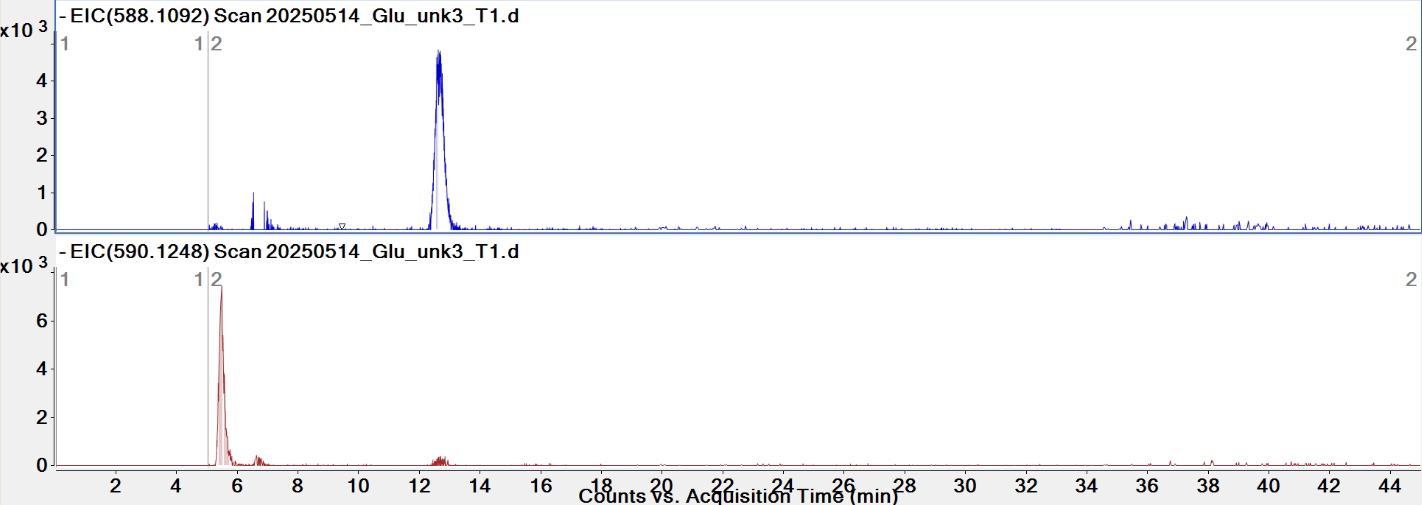

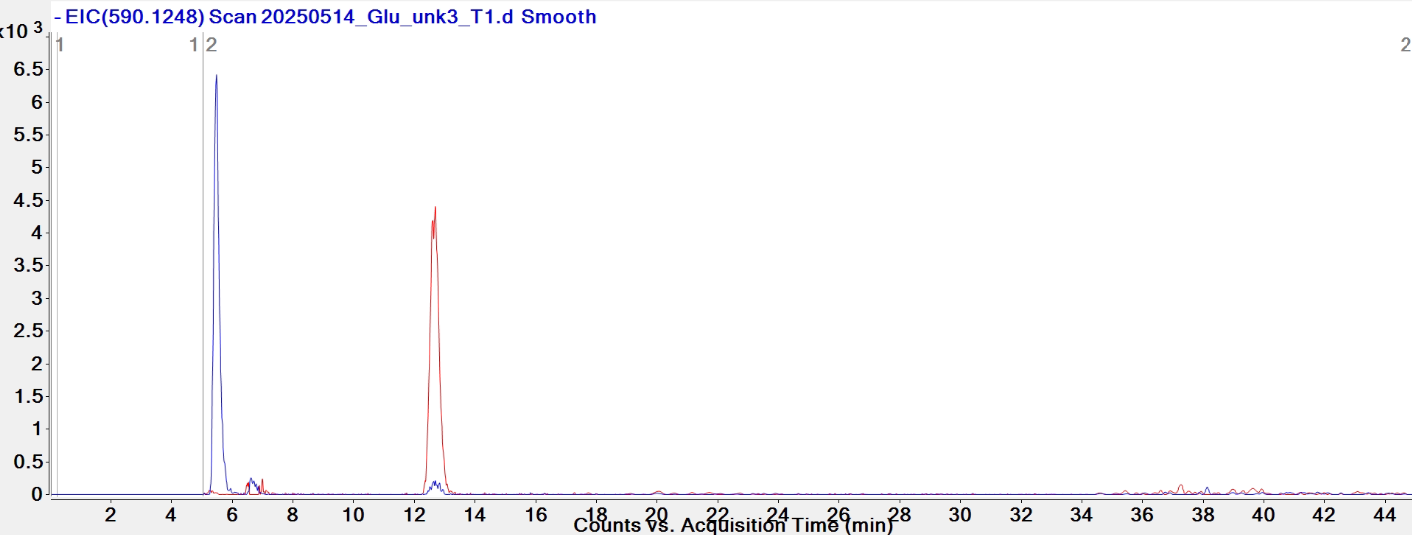

accessory genital mass

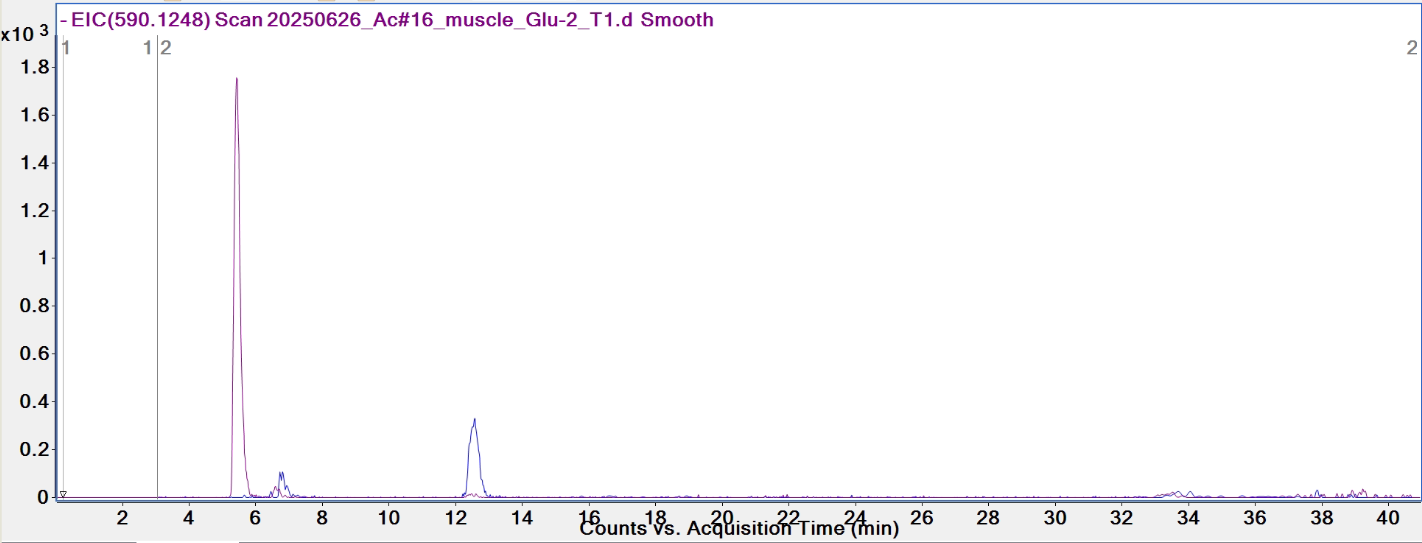

muscle

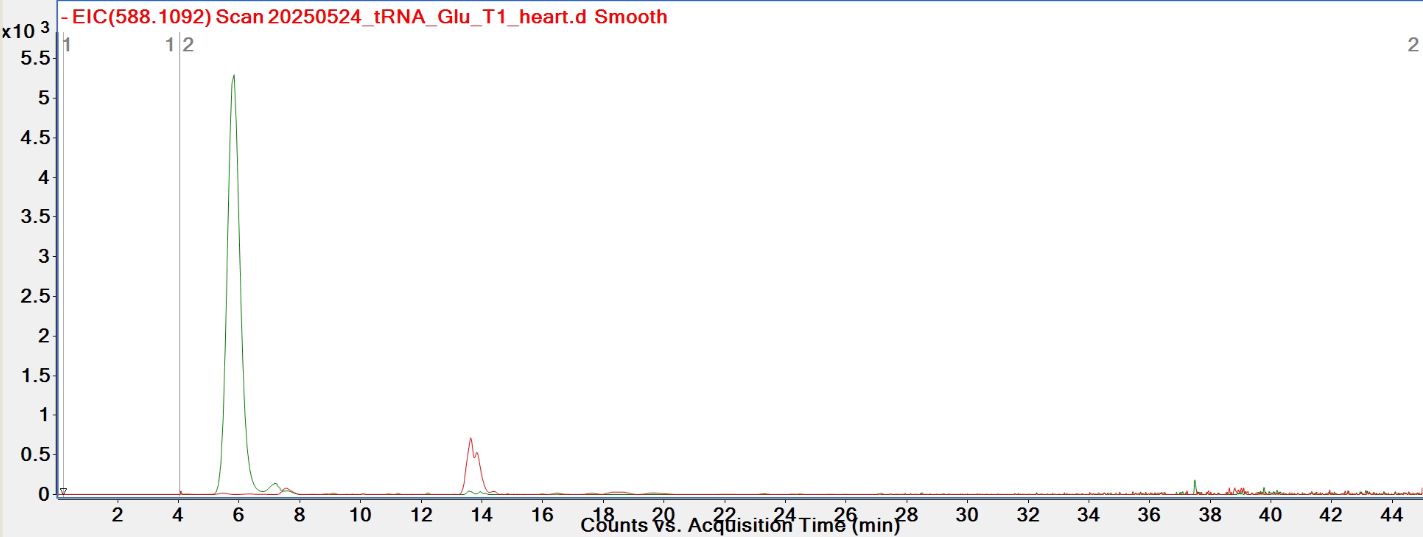

heart

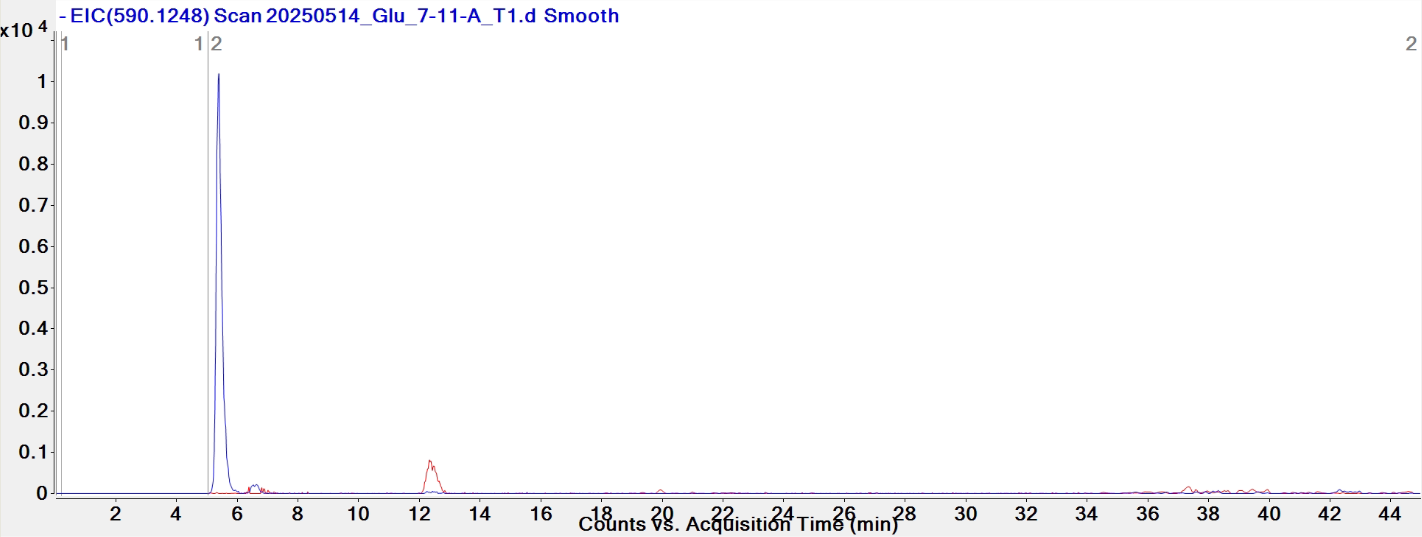

heart

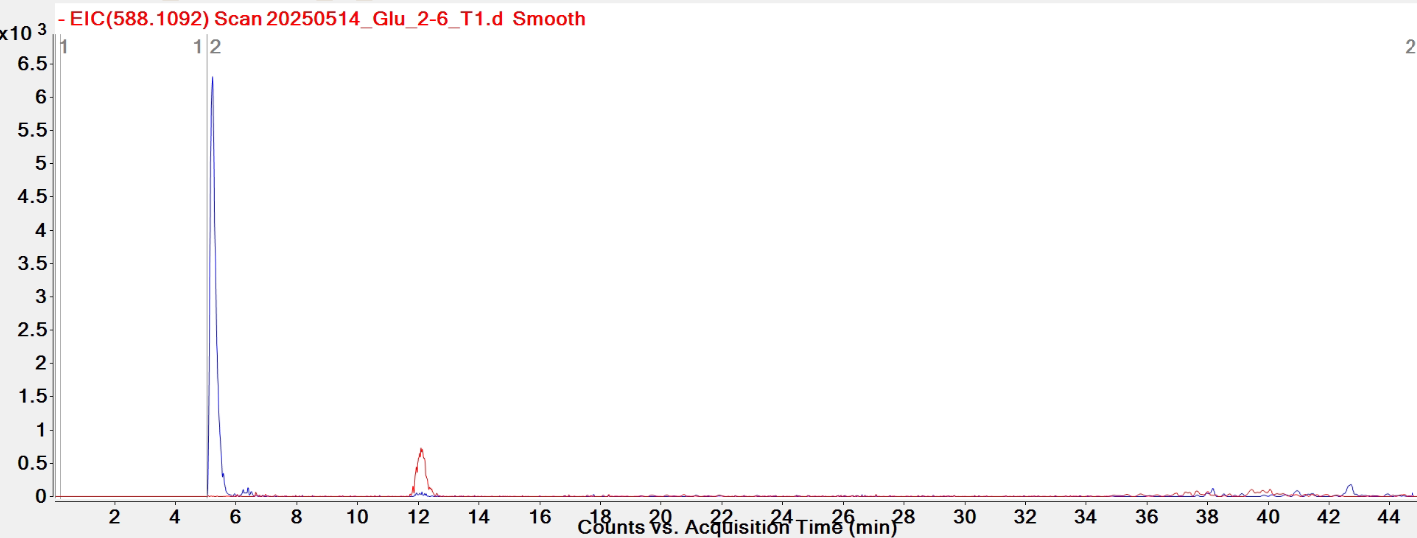

heart

| **digested fragment information** | |
| --- | --- |
| **enzyme** | RNase T1 |
| **unmodified sequence** | UUAG |
| **modified sequence** | DDAG |
| **position on *A. c* tRNA^Glu^CUC** | [19-22] |
| **5' end** | OH |
| **3' end** | OH |
| **charge state detected** | -2 |
| **theoretical m/z** | 613.1052 |
| **ppm used for EIC extraction** | 20 |
| **detection of MS1 signal in data files** | |
| 20250514_Glu_2-6_T1 | Y |
| 20250514_Glu_7-11-A_T1 | Y |
| 20250514_Glu_unk3_T1 | Y |
| 20250518_Glu_combined | Y |
| 20250524_tRNA_Glu_T1_heart | Y |
| 20250626_Ac#16_muscle_Glu-2_T1 | Y |

**EICs**

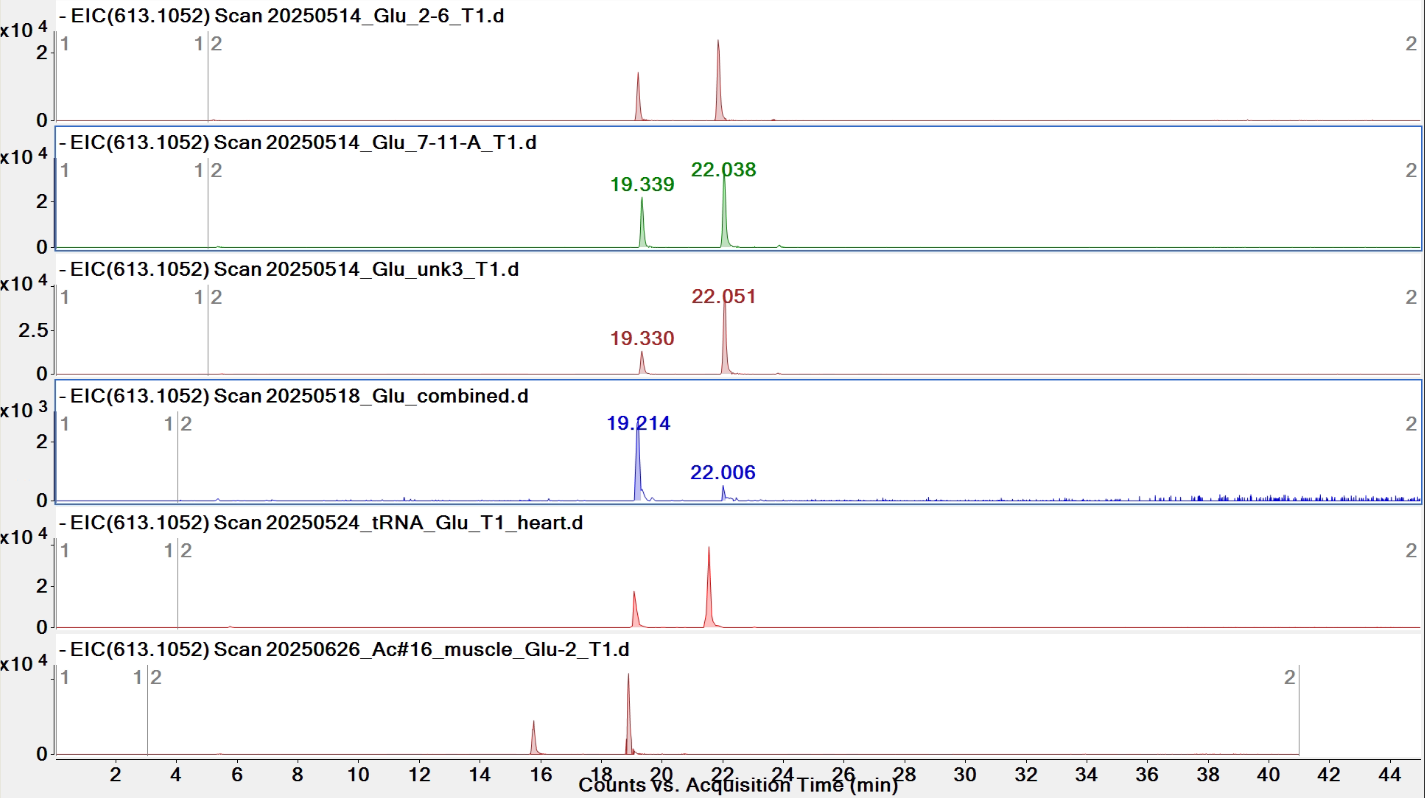

*The later peak is for [m5Um]UCG (*z* = -2)

**Pytheas results**

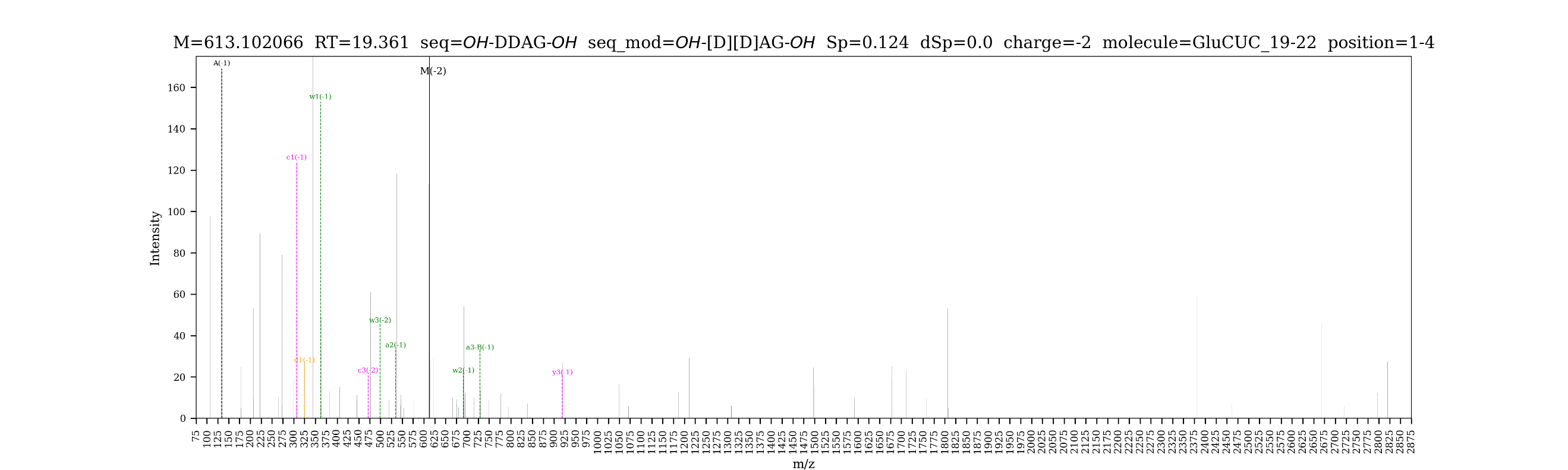

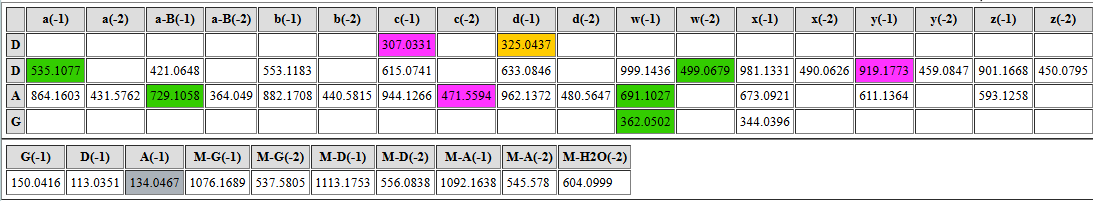

| **digested fragment information** | |
| --- | --- |
| **enzyme** | RNase T1 |
| **unmodified sequence** | AUUUCG |
| **modified sequence** | N/A |
| **position on *A. c* tRNA^Glu^CUC** | [24-29] |
| **5' end** | OH |
| **3' end** | OH |
| **charge state detected** | -2, -3 |
| **theoretical m/z** | 916.6229, 610.7460 |
| **ppm used for EIC extraction** | 20 |
| **detection of MS1 signal in data files** | |
| 20250514_Glu_2-6_T1 | Y, Y |
| 20250514_Glu_7-11-A_T1 | Y, Y |
| 20250514_Glu_unk3_T1 | Y, Y |
| 20250518_Glu_combined | Y, Y |
| 20250524_tRNA_Glu_T1_heart | Y, Y |
| 20250626_Ac#16_muscle_Glu-2_T1 | Y, Y |

**EICs**

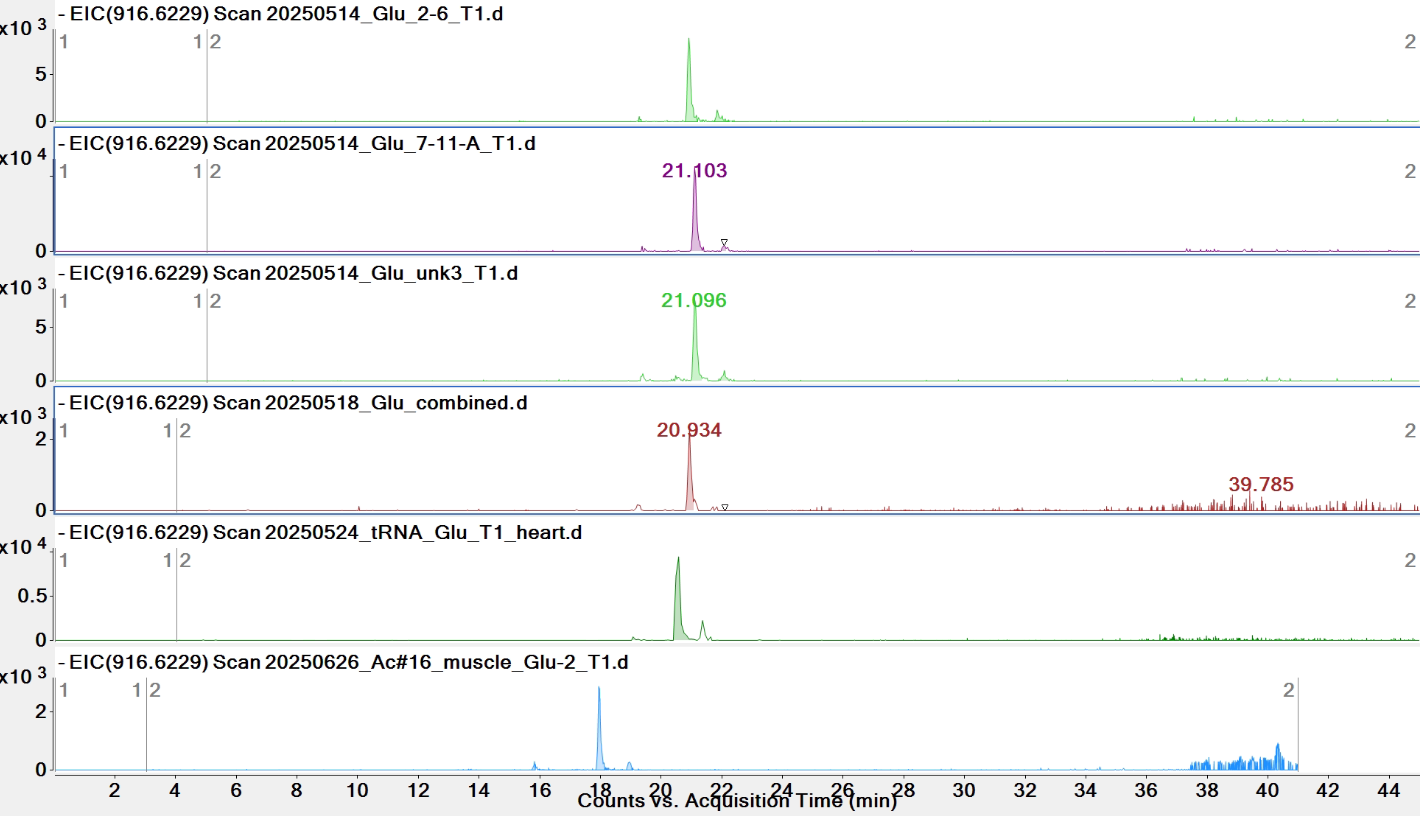

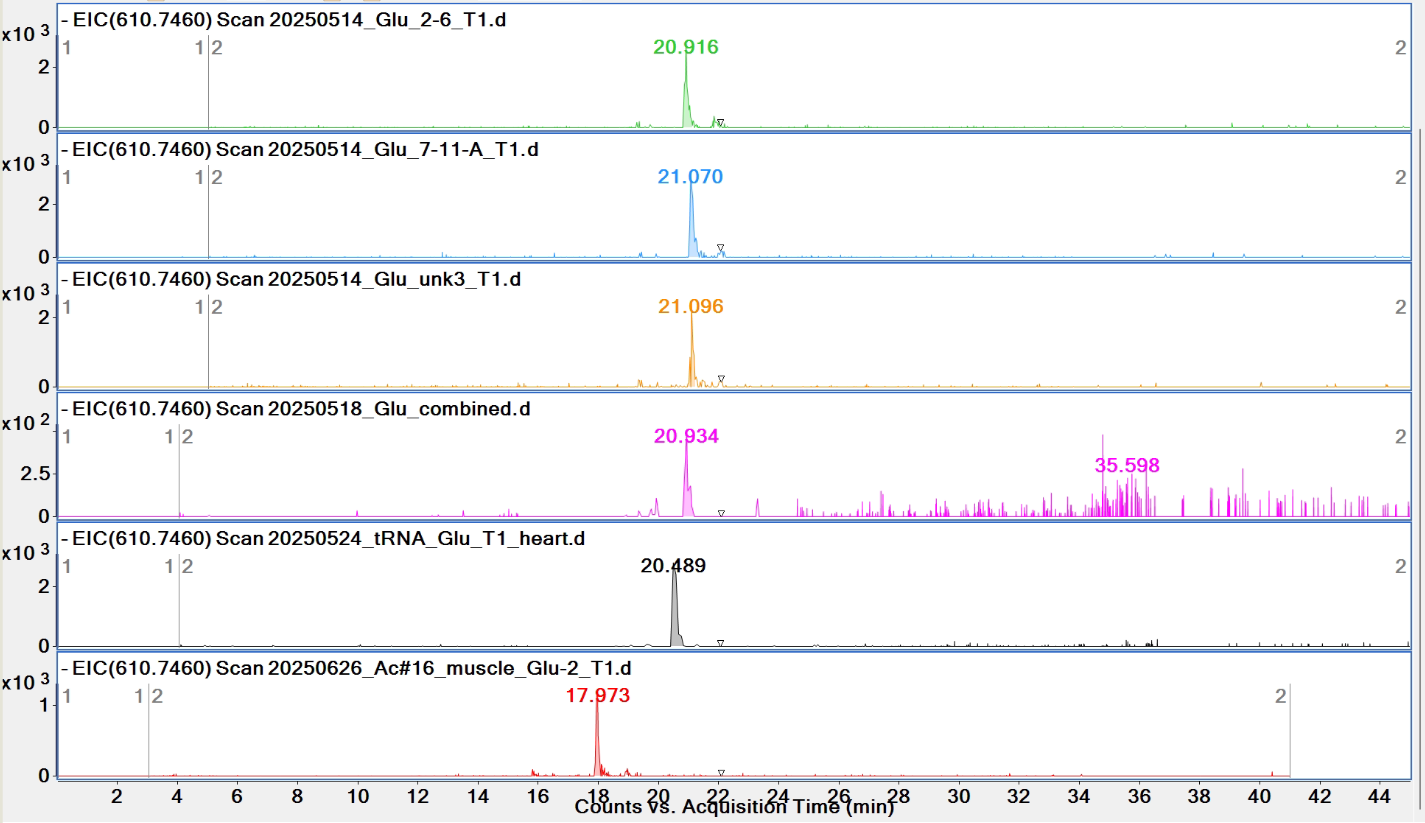

**Pytheas results**

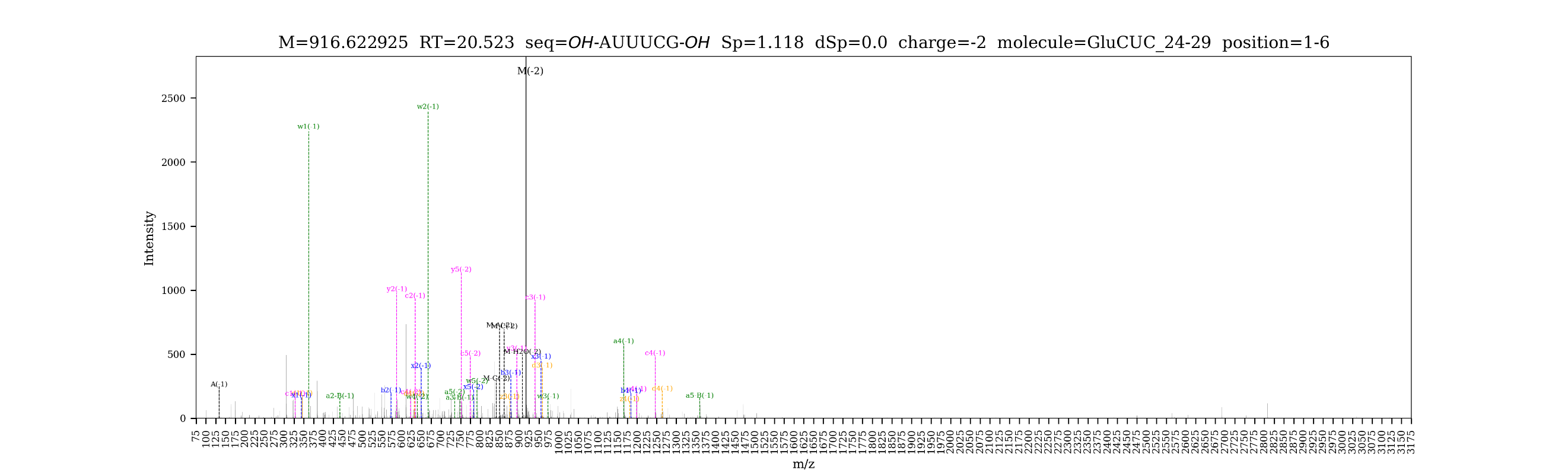

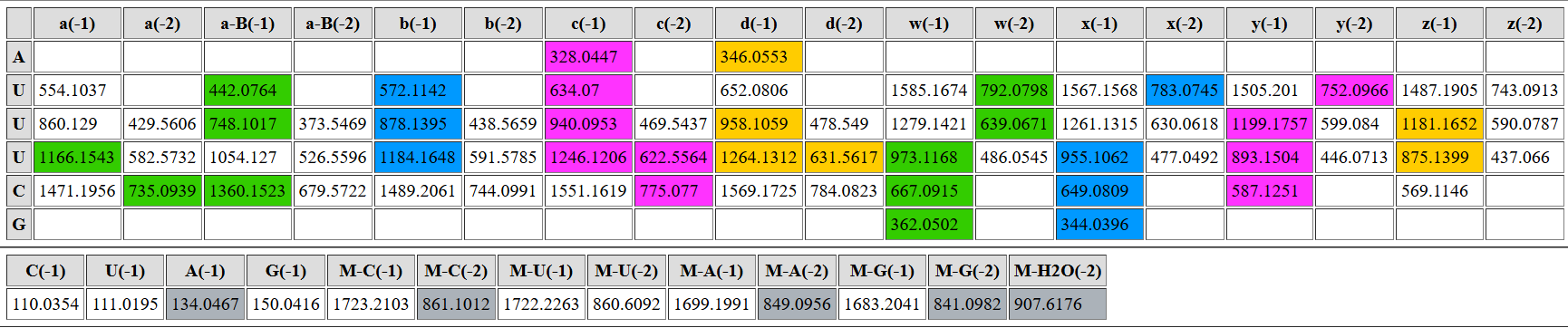

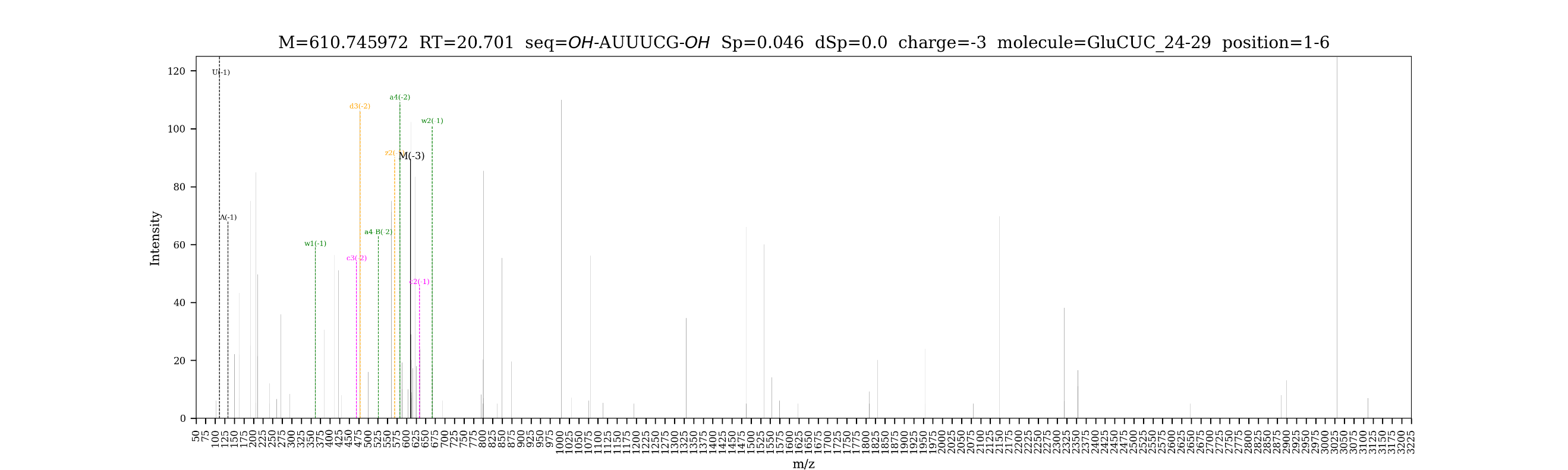

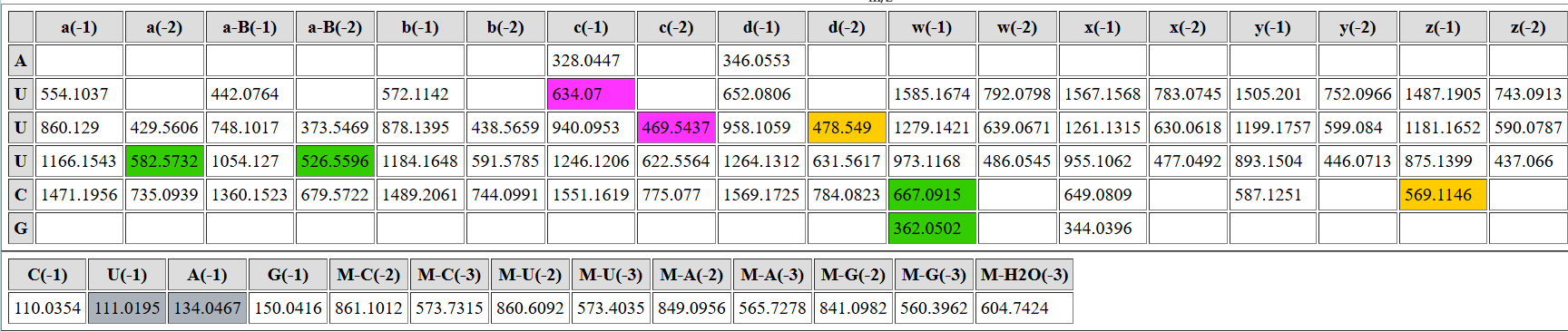

| **digested fragment information** | |
| --- | --- |
| **enzyme** | RNase T1 |
| **unmodified sequence** | CG |
| **modified sequence** | [mC]G |
| **position on *A. c* tRNA^Glu^CUC** | [30-31] |
| **5’ end** | OH |
| **3’ end** | OH |
| **charge state detected** | -1 |
| **theoretical m/z** | unmodified: 587.1251  modified: 601.1408 |
| **ppm used for EIC extraction** | 20 |
| **detection of MS1 signal in data files** | |
| 20250514_Glu_2-6_T1 | Y; N |
| 20250514_Glu_7-11-A_T1 | Y; N |
| 20250514_Glu_unk3_T1 | Y; N |
| 20250518_Glu_combined | Y; N |
| 20250524_tRNA_Glu_T1_heart | Y; N |
| 20250626_Ac#16_muscle_Glu-2_T1 | Y; N |

**EICs**

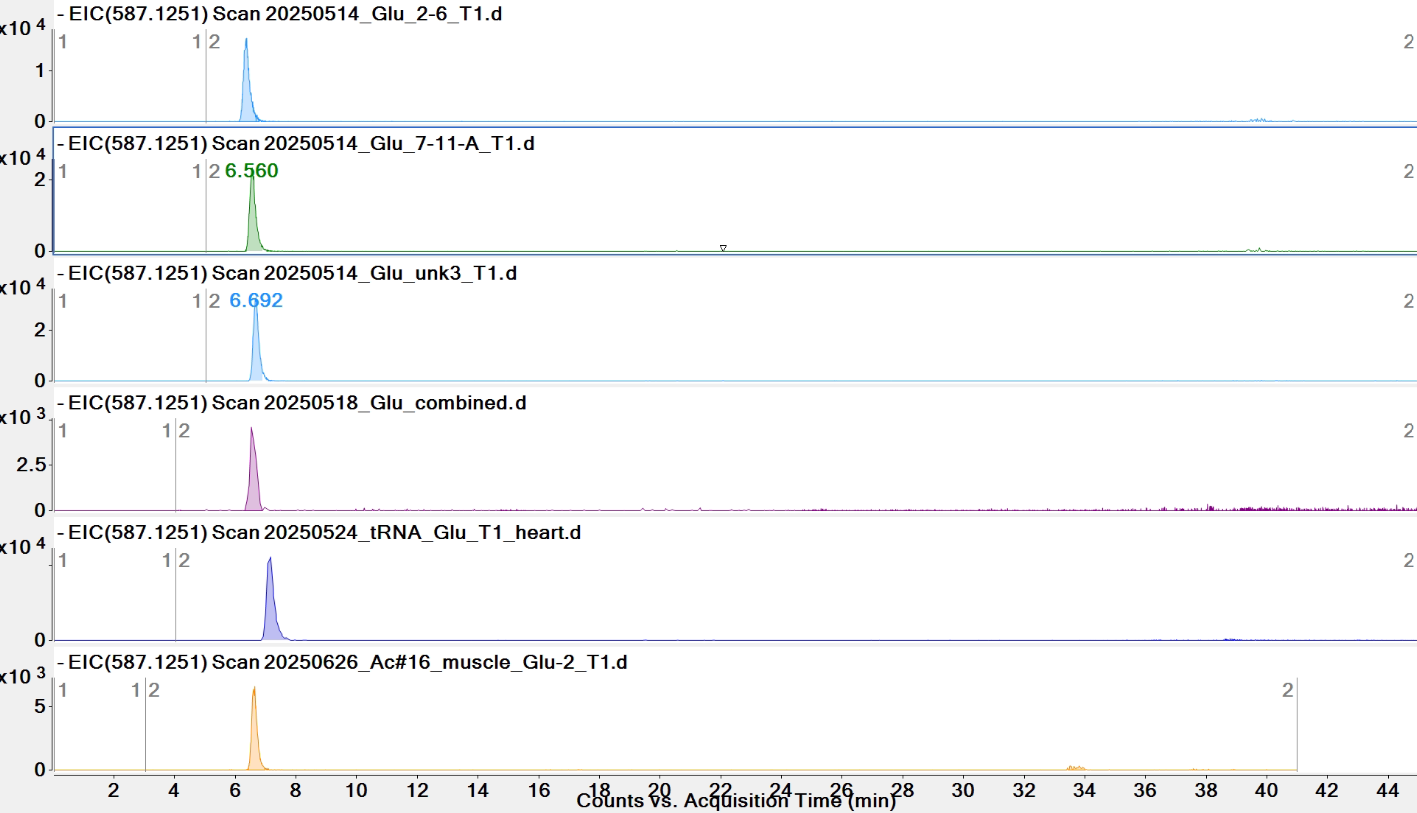

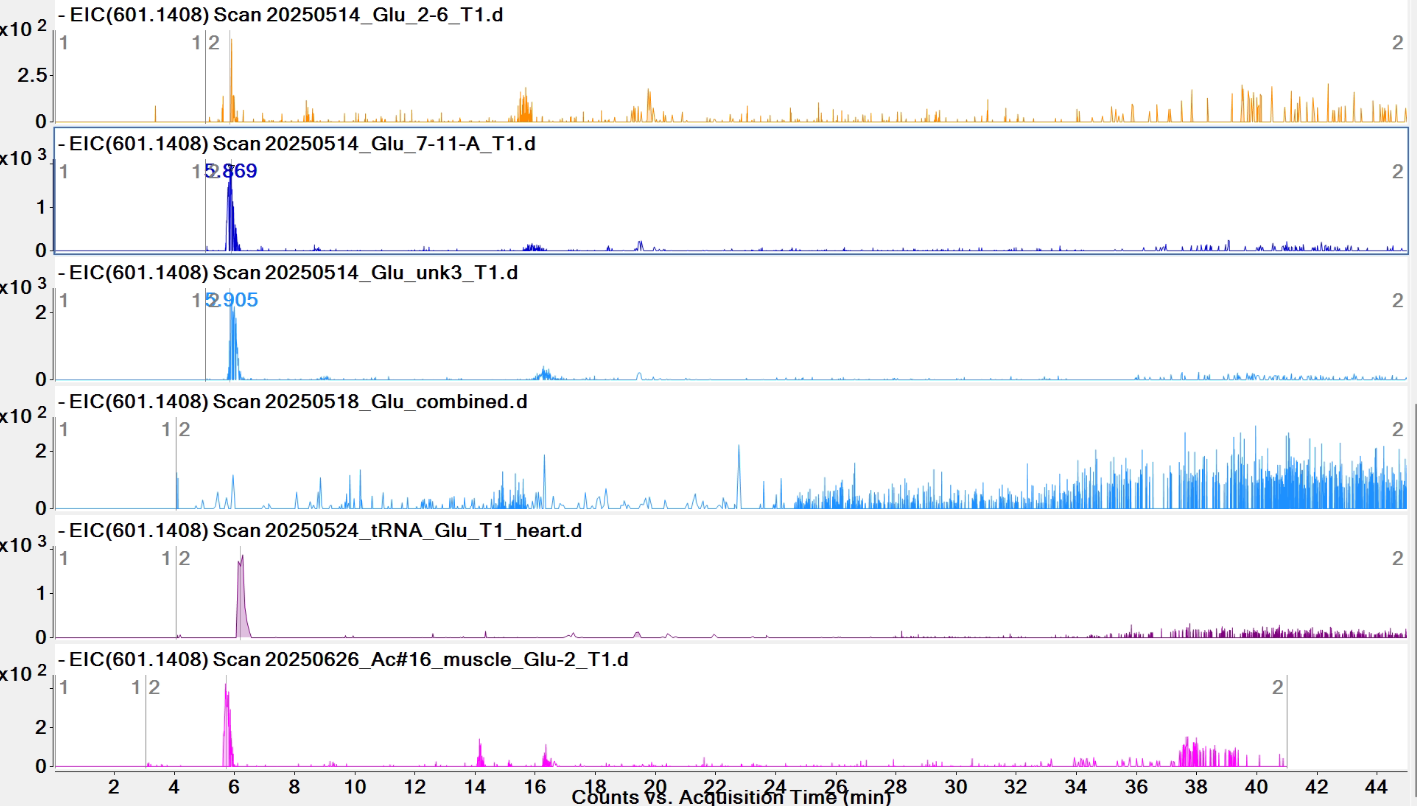

**Observed mass spectra for *m/z* = 601.1408**

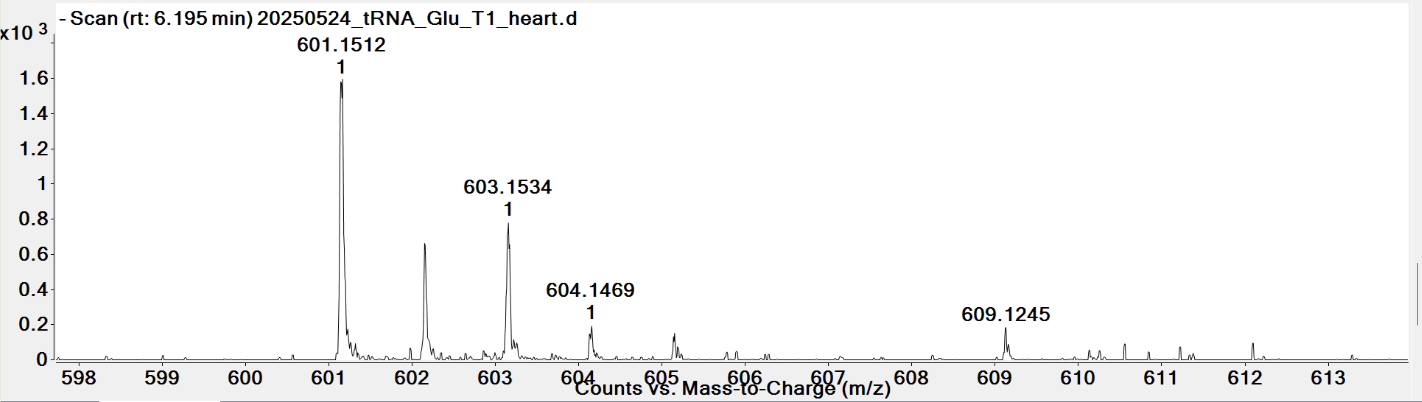

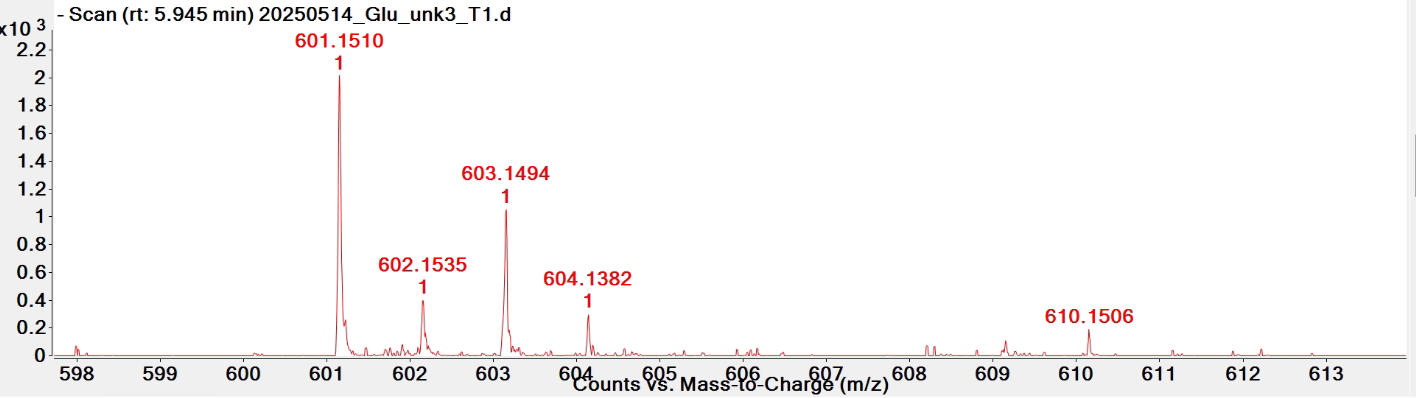

**Theoretical isotopic envelope for [mC]G (z = -1)**

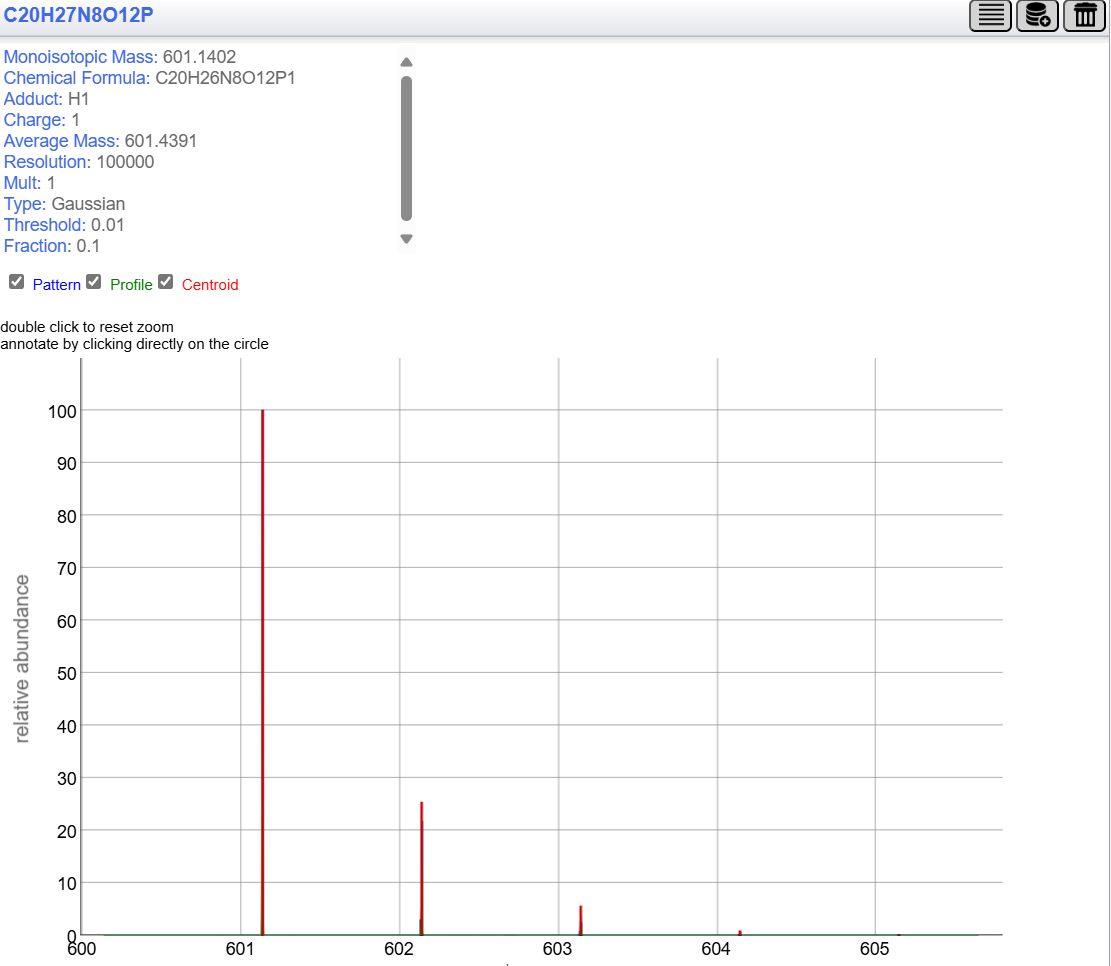

*The observed mass spectra for *m/z* = 601.1408 (z = -1) doesn’t match theoretical isotopic envelope for [mC]G (*z* = -1), so could not verify this signal was from [mC]G. Signal from [s^2^C]G was suspected, but s^2^C was not detected in nucleoside profile of tRNA^Glu^CUC.

**Pytheas results: no match**

| **digested fragment information** | |
| --- | --- |
| **enzyme** | RNase T1 |
| **unmodified sequence** | CUCUCACCG |
| **modified sequence** | [Cm]UCUCACCG |
| **position on *A. c* tRNA^Glu^CUC** | [32-40] |
| **5' end** | OH |
| **3' end** | OH |
| **charge state detected** | -3 |
| **theoretical m/z** | 920.1311 |
| **ppm used for EIC extraction** | 20 |
| **detection of MS1 signal in data files** | |
| 20250514_Glu_2-6_T1 | Y |
| 20250514_Glu_7-11-A_T1 | Y |
| 20250514_Glu_unk3_T1 | Y |
| 20250518_Glu_combined | Y |
| 20250524_tRNA_Glu_T1_heart | Y |
| 20250626_Ac#16_muscle_Glu-2_T1 | Y |

**EICs**

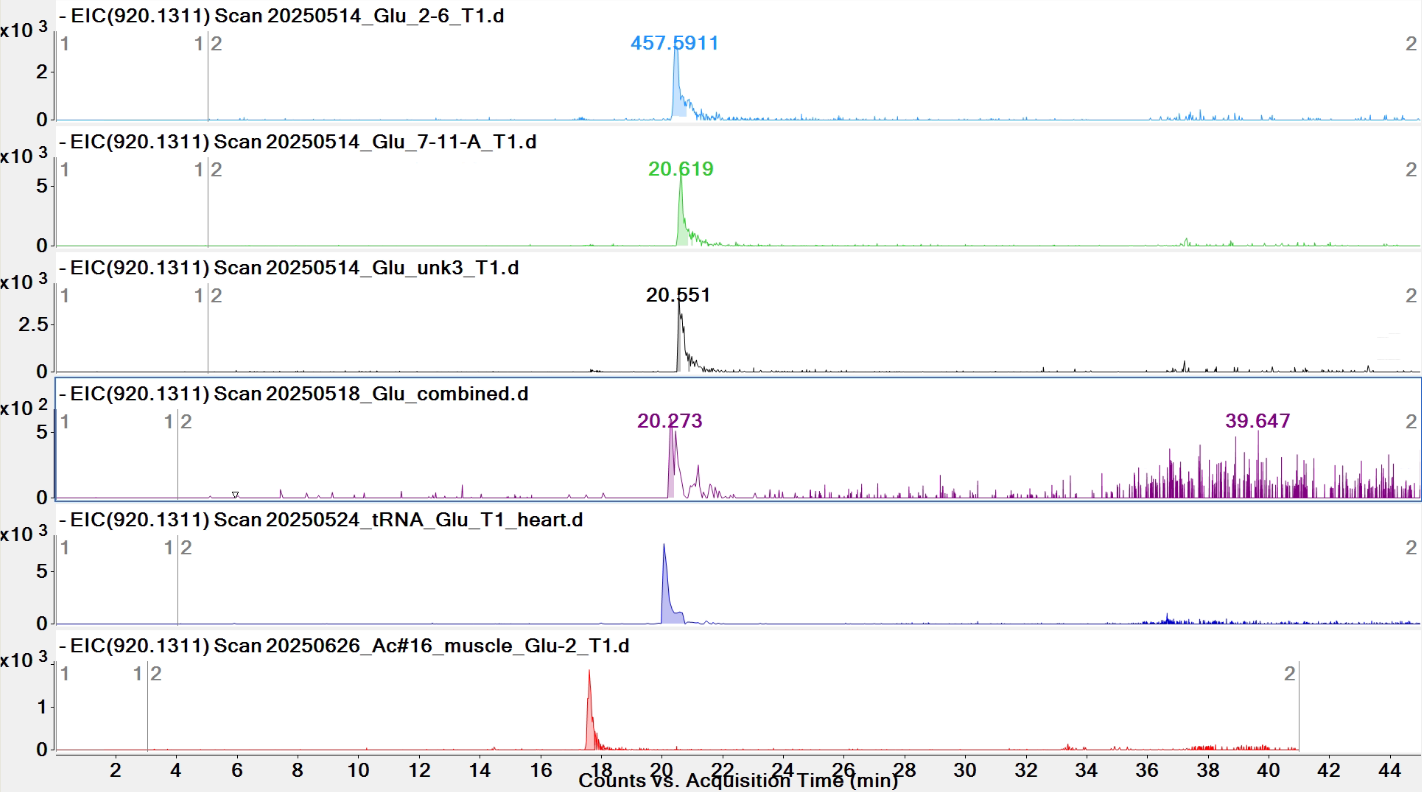

**Pytheas results:**

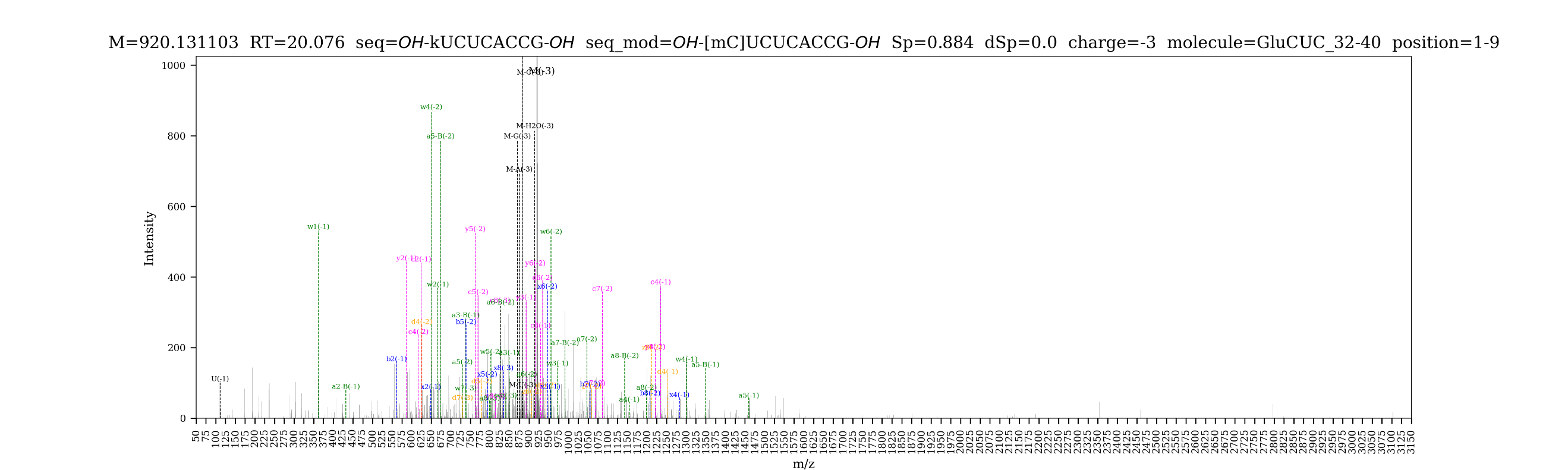

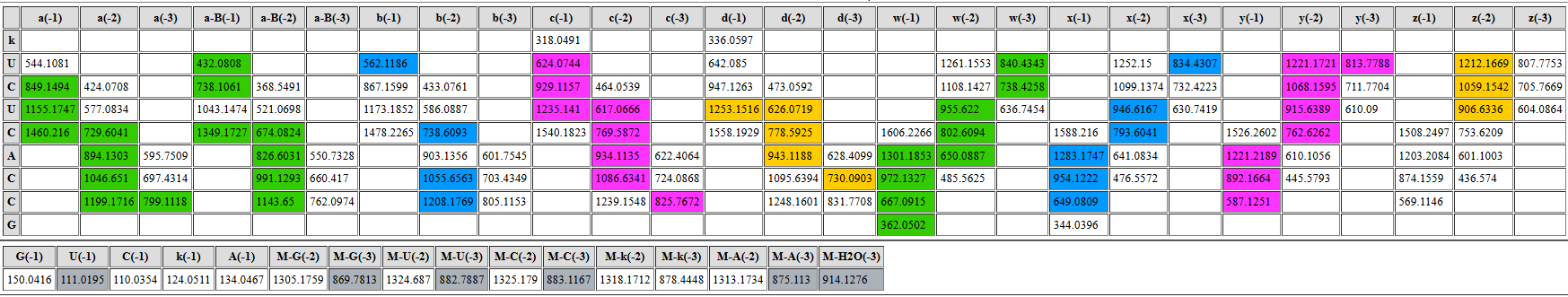

| **digested fragment information** | |
| --- | --- |
| **enzyme** | RNase T1 |
| **unmodified sequence** | CG |
| **modified sequence** | N/A |
| **position on *A. c* tRNA^Glu^CUC** | [41-42] |
| **5' end** | OH |
| **3' end** | OH |
| **charge state detected** | -1 |
| **theoretical m/z** | 587.1251 |
| **ppm used for EIC extraction** | 20 |
| **detection of MS1 signal in data files** | |
| 20250514_Glu_2-6_T1 | Y |
| 20250514_Glu_7-11-A_T1 | Y |
| 20250514_Glu_unk3_T1 | Y |
| 20250518_Glu_combined | Y |
| 20250524_tRNA_Glu_T1_heart | Y |
| 20250626_Ac#16_muscle_Glu-2_T1 | Y |

**EICs**

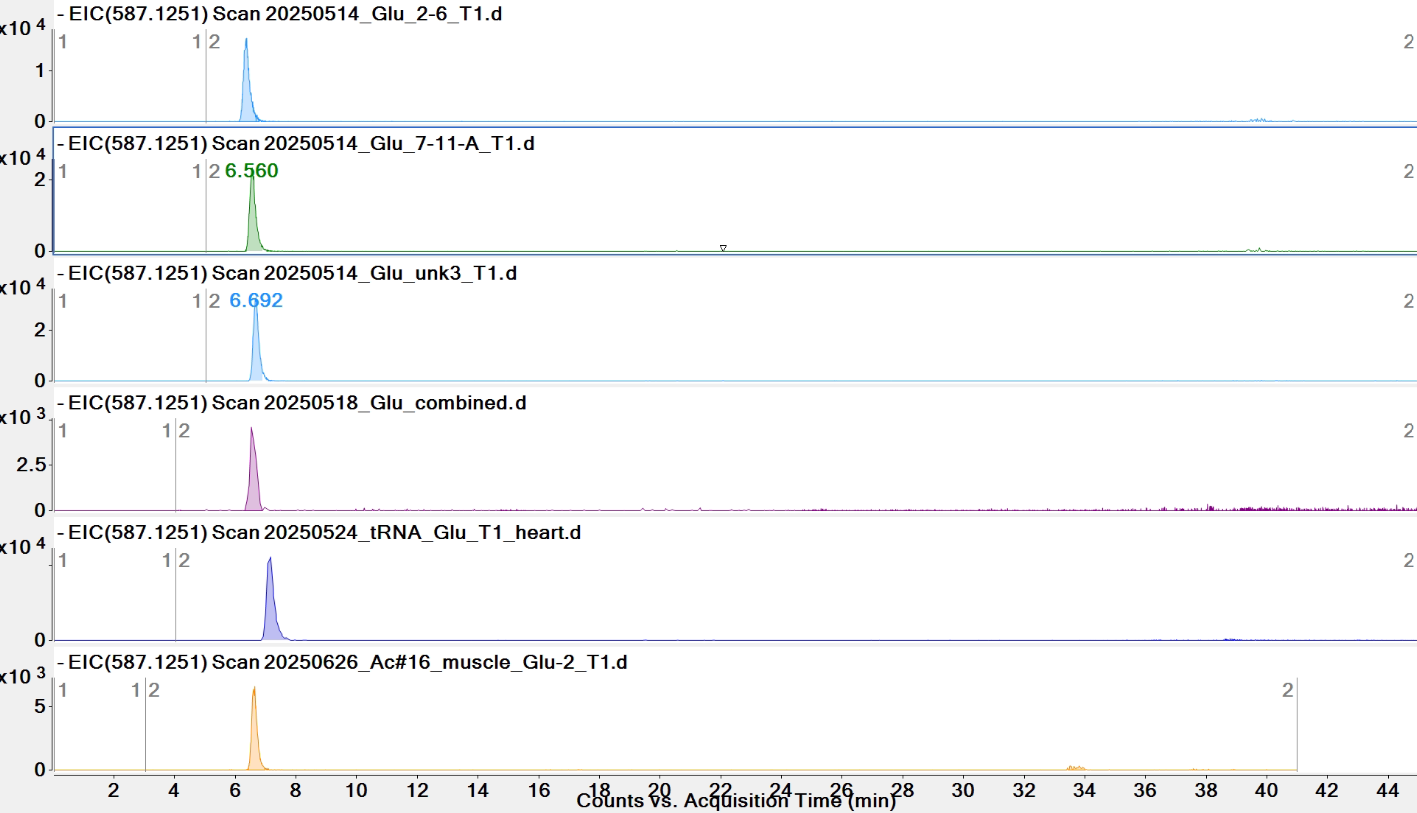

**Pytheas results: no match**

| **digested fragment information** | |
| --- | --- |
| **enzyme** | RNase T1 |
| **unmodified sequence** | ACG |
| **modified sequence** | N/A |
| **position on *A. c* tRNA^Glu^CUC** | [43-45] |
| **5' end** | OH |
| **3' end** | OH |
| **charge state detected** | -2 |
| **theoretical m/z** | 457.5849 |
| **ppm used for EIC extraction** | 20 |
| **detection of MS1 signal in data files** | |
| 20250514_Glu_2-6_T1 | Y |
| 20250514_Glu_7-11-A_T1 | Y |
| 20250514_Glu_unk3_T1 | Y |
| 20250518_Glu_combined | Y |
| 20250524_tRNA_Glu_T1_heart | Y |
| 20250626_Ac#16_muscle_Glu-2_T1 | Y |

*For all samples, 464 (with one methyl) signal is very weak and only z = -2 can be verified (z = -1 cannot be verified). So, 464 was deleted.

**EICs**

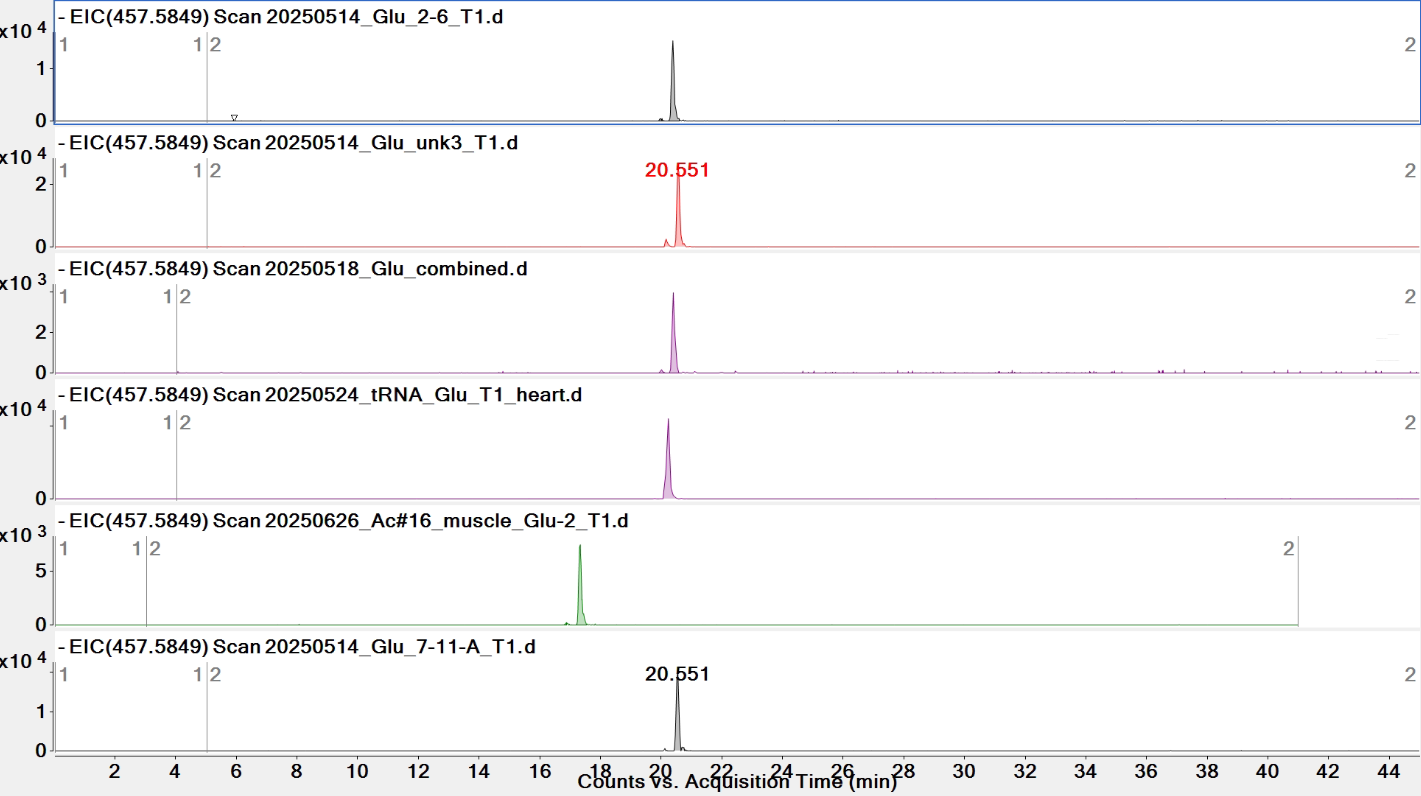

**Pytheas results**

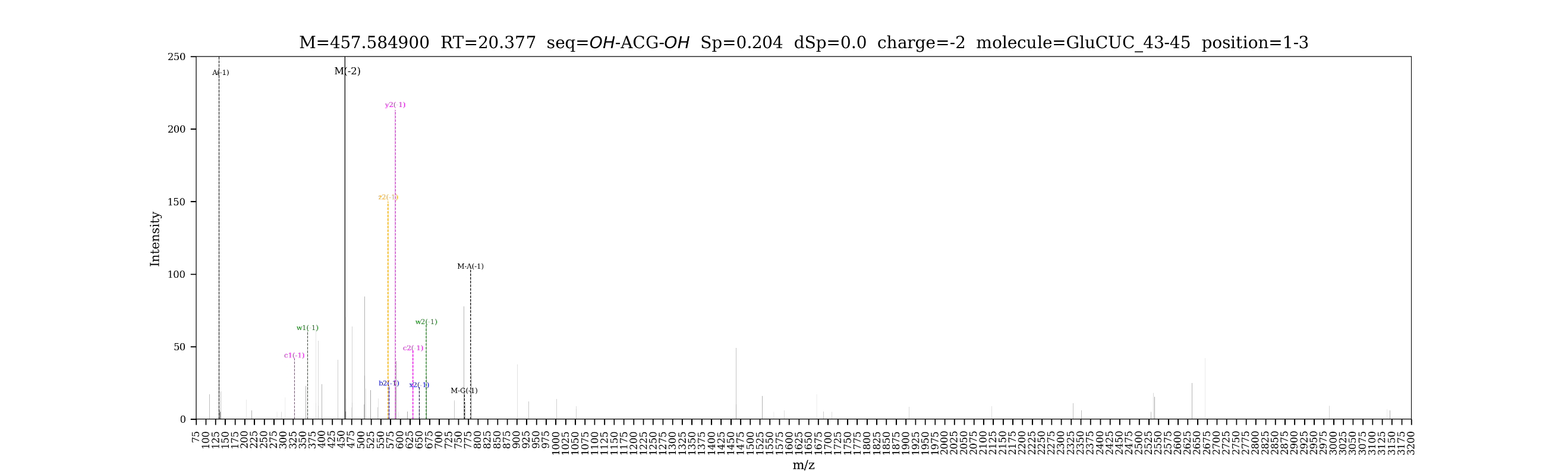

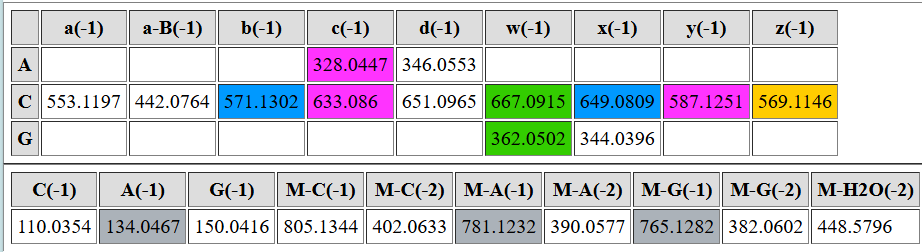

| **digested fragment information** | |
| --- | --- |
| **enzyme** | RNase T1 |
| **unmodified sequence** | CCG |
| **modified sequence** | [mC][mC]G |
| **position on *A. c* tRNA^Glu^CUC** | [47-49] |
| **5’ end** | OH |
| **3’ end** | OH |
| **charge state detected** | -2 |
| **theoretical m/z** | unmodified: 445.5793  modified: 452.5871, 459.5950 |
| **ppm used for EIC extraction** | 20 |
| **detection of MS1 signal in data files** | |
| 20250514_Glu_2-6_T1 | Y; Y, Y |
| 20250514_Glu_7-11-A_T1 | Y; Y, Y |
| 20250514_Glu_unk3_T1 | Y; Y, Y |
| 20250518_Glu_combined | Y; Y, Y |
| 20250524_tRNA_Glu_T1_heart* | Y; Y, Y |
| 20250626_Ac#16_muscle_Glu-2_T1 | Y; Y, Y |

**EICs**

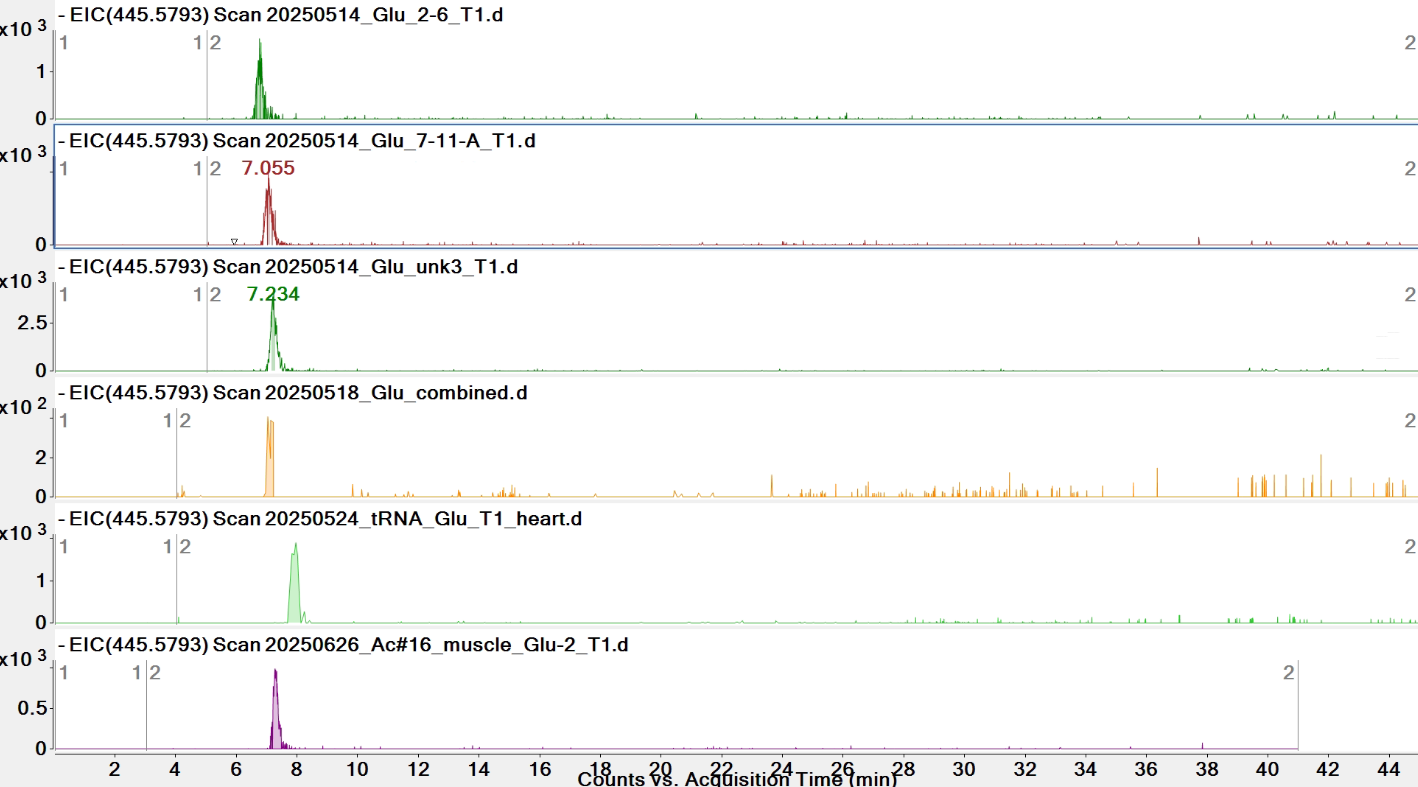

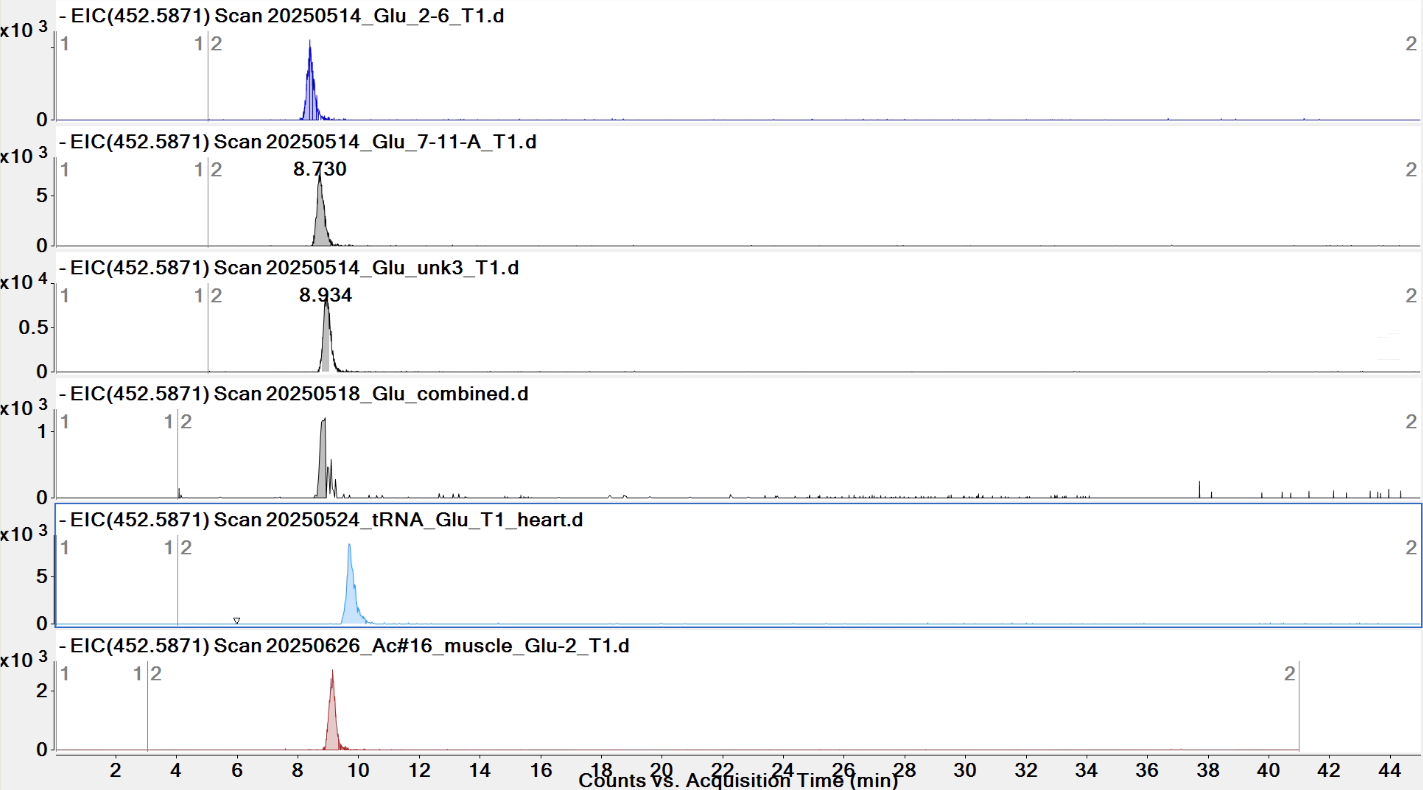

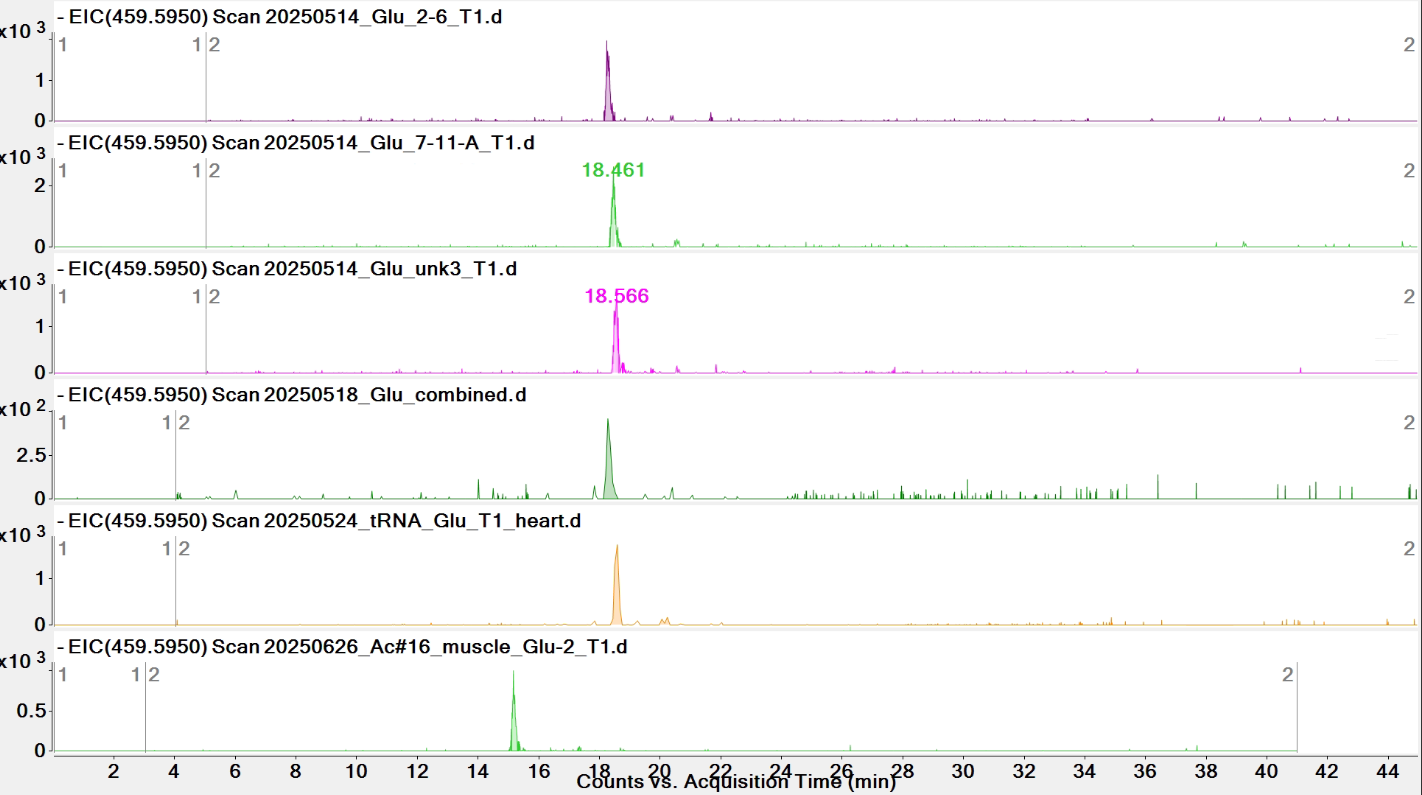

**Comparison of EICs in heart sample**
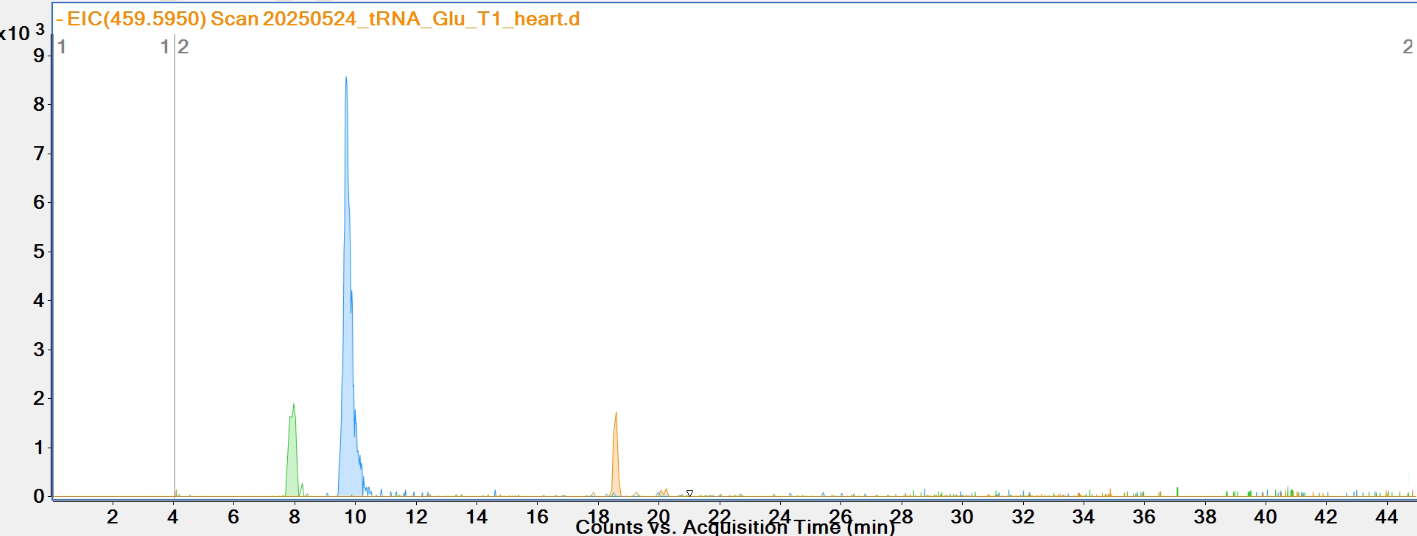

459.5950

445.5793

452.5871

**Pytheas results**

| **digested fragment information** | |
| --- | --- |
| **enzyme** | RNase T1 |
| **unmodified sequence** | UUCG |
| **modified sequence** | [m5Um]UCG |
| **position on *A. c* tRNA^Glu^CUC** | [53-56] |
| **5' end** | OH |
| **3' end** | OH |
| **charge state detected** | -2 |
| **theoretical m/z** | unmodified: 599.0840  modified: 613.0996 |
| **ppm used for EIC extraction** | 20 |
| **detection of MS1 signal in data files** | |
| 20250514_Glu_2-6_T1 | Y, Y |
| 20250514_Glu_7-11-A_T1 | Y, Y |
| 20250514_Glu_unk3_T1 | Y, Y |
| 20250518_Glu_combined | Y, Y |
| 20250524_tRNA_Glu_T1_heart | Y, Y |
| 20250626_Ac#16_muscle_Glu-2_T1 | Y, Y |

**EICs**

*The earlier peak is for DDAG (z = -2)

peak for DDAG*

599.0840

613.0996

*The peak for DDAG (z = -2) has a very close *m/z* = 613.1052 but can be distinguished by manual inspection of spectra.

**Pytheas results**

| **digested fragment information** | |
| --- | --- |
| **enzyme** | RNase T1 |
| **unmodified sequence** | AUUCCCCG |
| **modified sequence** | [mA]UUCCCCG |
| **position on *A. c* tRNA^Glu^CUC** | [57-64] |
| **5’ end** | OH |
| **3’ end** | OH |
| **charge state detected** | -3, -2 |
| **theoretical m/z** | unmodified: 813.7788  modified: 818.4507, 1228.1800 |
| **ppm used for EIC extraction** | 20 |
| **detection of MS1 signal in data files** | |
| 20250514_Glu_2-6_T1 | Y; Y, Y |
| 20250514_Glu_7-11-A_T1 | Y; Y, Y |
| 20250514_Glu_unk3_T1 | Y; Y, Y |
| 20250518_Glu_combined | Y; Y, Y |
| 20250524_tRNA_Glu_T1_heart | Y; Y, Y |
| 20250626_Ac#16_muscle_Glu-2_T1 | Y; Y, Y |

**EICs**

*For the second peak of 813, UCUCACCG (#33-40) was suspected, but RNase T1 should not cut after Cm, and tiRNA^Glu^CUC of 43 nt was not detected.

**A comparison of EICs for modified and unmodified variants in heart sample**

813.7788

818.4507

**Pytheas results**

| **digested fragment information** | |
| --- | --- |
| **enzyme** | RNase T1 |
| **unmodified sequence** | CUUG |
| **modified sequence** | N/A |
| **position on *A. c* tRNA^Glu^CUC** | [65-68] |
| **5' end** | OH |
| **3' end** | OH |
| **charge state detected** | -2 |
| **theoretical m/z** | 599.0840 |
| **ppm used for EIC extraction** | 20 |
| **detection of MS1 signal in data files** | |
| 20250514_Glu_2-6_T1 | Y |
| 20250514_Glu_7-11-A_T1 | Y |
| 20250514_Glu_unk3_T1 | Y |
| 20250518_Glu_combined | Y |
| 20250524_tRNA_Glu_T1_heart | Y |
| 20250626_Ac#16_muscle_Glu-2_T1 | Y |

**EICs**

**Pytheas results**

| **digested fragment information** | |
| --- | --- |
| **enzyme** | RNase T1 |
| **unmodified sequence** | AACCA |
| **modified sequence** | N/A |
| **position on *A. c* tRNA^Glu^CUC** | [71-75] |
| **5' end** | OH |
| **3' end** | OH |
| **charge state detected** | -2 |
| **theoretical m/z** | 766.6344 |
| **ppm used for EIC extraction** | 20 |
| **detection of MS1 signal in data files** | |
| 20250514_Glu_2-6_T1 | Y |
| 20250514_Glu_7-11-A_T1 | Y |
| 20250514_Glu_unk3_T1 | Y |
| 20250518_Glu_combined | Y |
| 20250524_tRNA_Glu_T1_heart | Y |
| 20250626_Ac#16_muscle_Glu-2_T1 | Y |

**EICs**

**Pytheas results**

| **digested fragment information** | |
| --- | --- |
| **enzyme** | RNase T1 |
| **unmodified sequence** | AACC |
| **modified sequence** | N/A |
| **position on *A. c* tRNA^Glu^CUC** | [71-74] |
| **5' end** | OH |
| **3' end** | OH |
| **charge state detected** | -2 |
| **theoretical m/z** | 602.1081 |
| **ppm used for EIC extraction** | 20 |
| **detection of MS1 signal in data files** | |
| 20250514_Glu_2-6_T1 | N |
| 20250514_Glu_7-11-A_T1 | Y |
| 20250514_Glu_unk3_T1 | Y |
| 20250518_Glu_combined | N |
| 20250524_tRNA_Glu_T1_heart | Y |
| 20250626_Ac#16_muscle_Glu-2_T1 | Y |

**EICs**

**Pytheas results**

| **digested fragment information** | |
| --- | --- |
| **enzyme** | RNase T1 |
| **unmodified sequence** | AAC |
| **modified sequence** | N/A |
| **position on *A. c* tRNA^Glu^CUC** | [71-73] |
| **5' end** | OH |
| **3' end** | OH |
| **charge state detected** | -1 |
| **theoretical m/z** | 900.1827 |
| **ppm used for EIC extraction** | 20 |
| **detection of MS1 signal in data files** | |
| 20250514_Glu_2-6_T1 | N |
| 20250514_Glu_7-11-A_T1 | N |
| 20250514_Glu_unk3_T1 | Y |
| 20250518_Glu_combined | N |
| 20250524_tRNA_Glu_T1_heart | N |
| 20250626_Ac#16_muscle_Glu-2_T1 | N |

**EICs**

**Pytheas results:** no match

| **digested fragment information** | |
| --- | --- |
| **enzyme** | RNase T1 |
| **unmodified sequence** | AA |
| **modified sequence** | N/A |
| **position on *A. c* tRNA^Glu^CUC** | [71-72] |
| **5' end** | OH |
| **3' end** | OH |
| **charge state detected** | -1 |
| **theoretical m/z** | 595.1415 |
| **ppm used for EIC extraction** | 20 |
| **detection of MS1 signal in data files** | |
| 20250514_Glu_2-6_T1 | Y |
| 20250514_Glu_7-11-A_T1 | Y |
| 20250514_Glu_unk3_T1 | Y |
| 20250518_Glu_combined | Y |
| 20250524_tRNA_Glu_T1_heart | Y |
| 20250626_Ac#16_muscle_Glu-2_T1 | N |

EICs

**Pytheas results:** no MS/MS spectra (2-nt fragment is too short)

**Part 3. RNase T1 digested fragments from tiRNA^Glu^CUC**

**Note:**

Theoretically, these fragments are only produced by T1 digest on tiRNA^Glu^CUC but not intact tRNA^Glu^CUC

| **digested fragment information** | |
| --- | --- |
| **enzyme** | RNase T1 |
| **unmodified sequence** | ACCG |
| **modified sequence** | N/A |
| **position on *A. c* tRNA^Glu^CUC** | [37-40] |
| **5' end** | OH |
| **3' end** | OH |
| **charge state detected** | -2 |
| **theoretical m/z** | 610.1056 |
| **ppm used for EIC extraction** | 20 |
| **detection of MS1 signal in data files** | |
| 20250514_Glu_2-6_T1 | Y |
| 20250514_Glu_7-11-A_T1 | Y |
| 20250514_Glu_unk3_T1 | Y |
| 20250518_Glu_combined | Y |
| 20250524_tRNA_Glu_T1_heart | Y |
| 20250626_Ac#16_muscle_Glu-2_T1 | Y |

**EICs**

**Pytheas results** (from 20250514_Glu_unk3_T1)

| **digested fragment information** | |
| --- | --- |
| **enzyme** | RNase T1 |
| **unmodified sequence** | CACCG |
| **modified sequence** | N/A |
| **position on *A. c* tRNA^Glu^CUC** | [36-40] |
| **5’ end** | OH |
| **3’ end** | OH |
| **charge state detected** | -2 |
| **theoretical m/z** | 762.6262 |
| **ppm used for EIC extraction** | 20 |
| **detection of MS1 signal in data files** | |
| 20250514_Glu_2-6_T1 | Y |
| 20250514_Glu_7-11-A_T1 | Y |
| 20250514_Glu_unk3_T1 | Y |
| 20250518_Glu_combined | Y |
| 20250524_tRNA_Glu_T1_heart | Y |
| 20250626_Ac#16_muscle_Glu-2_T1 | Y |

**EICs**

**Pytheas results**

| **digested fragment information** | |
| --- | --- |
| **enzyme** | RNase T1 |
| **unmodified sequence** | UCACCG |
| **modified sequence** | N/A |
| **position on *A. c* tRNA^Glu^CUC** | [35-40] |
| **5' end** | OH |
| **3' end** | OH |
| **charge state detected** | -2 |
| **theoretical m/z** | 915.6389 |
| **ppm used for EIC extraction** | 20 |
| **detection of MS1 signal in data files** | |
| 20250514_Glu_2-6_T1 | Y |
| 20250514_Glu_7-11-A_T1 | Y |
| 20250514_Glu_unk3_T1 | Y |
| 20250518_Glu_combined | N |
| 20250524_tRNA_Glu_T1_heart | Y |
| 20250626_Ac#16_muscle_Glu-2_T1 | Y |

**EICs**

**Pytheas results**

| **digested fragment information** | |
| --- | --- |
| **enzyme** | RNase T1 |
| **unmodified sequence** | CUCACCG |
| **modified sequence** | N/A |
| **position on *A. c* tRNA^Glu^CUC** | [34-40] |
| **5’ end** | OH |
| **3’ end** | OH |
| **charge state detected** | -2, -3 |
| **theoretical m/z** | 1068.1595, 711.7704 |
| **ppm used for EIC extraction** | 20 |
| **detection of MS1 signal in data files** | |
| 20250514_Glu_2-6_T1 | Y, Y |
| 20250514_Glu_7-11-A_T1 | Y, Y |
| 20250514_Glu_unk3_T1 | Y, Y |
| 20250518_Glu_combined | N, N |
| 20250524_tRNA_Glu_T1_heart | Y, Y |
| 20250626_Ac#16_muscle_Glu-2_T1 | N, N |

**EICs**

**Pytheas results**
