## Supplementary material for "Sub-stoichiometric modifications of *Aplysia californica* tRNAs and tRNA fragments revealed by integrating intact and bottom-up mass spectrometry": Supplementary_material_3_tRNA_LysUUU.docx

**Supplementary material 3: tRNA^Lys^UUU information**

**Sequence information:**

Organism: *Aplysia californica*

RNA type: tRNA^Lys^UUU

RNAcentral link: <https://rnacentral.org/rna/URS0000947ED9/6500>

RNA sequence (unmodified, with CCA tail, 5'-3'): GCCCGGUUAGCUCAGUCGGUAGAGCAUCAGACUUUUAAUCUGAGGGUCAAGGGUUCGAGUCCCUUAUUGGGCG-CCA

Length: 76

| **List of ribonucleosides detected in *A. californica* tRNA^Lys^UUU digest** | | | | | | |
| --- | --- | --- | --- | --- | --- | --- |
| **RNAMods code** | **short name*** | **full name** | **theoretical [M+H]^+^** | **observed [M+H]^+^** | **mass error (ppm)** | **product ion(s) detected** |
| P | Y | pseudouridine | 245.0773 | 245.0732 | 16.73 | 155, 179 |
| D | D | dihydrouridine | 247.0930 | 247.0883 | 19.02 | 115 |
| C | C | cytidine | 244.0933 | 244.0896 | 15.16 | 112 |
| U | U | uridine | 245.0773 | 245.0735 | 15.51 | 113 |
| % | s^2^C | 2-thiocytidine | 260.0705 | 260.0662 | 16.53 | 128 |
| " | m^1^A | 1-methyladenosine | 282.1202 | 282.1152 | 17.72 | 150 |
| ? | m^5^C | 5-methylcytidine | 258.1090 | 258.1042 | 18.60 | 126 |
| 7 | m^7^G | 7-methylguanosine | 298.1151 | 298.1103 | 16.10 | 166 |
| G | G | guanosine | 284.0995 | 284.0949 | 16.19 | 152 |
| L | m^2^G | N2-methylguanosine | 298.1151 | 298.1097 | 18.11 | 166 |
| A | A | adenosine | 268.1046 | 268.0997 | 18.28 | 136 |
| \ | m^5^Um | 5,2'-O-dimethyluridine | 273.1086 | 273.1037 | 17.94 | 127 |
| 3 | mcm^5^s^2^U | 5-methoxycarbonylmethyl-2-thiouridine | 333.0756 | 333.0705 | 15.31 | 141, 169, 201 |
| = | m^6^A | N6-methyladenosine | 282.1202 | 282.1148 | 19.14 | 150 |
| [ | ms^2^t^6^A | 2-methylthio-N6-threonylcarbamoyladenosine | 459.1298 | 459.1214 | 18.30 | 327 |

*mass error was calculated using the observed [M+H]^+^ value at the peak maximum: mass error = (observed [M+H]^+^ - theoretical[M+H]^+^)/ theoretical[M+H]^+^ × 10^6^. The dataset used was: 20250418_Ac#2muscletRNALys (muscle).

**Overlaid EICs for ribonucleosides detected in *A. californica* tRNA^Lys^UUU digest**

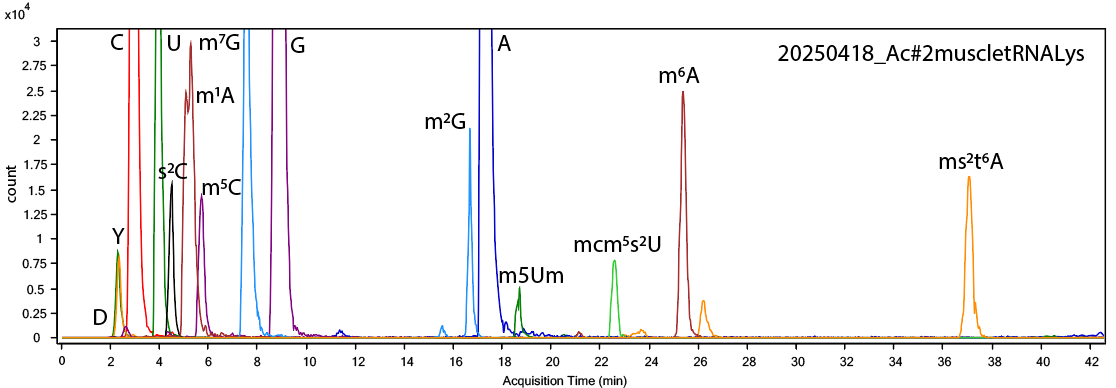

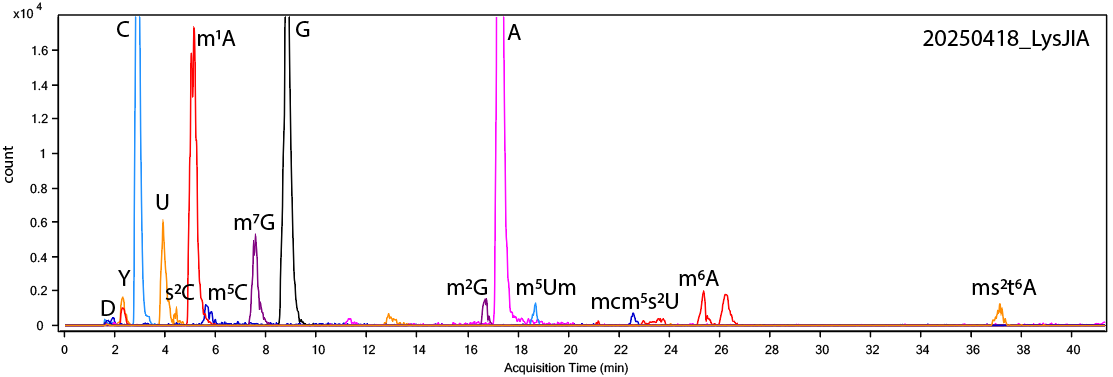

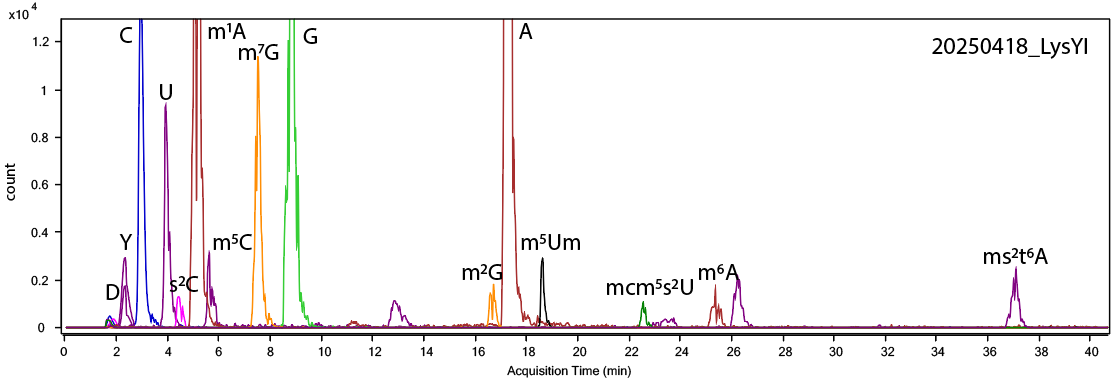

**Part 1. RNase 4 data**

| **list of LC-MS/MS data files used for analysis by RNase 4 digestion** | |
| --- | --- |
| **data file name** | **tissue** |
| 20250216_Lys-1+2-R4 | male reproductive organ pooled from multiple slugs |
| 20250216_Lys-JIA+YI-R4 | male reproductive organ pooled from multiple slugs |
| 20250227_Lys1+2_R4* | male reproductive organ pooled from multiple slugs |
| *Pytheas results are based on this data file unless otherwise indicated. | |

| **digested fragment information** | |
| --- | --- |
| **enzyme** | RNase 4 |
| **unmodified sequence** | GCCCGGUU |
| **modified sequence** | N/A |
| **position on *A. c* tRNA Lys UUU** | [1-8] |
| **5’ end** | P |
| **3’ end** | OH |
| **charge state detected** | -3 |
| **theoretical m/z** | 859.1213 |
| **ppm used for EIC extraction** | 20 |
| **detection of MS1 signal in data files** | |
| 20250216_Lys-1+2-R4 | Y |
| 20250216_Lys-JIA+YI-R4 | N* |
| 20250227_Lys1+2_R4 | Y* |

*Y = Yes, MS1 signal detected; N = No, MS1 signal not detected

**Corresponding MS1 EIC**

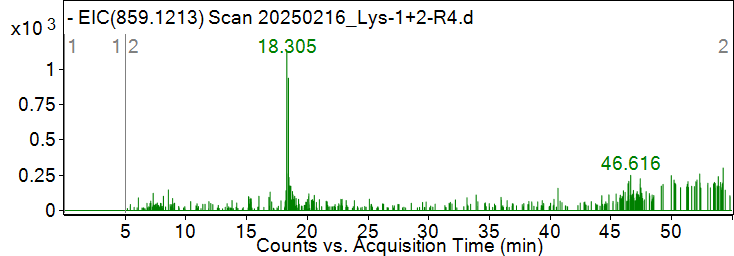

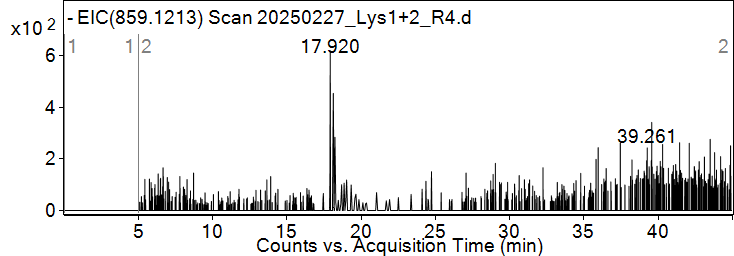

**Pytheas results**

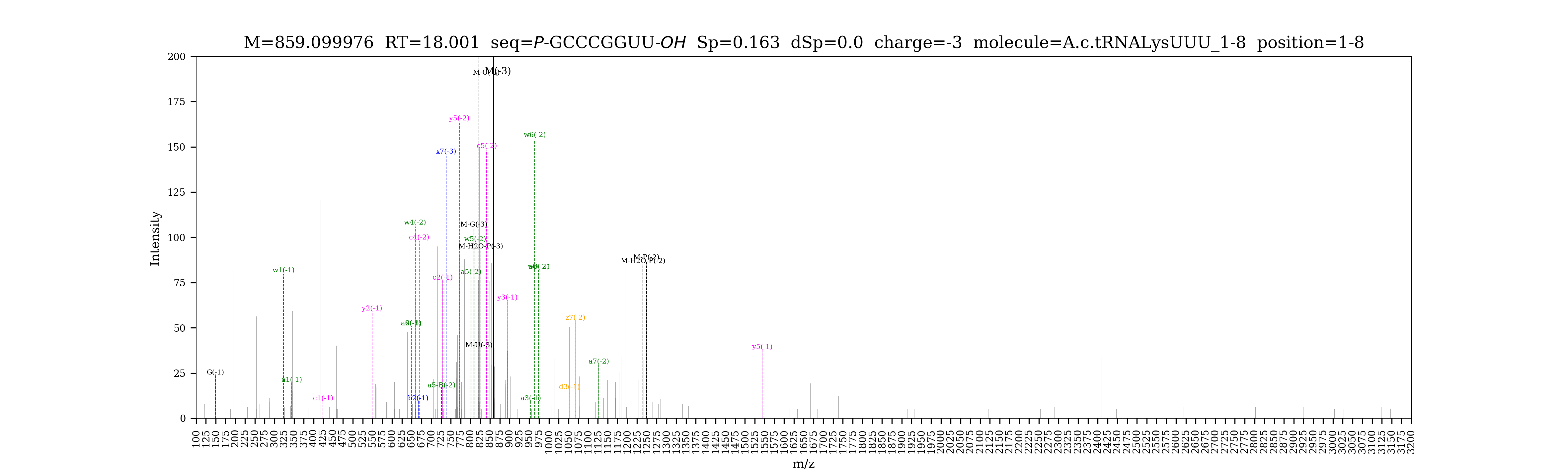

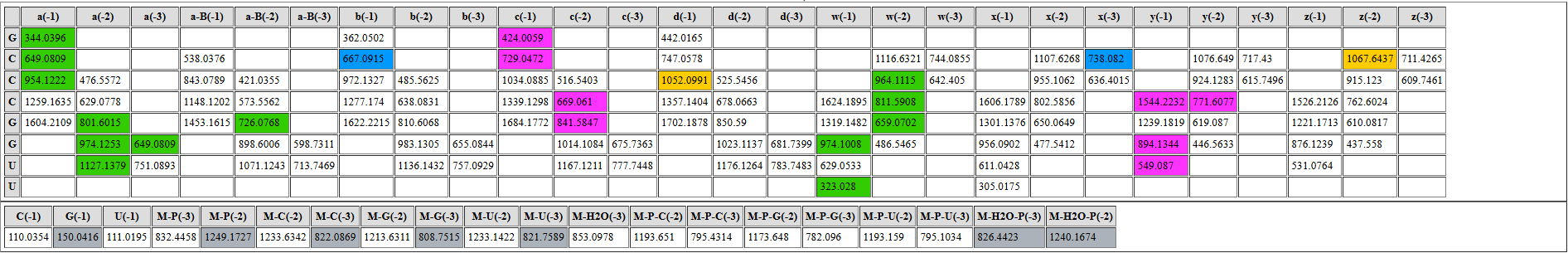

| **digested fragment information** | |
| --- | --- |
| **enzyme** | RNase 4 |
| **unmodified sequence** | AGCU |
| **modified sequence** | A[m2G]CU |
| **position on *A. c* tRNA^Lys^UUU** | [9-12] |
| **5’ end** | OH |
| **3’ end** | OH |
| **charge state detected** | -2 |
| **theoretical m/z** | 617.6054 |
| **ppm used for EIC extraction** | 20 |
| **detection of MS1 signal in data files** | |
| 20250216_Lys-1+2-R4 | Y |
| 20250216_Lys-JIA+YI-R4 | Y |
| 20250227_Lys1+2_R4 | Y |

**Corresponding MS1 EIC**

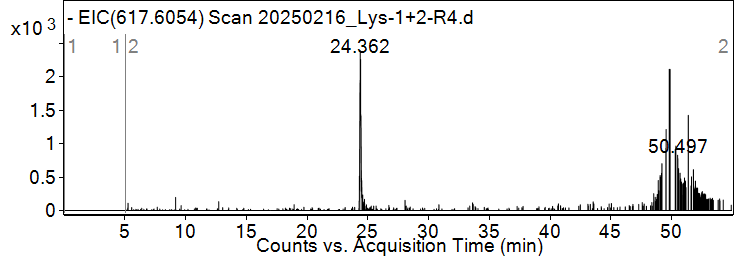

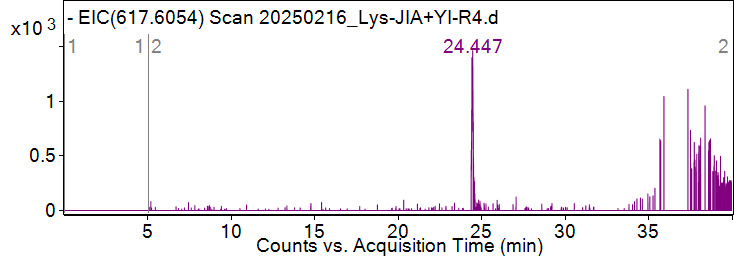

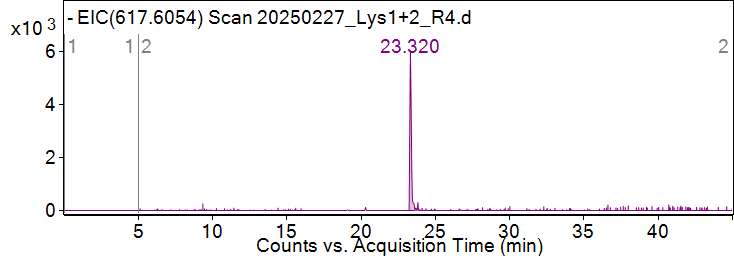
**Pytheas results**

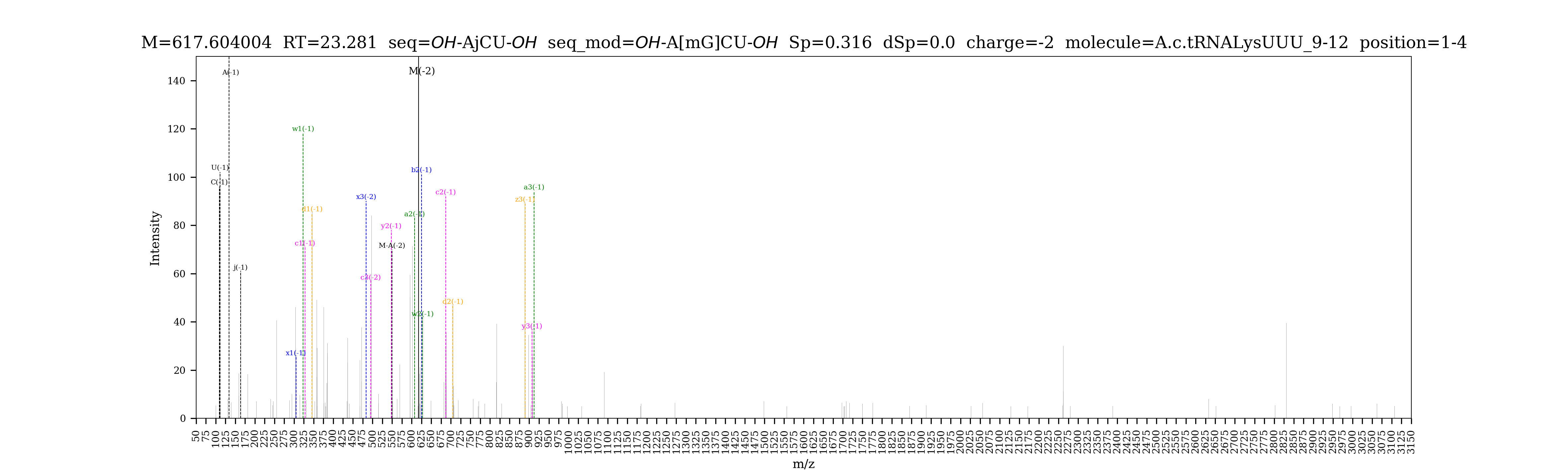

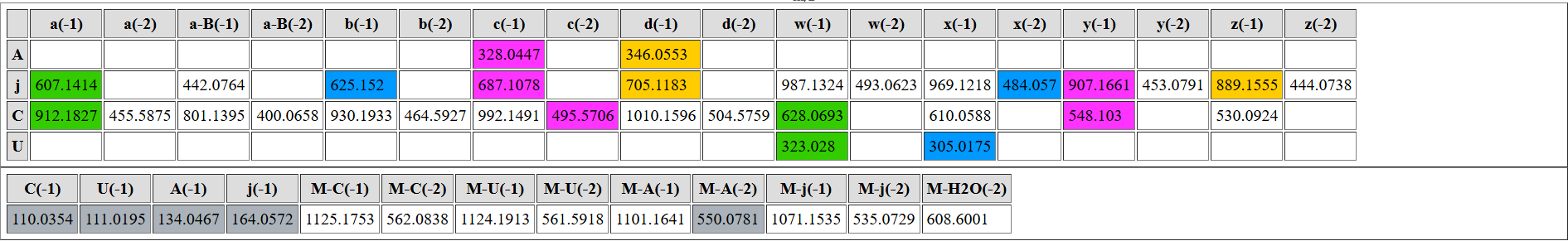

| **digested fragment information** | |
| --- | --- |
| **enzyme** | RNase 4 |
| **unmodified sequence** | CAGU |
| **modified sequence** | CAGD |
| **position on *A. c* tRNA^Lys^UUU** | [13-16] |
| **5’ end** | OH |
| **3’ end** | OH |
| **charge state detected** | -2 |
| **theoretical m/z** | modified: 611.6054 |
| **ppm used for EIC extraction** | 20 |
| **detection of MS1 signal in data files** | |
| 20250216_Lys-1+2-R4 | Y |
| 20250216_Lys-JIA+YI-R4 | N |
| 20250227_Lys1+2_R4 | Y |

**Corresponding MS1 EICs**

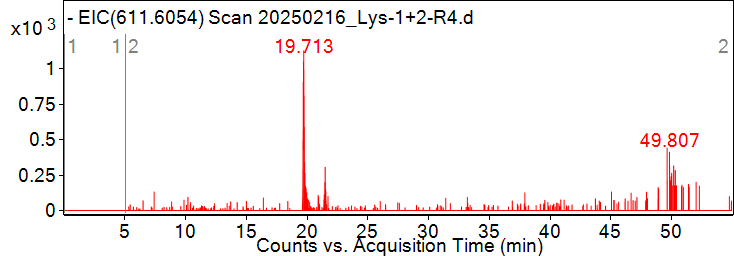

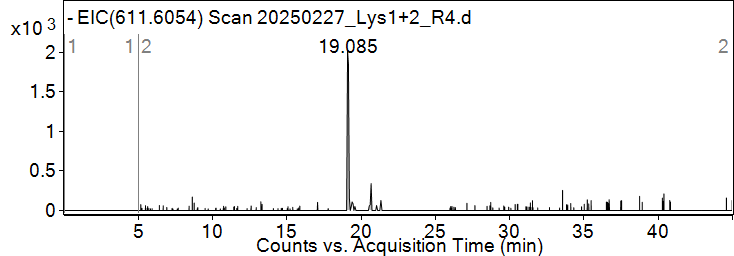

**Pytheas results**

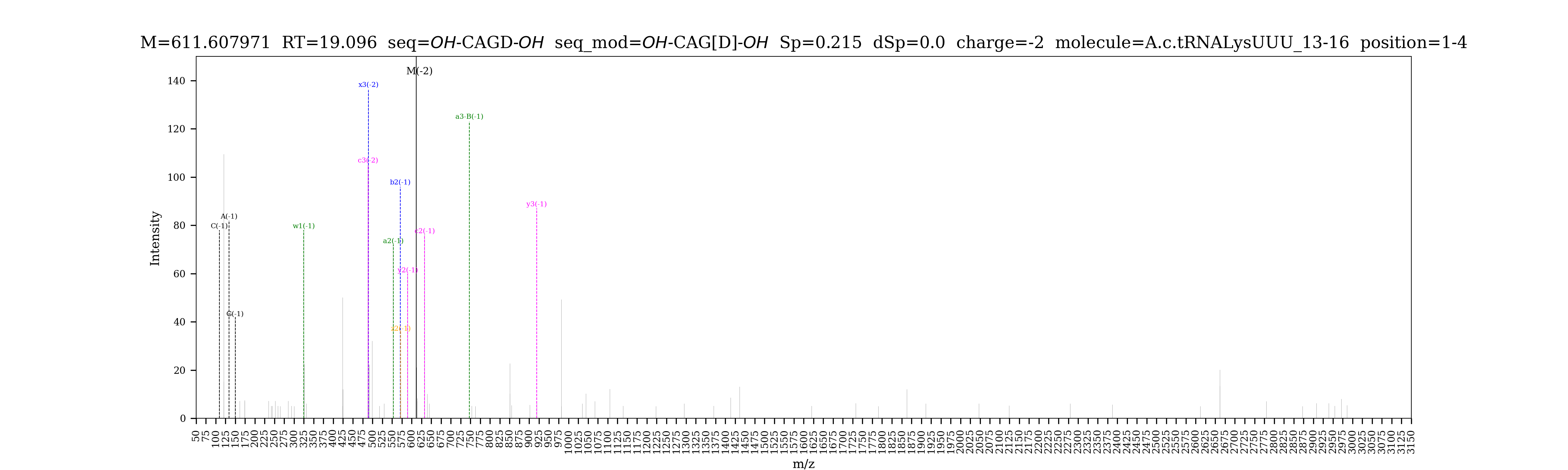

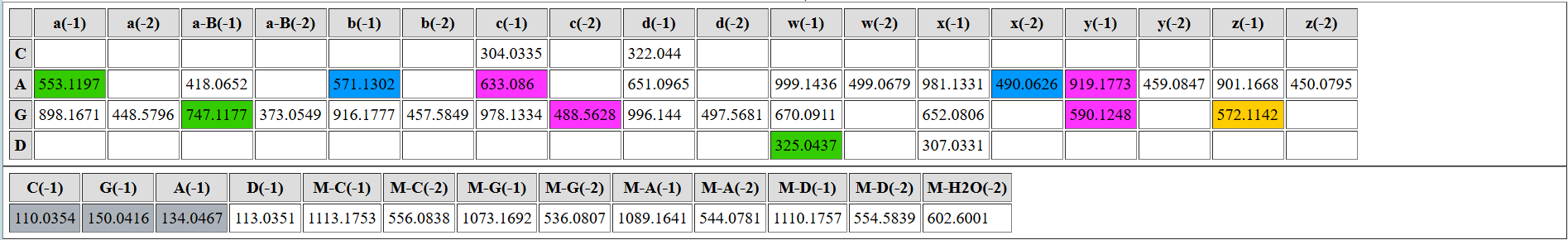

| **digested fragment information** | |
| --- | --- |
| **enzyme** | RNase 4 |
| **unmodified sequence** | CGGU |
| **modified sequence** | N/A |
| **position on *A. c* tRNA^Lys^UUU** | [17-20] |
| **5’ end** | OH |
| **3’ end** | OH |
| **charge state detected** | -2 |
| **theoretical m/z** | 618.5950 |
| **ppm used for EIC extraction** | 20 |
| **detection of MS1 signal in data files** | |
| 20250216_Lys-1+2-R4 | Y |
| 20250216_Lys-JIA+YI-R4 | Y |
| 20250227_Lys1+2_R4 | Y |

**Corresponding MS1 EIC**

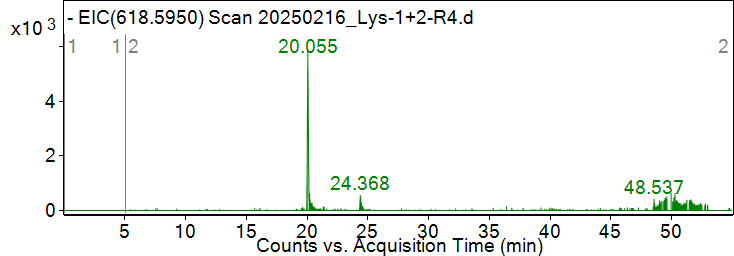

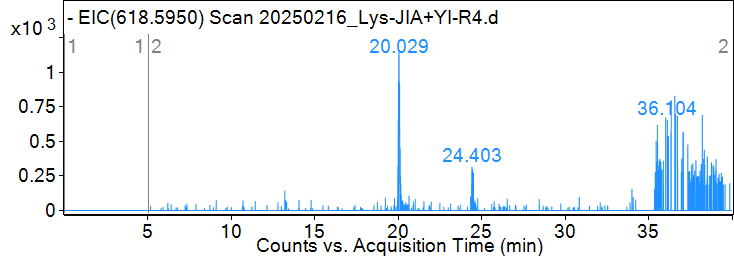

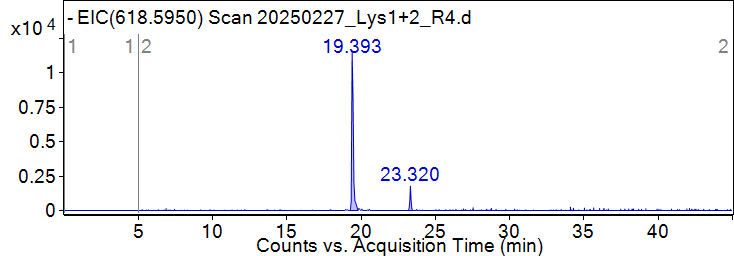
**Pytheas results**

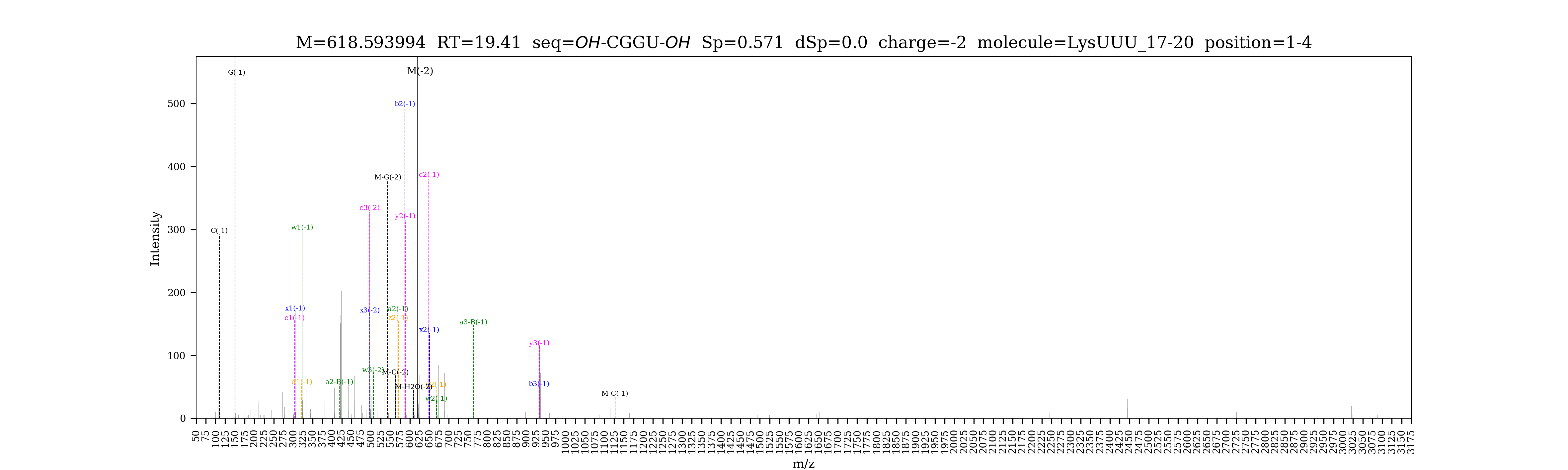

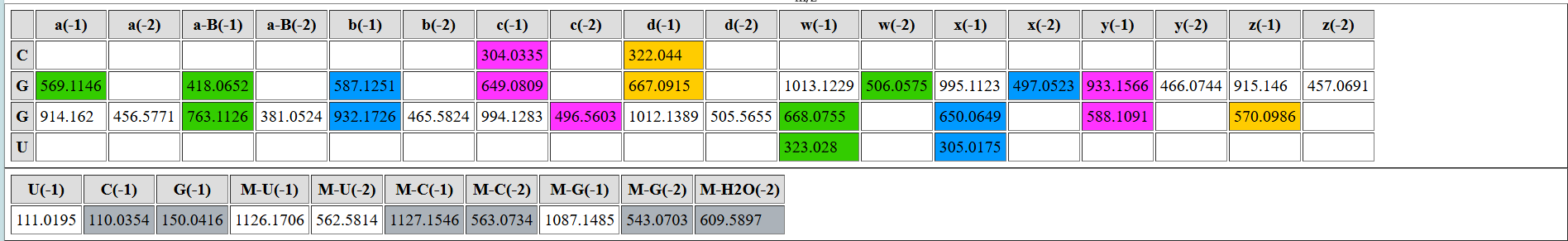

| **digested fragment information** | |
| --- | --- |
| **enzyme** | RNase 4 |
| **unmodified sequence** | AGAGCAU |
| **modified sequence** | N/A |
| **position on *A. c* tRNA^Lys^UUU** | [21-27] |
| **5’ end** | OH |
| **3’ end** | OH |
| **charge state detected** | -3, -2 |
| **theoretical m/z** | 741.1133, 1112.1738 |
| **ppm used for EIC extraction** | 20 |
| **detection of MS1 signal in data files** | |
| 20250216_Lys-1+2-R4 | Y, Y |
| 20250216_Lys-JIA+YI-R4 | N, N |
| 20250227_Lys1+2_R4 | Y, Y |

**Corresponding MS1 EIC**

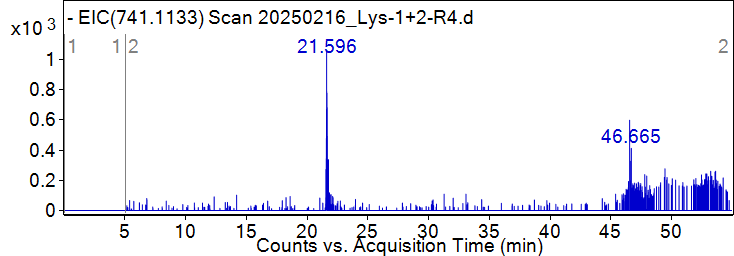

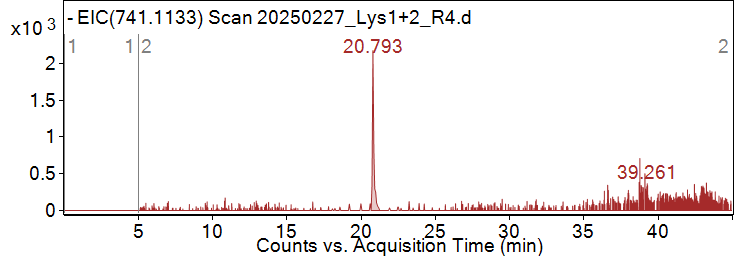

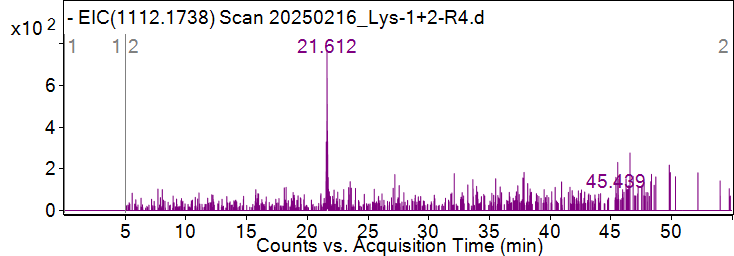

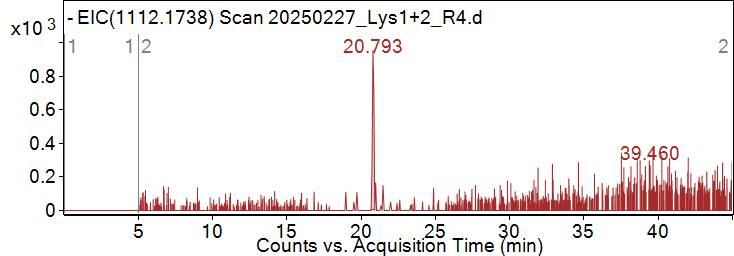

**Pytheas results**

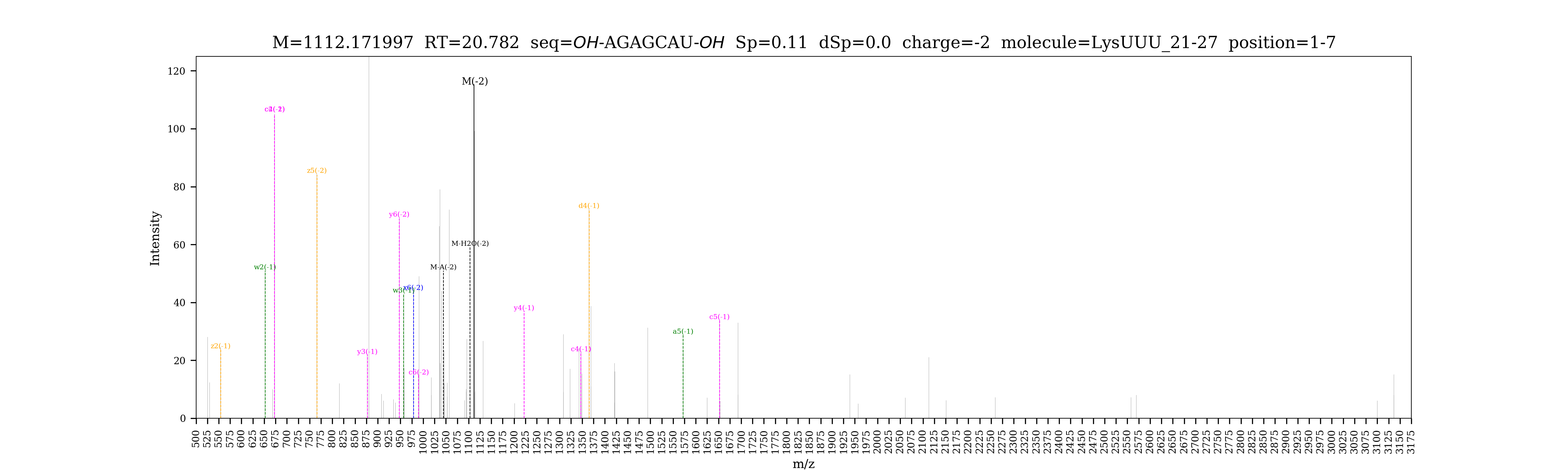

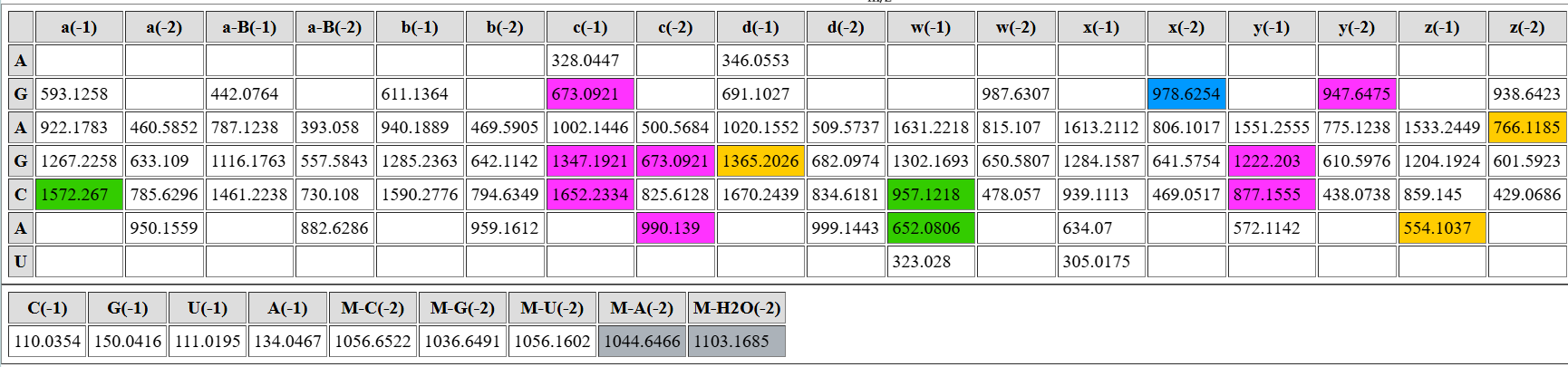

| **digested fragment information** | |
| --- | --- |
| **enzyme** | RNase 4 |
| **unmodified sequence** | CAGACU |
| **modified sequence** | CAGA[s2C]U |
| **position on *A. c* tRNA^Lys^UUU** | [28-33] |
| **5’ end** | OH |
| **3’ end** | OH |
| **charge state detected** | -2 |
| **theoretical m/z** | 935.6331 |
| **ppm used for EIC extraction** | 20 |
| **detection of MS1 signal in data files** | |
| 20250216_Lys-1+2-R4 | Y |
| 20250216_Lys-JIA+YI-R4 | N |
| 20250227_Lys1+2_R4 | Y |

**Corresponding MS1 EIC**

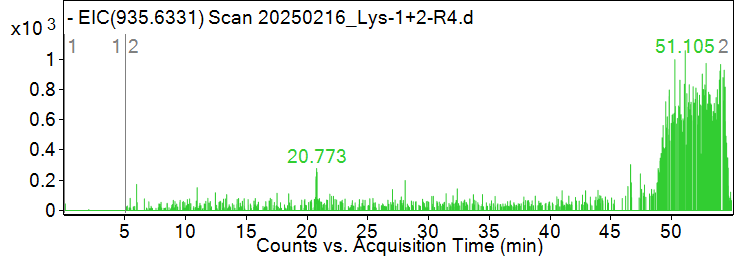

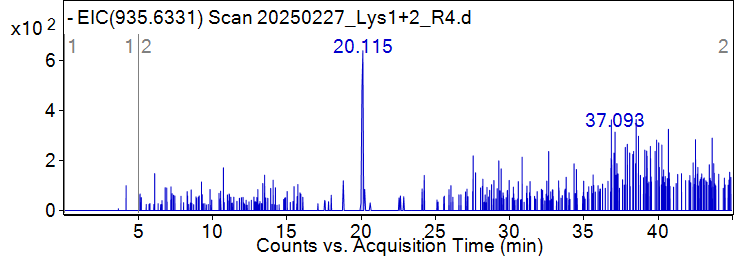
**Pytheas results: No MS2 data to match**

| **digested fragment information** | |
| --- | --- |
| **enzyme** | RNase 4 |
| **unmodified sequence** | CAGACUUU |
| **modified sequence** | CAGA[s2C]U[mcm5s2U]U |
| **position on *A. c* tRNA^Lys^UUU** | [28-35] |
| **5’ end** | OH |
| **3’ end** | OH |
| **charge state detected** | -3 |
| **theoretical m/z** | 856.7691 |
| **ppm used for EIC extraction** | 20 |
| **detection of MS1 signal in data files** | |
| 20250216_Lys-1+2-R4 | Y |
| 20250216_Lys-JIA+YI-R4 | N |
| 20250227_Lys1+2_R4 | Y |

**Corresponding MS1 EIC**

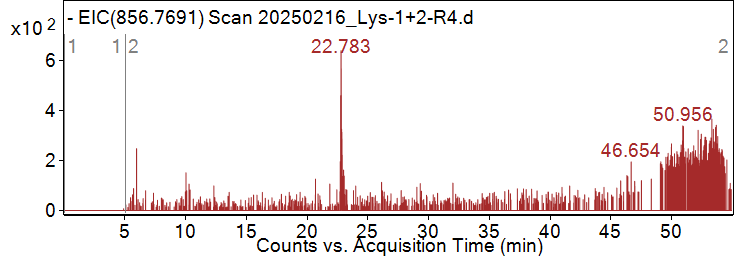

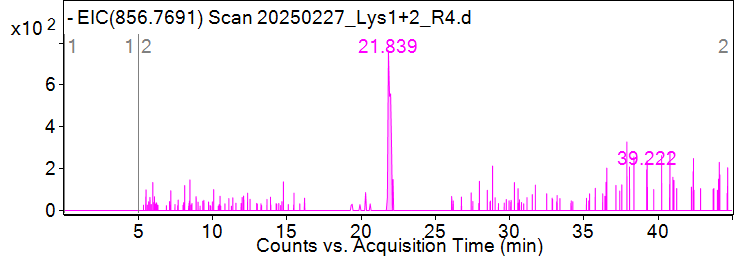

**Pytheas results**

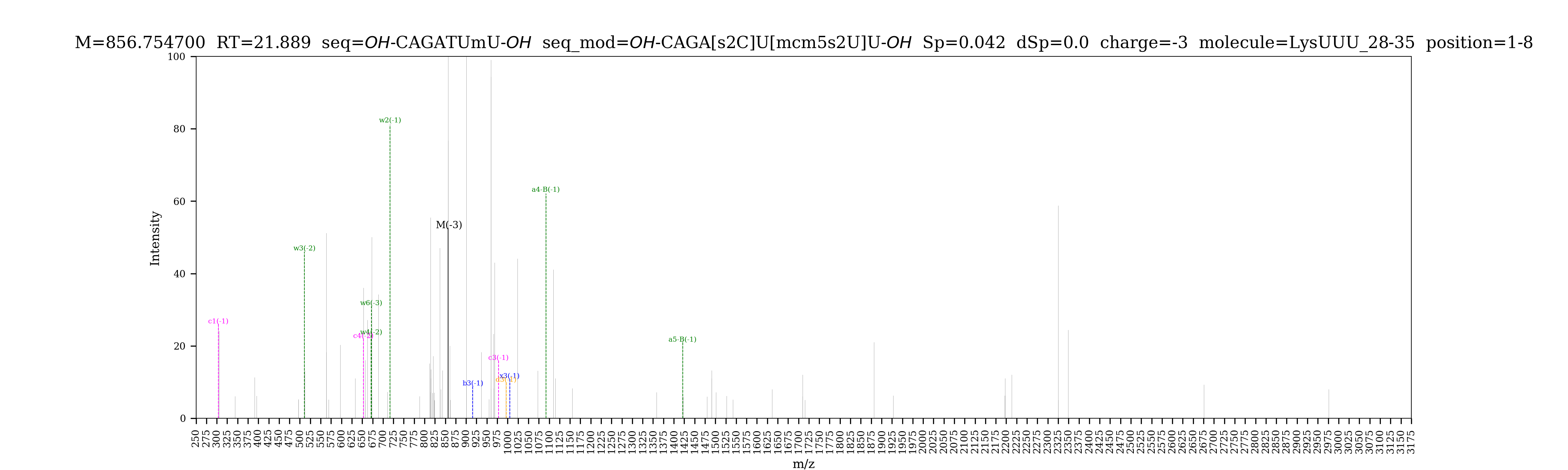

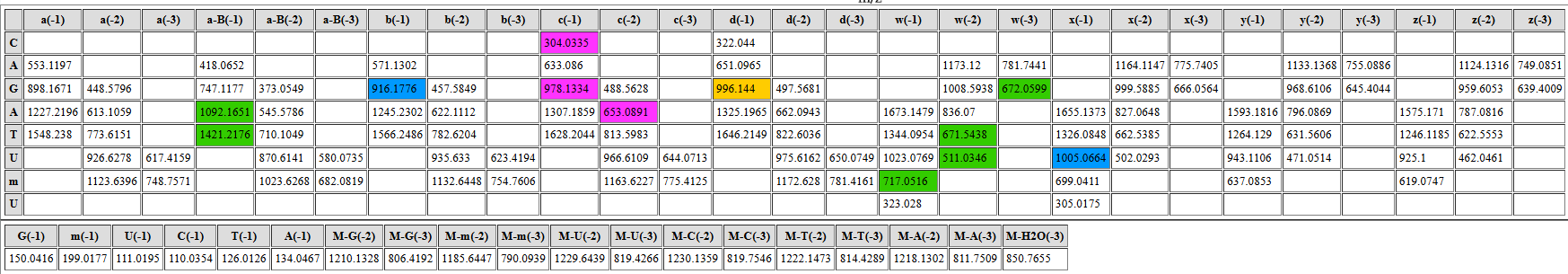

| **digested fragment information** | |
| --- | --- |
| **enzyme** | RNase 4 |
| **unmodified sequence** | UAAUCU |
| **modified sequence** | U[ms2t6A]AU[m5C]U |
| **position on *A. c* tRNA^Lys^UUU** | [36-41] |
| **5’ end** | OH |
| **3’ end** | OH |
| **charge state detected** | -2, -3 |
| **theoretical m/z** | 1011.1458/673.7613 |
| **ppm used for EIC extraction** | 20 |
| **detection of MS1 signal in data files** | |
| 20250216_Lys-1+2-R4 | Y, Y |
| 20250216_Lys-JIA+YI-R4 | Y, Y |
| 20250227_Lys1+2_R4 | Y, Y |

**Corresponding MS1 EIC**

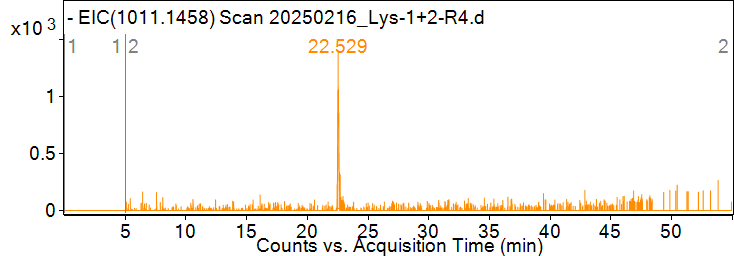

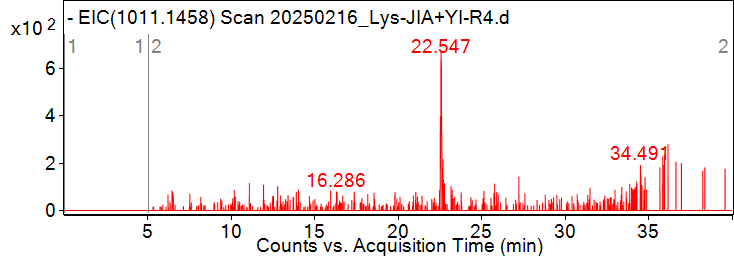

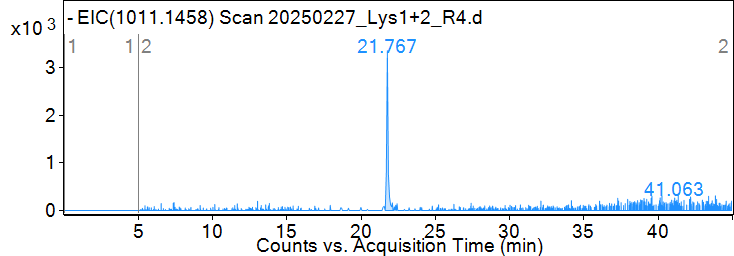

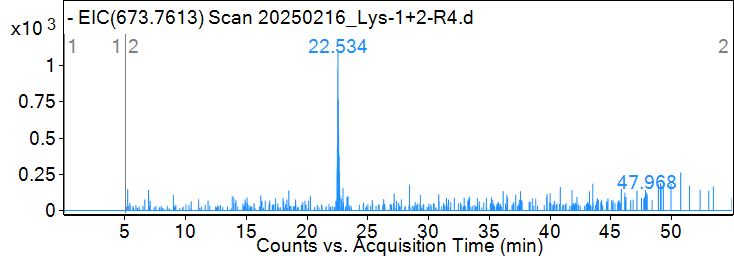

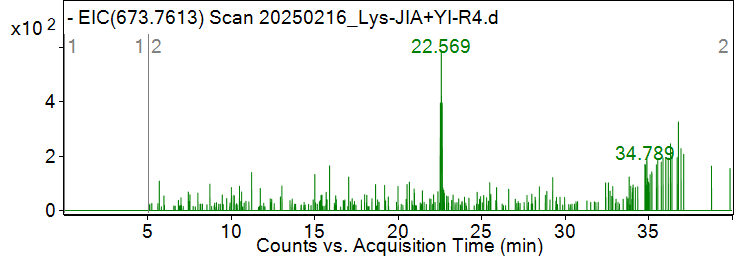

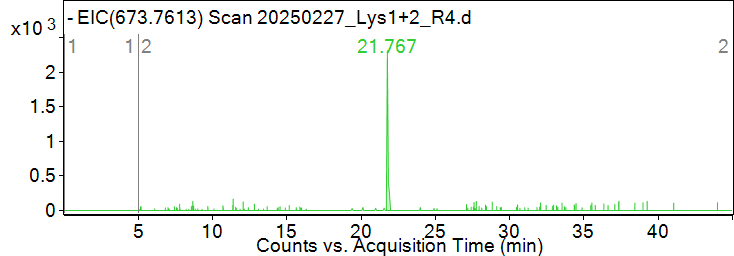

**Pytheas results: no MS2 data to be matched**

| **digested fragment information** | |
| --- | --- |
| **enzyme** | RNase 4 |
| **unmodified sequence** | GAGGGU |
| **modified sequence** | GAGG[m7G]D |
| **position on *A. c* tRNA^Lys^UUU** | [42-47] |
| **5’ end** | OH |
| **3’ end** | OH |
| **charge state detected** | -2 |
| **theoretical m/z** | 983.6637 |
| **ppm used for EIC extraction** | 20 |
| **detection of MS1 signal in data files** | |
| 20250216_Lys-1+2-R4 | N |
| 20250216_Lys-JIA+YI-R4 | N |
| 20250227_Lys1+2_R4 | Y |

**Corresponding MS1 EIC**

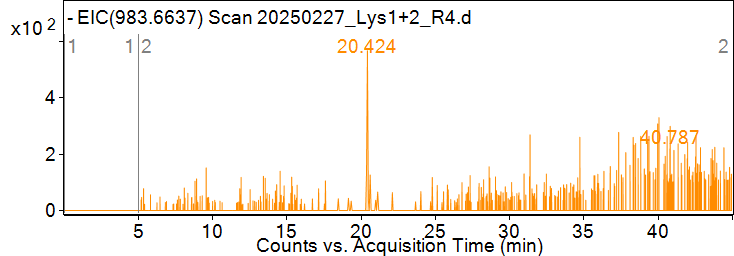

**Pytheas results: no MS2 data to be matched**

| **digested fragment information** | |
| --- | --- |
| **enzyme** | RNase 4 |
| **unmodified sequence** | CAAGGGUU |
| **modified sequence** | [m5C]AAGGG[m5Um]Y |
| **position on *A. c* tRNA^Lys^UUU** | [48-55] |
| **5’ end** | OH |
| **3’ end** | OH |
| **charge state detected** | N/A |
| **theoretical m/z** | 862.4690 (z = -3) |
| **ppm used for EIC extraction** | 20 |
| **detection of MS1 signal in data files** | |
| 20250216_Lys-1+2-R4 | N |
| 20250216_Lys-JIA+YI-R4 | N |
| 20250227_Lys1+2_R4 | N |

**Corresponding MS1 EIC: not detected**

**Pytheas results: no MS2 data to be matched**

| **digested fragment information** | |
| --- | --- |
| **enzyme** | RNase 4 |
| **unmodified sequence** | CGAGU |
| **modified sequence** | CG[m1A]GU |
| **position on *A. c* tRNA^Lys^UUU** | [56-60] |
| **5’ end** | OH |
| **3’ end** | OH |
| **charge state detected** | -2 |
| **theoretical m/z** | 783.1213 (unmodified);  790.1291 (modified) |
| **ppm used for EIC extraction** | 20 |
| **detection of MS1 signal in data files** | |
| 20250216_Lys-1+2-R4 | Y, Y |
| 20250216_Lys-JIA+YI-R4 | Y, Y |
| 20250227_Lys1+2_R4 | Y, Y |

**Corresponding MS1 EIC**

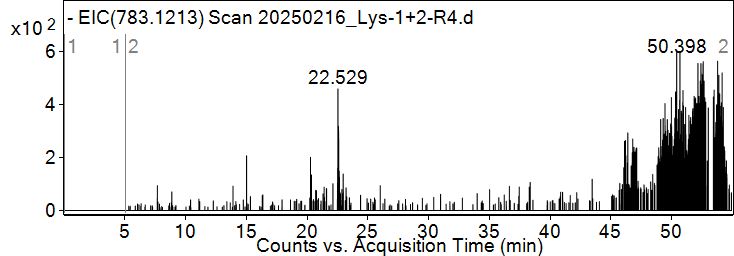

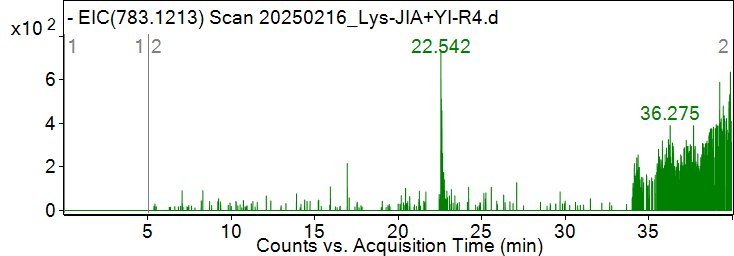

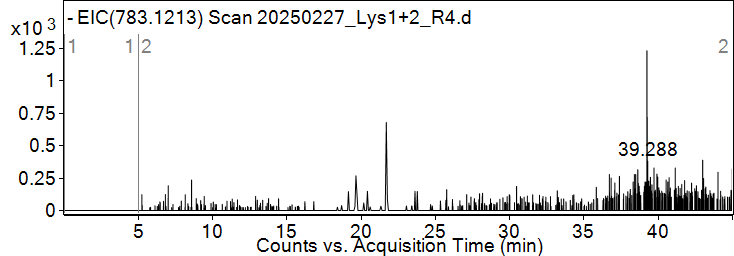

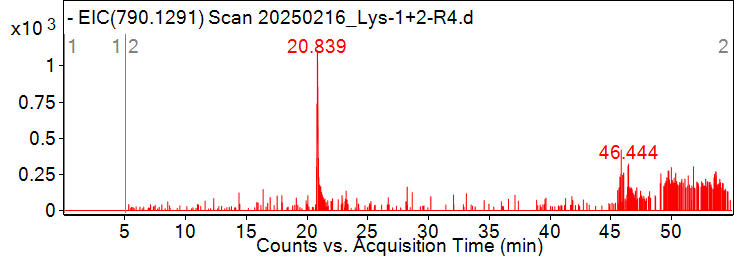

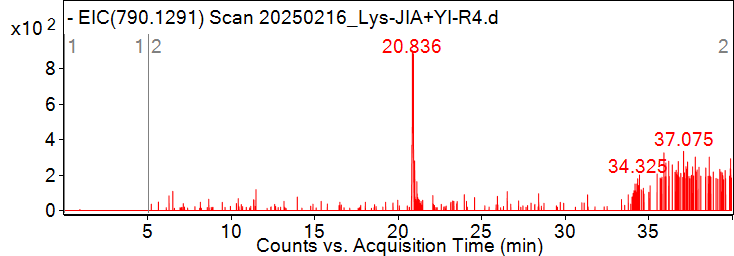

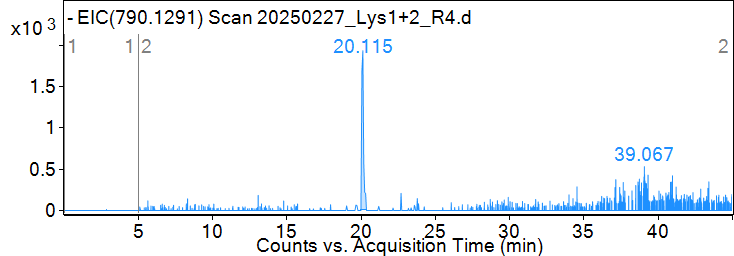

**Pytheas results: matched for CG[m1A]GU**

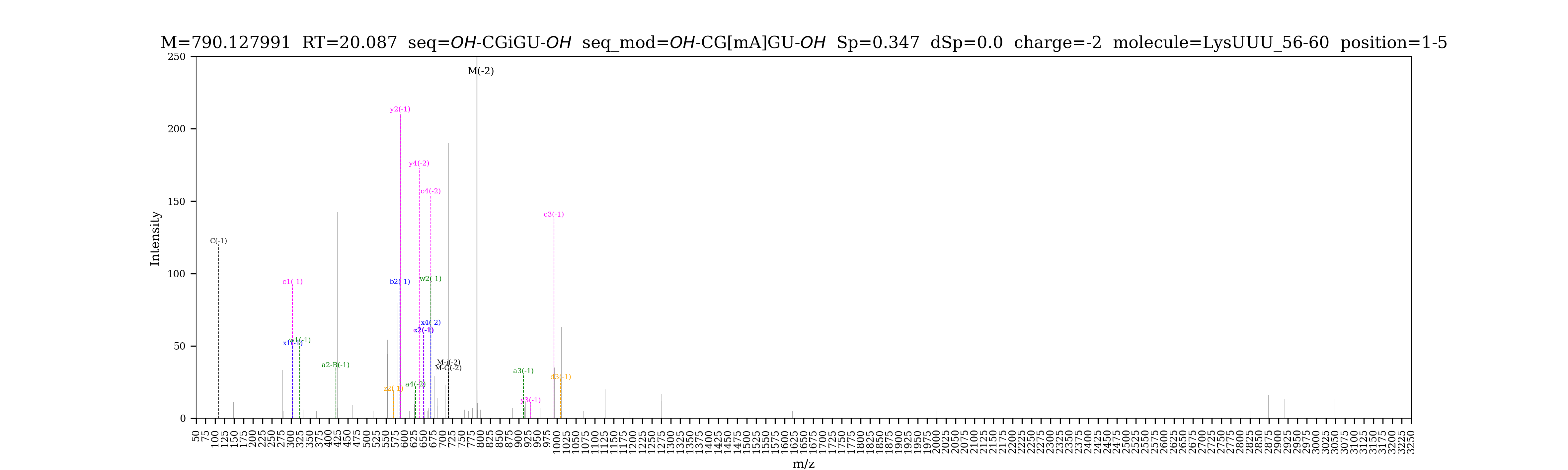

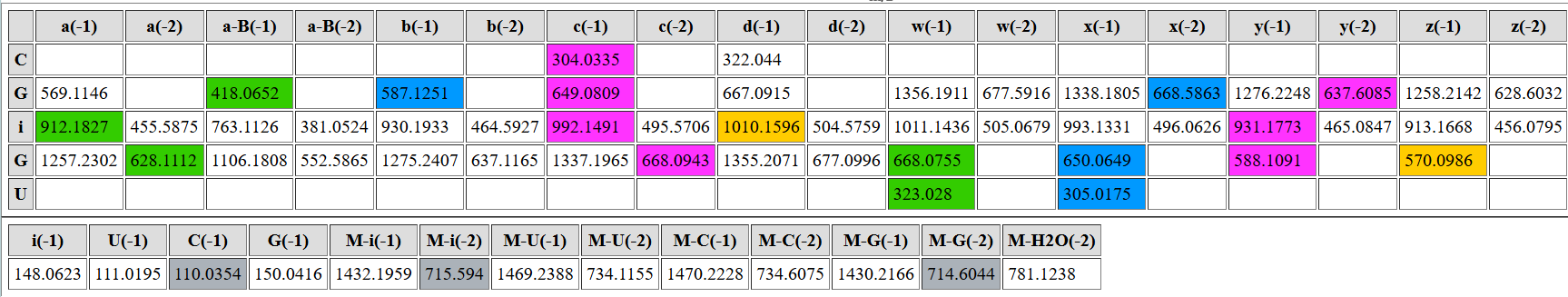

| **digested fragment information** | |
| --- | --- |
| **enzyme** | RNase 4 |
| **unmodified sequence** | CCCUU |
| **modified sequence** | N/A |
| **position on *A. c* tRNA^Lys^UUU** | [61-65] |
| **5’ end** | OH |
| **3’ end** | OH |
| **charge state detected** | -2 |
| **theoretical m/z** | 731.6015 |
| **ppm used for EIC extraction** | 20 |
| **detection of MS1 signal in data files** | |
| 20250216_Lys-1+2-R4 | Y |
| 20250216_Lys-JIA+YI-R4 | Y |
| 20250227_Lys1+2_R4 | Y |

**Corresponding MS1 EIC**

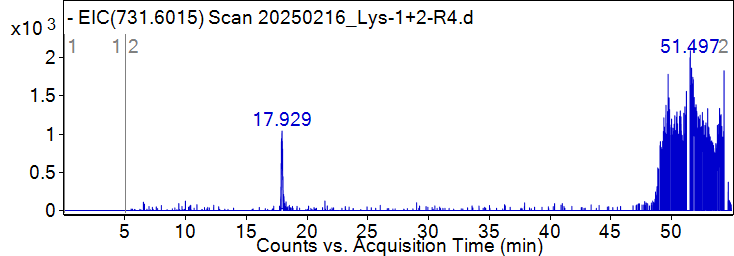

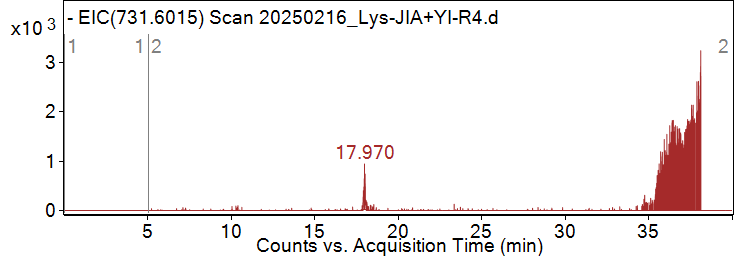

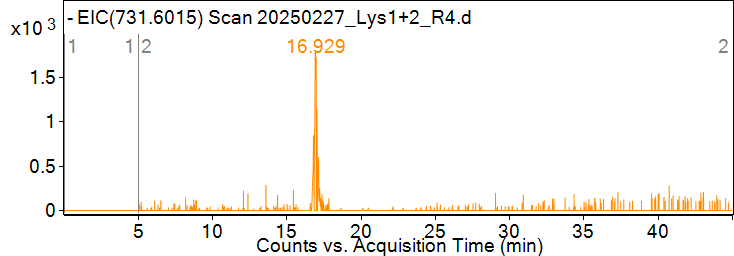
**Pytheas results**

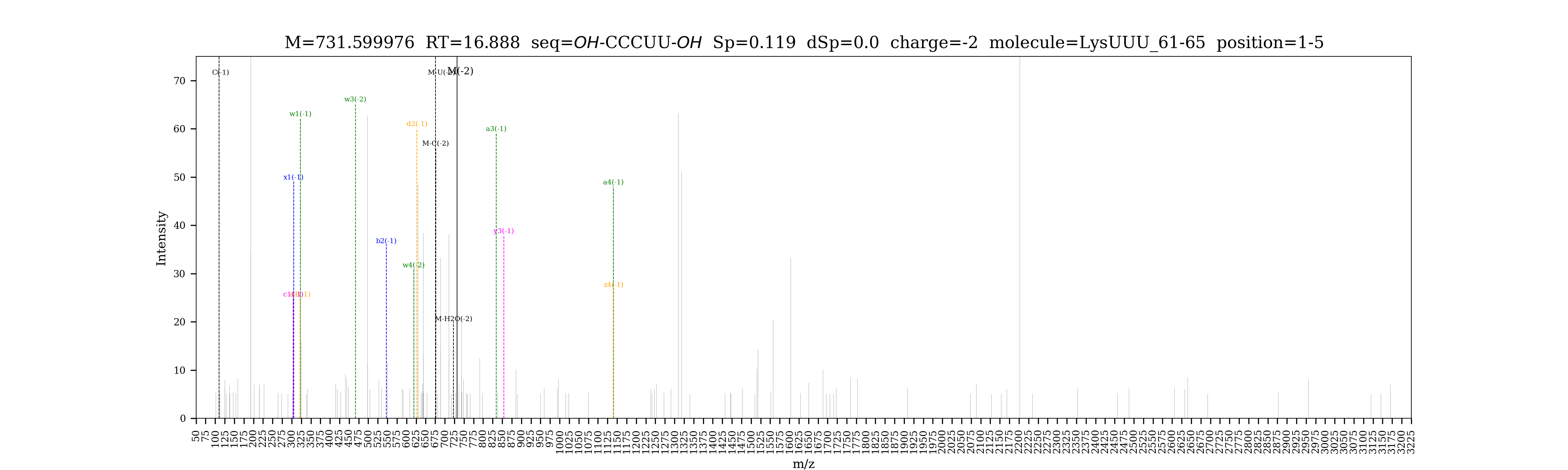

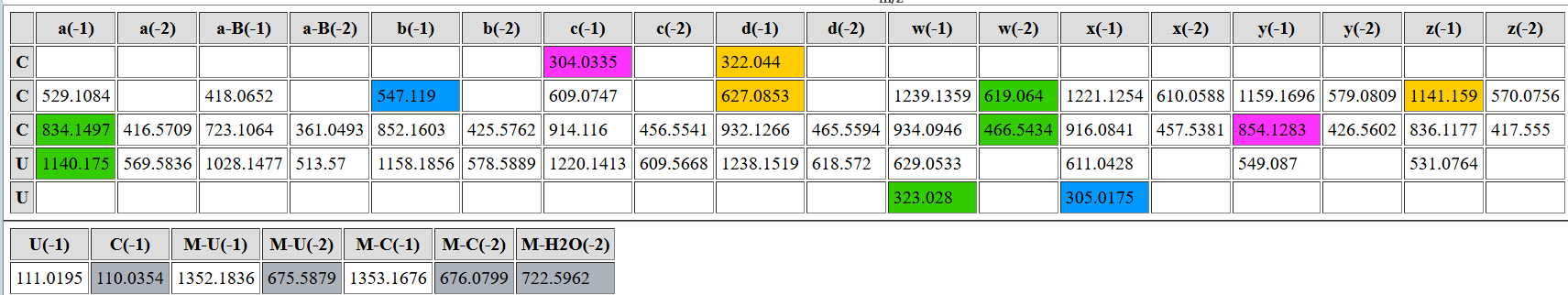

| **digested fragment information** | |
| --- | --- |
| **enzyme** | RNase 4 |
| **unmodified sequence** | AUU |
| **modified sequence** | N/A |
| **position on *A. c* tRNA^Lys^UUU** | [66-68] |
| **5’ end** | OH |
| **3’ end** | OH |
| **charge state detected** | -2 |
| **theoretical m/z** | 438.5659 |
| **ppm used for EIC extraction** | 20 |
| **detection of MS1 signal in data files** | |
| 20250216_Lys-1+2-R4 | Y |
| 20250216_Lys-JIA+YI-R4 | Y |
| 20250227_Lys1+2_R4 | Y |

**Corresponding MS1 EIC**

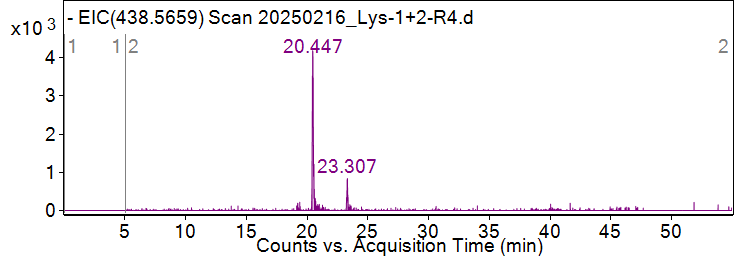

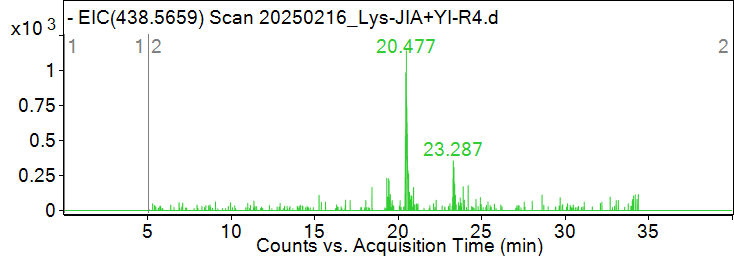

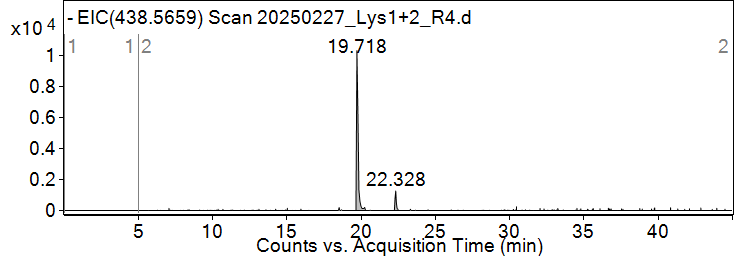
**Pytheas results**

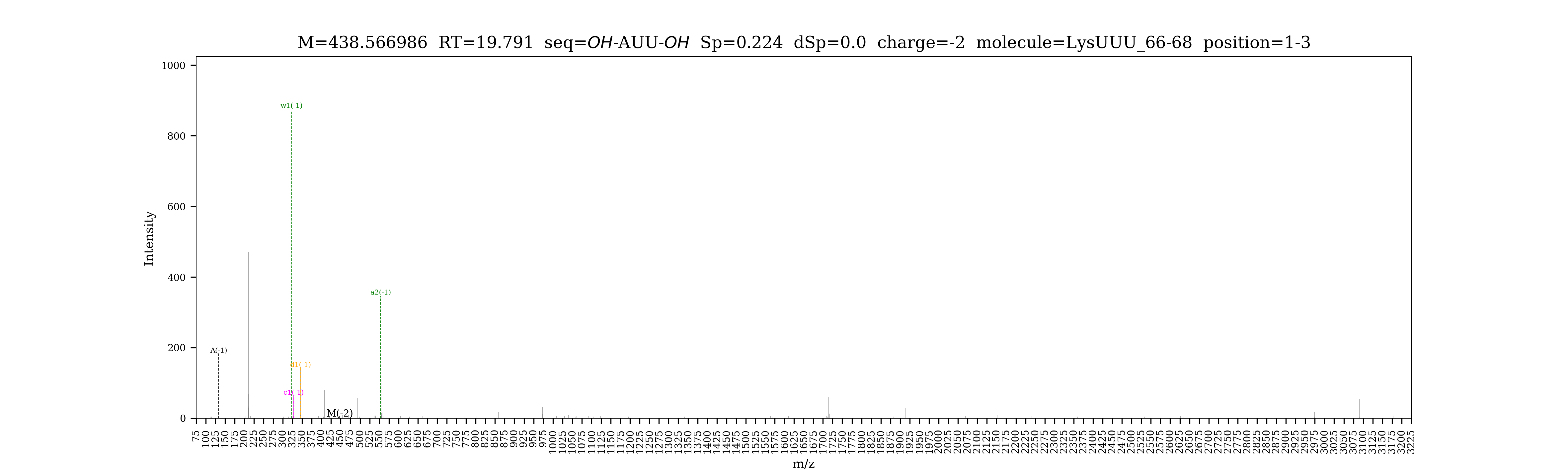

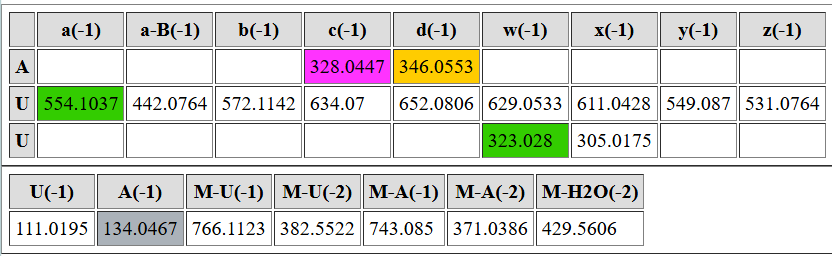

| **digested fragment information** | |
| --- | --- |
| **enzyme** | RNase 4 |
| **unmodified sequence** | GGGCGCCA |
| **modified sequence** | N/A |
| **position on *A. c* tRNA^Lys^UUU** | [69-76] |
| **5’ end** | OH |
| **3’ end** | OH |
| **charge state detected** | -3 |
| **theoretical m/z** | 853.1290 |
| **ppm used for EIC extraction** | 20 |
| **detection of MS1 signal in data files** | |
| 20250216_Lys-1+2-R4 | Y |
| 20250216_Lys-JIA+YI-R4 | Y |
| 20250227_Lys1+2_R4 | Y |

**Corresponding MS1 EIC**

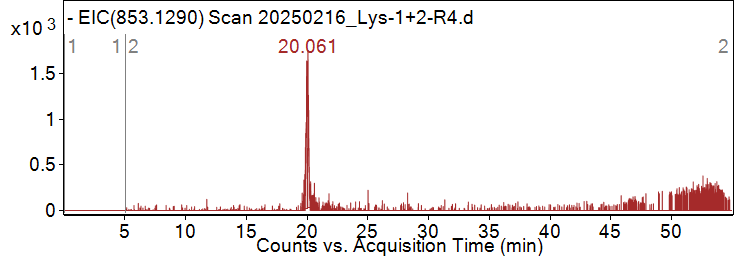

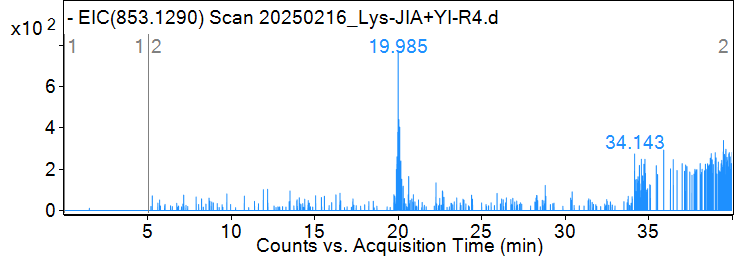

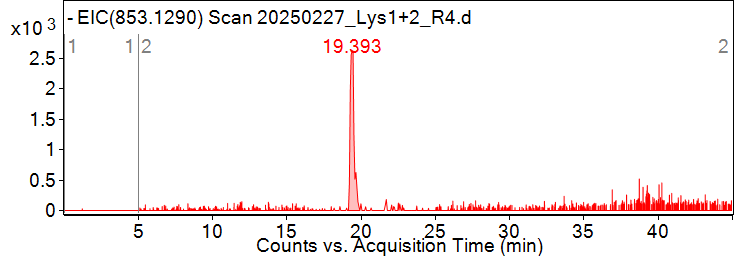
**Pytheas results**

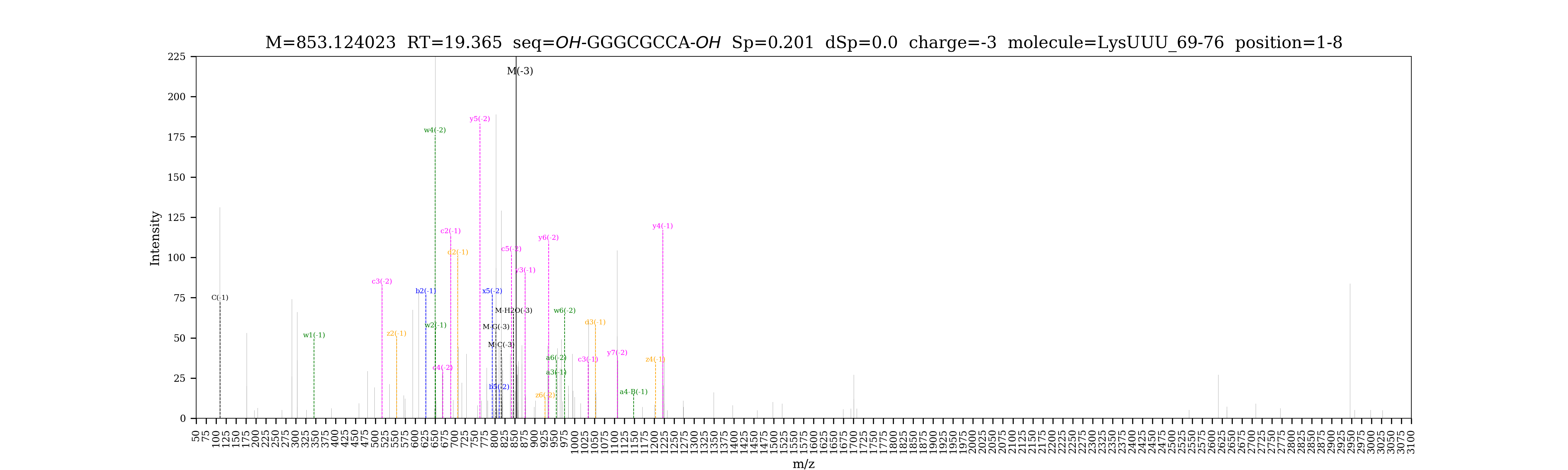

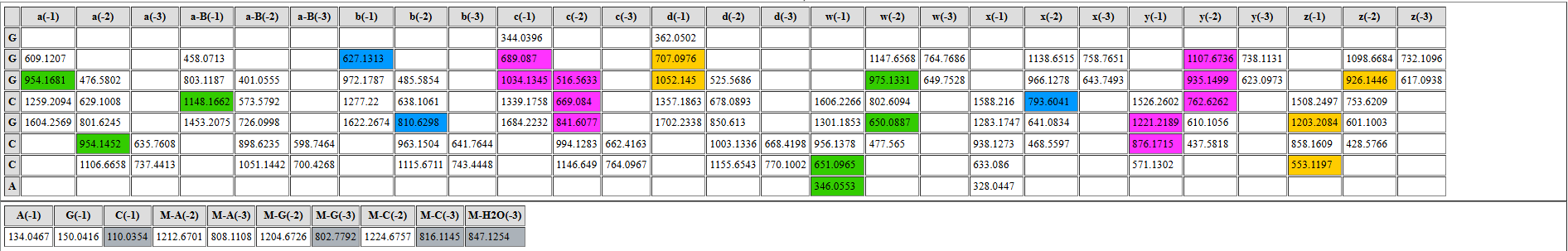

**Part 2. RNase T1 data**

| **list of LC-MS/MS data files used for analysis by RNase T1 digestion** | |
| --- | --- |
| **data file name** | **tissue** |
| 20250504_Lys_T1 | penis/heart pooled from multiple slugs |
| 20250528_Lys_CNS | pooled neurons from multiple slugs |
| 20250528_Lys_Ac#11_unk3* | accessory genital mass from slug #11 |
| *Pytheas results are based on this file unless otherwise indicated. | |

| **digested fragment information** | |
| --- | --- |
| **enzyme** | RNase T1 |
| **unmodified sequence** | CCCG |
| **modified sequence** | N/A |
| **position on *A. c* tRNA^Lys^UUU** | [2-5] |
| **5’ end** | OH |
| **3’ end** | OH |
| **charge state detected** | -2 |
| **theoretical m/z** | 598.0999 |
| **ppm used for EIC extraction** | 20 |
| **detection of MS1 signal in data files** | |
| 20250504_Lys_T1 | Y |
| 20250528_Lys_CNS | Y |
| 20250528_Lys_Ac#11_unk3 | Y |

**Corresponding MS1 EIC**

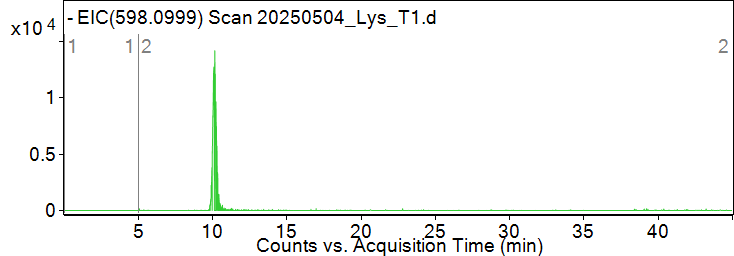

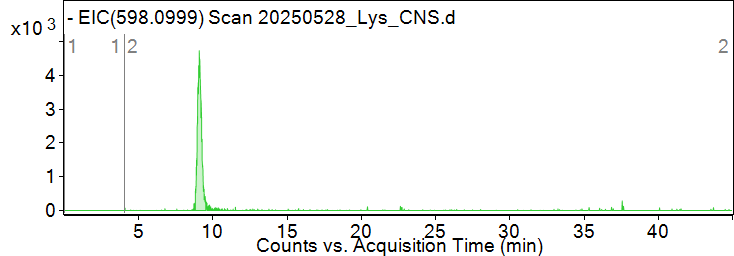

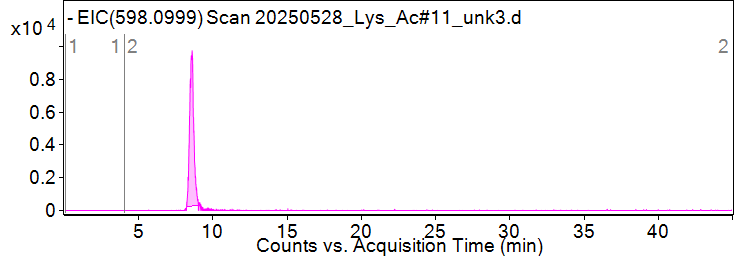

**Pytheas results**

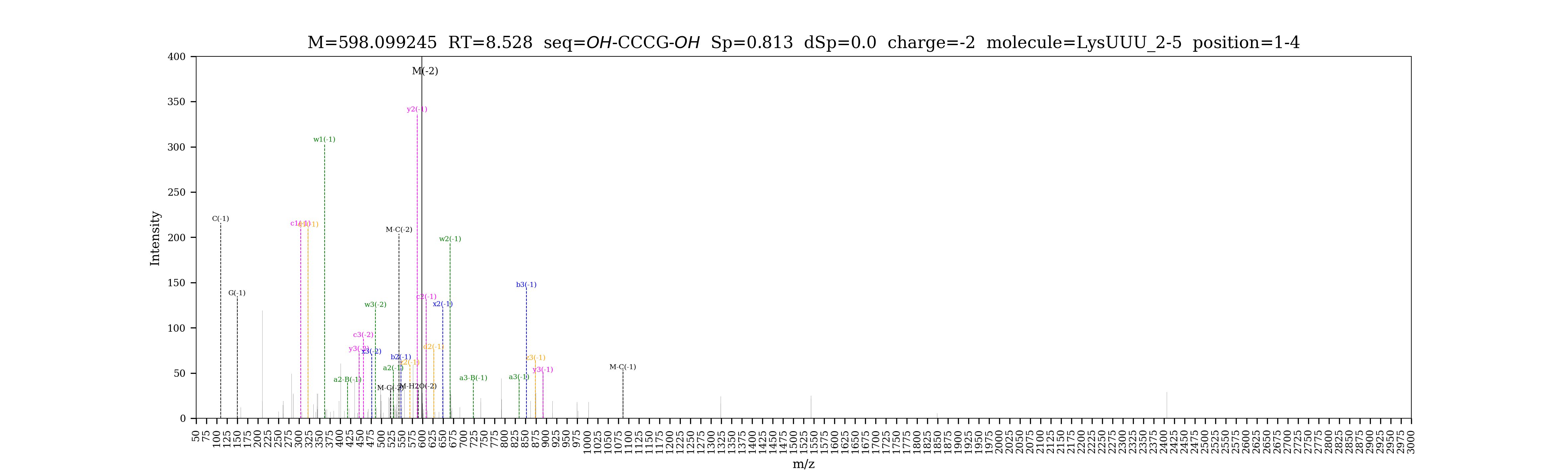

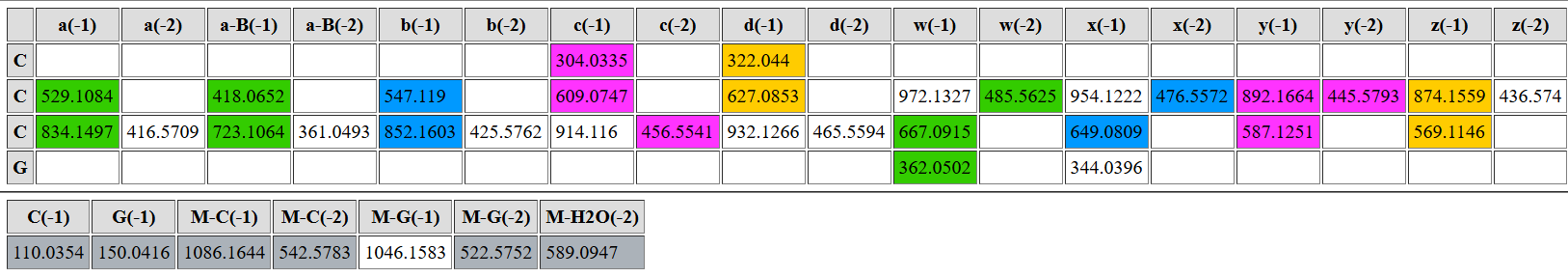

| **digested fragment information** | |
| --- | --- |
| **enzyme** | RNase T1 |
| **unmodified sequence** | UUAG |
| **modified sequence** | UUA[m2G] |
| **position on *A. c* tRNA^Lys^UUU** | [7-10] |
| **5’ end** | OH |
| **3’ end** | OH |
| **charge state detected** | -2 |
| **theoretical m/z** | 618.0974 |
| **ppm used for EIC extraction** | 20 |
| **detection of MS1 signal in data files** | |
| 20250504_Lys_T1 | Y |
| 20250528_Lys_CNS | Y |
| 20250528_Lys_Ac#11_unk3 | Y |

**Corresponding MS1 EIC**

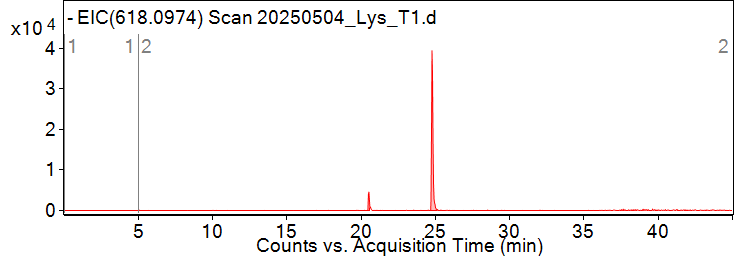

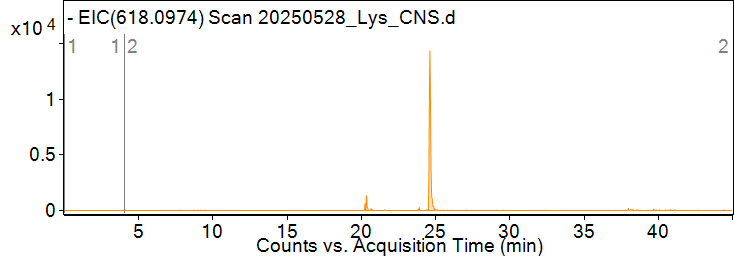

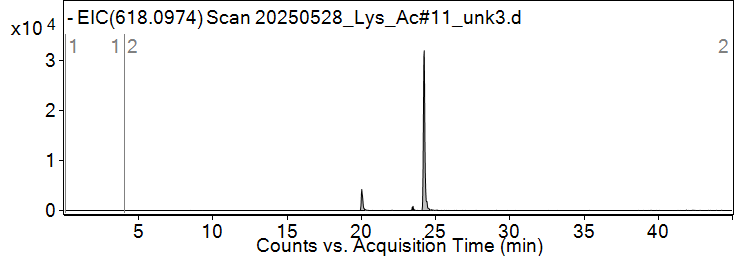

*The smaller, earlier peak is for CAUCAG (charge = -3, m/z = 618.0937)

**Pytheas results**

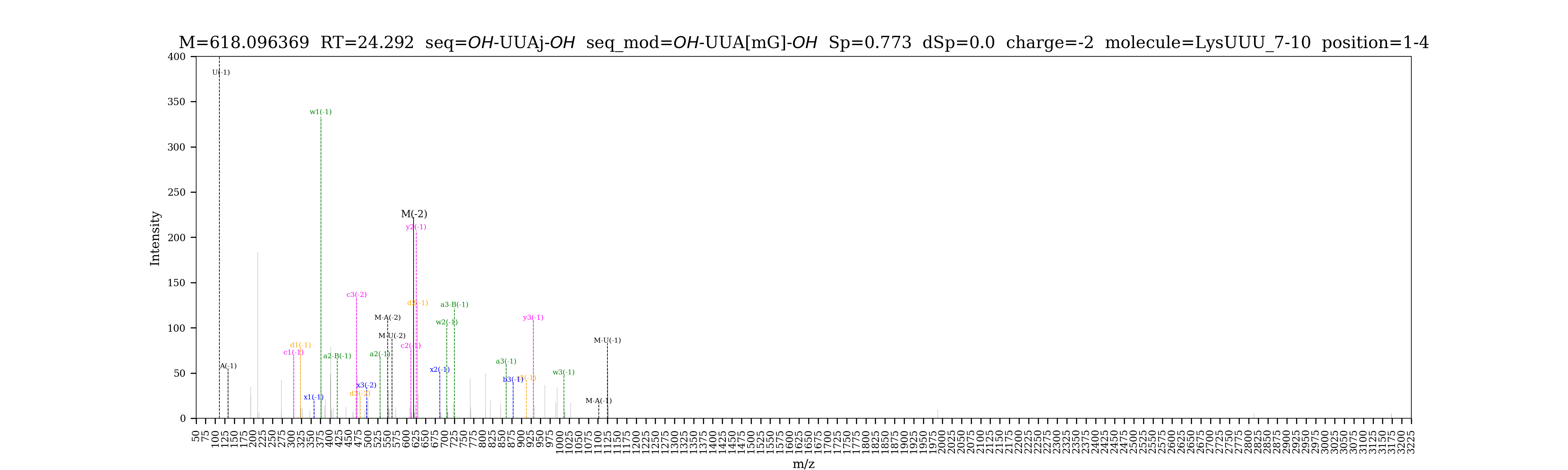

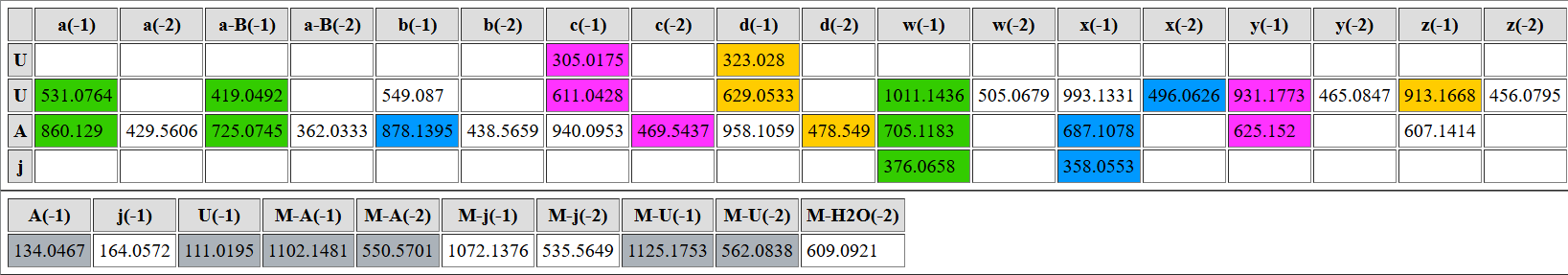

| **digested fragment information** | |
| --- | --- |
| **enzyme** | RNase T1 |
| **unmodified sequence** | CUCAG |
| **modified sequence** | N/A |
| **position on *A. c* tRNA^Lys^UUU** | [11-15] |
| **5’ end** | OH |
| **3’ end** | OH |
| **charge state detected** | -2 |
| **theoretical m/z** | 763.1182 |
| **ppm used for EIC extraction** | 20 |
| **detection of MS1 signal in data files** | |
| 20250504_Lys_T1 | Y |
| 20250528_Lys_CNS | Y |
| 20250528_Lys_Ac#11_unk3 | Y |

**Corresponding MS1 EIC**

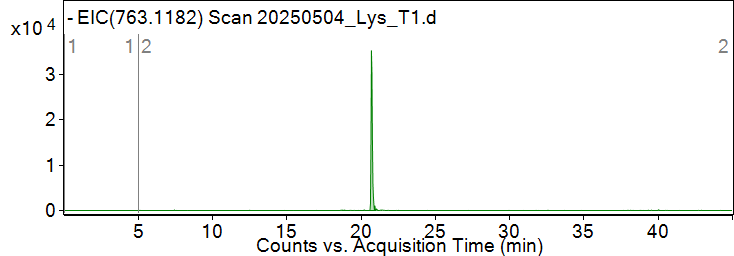

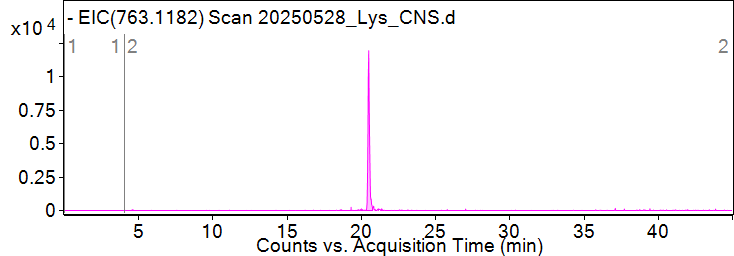

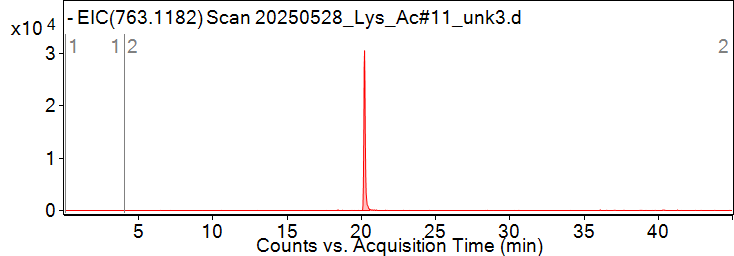

**Pytheas results**

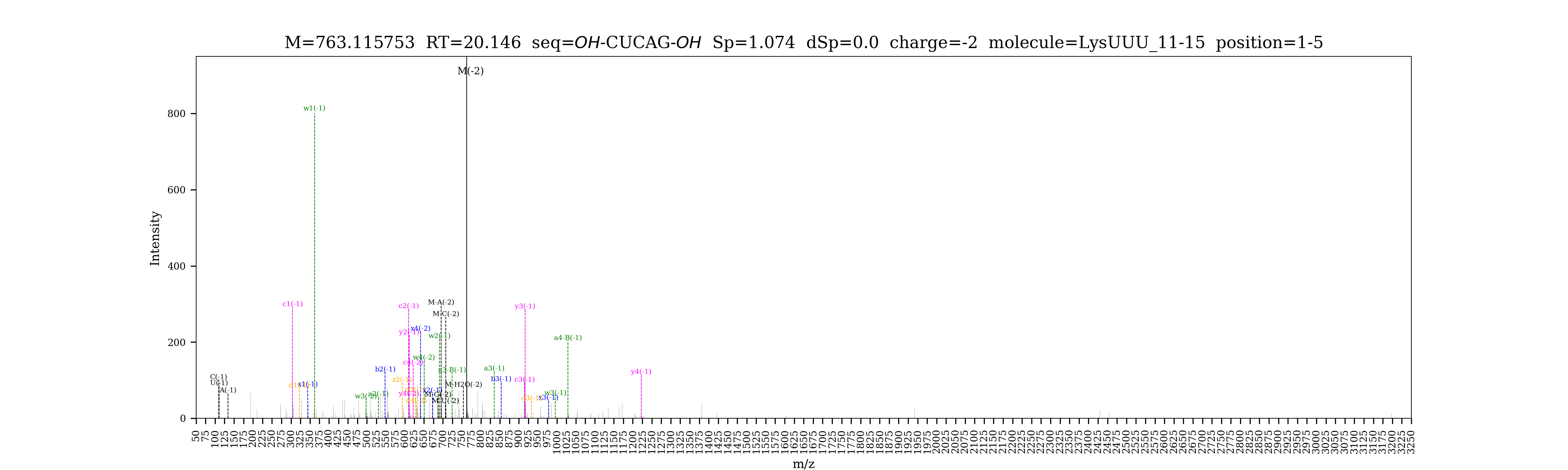

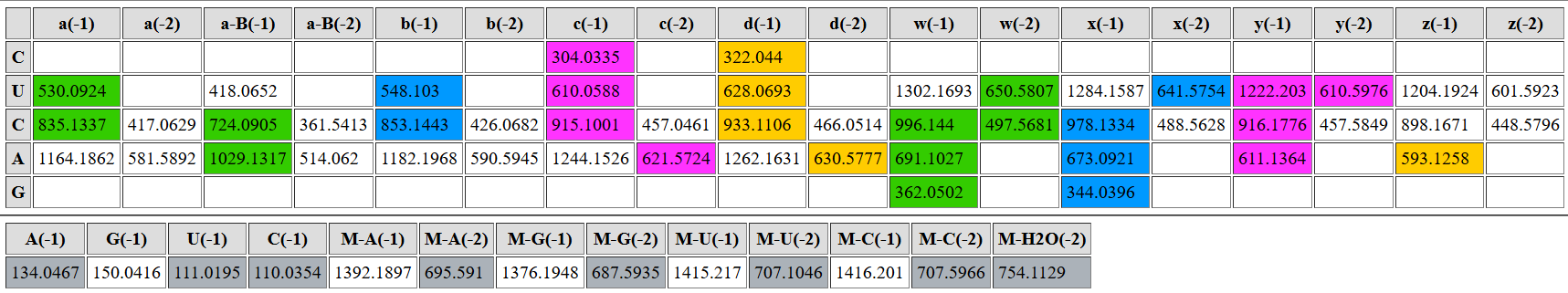

| **digested fragment information** | |
| --- | --- |
| **enzyme** | RNase T1 |
| **unmodified sequence** | UCG |
| **modified sequence** | DCG |
| **position on *A. c* tRNA^Lys^UUU** | [16-18] |
| **5’ end** | OH |
| **3’ end** | OH |
| **charge state detected** | -2, -1 |
| **theoretical m/z** | 446.0713 (unmodified);  447.0791 (modified); 895.1661 (modified) |
| **ppm used for EIC extraction** | 20 |
| **detection of MS1 signal in data files** | |
| 20250504_Lys_T1 | Y, Y, Y |
| 20250528_Lys_CNS | Y, Y, Y |
| 20250528_Lys_Ac#11_unk3 | Y, Y, Y |

*Signal for unmodified variant is weak

**Corresponding MS1 EIC**

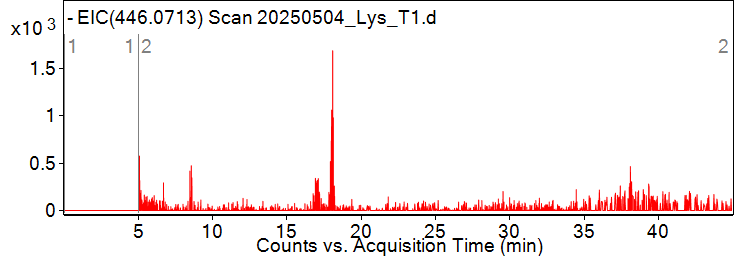

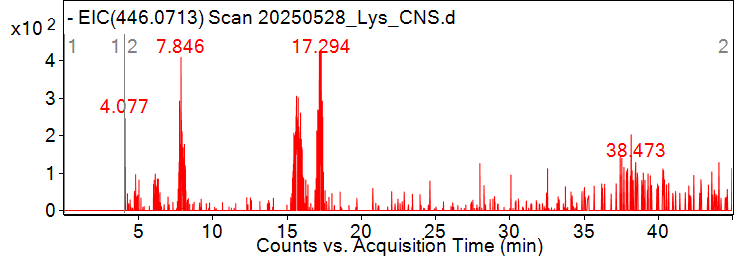

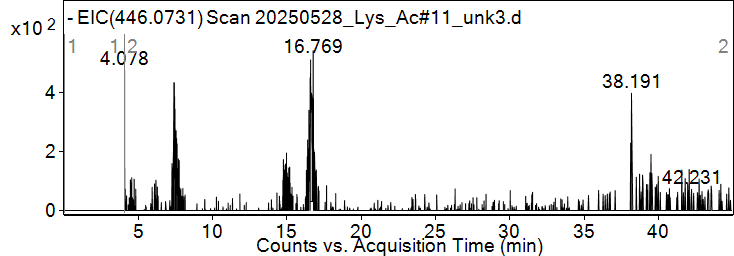

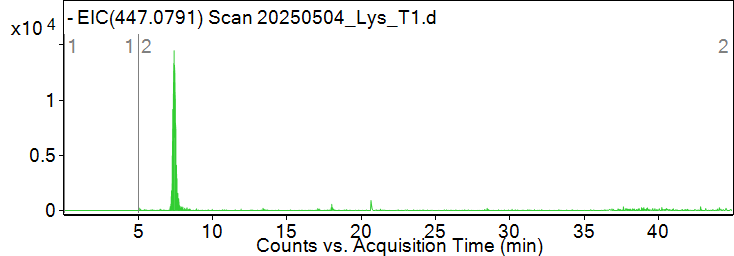

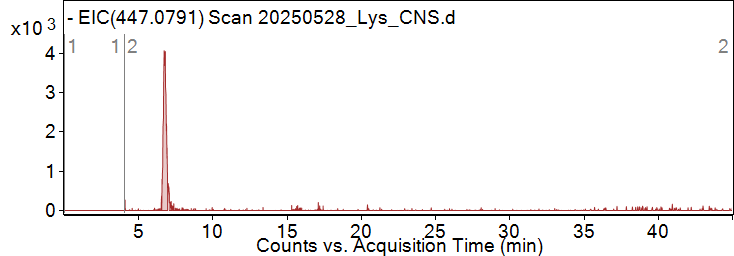

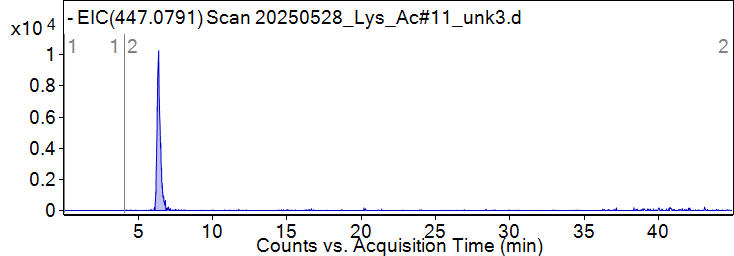

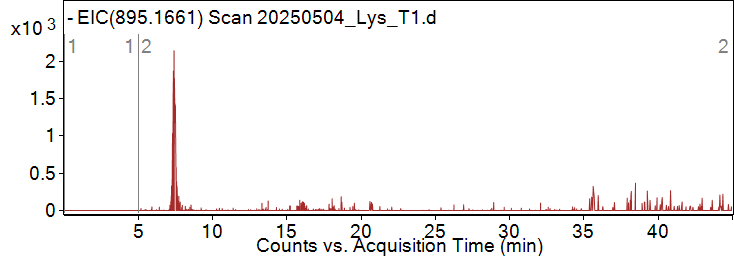

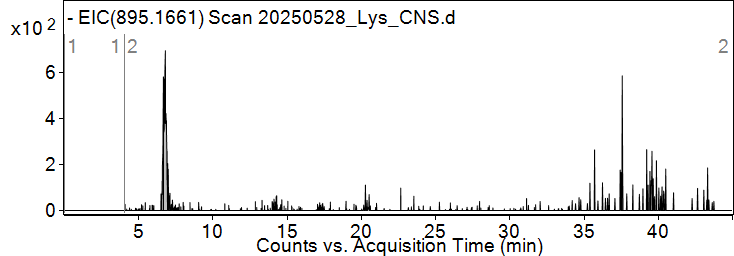

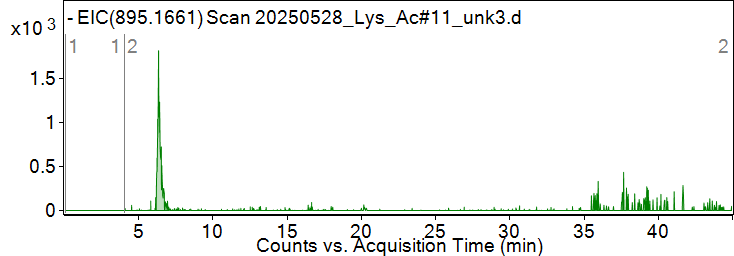

**Pytheas results**

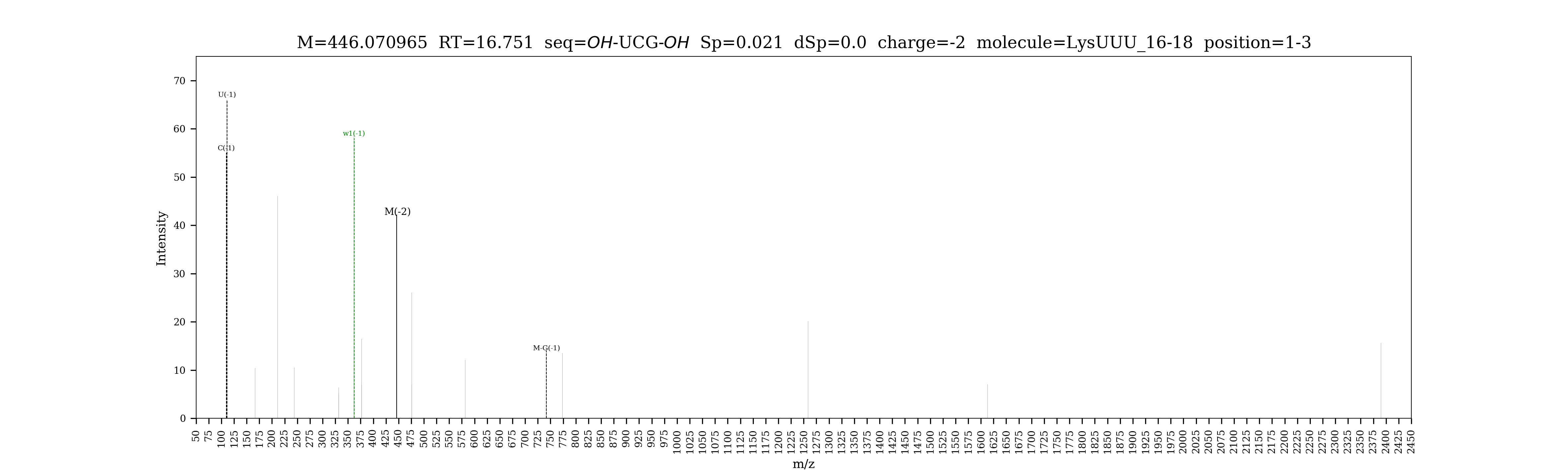

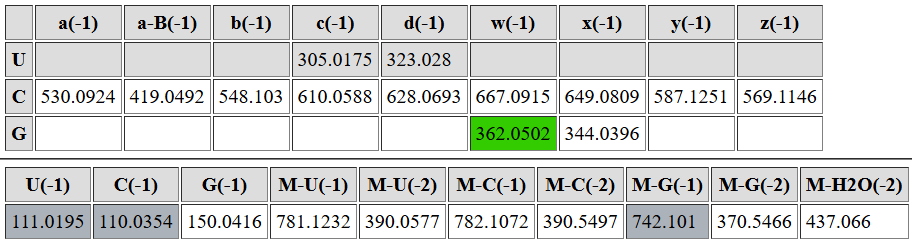

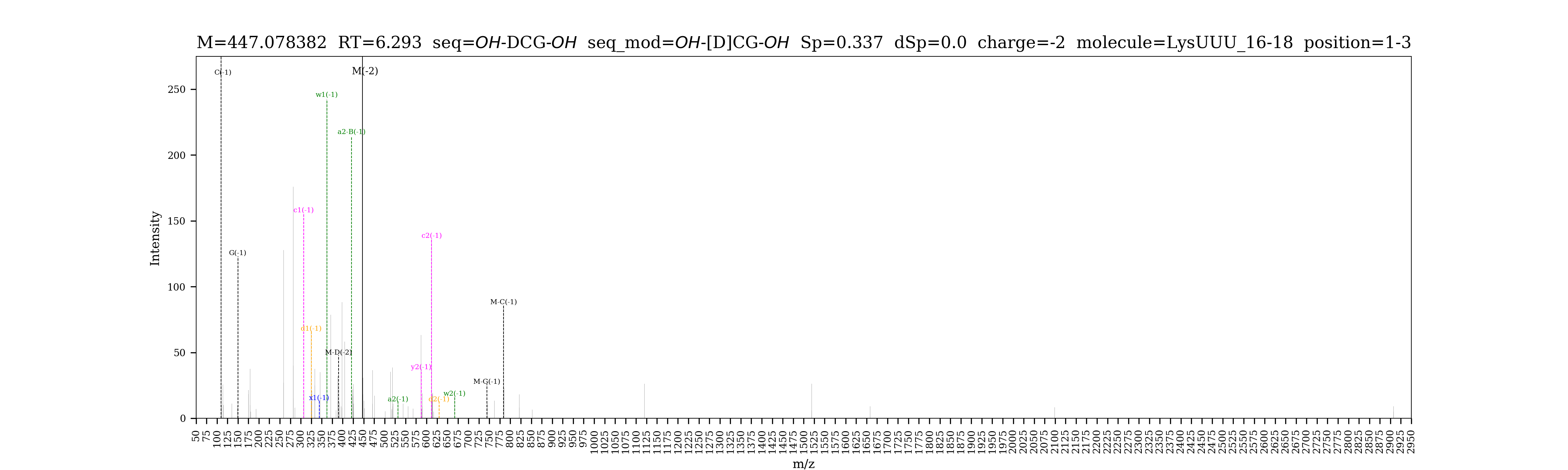

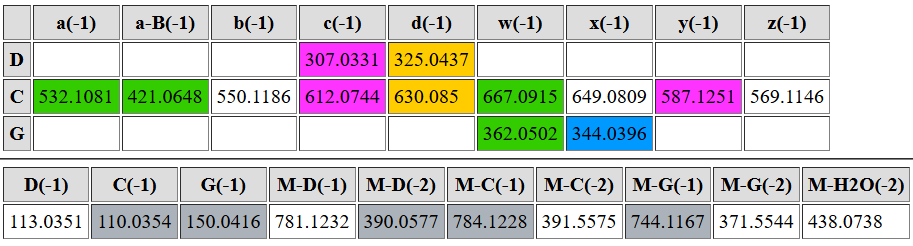

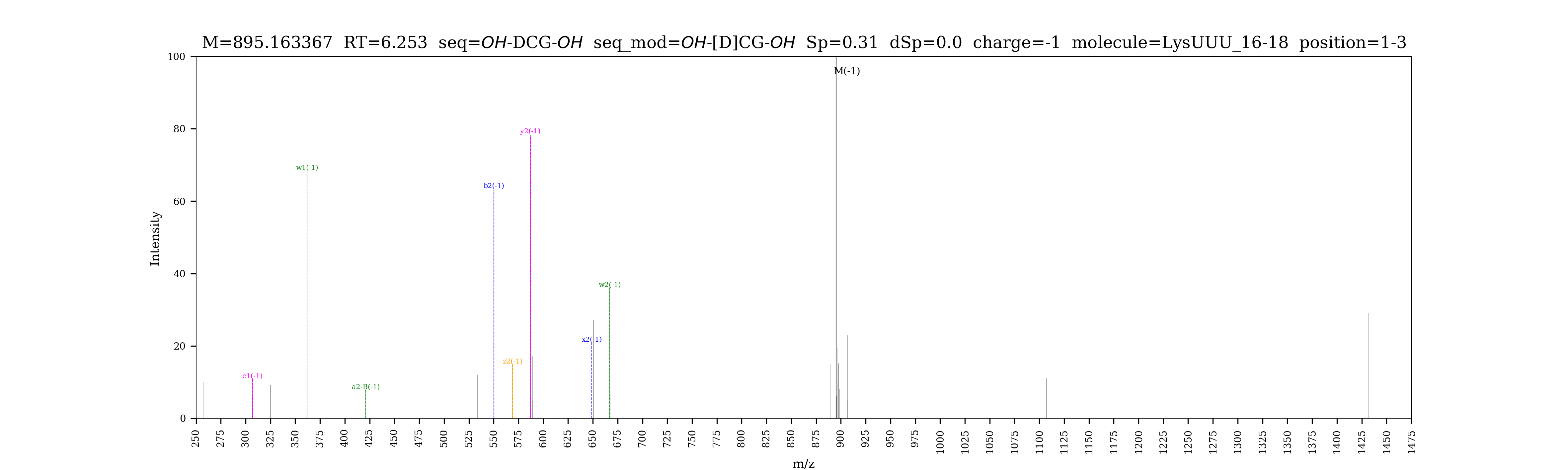

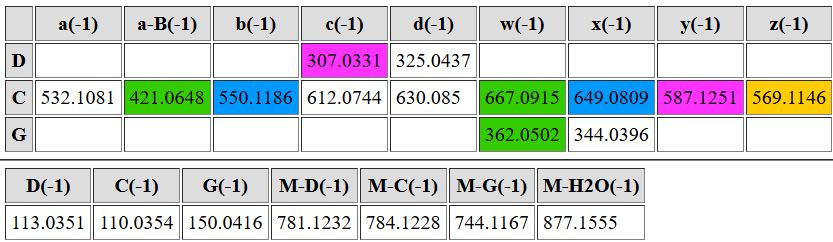

| **digested fragment information** | |
| --- | --- |
| **enzyme** | RNase T1 |
| **unmodified sequence** | UAG |
| **modified sequence** | N/A |
| **position on *A. c* tRNA^Lys^UUU** | [20-22] |
| **5’ end** | OH |
| **3’ end** | OH |
| **charge state detected** | -2, -1 |
| **theoretical m/z** | 458.0769, 917.1617 |
| **ppm used for EIC extraction** | 20 |
| **detection of MS1 signal in data files** | |
| 20250504_Lys_T1 | Y, Y |
| 20250528_Lys_CNS | Y, Y |
| 20250528_Lys_Ac#11_unk3 | Y, Y |

**Corresponding MS1 EIC**

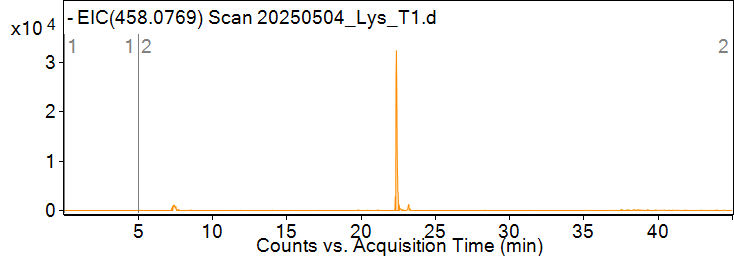

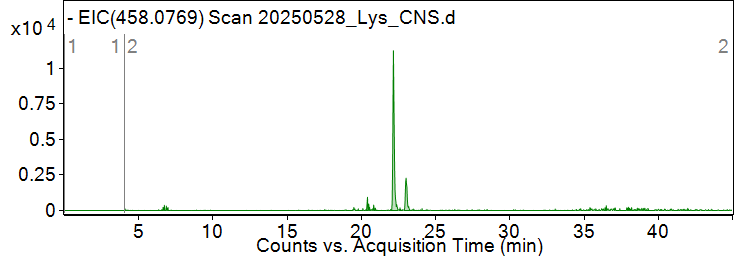

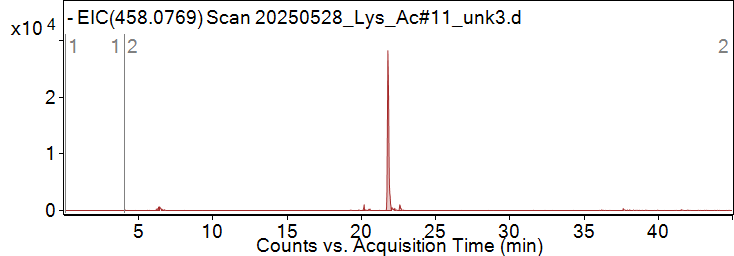

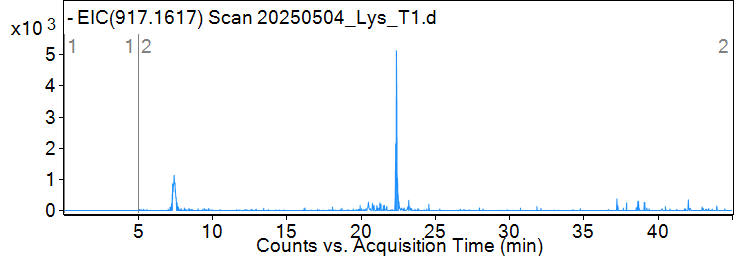

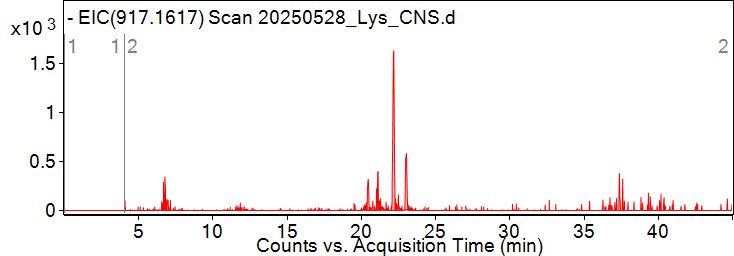

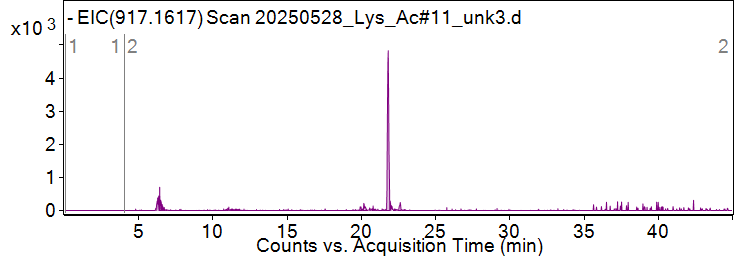

**Pytheas results**

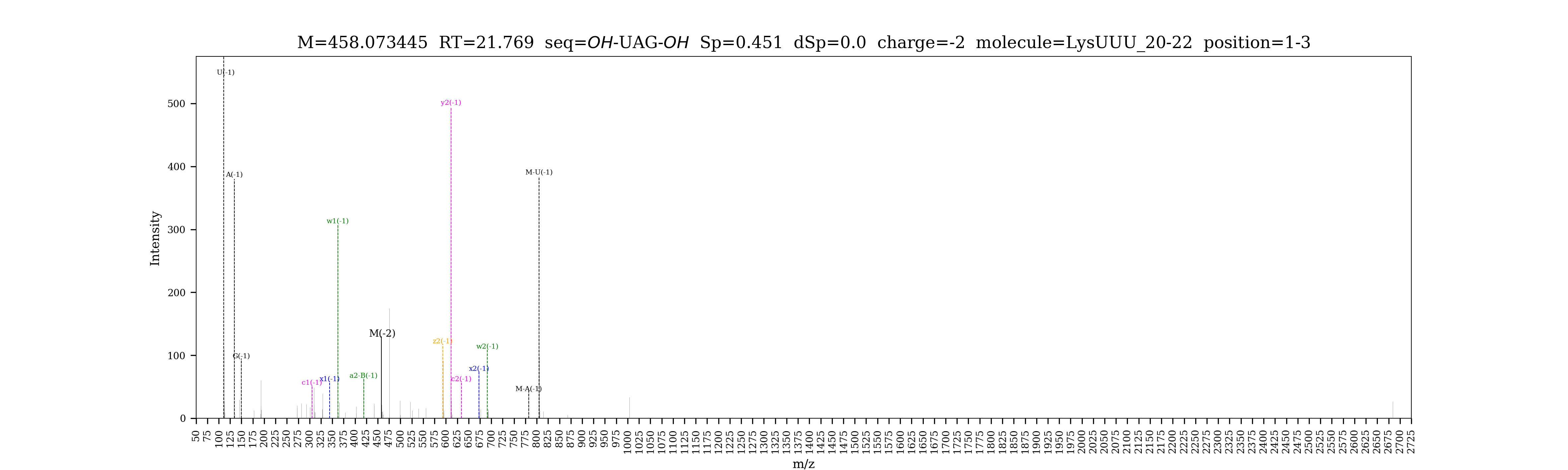

| **digested fragment information** | |
| --- | --- |
| **enzyme** | RNase T1 |
| **unmodified sequence** | AG |
| **modified sequence** | N/A |
| **position on *A. c* tRNA^Lys^UUU** | [23-24] |
| **5’ end** | OH |
| **3’ end** | OH |
| **charge state detected** | -1 |
| **theoretical m/z** | 611.1364 |
| **ppm used for EIC extraction** | 20 |
| **detection of MS1 signal in data files** | |
| 20250504_Lys_T1 | Y |
| 20250528_Lys_CNS | Y |
| 20250528_Lys_Ac#11_unk3 | Y |

**Corresponding MS1 EIC**

**Pytheas results: no MS2 data to be matched**

| **digested fragment information** | |
| --- | --- |
| **enzyme** | RNase T1 |
| **unmodified sequence** | CAUCAG |
| **modified sequence** | N/A |
| **position on *A. c* tRNA^Lys^UUU** | [25-30] |
| **5’ end** | OH |
| **3’ end** | OH |
| **charge state detected** | -2, -3 |
| **theoretical m/z** | 927.6445, 618.0937 |
| **ppm used for EIC extraction** | 20 |
| **detection of MS1 signal in data files** | |
| 20250504_Lys_T1 | Y, Y |
| 20250528_Lys_CNS | Y, Y |
| 20250528_Lys_Ac#11_unk3 | Y, Y |

**Corresponding MS1 EIC**

**Pytheas results**

| **digested fragment information** | |
| --- | --- |
| **enzyme** | RNase T1 |
| **unmodified sequence** | ACUUUUAAUCUG |
| **modified sequence** | A[s^2^C]U[mcm^5^s^2^U]UU[ms^2^t^6^A]AU[m^5^C]UG;  ACU[mcm^5^s^2^U]UU[ms^2^t^6^A]AU[m^5^C]UG |
| **position on *A. c* tRNA^Lys^UUU** | [31-42] |
| **5’ end** | OH |
| **3’ end** | OH |
| **charge state detected** | -3, -4 |
| **theoretical m/z** | 1340.8255, 1005.3671;  1335.4998, 1001.3729 |
| **ppm used for EIC extraction** | 20 |
| **detection of MS1 signal in data files** | |
| 20250504_Lys_T1 | Y, Y; N/A |
| 20250528_Lys_CNS | Y, Y; N/A |
| 20250528_Lys_Ac#11_unk3 | Y, Y; N/A |

*No M-16 Da variant of this fragment was detected

**Corresponding MS1 EICs** (EICs for 1335.4998 and 1001.3729 were too low, but signals can be identified by averaged mass spectra)

**Mass spectra** (averaged over scans)

M

M-16 Da

M-32 Da

**Pytheas results**

| **digested fragment information** | |
| --- | --- |
| **enzyme** | RNase T1 |
| **unmodified sequence** | AG |
| **modified sequence** | N/A |
| **position on *A. c* tRNA^Lys^UUU** | [43-44] |
| **5’ end** | OH |
| **3’ end** | OH |
| **charge state detected** | -1 |
| **theoretical m/z** | 611.1364 |
| **ppm used for EIC extraction** | 20 |
| **detection of MS1 signal in data files** | |
| 20250504_Lys_T1 | Y |
| 20250528_Lys_CNS | Y |
| 20250528_Lys_Ac#11_unk3 | Y |

**Corresponding MS1 EIC**

**Pytheas results: no MS2 data to be matched**

| **digested fragment information** | |
| --- | --- |
| **enzyme** | RNase T1 |
| **unmodified sequence** | GUCAAG |
| **modified sequence** | [m7G]D[m5C]AAG |
| **position on *A. c* tRNA^Lys^UUU** | [46-51] |
| **5’ end** | OH |
| **3’ end** | OH |
| **charge state detected** | -2, -3 |
| **theoretical m/z** | 962.671, 641.4447 |
| **ppm used for EIC extraction** | 20 |
| **detection of MS1 signal in data files** | |
| 20250504_Lys_T1 | Y |
| 20250528_Lys_CNS | Y |
| 20250528_Lys_Ac#11_unk3 | Y |

*Unmodified variants were not detected

**Corresponding MS1 EIC**

**Pytheas results**

| **digested fragment information** | |
| --- | --- |
| **enzyme** | RNase T1 |
| **unmodified sequence** | UUCG |
| **modified sequence** | [m5Um]PCG |
| **position on *A. c* tRNA^Lys^UUU** | [54-57] |
| **5’ end** | OH |
| **3’ end** | OH |
| **charge state detected** | -2 |
| **theoretical m/z** | 599.0840 (unmodified);  613.0996 (modified) |
| **ppm used for EIC extraction** | 20 |
| **detection of MS1 signal in data files** | |
| 20250504_Lys_T1 | Y, Y |
| 20250528_Lys_CNS | Y, Y |
| 20250528_Lys_Ac#11_unk3 | Y, Y |

*Signal for unmodified variant was weak.

**Corresponding MS1 EIC**

**Pytheas results**

| **digested fragment information** | |
| --- | --- |
| **enzyme** | RNase T1 |
| **unmodified sequence** | AG |
| **modified sequence** | [m1A]G |
| **position on *A. c* tRNA^Lys^UUU** | [58-59] |
| **5’ end** | OH |
| **3’ end** | OH |
| **charge state detected** | -1 |
| **theoretical m/z** | 625.152 |
| **ppm used for EIC extraction** | 20 |
| **detection of MS1 signal in data files** | |
| 20250504_Lys_T1 | Y |
| 20250528_Lys_CNS | Y |
| 20250528_Lys_Ac#11_unk3 | Y |

**Corresponding MS1 EIC**

**Pytheas results: Pytheas does not map 2-nt oligo**

| **digested fragment information** | |
| --- | --- |
| **enzyme** | RNase T1 |
| **unmodified sequence** | UCCCUUAUUG |
| **modified sequence** | N/A |
| **position on *A. c* tRNA^Lys^UUU** | [60-69] |
| **5’ end** | OH |
| **3’ end** | OH |
| **charge state detected** | -3 |
| **theoretical m/z** | 1018.1237 |
| **ppm used for EIC extraction** | 20 |
| **detection of MS1 signal in data files** | |
| 20250504_Lys_T1 | Y |
| 20250528_Lys_CNS | Y |
| 20250528_Lys_Ac#11_unk3 | Y |

**Corresponding MS1 EIC**

**Pytheas results**

| **digested fragment information** | |
| --- | --- |
| **enzyme** | RNase T1 |
| **unmodified sequence** | CG |
| **modified sequence** | N/A |
| **position on *A. c* tRNA^Lys^UUU** | [72-73] |
| **5’ end** | OH |
| **3’ end** | OH |
| **charge state detected** | -1 |
| **theoretical m/z** | 587.1251 |
| **ppm used for EIC extraction** | 20 |
| **detection of MS1 signal in data files** | |
| 20250504_Lys_T1 | Y |
| 20250528_Lys_CNS | Y |
| 20250528_Lys_Ac#11_unk3 | Y |

**Corresponding MS1 EIC**

**Pytheas results: no MS2 data to be matched**

| **digested fragment information** | |
| --- | --- |
| **enzyme** | RNase T1 |
| **unmodified sequence** | CCA |
| **modified sequence** | N/A |
| **position on *A. c* tRNA^Lys^UUU** | [74-76] |
| **5’ end** | OH |
| **3’ end** | OH |
| **charge state detected** | -2 |
| **theoretical m/z** | 437.5818 |
| **ppm used for EIC extraction** | 20 |
| **detection of MS1 signal in data files** | |
| 20250504_Lys_T1 | Y |
| 20250528_Lys_CNS | Y |
| 20250528_Lys_Ac#11_unk3 | Y |

*shorter tails were not detected

**Corresponding MS1 EIC**

**Pytheas results**
