## Supplementary material for "Sub-stoichiometric modifications of *Aplysia californica* tRNAs and tRNA fragments revealed by integrating intact and bottom-up mass spectrometry": Supplementary_material_6_IscS-TtcA.docx

| *Escherichia coli* (str. K-12 substr. MG1655) | | |
| --- | --- | --- |
| **protein name** | **protein accession (NCBI)** | **gene ID** |
| IscS | YP_026169.1 | 947004 |
| TtcA | NP_415860.1 | 948967 |
| *Aplysia californica* |  |  |
| **protein name** | **protein accession (NCBI)** | **gene ID** |
| IscS-TtcA | XP_005111426.3 | LOC101861551 |
| IscS/CsdA-like | XP_035824879.1 | LOC101846496 |

**Conserved domains on *A. californica* IscS-TtcA**

CsdA: cysteine desulfurase, equivalent to IscS

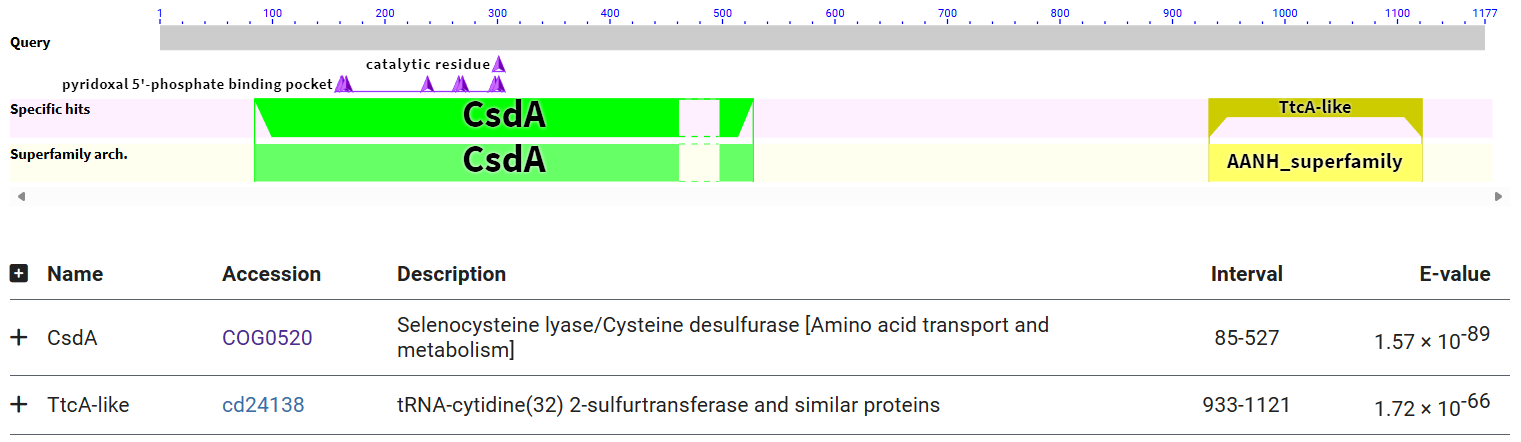

***A. californica* IscS-TtcA vs CsdA domain**

First line in alignment is query sequence; second line is a consensus sequence from multiple species, generated by NCBI.

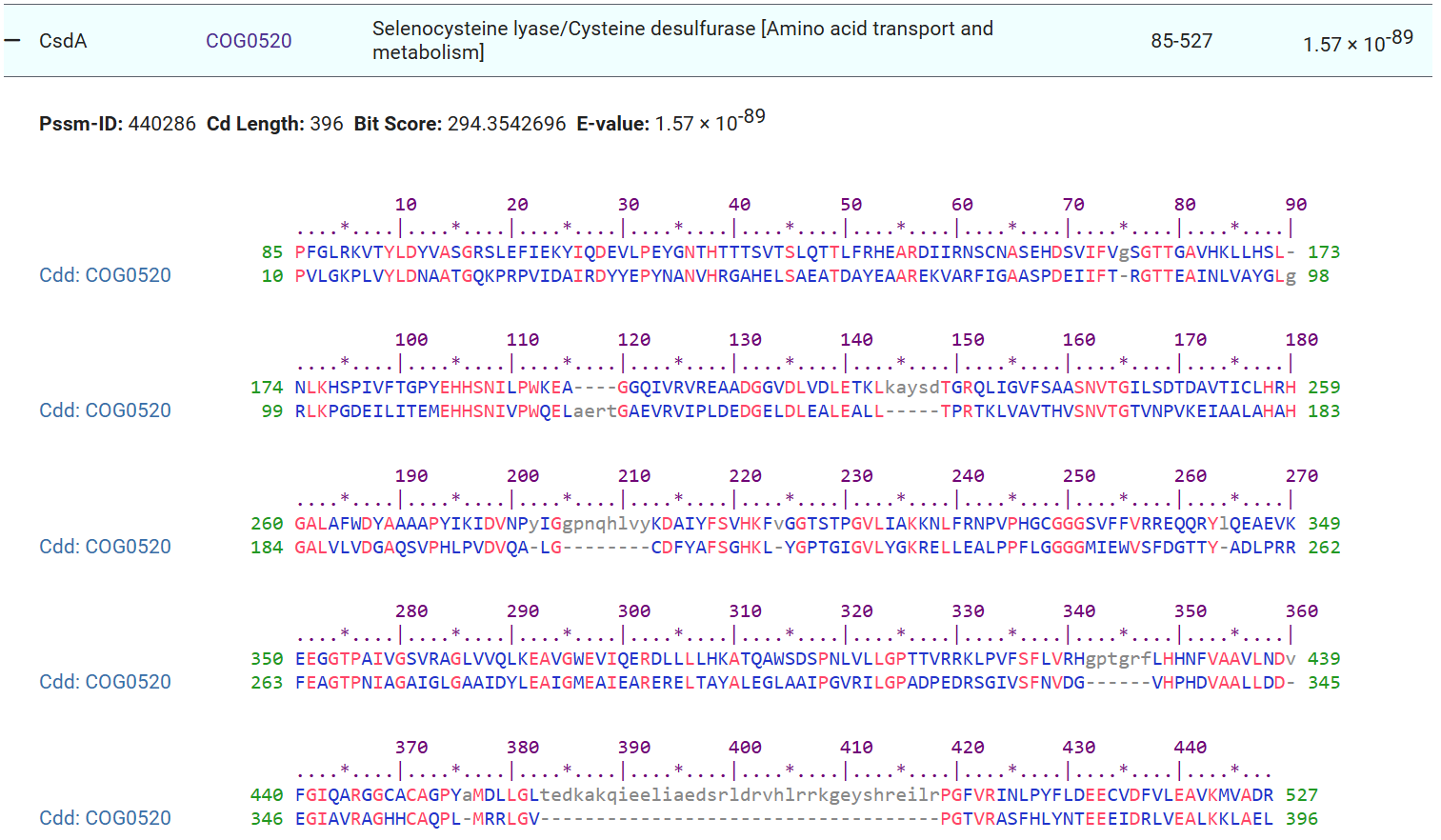

***A. californica* IscS-TtcA vs TtcA domain**

First line in alignment is query sequence; second line is a consensus sequence from multiple species.

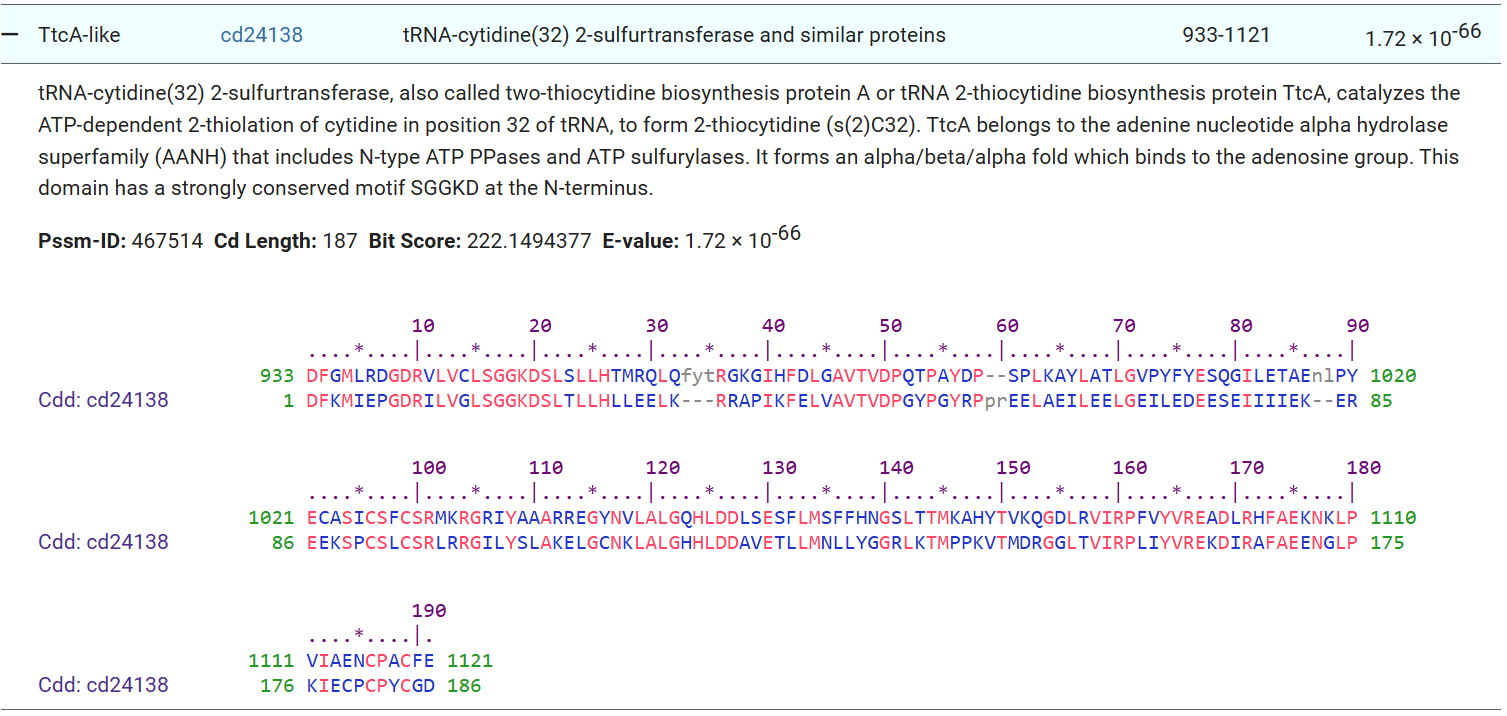

***E. coli IscS* vs *A. californica* IscS-TtcA**

Blastp search using *E. coli* IscS as the query did not find the same protein in *A. californica* (encoded by LOC101861551), but there is a IscS targeted to mitochondria (see below). The CsdA domain itself in *A. californica* IscS-TtcA is a little more divergent and cannot be aligned very well with *E. coli* IscS.

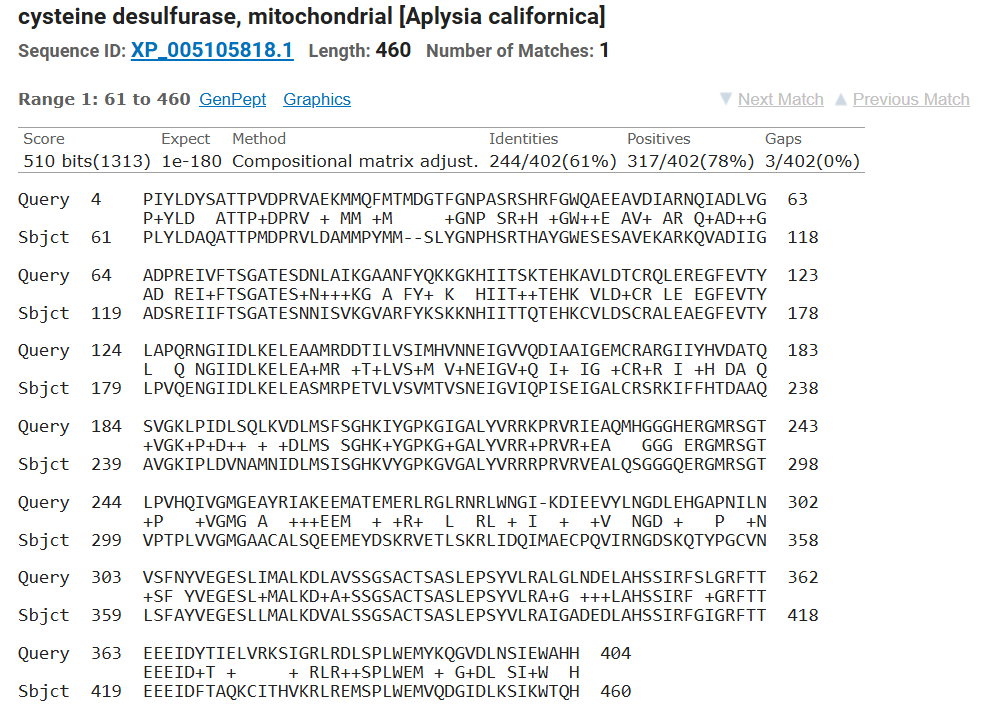

***E. coli* TtcA vs *A. californica* IscS-TtcA**

Blastp search using *E. coli* TtcA as the query found a hit in *A. californica* (encoded by LOC101861551), which is the fused IscS-TtcA protein.

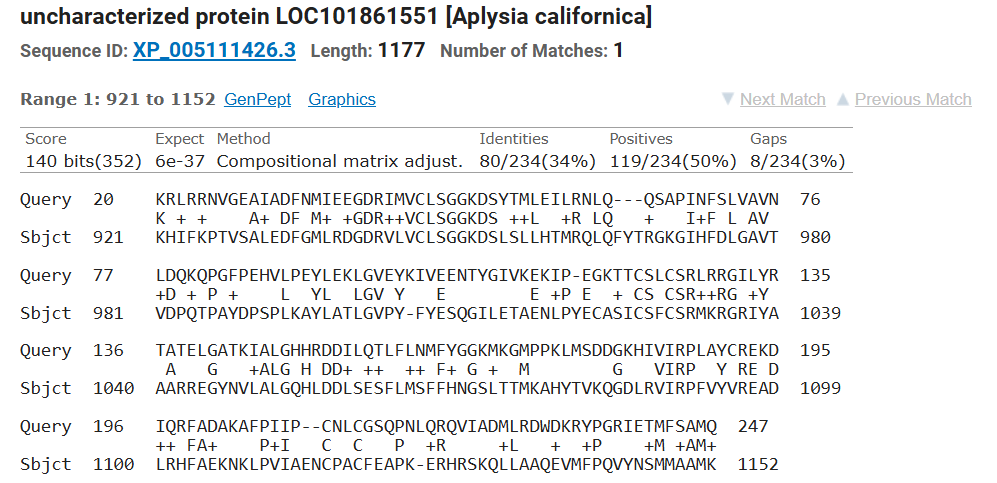

**Expression of IscS-TtcA as confirmed by RNA-seq data**

The gene LOC101861551 (IscS-TtcA) is indeed expressed in gills (lower panel) and hepatopancreas (middle panel), as seen by RNA-seq mapping data visualized by IGV. Whether it is expressed as a single transcript or as multiple needs further investigation.

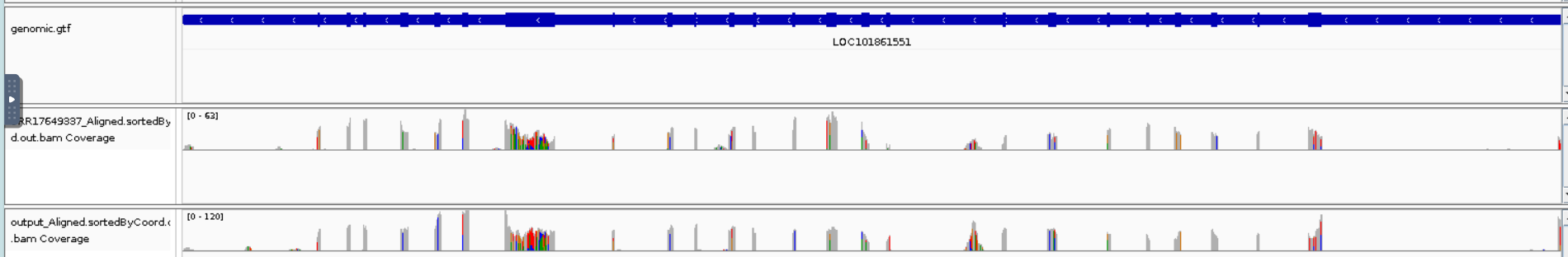

RNA-seq coverage can also be found on the gene’s page: <https://www.ncbi.nlm.nih.gov/gene/?term=LOC101861551>
