## Supplementary figures and images for "Sub-stoichiometric modifications of *Aplysia californica* tRNAs and tRNA fragments revealed by integrating intact and bottom-up mass spectrometry"

### Supplementary_material_4_A_californica_tRNAs-NJ-tree.pdf

Supplementary material 4. Neighbor-joining tree of Aplysia californica tRNA sequences

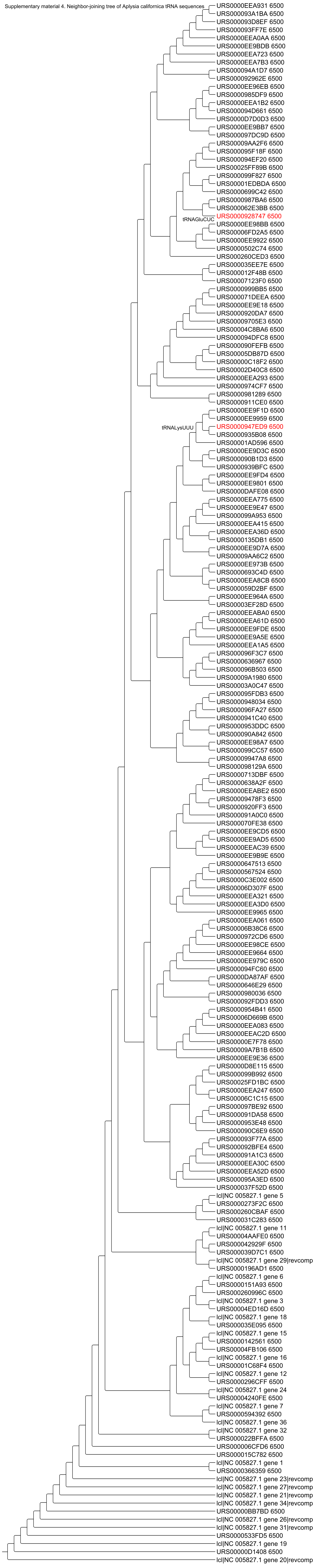

### Supplementary_material_5_A_californica_tRNAs_aligned.pdf

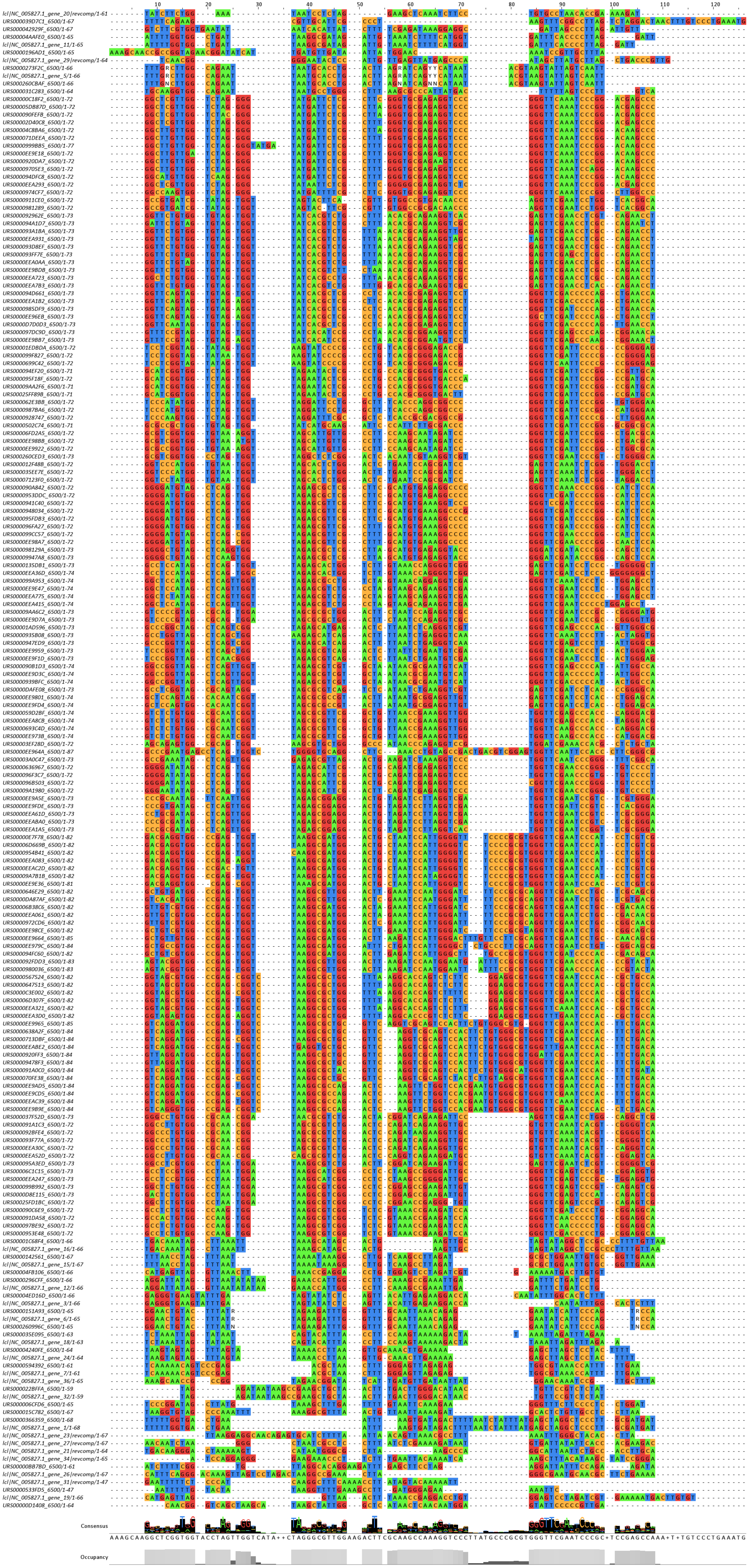
